# Chimeric Induced Cooperativity Opens the Design Space of Eukaryotic Gene Regulation

**DOI:** 10.64898/2026.08.28.747696

**Authors:** Yi Zhan, Zengli Li, Xinrui Li, Gengjiang Liu, Jinghui Xiong, Boyan Wei, Yanjun Yi, Ran Wang, Fangxin Wang, Bin Shao, ShuYi Zhang, Ye Chen

**Affiliations:** Key Laboratory of Quantitative Synthetic Biology, Shenzhen Institute of Synthetic Biology, Shenzhen Institutes of Advanced Technology, Chinese Academy of Sciences, Shenzhen, China; Colleges of Biological Sciences, China Agricultural University, Beijing, China; School of Pharmaceutical Sciences, Tsinghua University, Beijing, China; Advanced Research Institute of Multidisciplinary Science, Beijing Institute of Technology, Beijing, China; School of Science and Engineering, The Chinese University of Hong Kong, Shenzhen, China; University of Chinese Academy of Sciences, Beijing, China

**Keywords:** induced cooperativity, synthetic transcription factor, predictive gene regulation, thermodynamic modeling, multiplexed gene regulation

## Abstract

Predictive engineering of eukaryotic transcription is limited by the coupling of signal sensing, DNA binding, TF abundance and promoter output. Here we establish chimeric induced cooperativity (CIC), a modular architecture that separates LBD, DBD and AD functions and links them to promoters with tunable basal and maximal output. Module parameters can be recombined to predict new CIC-TF configurations and guide design before construction. Ligand-induced cooperativity reduces basal DNA occupancy while increasing induced occupancy, and effective DBDs combine low OFF-state activity with strong ON-state promoter occupancy rather than binding strength alone. Synthetic promoters independently control occupancy gain and output range. The same framework extends to repression and can be recalibrated with limited measurements in mammalian cells. In yeast, CIC-12 achieved a mean fold induction of 298-fold across 12 orthogonal sensors; an earlier CIC-10 chassis enabled model-guided optimization of an eight-gene vitamin B5 biosynthetic pathway. CIC establishes a programmable, model-guided design space for eukaryotic transcriptional control.

## Introduction

Cells do not simply turn genes on and off; they compute signals through genetically encoded regulatory networks. In natural systems, this computation often emerges from cooperative assembly of transcription factors, cofactors and chromatin-associated machinery at regulatory DNA, allowing multiple inputs to be integrated before transcriptional output is produced^1–7^. Synthetic biology seeks to reproduce these interactions as quantitative transfer functions^8–11^: input–output responses with specified basal leakage, maximal output, dynamic range, sensitivity, burden and orthogonality. These transfer functions determine whether a regulator can support a single inducible gene, a multigene metabolic pathway, a therapeutic circuit or a larger cellular program. A practical challenge is how to use measurements from a limited number of regulatory components to predict behavior across a much larger design space. Addressing this challenge would reduce the need to test each combination as a complete construct.

One limitation is that cooperativity is difficult to make modular. Classical allosteric regulators, including Tet-, Lac- and steroid receptor-derived systems^12–18^, have become foundational tools because they couple ligand state to DNA binding or regulatory output. However, in these Type I architectures, signal sensing and DNA recognition are encoded within the same structural framework. Domain exchange, affinity tuning or logic inversion can be achieved in selected cases, but typically requires extensive engineering, mutagenesis or directed evolution^19–23^ because DNA binding, ligand response and conformational switching are not independently adjustable. Moreover, many natural allosteric systems are optimized for ligand-induced release, repression or intramolecular switching rather than modular ligand-induced binding and activation, restricting the accessible design space for synthetic eukaryotic transcriptional control.

A second strategy has been to build chimeric or split regulators by coupling a strong DNA-binding module to a ligand-controlled interaction or effector-recruitment module^24–26^. Chemically induced dimerization systems, dCas-based recruitment platforms, zinc-finger and TALE effectors, and related Type II designs^27–41^ substantially improve modularity because DNA-binding, sensing and effector domains can be exchanged more independently. However, this modularity usually comes at the cost of signal-induced cooperative DNA occupancy. The DNA-binding component is often present at its target site before induction, while the ligand primarily recruits or activates an effector after DNA binding has occurred. Previous studies have shown that such persistent occupancy can increase basal expression, create genomic roadblocks, impose host burden and compress dynamic range, particularly when multiple regulators are expressed simultaneously. Strategies such as attenuating DNA affinity, lowering TF abundance, scaffolding components or adding generic multimerization domains^36–38, 42^ can mitigate specific failure modes, but they often trade off output, targeting efficiency^43^, composability or quantitative control rather than defining a general transfer function design rule. Type I systems therefore provide cooperativity with limited modularity, whereas Type II systems provide modularity without induced cooperative DNA occupancy. A Type III architecture should make DNA occupancy conditional on ligand-dependent formation of the active TF pool, limiting promoter occupancy in the OFF state while allowing high occupancy after induction.

Promoter context introduces an additional source of variation: operator placement, core promoter syntax, spacer sequence, nucleosome organization, cryptic regulatory sites, interactions with endogenous host factors and sequence repetition can all reshape the observed response^44–54^. Operator insertion into native promoters, hybrid promoter construction and synthetic promoter libraries have produced many useful parts^55–61^, but their performance is often optimized as sequence activity rather than parameterized as part of a TF–promoter transfer function. Native or semi-synthetic promoters can also carry context dependence or recombination risk, limiting scalability in large genetic systems. Existing promoter models and activity maps support descriptive analysis and local optimization, but they rarely define the feasible performance boundary of a complete regulatory architecture. Without an integrated TF–promoter transfer function model, each new DBD–LBD–promoter combination becomes a separate empirical search.

The same combinatorial problem limits model-guided design. If each regulator must be characterized directly in part or sequence space, the search expands across DBD identity, LBD identity, activation domain, TF expression level, operator arrangement and promoter context. Even large screens therefore sample only a small fraction of the possible transfer functions. Quantitative frameworks have begun to make such parameterized design practical in bacterial hosts by updating host-specific transcriptional models from limited measurements^62–64^. An alternative is to estimate reusable biophysical parameters from a limited set of component measurements and use them to calculate responses for unmeasured combinations. These parameters could be used to calculate design landscapes spanning DBD identity, LBD identity, TF expression and promoter architecture. The resulting landscapes could identify feasible regions and Pareto frontiers and help select TF–promoter designs that meet the performance criteria of a given application^10, 11, 65, 66^.

Here we introduce and experimentally validate **chimeric induced cooperativity** (CIC) as an architecture that combines modular domain exchange with signal-induced cooperative DNA occupancy. The architecture allows LBD-dependent active pool formation, DBD–operator coupling, TF dosage, activation strength and promoter output capacity to be measured separately. These parameters can then be recombined to calculate responses for TF–promoter configurations that were not measured as complete constructs.

We first characterize cis- and trans-regulatory parameters in yeast and test whether they can predict additional CIC-TF configurations and guide sensor design. We then examine whether the same occupancy model can be used for repression and, after host-specific calibration, in mammalian cells. Finally, we apply the framework to the design of multi-sensor panels and the optimization of a multi-gene pathway.

## Results

### Induced cooperativity separates the determinants of CIC-TF response

We first compared three inducible regulatory topologies: Type I allosteric regulators, Type II ligand-induced recruitment systems and Type III ligand-induced cooperative DNA-binding systems (Fig. 1A). These architectures differ in when the DNA-binding module occupies the promoter, whether ligand input changes the effective DNA-binding-competent TF pool, and whether the DNA-binding and ligand-sensing functions can be independently exchanged.

**Fig. 1.**
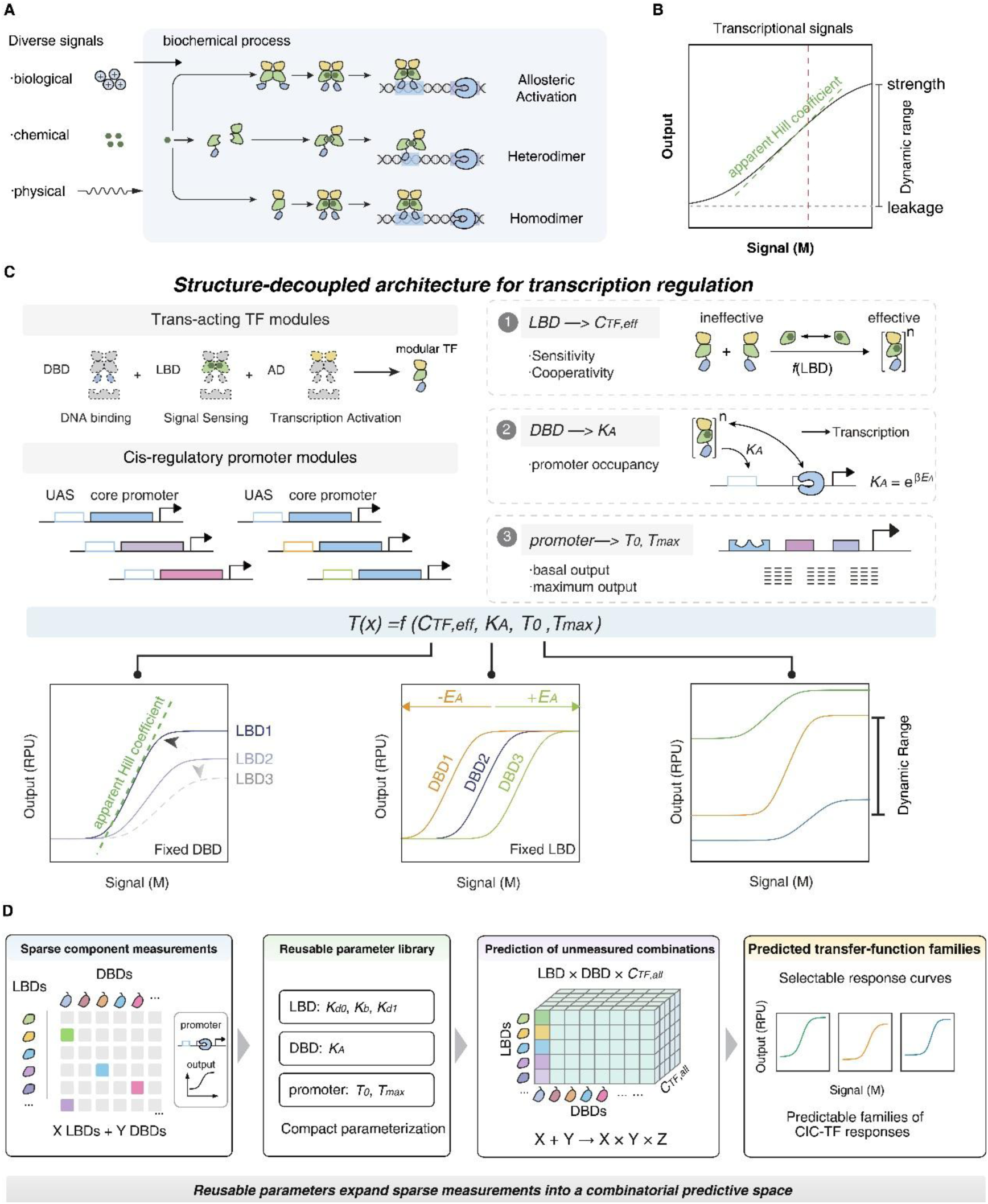
Induced cooperativity separates the parameters that define the eukaryotic transcriptional design space. **(A)** Schematic comparison of inducible transcriptional regulation architectures that convert diverse biological, chemical, or physical inputs into transcriptional outputs. Representative architectures include allosteric regulators, ligand-induced heterodimerization systems, and ligand-induced homodimerization systems. **(B)** Key quantitative features of a transcriptional response. Basal leakiness, output strength, dynamic range, sensitivity, and cooperativity define the performance of an inducible transcriptional signal. **(C)** Modular CIC architecture for transcriptional regulation. Trans-acting TF modules are separated into DNA-binding domains (DBDs), ligand-binding domains (LBDs), and activation domains (ADs), while cis-regulatory promoter modules define the input and output window. In this framework, LBD properties determine the apparent DNA-binding-competent TF pool (effective TF, *C_TF,eff_*), DBD affinity (*K_A_*) controls promoter occupancy, and promoter architecture sets basal and maximal output (*T_0_*, *T_max_*). Together, these components define the transfer function T(x) = f(*C_TF,eff_*, *K_A_*, *T_0_*, *T_max_*). Bottom schematics illustrate how LBD properties, DBD affinity, and promoter output capacity independently tune response shape, signal threshold, and dynamic range. **(D)** Measurements of individual modules define the broader CIC-TF design space. Characterization of individual LBDs, DBDs and promoter output windows yields a compact parameter library, including LBD-associated parameters (*K_d0_*, *K_b_*, *K_d1_*), DBD affinity (*K_A_*), and promoter parameters (*T_0_*, *T_max_*). Measurements of X LBDs and Y DBDs can be combined to calculate responses across a design space spanning LBD identity, DBD identity and TF abundance *C_TF,all_*, thereby defining families of CIC-TF transfer functions without exhaustive combinatorial screening.

To compare these designs in transfer function space, we defined the key performance features of an inducible transcriptional response: basal leakage, output strength, dynamic range, sensitivity and cooperativity (Fig. 1B). Architectures lacking signal-dependent cooperative DNA occupancy can be modular, but they often place a DNA-binding module at the promoter before induction. This constitutive promoter occupancy creates a tradeoff: reducing DBD affinity or TF abundance can lower leakage and burden, but also reduces induced output and can narrow the useful dynamic range. Allosteric regulators can show strong input-dependent behavior, but their DBD and LBD functions are often structurally coupled, making it difficult to independently retune DNA affinity, ligand response and regulatory logic. A modular eukaryotic regulator would therefore need to combine chimeric domain exchange and ligand-induced cooperative DNA occupancy.

We therefore built the CIC architecture around a separable transfer function model. In this model, the LBD controls formation of an apparent cooperative TF pool capable of DNA binding, denoted *C_TF,eff_*; the total TF abundance *C_TF,all_* sets the operating point; the DBD–operator affinity *K_A_* converts *C_TF,eff_* into promoter occupancy; and the promoter defines the basal and maximal output window, *T_0_* and *T_max_* (Fig. 1C and Supplementary Fig. 1). For modules derived from quorum-sensing systems, *C_TF,eff_* can be interpreted through ligand binding and induced dimerization parameters. For chemically diverse modules, including those derived from nuclear receptors, we treat *C_TF,eff_* operationally as an apparent active pool that captures all upstream processes required to produce DNA-binding-competent TF.

The model represents the CIC design space using parameters that describe LBD activity, DBD affinity, TF abundance and promoter output capacity (Fig. 1C,D). Measurements of individual components can therefore be combined to calculate transfer functions across this design space (Fig. 1D). LBD parameters determine a family of response curves, DBD affinity selects the corresponding occupancy regime, and the promoter sets the available output range. Recombining these parameters allows transfer functions to be calculated for TF–promoter constructs that were not measured directly.

### Synthetic promoter reconstruction separates input gain from output window capacity

Quantitative prediction also requires a defined relationship between TF occupancy and promoter output. We therefore designed a synthetic promoter architecture in which input recognition and output capacity are separable. The promoter was divided into an upstream input module containing operator sites, a core promoter module containing transcription initiation elements, and spacer sequences that separate these regulatory elements and tune basal and maximal output (Fig. 2A). In this architecture, operator identity and copy number determine how the promoter receives CIC-TF input, whereas spacer and core promoter syntax define the output window represented by *T_0_* and *T_max_*.

**Figure 2.**
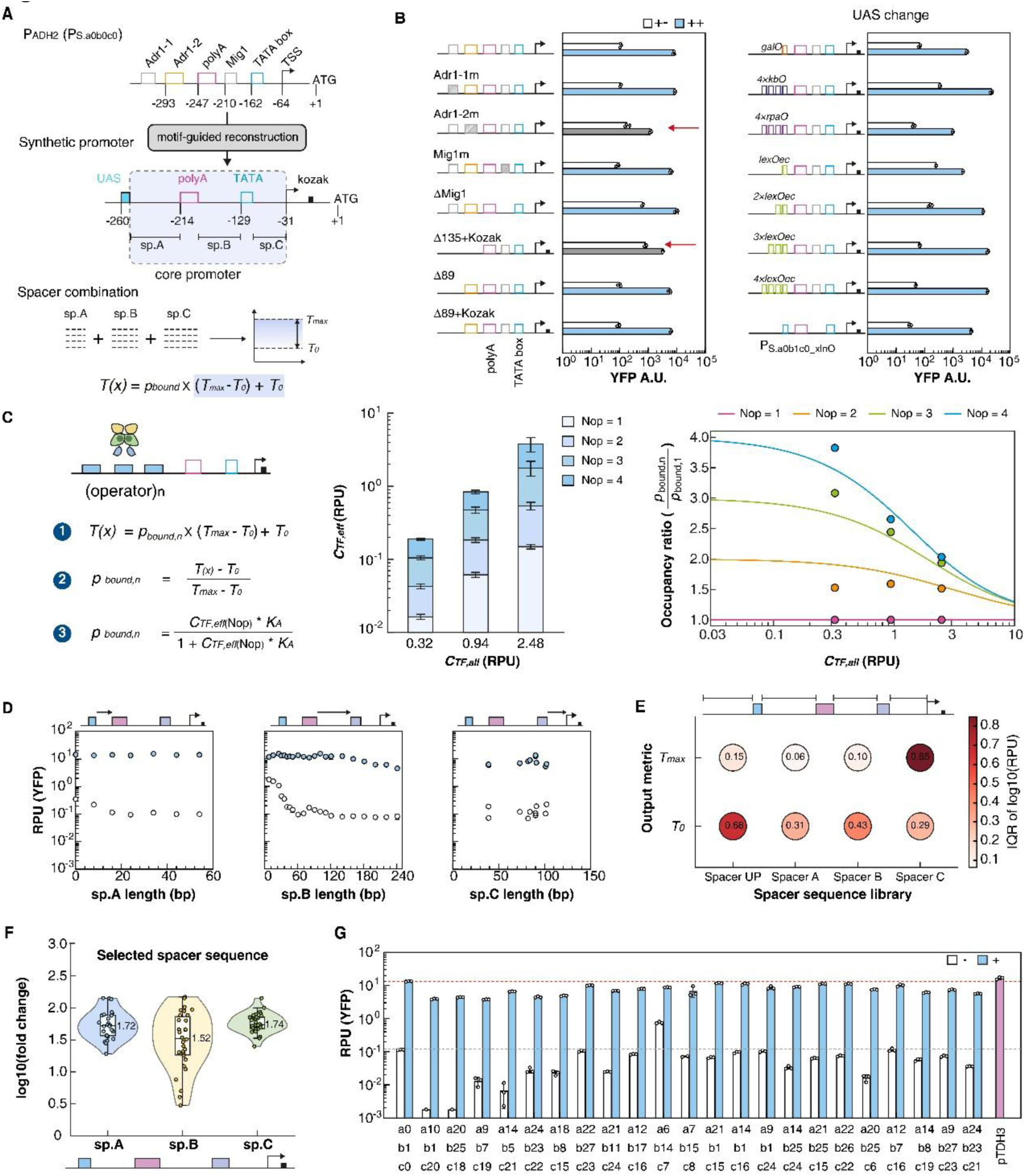
A limited set of promoter measurements defines reusable parameters for input gain and output range. **(A)** Workflow for resolving native promoter architecture and constructing a modular synthetic promoter. Functional elements in pADH2 were mapped, and spacer length and sequence were varied to tune the promoter output window. **(B)** Core and synthetic inducible promoter architectures derived from the native S. cerevisiae pADH2 promoter. Functional motifs and regulatory elements are identified through mutations or deletions. A Kozak sequence is positioned immediately downstream of the TSS. Δ89 and Δ89+Kozak differ in the Kozak element, indicated by the black box. The native Adr1 binding site is placed with different types and numbers of operators and can respond to different signals. Red arrows indicate constructs in which deletion or mutation of the indicated native promoter element substantially reduced the regulatory range. In B, ‘+−’ denotes induction of TF expression without ligand, whereas ‘++’ denotes induction of TF expression together with the cognate ligand. **(C)** Quantitative model and experimental analysis of operator copy-number effects. Promoter output was modeled as T(x) = *p_bound,n_* × (*T_max_* - *T_0_*) + *T_0_*, where *p_bound,n_* depends on effective TF concentration, DBD affinity, and operator copy number. Operator copy number controls occupancy gain; its benefit is largest when TF input is limiting and diminishes near promoter saturation. Experiments were performed using LexA*_mm_*_115_-DHBR_282-595_- VP16 as the TF, with TF input levels of 0.32, 0.94, and 2.48 RPU and DHB fixed at 100 μM. **(D)** Spacer-length optimization for sp.A, sp.B and sp.C, showing the effects of spacer length on basal and induced promoter output. The selected architecture contained 24-bp sp.A, 61-bp sp.B and 90- bp sp.C segments. **(E)** Bubble summary of spacer-sequence effects on basal (*T_0_*) and maximal (*T_max_*) promoter output. For each spacer library, sequence-dependent variability was calculated as IQR(log_10_ X) = Q75(log_10_ X) − Q25(log_10_ X), where X is the variant-level *T_0_* or *T_max_* in RPU. Values from the main and supplementary screens were pooled; main-dataset replicates were summarized by their median, and Spacer C included the C1, C2, C3, and full-SpacerC libraries. Bubble color indicates the magnitude of the sequence-dependent effect. **(F)** Fold induction distributions of screened effective spacer sequences for sp.A, sp.B and sp.C (n = 26, 30 and 29 variants, respectively). **(G)** Synthetic promoters generated by assembling selected spacer combinations with reduced sequence similarity. Basal and induced YFP outputs were measured. The gray and orange dashed horizontal lines indicate the mean basal and induced outputs, respectively, of the P*_S. a0b1c0_* promoter, whereas the magenta bar indicates the output of the constitutive pTDH3 reference promoter. In panels D and G, blue-filled circles and bars indicate the presence of the cognate ligand, whereas unfilled circles and bars indicate TF expression in the absence of ligand and therefore represent basal leakage.

To construct a promoter scaffold with separable parameters, we used the native yeast pADH2 promoter^67–71^ as a motif-guided reconstruction template. Systematic mutation of its regulatory motifs showed that the Adr1-2 binding site was the dominant native input element under our assay conditions, whereas other candidate motifs contributed little to activation in this context (Fig. 2B). Replacing this input site with heterologous operator sequences, including NF-κB^72^, rpaO, LexO_rec_ and galO sites, rewired promoter responsiveness to synthetic TF inputs (Fig. 2B). Thus, the promoter input module could be exchanged while preserving a common output scaffold, enabling operator identity to be treated as a programmable promoter input parameter.

We next quantified how operator copy number changes promoter behavior within the CIC transfer function model. By inverting measured promoter responses into an equivalent *C_TF,eff_*, we quantified the gain produced by different operator configurations while keeping operator identity constant (Fig. 2C). This analysis showed that operator copy number is best interpreted as an occupancy gain parameter rather than as an intrinsic promoter strength parameter. At low *C_TF,all_*, additional operators increased equivalent *C_TF,eff_*, indicating improved promoter occupancy when TF input was limiting. At higher *C_TF,all_*, this gain diminished as promoter occupancy approached saturation, such that one operator could be sufficient to drive activation close to the promoter maximum at the matched CIC-TF operating point (Fig. 2C). Operator number could therefore be used as a tunable determinant of input gain in the promoter model.

We then optimized the promoter output window by systematically changing spacer length and sequence. Length tuning identified spacer architectures that minimized basal expression while preserving high induced output, including optimized spacer A, spacer B and spacer C configurations (Fig. 2D). To separate length effects from sequence effects, we screened randomized spacer libraries. Spacer UP and spacer B primarily affected basal leakage *T_0_*, whereas spacer C strongly influenced maximal output *T_max_* (Fig. 2E,F and Supplementary Fig. 2). Nucleosome occupancy analysis and models based on sequence features^47, 49, 50, 52, 53^ provided possible explanations for these effects on the output window. Chromatin features associated with spacer B correlated with basal expression, whereas sequence variation in spacer C correlated with induced output (Supplementary Figs. 3 and 4). These analyses do not require treating the promoter as a black box: spacer and core promoter syntax can be mapped onto operational *T_0_* and *T_max_* parameters (Fig. 2D–F).

**Figure 3.**
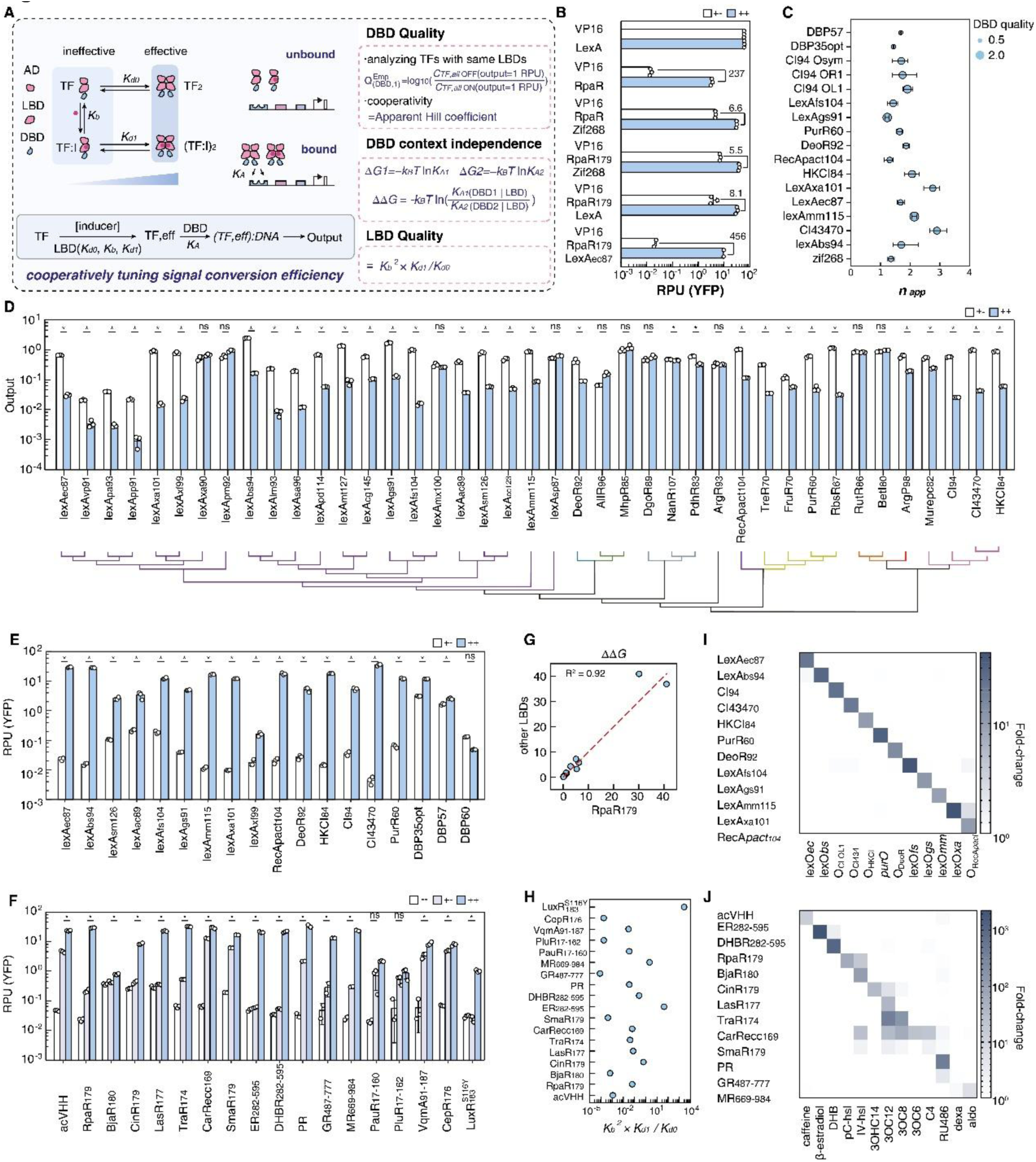
D**B**D **and LBD measurements define reusable parameters for CIC-TF design. (A)** General scheme of CIC activator function and evaluation of module performance. A monomeric CIC activator can undergo basal or ligand-dependent dimerization to generate a DNA-binding- competent TF pool, which binds its cognate operator and activates transcription. DBD modules were evaluated along three complementary dimensions. First, DBD signal-transmission quality was defined as the horizontal separation between the OFF and ON TF input curves at an output of 1 RPU; larger values indicate a greater separation between the two states. 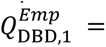 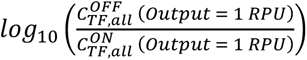, Second, apparent cooperativity was quantified by the apparent Hill coefficient of the fitted TF input response. Third, DBD context independence was evaluated by testing whether the relative DBD–operator binding free energy, ΔΔ*G* = 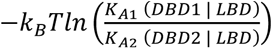, was conserved across different LBD backgrounds. LBD modules were independently ranked using 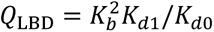, which quantifies the ligand-dependent gain in the DNA-binding-competent TF pool relative to basal active pool formation. **(B)** CIC architecture proof-of-principle in *S. cerevisiae*. Activator designs with different architecture-level cooperativity were compared, including non-cooperative, partially cooperative, and cooperative CIC configurations. Leakiness was measured in the absence of pC-HSL, and maximal induced output was measured with 100 μM pC-HSL. This panel validates the requirement for ligand-induced cooperative regulation at the architecture level. **(C)** Empirical ranking of DBD signal-transmission quality and apparent cooperativity. Each row represents one DBD. Bubble position along the x axis denotes the apparent Hill coefficient *n_app_*, estimated by power-law fitting in the low-output regime, whereas bubble area represents the corresponding DBD quality score (*Q*^emp^ , as defined in A). Horizontal grey-blue error bars indicate 95% confidence intervals obtained by bootstrap resampling. Larger *Q*^emp^ values indicate greater ligand-dependent separation between the OFF and ON transfer functions, whereas *n_app_* values indicate stronger apparent cooperativity. Zif268 was included as a non-cooperative DNA-binding control. DBP35opt and DBP57 were characterized using 1 mM pC-HSL, whereas the remaining DBD–RpaR_179_ constructs were evaluated using 100 μM pC-HSL. **(D)** Screening of DNA-binding domains (DBDs) in *E. coli* and their phylogenetic relationships, shown as a neighbor-joining tree. DBDs were evaluated in the DBD-RpaR_179_ repressor architecture in the absence (0 μM; open bars) or presence (20 μM; blue bars) of pC-HSL. Each point represents an independent biological replicate (n = 3 per condition), and bars represent arithmetic means of untransformed RPU. For each DBD, a one-sided Welch’s t-test on untransformed RPU values tested the a priori hypothesis that 20 μM pC-HSL decreases output relative to 0 μM pC-HSL. P values from 41 DBD-wise comparisons were adjusted using the Benjamini-Hochberg procedure. Statistical significance is indicated as follows: *q < 0.05; ns, q ≥ 0.05. **(E, F)** Screening of functional DBDs and LBDs in *S. cerevisiae*. DBD activity was evaluated using the DBD–RpaR_179_–VP16 activator architecture (E), whereas LBD activity was evaluated using the LexAbs94–LBD–VP16 architecture (F). RPU was measured under uninduced and fully induced ligand conditions. For DBDs, log_10_-transformed RPU values were compared between + and − inducer conditions using one-sided unpaired Welch’s *t*-tests (pre-specified alternative: RPU (+inducer) > RPU (-inducer)); P values from 18 comparisons were adjusted by the Benjamini– Hochberg procedure. For LBDs, RPU was measured without TF expression or ligand (--), with TF expression but without ligand (+-), and with TF expression plus ligand at its fully inducing concentration (++). Ligand responsiveness was assessed by comparing ++ and +- using one-sided Welch’s *t*-tests on log10-transformed RPU values (pre-specified alternative: ++ > +-), followed by Benjamini–Hochberg adjustment across 18 comparisons. Bars show arithmetic means ± sample SD; circles indicate independent biological replicates. *, q < 0.05; ns, q ≥ 0.05. **(G)** DBD context independence. Agreement of ΔΔ*G* across LBD backgrounds indicates context-independent DBD– operator coupling. **(H)** LBD quality ranking. The quality of each LBD was quantified as the inducer- dependent gain in DNA-binding-competent TF pool relative to its basal activity. Briefly, measured RPU values were normalized to promoter occupancy and converted to promoter-binding odds (Z=p/(1-p)). LBD quality was then calculated as *Q_LBD_* = *K_b_*_2_ × *K_d1_*/*K_d0_* according to the apparent cooperative transfer model described in Supplementary Note 4. Higher values indicate stronger ligand-dependent activation with lower basal active pool formation. (**I)** Combinatorial pairing of selected DBDs and operators identified in *S. cerevisiae* was tested in *E. coli*. Heatmap color indicates the fold change in RPU before and after induction with 100 μM IPTG. Values represent the mean of each tested combination. The response of DBD–operator combinations to 20 μM pC- HSL was further evaluated under the *E. coli* repressor architecture. No substantial crosstalk was observed among the tested DBD–operator pairs. **(J)** Orthogonality profiling of ligand-binding domains across a panel of chemical inducers. Heatmap showing the fold change of each ligand- binding domain (LBD) in response to cognate and non-cognate chemical inducers. All CIC-TFs were tested at a fixed TF input level of 1.24 RPU. Rows indicate LBD modules, and columns indicate inducers tested at the concentrations shown above the matrix. Values represent fold change. The inducer panel included 1 μM caffeine, 1 μM β-estradiol, 10 μM DHB, 100 μM pC-HSL, 100 μM IV- HSL, 1 μM 3OHC14-HSL, 1 μM 3OC12-HSL, 100 μM 3OC8-HSL, 1 μM 3OC6-HSL, 100 μM C4-HSL, 10 μM RU486, 100 μM dexamethasone, and 1 μM aldosterone. Unless otherwise indicated, data are shown as mean ± SD from three independent cultures (n = 3). Heatmap values represent log_10_(fold change) calculated from replicate means.

**Figure 4.**
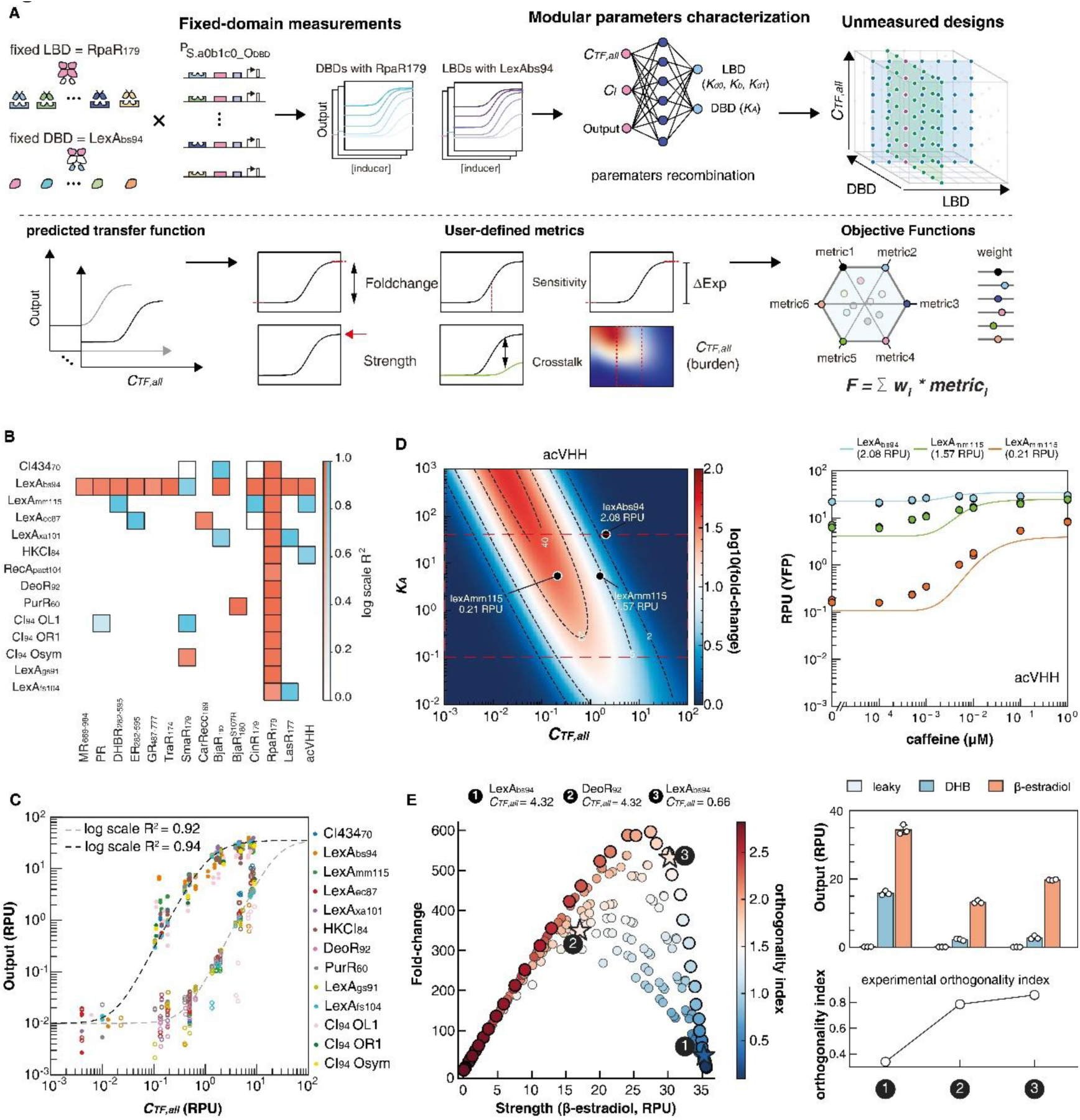
M**e**asurements **of individual modules predict additional CIC-TF combinations and guide prospective design. (A)** Workflow for parameter estimation and CIC-TF design. DBD and LBD modules were characterized in fixed-domain backgrounds, using RpaR_179_ for DBD screening and LexA_bs94_ for LBD screening. Model fitting of these induction datasets yielded domain-specific LBD parameters (*K_d0_*, *K_b_*, *K_d1_*), DBD affinity (*K_A_*), and TF input-output relationships, which were further evaluated using additional DBD–LBD combinations for cross-validation. The fitted parameters are recombined to calculate responses across the design space defined by LBD identity, DBD identity and total TF abundance *C_TF,all_*. Predicted transfer functions can then be scored using specified performance metrics, including output, fold change, ΔExp, burden, sensitivity, and crosstalk, to identify Pareto-optimal sensor designs. **(B)** Transferability of DBD and LBD parameters across CIC- TF combinations. Heatmap cells report the coefficient of determination (R²), calculated from log- transformed model-predicted and experimentally measured reporter outputs, for each tested DBD-LBD sensor. Orange fills denote sensors used for parameter fitting, whereas blue fills denote sensors withheld from the corresponding parameter estimation and used for cross-validation. Four sensors in the cross-validation set were drawn from the 12s set. For these four sensors, CIC- TF expression was calculated rather than directly measured. Their prediction-to-experiment discrepancy therefore contains two components: error in the model prediction and error arising from the difference between the calculated and actual CIC-TF expression levels. **(C)** Master-curve validation for CIC-TFs carrying the RpaR_179_ LBD. Experimental dose-response data from DBD– RpaR_179_–VP16 variants were normalized to a common reference sensor using the deformation- based correction described in Supplementary Note 8.2, thereby accounting for DBD-specific *K_A_* differences. Each color represents a different DBD. Open circles denote inducer-free measurements (*C_I_* = 0 μM pC-HSL), and filled circles denote induced measurements (*C_I_* = 100 μM pC-HSL). The two dashed lines indicate model-predicted master curves for the uninduced and induced states, respectively, plotted as output versus total TF abundance *C_TF,all_*. Log scale *R^2^* values indicate the agreement between corrected experimental measurements and the predicted master curves for each inducer condition. **(D)** Model-guided optimization of acVHH-based CIC-TFs across total TF abundance *C_TF,all_* and DBD affinity *K_A_*. The heatmap shows predicted fold change, and marked points indicate experimentally tested designs. **(E)** Model-guided Pareto optimization of ER-based CIC-TFs to reduce DHB crosstalk. Left, predicted design landscape across DBD identities and *C_TF,all_* levels. Each point represents one simulated design, with predicted strength under maximal β-estradiol induction on the x-axis and fold change on the y-axis. Point color indicates the predicted orthogonality index, calculated as log_10_(ΔExp β-estradiol/ΔExp DHB). Enlarged points denote Pareto-optimal designs, and stars mark three experimentally tested designs: 1, LexA_bs94_- ER_282-595_-VP16 at *C_TF,all_* = 4.32; 2, DeoR_92_-ER_282-595_-VP16 at *C_TF,all_* = 4.32; and 3, LexA_bs94_-ER_282-595_-VP16 at *C_TF,all_* = 0.66. Right, measured basal, DHB-induced, and β-estradiol-induced outputs for the selected designs, with the black line showing the measured orthogonality index.

Finally, we assembled 23 synthetic promoters from characterized operator, spacer and core promoter modules (Fig. 2G). These assembled promoters retained high inducibility, with an average induction exceeding 200-fold, while providing a set of promoter output windows for matching different CIC-TF operating points. Because the promoters were built by motif-guided reconstruction and modular assembly^51, 55, 58, 59, 61^ rather than by unstructured insertion into native promoter contexts, they also reduced reliance on native regulatory logic and repeated promoter sequences. The resulting promoter series provided independent control over input gain and the output window in subsequent CIC-TF analyses.

### DBD and LBD parameters are transferable across CIC-TF combinations

We next characterized how DBD and LBD identity affected ligand-dependent cooperative promoter occupancy (Fig. 3A). A CIC-TF contains three separable modules: a DBD that determines operator recognition and occupancy coupling, an LBD that controls ligand-dependent formation of the active TF pool, and an AD that recruits transcriptional activation machinery. This chimeric structure allows these functions to be exchanged independently, but high performance requires that the DBD transmit the cooperative state generated by the LBD into DNA occupancy.

We first tested this architecture using a set of proof-of-principle designs in yeast. A TF containing a non-cooperative monomeric DBD produced high basal activation because a single DNA-binding monomer could occupy the promoter before ligand input. In contrast, coupling an inducible dimerization module to a dimeric DNA-binding scaffold produced ligand-dependent activation with reduced basal output (Fig. 3B). The LexAec87–RpaR179 fusion served as a scaffold for testing the architecture; it was not intended to identify LexAec87 as the optimal DBD. This experiment established the qualitative requirement for CIC regulation: the DBD must couple promoter occupancy to ligand-induced cooperative assembly, rather than bind DNA productively in the basal state.

We then asked a separate quantitative question: which DBDs best transmit ligand-induced cooperativity into transcriptional output? To build a broad DBD library, we screened 41 prokaryotic repressor-derived DBDs in an E. coli repression assay and identified 31 domains with detectable function (Fig. 3D). From this set, we selected DBDs representing diverse half-site architectures and orthogonality across families, together with three designed DNA-binding proteins^73^, for characterization in the yeast CIC activation context (Supplementary Fig. 5). The E. coli assay provided a rapid filter for DNA-binding and repression function, but yeast activation revealed that repression strength, static binding capability and CIC performance are not equivalent.

Quantitative CIC characterization showed that DBDs differ in two separable properties: cooperative behavior and signal-transmission quality (Fig. 3C,E). Some DBDs supported strong induced output but also showed elevated basal activity or weak apparent cooperativity; others remained silent in the OFF state but failed to generate sufficient induced occupancy. DBD performance therefore depended on both OFF-state activity and ON-state occupancy rather than on binding strength alone. Quantitatively, a CIC-compatible DBD should keep *K_A_*·*C_TF,eff,OFF_* low enough to suppress basal promoter occupancy while placing *K_A_*·*C_TF,eff,ON_* in a responsive regime after induction; excessively weak or excessively strong coupling can both reduce transfer function performance (Fig. 3C,E). Static affinity alone is therefore insufficient to evaluate designed DNA-binding proteins in the CIC context. Phylogenetic and family-level analysis suggested that wHTH/LexA-like and CI-family scaffolds are particularly useful starting points for such DBD engineering (Fig. 3D,E).

In parallel, we assembled a chemically diverse LBD library responsive to bacterial quorum sensing molecules and mammalian hormone-like ligands^74–84^ (Fig. 3F and Supplementary Fig. 6). We evaluated these LBDs by basal activity, induced output and apparent LBD induction quality, and selected modules with strong ligand responses for the CIC platform (Fig. 3H,J). Because this LBD set includes both simple inducible dimerization modules and domains derived from nuclear receptors, we interpret LBD performance through the apparent active pool model rather than assuming a single microscopic mechanism for all modules. We also screened activation domains and selected VP16 as the default AD for subsequent CIC-TF construction because it provided strong ligand-dependent activation in the tested architecture (Supplementary Fig. 7).

To test whether DBD properties remain modular across LBD contexts, we compared relative DBD- associated binding energies when the same DBDs were fused to different LBDs. The resulting relationships were strongly conserved, indicating that apparent DBD–operator coupling is largely retained across LBD backgrounds (Fig. 3G). We then confirmed the orthogonality of the selected DBD/operator pairs in yeast, establishing a reusable set of DNA recognition modules for multi- channel regulation^85^ (Fig. 3I). Finally, we compared architectures that differed in DNA occupancy in the OFF state and measured both transcriptional output and growth effects across TF input levels (Supplementary Fig. 8). These data showed that maintaining CIC-TFs in an OFF state with low activity reduced the burden associated with the architecture under the tested conditions, whereas persistent DNA binding impaired growth at high TF input. To characterize the two module classes separately, DBDs were tested with RpaR179 as a common LBD, whereas LBDs were tested with LexAbs94 as a common DBD. These measurements provided DBD- and LBD-specific parameters that could then be tested in new pairings.

### Module parameters predict additional CIC-TF combinations and guide prospective design

With promoter output windows and DBD/LBD modules in hand, we tested whether parameters estimated in fixed-domain backgrounds could be recombined to predict complete CIC-TF transfer functions for DBD–LBD combinations not used for fitting. We established a quantitative yeast testing system in which TF expression level and inducer concentration could be independently varied (Supplementary Fig. 9). The DBD–RpaR_179_ series was used to estimate DBD-specific *K_A_* values, whereas the LexA_bs94_–LBD series was used to estimate LBD-specific active pool parameters *K_d0_*, *K_b_* and *K_d1_*; these 27 combinations were used to characterize parameters for the individual DBDs and LBDs (orange fills in Fig. 4B). The fitted parameters were then recombined to predict 15 additional DBD–LBD combinations. These combinations were excluded from the corresponding parameter estimation and used for cross-validation (blue fills in Fig. 4B). The predicted dose-response outputs agreed closely with experiment (median cross validation log-scale R² = 0.82 across the 15 combinations; Fig. 4B), showing that parameters measured for individual modules could be reused when those modules were paired with different partners. The sensor designs described below provide a separate prospective test because they were selected from the predicted design space before experimental testing.

The transferability of the module parameters was also examined by normalizing response curves for DBDs paired with the same LBD. For the RpaR_179_ series, correction for differences in DBD- associated occupancy brought the dose-response data onto a shared LBD-specific trend (Fig. 4C). The curves did not collapse perfectly, as expected from residual context effects and measurement variation, but they followed the same overall relationship. This result supports the separate measurement and subsequent recombination of LBD and DBD contributions across CIC-TFs.

We next used the fitted parameters to design sensors with specified performance. From each predicted transfer function, we calculated performance metrics including fold change, expression shift ΔExp, output strength, sensitivity, crosstalk and TF input as a proxy for burden (Fig. 4A). These metrics allowed the model to generate landscapes across *K_A_* and TF input for each LBD, and to select candidate designs according to specified performance criteria. As a first test, we considered acVHH, which produced high output but substantial basal leakage. The design landscape predicted that acVHH performance could be improved by selecting a DBD affinity and TF abundance that reduced basal leakage while retaining high output. The configuration selected by the model, pairing acVHH with lexA_mm115_ at 0.21 RPU, was experimentally validated and produced the expected combination of high output and low leakage (Fig. 4D).

We also tested whether the model could improve a sensor for combined dynamic range and absolute expression change. For BjaR_180_, we maximized a composite log(FC × ΔExp) objective across DBD affinity and *C_TF,all_*, identifying a configuration containing lexAbs94 at 4.59 RPU that improved the joint objective and was experimentally verified (Supplementary Fig. 11B). In both cases, the model identified combinations of DBD identity and TF expression that were then confirmed experimentally.

We next used the model to reduce ligand crosstalk in ER282–595, which responded to both β- estradiol and DHB. We generated a Pareto landscape that balanced cognate induction, fold change and off-target activation. Designs with higher DBD affinity and TF expression produced stronger output but also greater off-target activation, whereas lower operating points improved orthogonality at the cost of output strength. Three representative designs were selected for validation, and the lexA_bs94_ design at 0.66 RPU provided a favorable compromise between β- estradiol responsiveness and reduced DHB crosstalk (Fig. 4E). Additional designs intended to reduce crosstalk supported the same conclusion (Supplementary Fig. 11A). Because the three designs were chosen before experimental testing, they provide a prospective test of the crosstalk objective.

We implemented these calculations in CIC-Designer^10, 11, 65, 66^ (Supplementary Fig. 10). CIC- Designer fits module parameters from data supplied by the user, searches the resulting design space according to specified design criteria, maps target *K_A_* and *C_TF,all_* values to characterized DBDs and expression cassettes, and returns candidate TF–promoter configurations for assembly. As new DBDs, LBDs or promoters are characterized, their parameters can be added to the same library and used in subsequent designs.

### The CIC occupancy model extends to repression and hybrid promoter control

We tested parameter transfer across regulatory polarity by converting CIC activators into repressors. The AD was removed and operator sites were repositioned between the TATA box and the transcription start site, where TF binding can competitively block transcriptional initiation (Fig. 5A and Supplementary Note 6). This inversion changes how promoter occupancy contributes to output, but it preserves the upstream relationship among ligand input, *C_TF,eff_*, DBD affinity and TF dosage.

**Figure 5.**
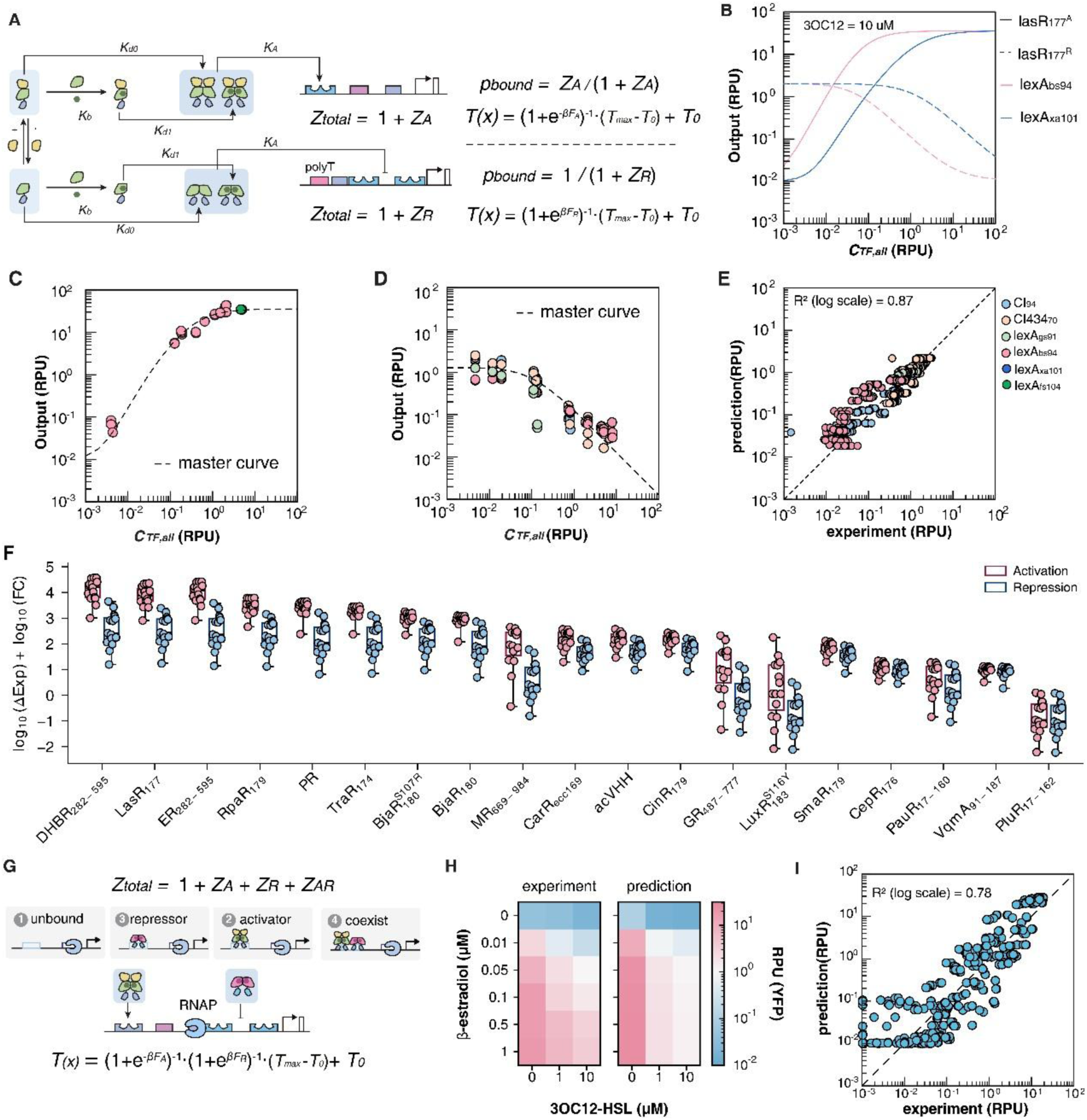
T**h**e **CIC occupancy model extends to repression and first-order hybrid regulation. (A)** Schematic and thermodynamic formulation of CIC-TF functional inversion from activator to repressor. In the activator configuration, ligand-induced active TF dimers bind upstream operator sites and increase transcriptional output. In the repressor configuration, the activation domain is removed and the operator is positioned between the TATA box and the TSS to block transcription. The same LBD-dependent effective TF pool and DBD/operator binding parameter are retained, while promoter occupancy is coupled oppositely to transcriptional output. For the repressor, *p_bound_* denotes the probability of the transcriptionally permissive, repressor-unbound promoter state, rather than repressor occupancy. **(B)** Theoretical comparison of activation and repression transfer functions for LasR_177_-based CIC-TFs. Solid lines indicate activator configurations and dashed lines indicate repressor configurations. Colors denote DBD/operator choices. The analysis illustrates how functional inversion changes the direction and dynamic range of regulation while preserving the underlying ligand-dependent TF activation model. **(C–D)** Master-curve comparison of LasR_177_- based CIC-TFs in activator and repressor architectures. Experimental responses from different DBD–LasR_177_ variants were normalized by their DBD-specific binding parameters and plotted against total TF abundance *C_TF,all_*. Dashed lines indicate the LasR177 transfer function predicted by the shared model for activator-type TFs **(C)** and repressor-type TFs **(D)**, showing that the dominant LBD-specific behavior is retained after functional inversion. **(E)** First-order evaluation of CIC-TF repressors using parameters obtained from activator data, without refitting the repressors. Scatter plot compares measured and model-predicted outputs across inverted repressor datasets. Colors indicate DBD identity, and the dashed red line denotes y = x. The R² value was calculated on untransformed outputs and summarizes how well the model captures output trends. **(F)** Predicted upper-bound regulatory capacity of LBD modules in activator and repressor configurations within the tested promoter geometry and TF input range. For each LBD, the feasible parameter space was scanned to estimate log_10_(ΔExp)+log_10_(FC). Box plots summarize simulated designs, with individual points representing candidate parameter combinations. Pink and blue indicate activation and repression architectures, respectively. **(G)** Extension of the thermodynamic model to hybrid promoter regulation containing both activator and repressor inputs. The promoter can occupy unbound, activator-bound, repressor-bound, or activator–repressor co-bound states. Hybrid promoter output was modeled using a first-order independence approximation, in which the activator-dependent and repressor-dependent permissive-state probabilities are combined multiplicatively. **(H)** Experimental validation of hybrid promoter regulation. Heatmaps compare measured and model-predicted outputs across two-dimensional inducer combinations controlling an activator input and a repressor input. β-estradiol controls the activator module, and 3OC12-HSL controls the repressor module. Color indicates output in RPU. **(I)** Evaluation of the first-order independence approximation for hybrid promoter regulation. Measured and model-predicted outputs are compared across all tested inducer combinations. The dashed diagonal line denotes y = x. Both axes are displayed on logarithmic scales. The displayed R² value was calculated on untransformed experimental and predicted outputs and summarizes model agreement across all tested hybrid promoter conditions, indicating that a simple multiplicative model captures first- order hybrid promoter behavior.

Using LasR_177_ as a representative LBD, we constructed and characterized repressive CIC-TFs across promoter output contexts (Fig. 5B). After normalization by promoter occupancy, the activation and repression data followed the corresponding master curves (Fig. 5C,D). Using parameters derived from activator characterization, the model captured repressor outputs with log-scale *R^2^* = 0.87 across the tested datasets (Fig. 5E and Supplementary Note 10). This result supports parameter reuse across regulatory polarity, while promoter geometry and steric blocking remained specific to each architecture.

We then compared the optimized design envelopes of activation and repression under the same accessible *C_TF,all_* range of 0.2–10 RPU. Using log_10_(FC × ΔExp) as a joint objective, activation generally provided a larger optimized performance envelope than repression for the tested LBD panel and promoter geometry (Fig. 5F). This likely reflects the difference between recruiting transcription above a low basal state and maintaining sufficient promoter occupancy to repress an already active promoter. Under these conditions, activation was favored for high-performance CIC sensing, although the same occupancy model could also be used to design ligand-dependent repression.

Finally, we asked whether activator and repressor modules could be combined within one promoter to implement multi-input control. We constructed hybrid promoters containing an upstream activator-binding site and two downstream repressor sites (Fig. 5G). Orthogonal CIC-TFs independently controlled the activator and repressor inputs, allowing output to be controlled across two input dimensions (Fig. 5H). A simple multiplicative model captured the first-order hybrid promoter behavior with *R^2^* = 0.78 (Fig. 5I and Supplementary Note 11). The hybrid data were therefore consistent with a first-order extension of the occupancy model, although additional promoter interactions will be needed to describe more complex logic.

### Host-specific measurements extend the CIC design space to mammalian cells

We next tested whether the CIC model could be transferred to mammalian cells without exhaustive screening. Host context can alter chromatin accessibility, nuclear transport, TF stability, cofactor availability, promoter output and endogenous transcriptional competition. We therefore retained the model structure but re-estimated its quantitative parameters from mammalian measurements (Fig. 6A).

**Figure 6.**
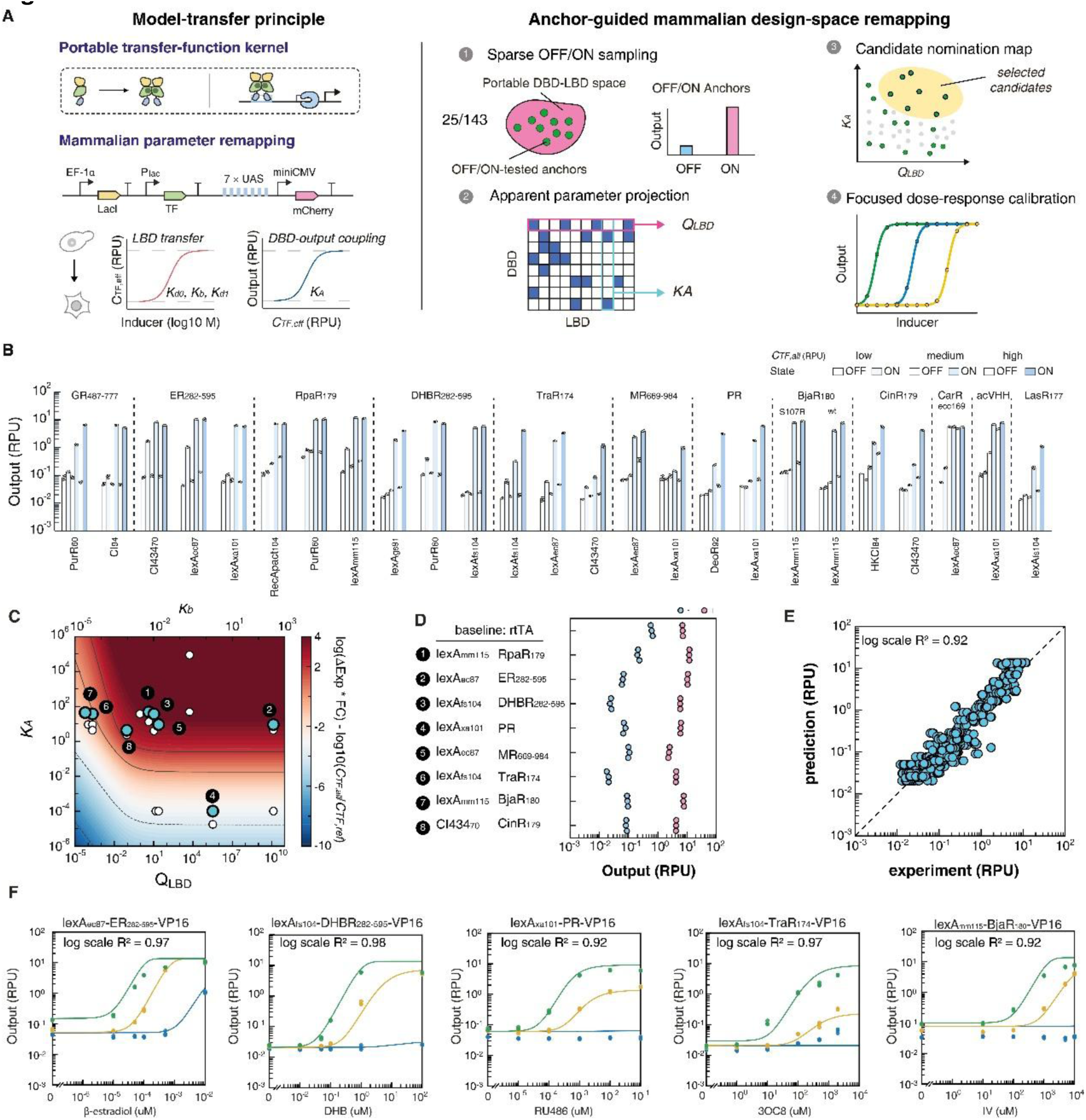
L**i**mited **host-specific measurements guide the selection and calibration of mammalian CIC-TFs. (A)** Host-specific calibration of the CIC-TF design space in mammalian cells. The model structure is retained across hosts, whereas its quantitative parameters are re-estimated from mammalian measurements. OFF/ON measurements of selected DBD–LBD combinations provide initial estimates of QLBD and KA and guide the selection of candidates for dose-response characterization. **(B)** OFF/ON measurements in stable cell lines provide an initial map of mammalian CIC-TF performance. A selected subset of the portable DBD–LBD design space was constructed as CIC-TF reporters in stable mammalian cell lines and measured under OFF and ON ligand conditions. Reporter output is shown for each DBD–LBD pair at three calibrated TF input levels. As the measured *C_TF,all_* input RPU values differ among sensors, exact sensor-specific values are provided in the Source Data. OFF indicates no cognate ligand, whereas ON indicates the high inducer concentration used for the corresponding LBD: dexamethasone for GR_487–777_ (10 μM), β- estradiol for ER_282–595_ (0.01 or 0.5 μM, as indicated for each construct), pC-HSL for RpaR_179_ (200 μM), DHB for DHBR_282–595_ (100 μM), 3OC8-HSL for TraR_174_ (2000 μM), aldosterone for MR_669–984_ (1 μM), RU486 for PR (0.1 μM), IV-HSL for BjaR_180_ and BjaR_180_^S107R^ (10,000 μM), 3OHC14-HSL for CinR_179_ (20 or 100 μM, as indicated for each construct), cognate HSL for CarR_ecc169_ (100 μM), caffeine for acVHH (10 μM), and 3OC12-HSL for LasR_177_ (10 μM). Bars represent mean output (RPU), overlaid points represent individual biological replicates, and error bars indicate SD (n = 3). DBDs are labeled beneath each bar group, and LBD groups are indicated above the plot. The resulting OFF/ON matrix provides the initial mammalian parameter estimates used in the design surface. **(C)** Mammalian CIC-TF objective landscape under fixed *K_d1_*/*K_d0_* and variable *K_b_*. Predicted mammalian CIC-TF design landscape calculated by fixing K_d1_/K_d0_ = 10^5^ and varying *K_b_*. The bottom x-axis denotes the LBD induction potential, (*Q_LBD_ = K_b_*_2_ *× K_d1_/K_d0_*), and the y-axis denotes the DBD–operator association parameter, (*K_A_*). Both axes are shown on logarithmic scales. The top x-axis indicates the corresponding (*K_b_*) value for each *Q_LBD_* under the fixed *K_d1_*/*K_d0_* setting. For each (*Q_LBD_*, *K_A_*) coordinate, OFF and ON outputs were computed using the mammalian CIC-TF transfer function model with (*c_I_* = 0) and (*c_I_ =* 100 *μM*), respectively. (*C_TF,all_*) was scanned over the mammalian calibrated expression range of 0.01–3.02 RPU, and the maximum objective value was plotted: J = max[log_10_(ΔExp × FC) − log10(*C_TF,all_*/*C_TF,ref_*)]. Background color represents the optimized objective value, with red indicating higher predicted performance and blue indicating lower performance. Gray contour lines indicate equal objective values. Open circles with black outlines indicate TF variants characterized only by OFF/ON measurements. Blue circles with black outlines indicate TF variants used for full dose-response characterization and parameter fitting. Black numbered labels identify the eight prioritized variants shown in panel D. **(D)** Mammalian CIC-TFs selected from candidates characterized with full dose-response curves. Among mammalian CIC-TFs subjected to full inducer dose-response characterization, the highest-scoring designs were selected using log10(ΔExp × FC). The selected TFs are shown relative to the rtTA benchmark under our assay conditions. Right, OFF and ON outputs for the prioritized designs under non-induced and maximally induced conditions, respectively. Each point represents an experimentally measured condition. **(E)** Model agreement after joint calibration. Experimentally measured outputs were compared with model-fitted outputs across a combined calibration dataset comprising 15 mammalian CIC-TFs characterized by full inducer dose-response curves and 10 additional CIC-TFs characterized by endpoint OFF/ON measurements (25 CIC-TFs in total). Both full dose-response and endpoint measurements contributed to parameter estimation. Each point represents one biological-replicate measurement at a defined TF input and inducer condition. The dashed line indicates (y = x). The pooled log-scale R^2^ across the combined calibration dataset was 0.92. **(F)** Representative dose-response fits for prioritized mammalian CIC-TFs. Inducer dose-response curves for selected mammalian CIC-TFs spanning distinct DBD–LBD combinations. Points indicate individual biological-replicate measurements (n = 3 per condition; some points overlap), and solid lines indicate model-fitted transfer functions. Colors denote different calibrated TF input levels. For LexA_ec87_-ER_282-595_-VP16, lexA_fs104_-DHBR_282-595_-VP16 and lexA_xa101_-PR-VP16 sensors, the input levels are: blue, 0.01 RPU; yellow, 0.48 RPU; green, 3.32 RPU. For LexA_fs104_-TraR_174_-VP16 and LexA_mm115_-BjaR_180_-VP16, the input levels are: blue, 0.01 RPU; yellow, 0.32 RPU; green, 2.72 RPU. These fits show the range of basal and ligand-induced responses captured after full dose-response calibration.

Using HEK293T reporter cells carrying a genomic landing pad, we profiled 25 representative DBD– LBD combinations selected from a design space of 11 DBDs × 13 LBDs (143 possible combinations). Each design was measured at three calibrated TF input levels under uninduced and saturating inducer conditions, producing a compact OFF/ON matrix of basal and maximally induced outputs (Fig. 6B and Supplementary Fig. 12). From these endpoint data, we estimated an apparent DBD– output coupling parameter *K_A_* and a metric of LBD induction quality *Q_LBD_* = *K_b_*_2_*K_d1_*/*K_d0_*, which reports the ligand-dependent enrichment of the DNA-binding-competent TF pool. The fitted parameter map reconstructed the OFF/ON measurements with high agreement (Supplementary Fig. 13A).

We used the remapped parameters to generate a mammalian design surface. For each *Q_LBD_*–*K_A_* coordinate, the model scanned the calibrated mammalian TF input range and maximized an objective that rewarded large expression shifts and high fold change while penalizing excessive TF input (Fig. 6C). We used the resulting surface to identify DBD–LBD combinations for detailed inducer dose-response characterization.

We then performed full inducer dose-response characterization for 15 selected CIC-TFs across three TF input levels. These measurements refined the mammalian parameters and tested whether the initial rankings from the endpoint survey were retained. LBD induction quality was highly conserved between estimates obtained from endpoint measurements and full dose-response curves after log10 transformation of the fitted parameters (Pearson r = 0.84; Supplementary Fig. 13C). In contrast, DBD KA showed a coarser correspondence under the same analysis (Pearson r = 0.87; Supplementary Fig. 13B), consistent with the lower identifiability of DBD–output coupling from endpoint measurements alone. These results indicate that the OFF/ON survey is useful for prioritizing candidates, whereas full dose-response measurements are required for final calibration.

This procedure yielded eight optimized mammalian CIC-TFs based on RpaR_179_, ER_282–595_, DHBR_282– 595_, PR, MR_669–984_, TraR_174_, BjaR_180_ and CinR_179_. Under the matched assay context, these CIC-TFs expanded the accessible dynamic range from approximately 16-fold for rtTA^16–18^ to 24–241-fold activation, while maintaining comparable maximal expression and lower basal leakage (Fig. 6D,F and Supplementary Fig. 13D–M). After calibration with the full dose-response data, the mammalian model captured reporter outputs spanning several orders of magnitude (log scale R² = 0.92, Fig. 6E). The refined mammalian DBD and LBD parameters differed from their yeast counterparts, confirming that portability depends on host-specific recalibration rather than transfer without further measurement (Supplementary Tables 2–5). A limited OFF/ON survey was therefore sufficient to identify promising combinations, whereas full dose-response measurements provided the quantitative calibration required for final sensor design.

### Model-guided panel optimization enables a 12-sensor yeast chassis

We next tested whether independently characterized CIC sensors could be combined in a multi- channel regulatory chassis. Strains carrying multiple sensors have been powerful in prokaryotic engineering^86^, but eukaryotic chassis construction is more difficult because many heterologous regulators must operate simultaneously without excessive leakage, crosstalk, repeated sequences or expression burden^87–89^. We therefore treated panel assembly as an optimization problem involving several performance criteria.

The CIC-12 sensor panel was designed in silico by assigning each of nine CIC LBDs to a distinct DBD/operator module and a TF input level within a biologically accessible range of 0.2–10 RPU (Fig. 7A). A total of 15,000 candidate nine-sensor panels were generated using a reproducible procedure based on random sampling; each panel contained all nine LBD sensors paired with distinct DBDs. Candidates were evaluated using mean log_10_(ΔExp), mean log_10_(FC), minimum orthogonality index, total TF input and maximum leak, and were compared by Pareto analysis across these five objectives and by a normalized objective score that penalized high TF input and leakage (Supplementary Note 8.4). This in silico search avoided constructing every DBD–LBD–input assignment experimentally.

**Figure 7.**
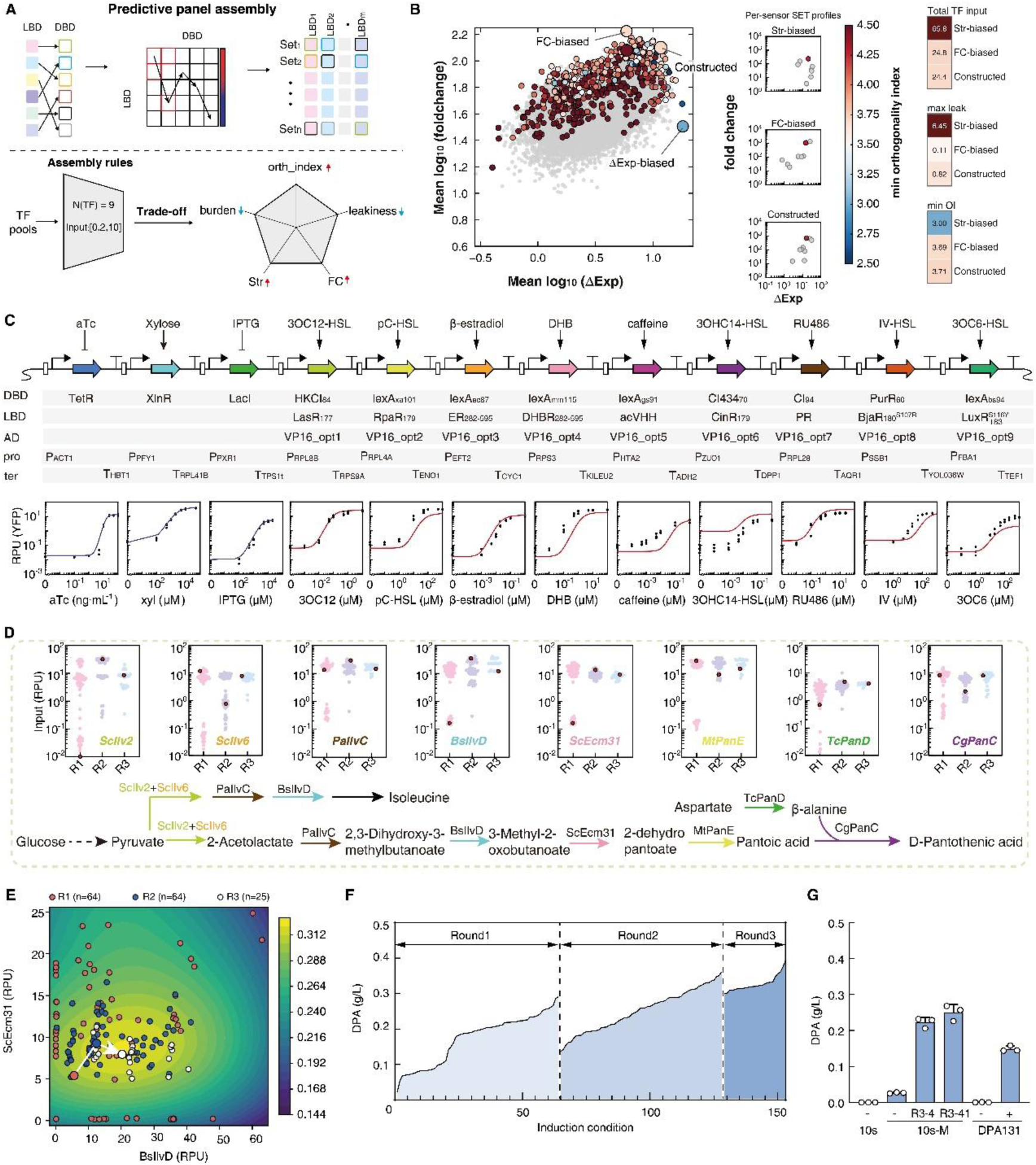
P**a**nel **design and GP-guided optimization extend CIC to multigene control. (A)** Strategy for model-guided assembly of the CIC-12 multisensor chassis. Nine CIC-TFs were selected in silico by assigning a fixed panel of nine ligand-binding domains (LBDs) to distinct DNA-binding domain (DBD)/operator modules and by selecting a total TF input level for each sensor within 0.2–10 RPU. The transfer function model was used to calculate OFF output, cognate-induced output, expression shift (ΔExp), fold change, sensor leakage, total TF input, and orthogonality for each combination of DBD, LBD and TF input. Complete nine-sensor panels were then compared using performance criteria and design constraints, including mean expression shift, mean fold change, minimum known orthogonality index, total TF input, and maximum leak. The model was used to select the CIC-TF panel; TetR, LacI, and XlnR were subsequently included as additional sensors in the final CIC-12 chassis. **(B)** Evaluation of candidate nine-sensor panels across multiple objectives. Nine fixed CIC LBDs were assigned to distinct DBD/operator modules and evaluated across TF input levels of 0.2–10 RPU. Each gray point represents one panel generated by the reproducible sampling procedure; colored points with black outlines indicate panels that are not dominated across five objectives: mean log_10_(ΔExp), mean log_10_(FC), minimum orthogonality index (OI), total TF input and maximum leakage; OI was calculated as described in Supplementary Note 8.4. Point color denotes minimum OI. Enlarged points indicate the ΔExp-biased SET, FC-biased SET, and the experimentally implemented Constructed SET evaluated at its implemented TF inputs. The Constructed SET values shown here are model predictions rather than experimental measurements. Middle, per-sensor ΔExp–fold change profiles for the four highlighted SETs; the red point marks the sensor with the lowest OI within each SET and gray points denote the remaining sensors. Right, total TF input, maximum leakage and minimum OI for each panel. Experimental characterization of the implemented CIC-12 chassis is shown in C and Supplementary Fig. 14. **(C)** Architecture and functional characterization of CIC-12. Twelve orthogonal sensors, including TetR, XlnR, LacI, and nine CIC-TFs selected by the model, were assembled using distinct DBDs, LBDs, activation domains, promoters, and terminators. Dose-response curves show YFP output for each sensor in response to its cognate inducer. **(D)** Application of CIC-12 to programmable control of the vitamin B5 biosynthetic pathway. Each of the eight pathway genes was controlled by a different sensor, allowing enzyme expression to be tuned independently. Small plots show input levels sampled across optimization rounds R1–R3, and the pathway diagram indicates the corresponding enzymatic steps from glucose/pyruvate-derived intermediates to D-pantothenic acid. **(E)** Gaussian process (GP)-guided search paths for vitamin B5 pathway optimization. Two-dimensional projections of the GP-predicted DPA production landscape are shown for selected pairs of pathway control inputs. Symbols indicate experimentally tested induction conditions from the initial design-of-experiments (DOE) round followed by two GP-guided rounds. The color scale represents the GP-predicted yield (g L⁻¹). Each circle represents one tested condition; red, blue and white circles denote Rounds 1, 2 and 3, respectively. The white line connects the coordinate-wise geometric means of Rounds 1, 2 and 3, with each path vertex colored according to its corresponding round. **(F)** Measured DPA production across induction conditions tested during the iterative optimization in microplates. Each point represents one experimentally tested induction condition from the initial DOE followed by two GP-guided optimization rounds. Vertical dashed lines separate the initial DOE round and the two subsequent GP-guided optimization rounds. Together with the projections of the search paths in **E** and Supplementary Fig. 15, this panel summarizes the optimization trajectory; final quantitative production comparisons are supported by the independent validation experiment shown in **G. (G)** Independent validation of selected production conditions in shake flasks. DPA titres are shown for the parental CIC-10 chassis and the DPA131 reference strain under the indicated induction conditions, including the model-guided conditions R3-4 and R3-41. Each data point represents one independent biological replicate (n = 3 for each strain–induction condition combination). Bars indicate the mean, and error bars indicate the standard deviation (SD).

The resulting design landscape revealed structured tradeoffs among strength, fold change, orthogonality and TF burden (Fig. 7B). We used this landscape to select a ΔExp-biased panel, an FC-biased panel and a panel chosen using the balanced objective. The experimentally constructed panel was chosen as a balanced, buildable design within this predicted landscape, rather than as a claim of a strict optimum predicted solely by the model. This selection preserved high predicted strength and fold change, maintained a favorable minimum orthogonality index, avoided DBD reuse and excessive TF input burden, and provided a blueprint for selecting non-repetitive promoter and terminator sequences^90^ (Fig. 7B and Supplementary Note 8.4).

We constructed CIC-12 in two yeast strain backgrounds, BY4741 12s and CEN.PK2-1C 12s, by iterative chromosomal integration of 12 orthogonal sensors: nine newly designed CIC-TFs plus TetR, LacI and XlnR^10, 91^ (Fig. 7C and Supplementary Fig. 14A). Characterization of the final strains showed that each sensor responded specifically to its cognate inducer with minimal functional crosstalk across the panel (Supplementary Fig. 14B,C). In the BY4741 12s strain, the arithmetic mean of the background-corrected ON/OFF fold changes across the 12 sensors was 298-fold (Fig. 7C and Source Data Fig. 7C). Growth effects were small and depended on the strain background under the tested conditions, consistent with the low basal activity of the CIC modules (Supplementary Fig. 14D). Additional non-repetitive reporter tests further supported the orthogonality of the chassis design (Supplementary Fig. 14E,F).

Each of the 12 sensors responded to a distinct chemical input, allowing multiple genes to be tuned independently in the same strain. We applied CIC-10, an earlier version before CIC-12 construction, to optimize an eight-gene vitamin B5 biosynthetic pathway^92^ (Fig. 7D). Each pathway gene was controlled by a different sensor, allowing the expression of all eight enzymes to be adjusted independently in the same strain.

We then used an initial design-of-experiments (DOE) set followed by two rounds of Gaussian process (GP)-guided optimization to select inducer combinations for the eight-gene pathway^65, 66^ (Fig. 7E,F, Supplementary Fig. 15 and Supplementary Note 8.5). The model was updated after each experimental round and used to select the next set of inducer combinations. Final production was assessed in independent validation experiments (Fig. 7G). The GP-guided microplate screening identified induction programs with D-pantothenic acid (DPA) titres approaching 0.4 g l⁻¹ (Fig. 7F). Independent validation of the selected strains in shake flasks yielded approximately 0.25 g l⁻¹ DPA, representing a 1.7-fold increase over the DPA131 reference strain under matched fermentation conditions (Fig. 7G). These experiments show that the CIC platform can be used to tune an eight-gene pathway without rebuilding the underlying sensors.

## Discussion

Decades of work have clarified core biophysical rules of transcription, and in prokaryotes thermodynamic models coupled with host-specific calibration have enabled increasingly predictive regulatory control^62–64, 93–96^. In eukaryotes, however, chromatin context, promoter architecture, endogenous regulatory interactions and complex transcriptional recruitment have made it difficult to connect molecular parts to promoter-level transfer functions^44–47, 54^. Empirical promoter engineering, hybrid promoters, random mutagenesis and receptor-level pathway tuning can identify useful operating points^51, 55, 58–61, 97, 98^, but they rarely identify the quantitative parameters that define feasible responses. Here we show that CIC combines induced cooperativity with reusable quantitative parameters for eukaryotic transcriptional control. Parameters describing LBD activation, DBD–operator coupling and promoter output were measured separately and reused across additional DBD–LBD combinations, different regulatory modes and, after recalibration, mammalian cells. The predictive results therefore depend on the CIC architecture and its parameterization, rather than on chimeric assembly alone. Three observations emerge from these experiments.

### i. Induced cooperativity supports modular domain exchange and low basal occupancy

Natural allosteric regulators show how weak molecular states can remain silent until ligand binding, oligomerization or cofactor assembly shifts the system into an active configuration. Yet in many Type I systems, signal sensing, DNA recognition and regulatory output are structurally coupled, making exchange of DNA specificity, affinity tuning or logic inversion difficult without mutagenesis or directed evolution^19–23^. Type II systems address modularity in a different way. CID-, dCas-, ZF- and TALE-based architectures provide chimeric modularity, but their DNA-binding modules are often present at the target site before induction. This constitutive occupancy can redistribute failures among leakage, dynamic range, targeting efficiency and host burden rather than eliminating the trade-off^27–43^. CIC combines chimeric domain exchange with ligand-dependent formation of a DNA-binding-competent TF pool, which limits basal occupancy while preserving induced occupancy. The comparison between activation and repression further showed that the same upstream occupancy parameters can be retained when the effect of promoter binding is inverted. Activation can exploit thresholded recruitment^99^ above a low basal state, whereas repression requires sustained promoter occupancy to block an already active promoter. These results suggest that modularity is most useful when the architecture also controls the ligand dependence of DNA occupancy.

### ii. DBD state dependence and promoter output windows define complementary design axes

DBD specificity and affinity alone did not fully predict CIC-TF performance. Effective DBDs maintained low activity in the OFF state while efficiently converting the ligand-induced TF pool into promoter occupancy. Designed DNA-binding proteins should therefore be evaluated not only for affinity and specificity, but also for how promoter occupancy changes between regulatory states^73, 100, 101^. This state dependence adds a functional criterion to current computational and generative approaches for DBD design.

Promoters provide a complementary layer of control. Operator number tuned occupancy gain, whereas spacer and core promoter sequences determined *T_0_* and *T_max_.* These quantities provide explicit design targets that can be linked to existing promoter sequence models^47, 54, 61^. Together, DBD tuning and promoter design allow input coupling and output range to be adjusted separately, expanding the accessible design space without changing the ligand-sensing module.

### iii. Parameter reuse reduces the number of complete combinations that must be measured

The practical advantage of CIC parameterization is that measurements made in one construct can inform others. Parameters measured for an LBD, DBD or promoter can be recombined across a DBD × LBD × expression × promoter space. In yeast, parameters estimated from fixed DBD or LBD backgrounds predicted 15 additional DBD–LBD combinations in cross-validation. This reuse reduces the number of complete constructs that must be measured and expands the design space that can be examined experimentally, including feasible regions and Pareto frontiers.

The parameter library can also be used in automated design and active learning workflows^65, 66^. CIC-Designer fits parameters and selects TF–promoter configurations before construction. In mammalian cells, 25 combinations characterized by OFF/ON measurements were used to select candidates from a 143-combination design space before focused dose-response characterization.

CIC-12 extended the same approach to panel design by balancing output, leakage, orthogonality and total TF input across multi-channels. Prokaryotic systems with multiple sensors have been powerful but often depend on host-specific directed evolution; comparable eukaryotic systems have been constrained by leaky or burdensome regulators^86–89^. CIC-12 is comparable in scale to Marionette^86^ and was assembled in yeast by optimizing the complete sensor panel. The resulting strain supported coordinated control of a multi-gene expression program.

The current framework has several limitations. Chemically diverse LBDs, especially nuclear receptor-like modules, are represented by an apparent active pool model and need not share one microscopic dimerization mechanism. Quantitative parameters must be re-estimated when the host context changes. Genome-wide off-target binding by DBDs, response kinetics, protein stability and chromatin remodeling are not yet included, and the hybrid promoter model captures only first-order interactions. Future extensions could incorporate de novo DBDs designed to preserve state-dependent occupancy^101^, designed heterodimers for asymmetric AND/OR/XOR logic^102, 103^, improved promoter models and iterative active learning. Within these limits, CIC provides a quantitative route from component measurements to the design of inducible eukaryotic transcriptional regulators.

## Methods

### Strains, media, and reagents

*E. coli* MG1655 with AraR, LacI, and TetR integrated in the attB site (MG1655 ALT) was used for initial domain screening and functional validation. *S. cerevisiae* strain CYE72^10^, derived from BY4741 (S288C MATa his3Δ1 leu2Δ0 met15Δ0 ura3Δ0), was used for qualitative domain characterization and quantitative parameterization. *E. coli* cells were grown in M9 medium supplemented with 0.4% glycerol (Sangon Biotech, A501745). SD-Ura-Leu (Coolaber, PM2291) medium was used to grow *S. cerevisiae* cells with induced transcriptional systems. For vitamin B5 production, SD medium with 4% glucose was used.

Carbenicillin (100 mg/L, Sangon Biotech, A600469), kanamycin (50 mg/L, Aladdin, K103024), or chloramphenicol (25 mg/L, Sangon Biotech, A100230) was added to LB medium at the indicated final concentration for plasmids carrying sdAmpR, KanR, or sdCmR, respectively. Agar was added to 2% (w/v, Bacto, 214010) for solid medium. L-Leucine (Sigma-Aldrich, PHR1105) and uracil (Sigma-Aldrich, U0750) were supplemented as required for the corresponding auxotrophic strains.

The inducers used to control TF expression included IPTG (Isopropyl β-D-Thiogalactoside, Sigma-Aldrich, I6758) dissolved in water for 1.0 M stock; aTc (anhydrotetracycline hydrochloride, Aladdin, A276282) dissolved in DMSO for 100 μg/mL stock; D-xylose (Sangon Biotech, A600998) dissolved in water for 1 M stock. Ligands used to alter TF activity included caffeine (GLPBIO, R3136V); N-(3- Hydroxytetradecanoyl)-DL-homoserine lactone (3OHC14-HSL; Sigma-Aldrich, 51481); N-(3- Oxododecanoyl)-L-homoserine lactone (3OC12-HSL; Sigma-Aldrich, O9139); N-3-(Oxooctanoyl)-L-homoserine lactone (3OC8-HSL; APE×BIO, C4369); N-(β-Ketocaproyl)-L-homoserine lactone (3OC6-HSL; Sigma-Aldrich, K3007); N-Butanoyl-L-homoserine lactone (C4-HSL; SML3427); N-(p-Coumaroyl)-L-homoserine lactone (pC-HSL; Sigma-Aldrich, 07077); isovaleryl-homoserine lactone (IV-HSL; chemically synthesized); β-estradiol (TCI, E0025); mifepristone (RU486; Selleck, S2606); aldosterone (NEB, GC41390); dexamethasone (GLPBIO, GC40775); 1,2-bis(4- hydroxyphenyl)ethane-1,2-dione (DHB; Mreda, M093053). All chemical inducers were prepared as stock solutions.

### Plasmid construction

Backbone vectors were first constructed by placing ccdB between matching type IIS restriction sites (BsaI, New England Biolabs, R3733L or BpiI, Fermentas, ER1012) and assembly overhangs. Insert fragments were generated by PCR with the corresponding restriction sites and overhangs. The backbone vectors were transformed into E. coli TransDB3.1 (TransGen Biotech, CD531) for construction and amplification. Genes were synthesized after codon optimization. Internal BpiI or BsaI recognition sites were removed by synonymous substitution. Amplification was performed using Q5 DNA polymerase (New England Biolabs, M0491L). Golden Gate assembly was used to construct plasmids and the assembly products were then transformed into *E. coli* DH10B ALT (DH10B with AraR, LacI, and TetR integrated into the attB site) or Trans10 (TransGen Biotech, CD101) competent cells and plated on LB agar containing the appropriate antibiotic.

Based on the study by Guo et al^92^, we selected eight genes, including *CgPanC*, *TcPanD*, *MtPanE*, *PaIlvC*, *BsIlvD*, *ScIlv2*, *ScIlv6*, *ScEcm31*, for combinatorial metabolic optimization because they had been previously identified as important determinants of vitamin B5 production. For the eight genes in the core pathway of vitamin B5 synthesis, the expression strengths identified in the study were ranked. Sensors with regulatory ranges matching the expression levels for each gene were selected for precise expression control. All plasmid constructs were assembled by Golden Gate.

### Chromosomal integration of TF and reporter cassettes in yeast

For qualitative and quantitative tests of domains in *S. cerevisiae*, the TF cassette was integrated at the URA3 locus on chromosome V, and the reporter cassette was integrated at the LEU2 locus on chromosome III. Linear fragments for S. cerevisiae transformation were prepared by BsaI digestion.

The Frozen EZ Yeast Transformation Kit II (Zymo Research Corporation, T2001) was used for the competent cell preparation and yeast transformation. Transformants were selected on the appropriate SD plates and incubated at 30 °C for 2 d.

### Iterative chromosomal integration of multi-sensor arrays

Sensor arrays were integrated iteratively at a site on chromosome XV between the GPY1 and NRT1 loci in BY4741 and CEN.PK2-1C. The linear integration fragment used in each round consisted of four parts: a 5’ upstream homologous arm, transcription factor (TF) expression cassettes, a selection marker, and a 3’ downstream homologous arm. The integration process alternated the use of antibiotic selection markers (NAT and G418 sulfate) to allow recycling and reuse. For example, in the first round, the integration fragment included expression cassettes for three transcription factors (tetR, xlnR, and lacI), the NAT selection marker, and the homologous arms. The upstream homologous arm corresponded to the sequence of Chr XV 458693 to 459153, and the downstream homologous arm matched the sequence of Chr XV 458192 to 458669. Transformants were obtained by selection with 100 µg/mL NAT. Reporter constructs were subsequently integrated into the selected transformants, and flow cytometry analysis was conducted to verify the functionality of each integrated transcription factor. After reporter function was validated by flow cytometry, the next integration round was performed. For the subsequent integration of 4–10 sensors, G418 sulfate was used for selection. The 5’ homologous arm of the second-round integration fragment was designed to correspond to the upstream 500 bp of the selection marker from the previous round, enabling the replacement and recycling of the NAT selection marker. The 3’ homologous arm remained unchanged. The subsequent rounds of integration followed the same strategy.

### Mammalian cell culture, transient transfection, and stable cell line generation

HEK293T cells (ATCC CRL-3216) were maintained in Dulbecco’s modified Eagle’s medium (DMEM; Gibco) supplemented with 10% (v/v) fetal bovine serum (FBS; Biological Industries) and 1% (v/v) penicillin–streptomycin solution (Beyotime) at 37 °C in a humidified atmosphere containing 5% CO₂. For stable integration, 6–8 × 10⁵ cells suspended in 0.2 mL complete DMEM were seeded per well in 12-well plates (NEST) and incubated for approximately 24 h before transfection. Each transfection mixture contained 150 ng of donor plasmid carrying attB and the genetic circuit, 150 ng of Bxb1 integrase plasmid, and 0.6 µL of Lipofectamine 2000. At 12 h post-transfection, cells were trypsinized and transferred to 6-well plates (NEST) containing 3 mL complete DMEM per well. Puromycin was added 12 h after transfer to select stably integrated cells.

### Quantification of ethanol-responsive promoter activity in *S. cerevisiae*

For ethanol-responsive promoter activity quantification, single colonies of target strains (test strain, background strain, and reference strain) were picked and inoculated in SD or SDΔUra medium supplemented with 1% (v/v) ethanol, 1% (w/v) glucose, and the nutrients required for the corresponding auxotrophic genotype, and grown at 30 °C for 40–48 h to saturation. Saturated cultures were subsequently diluted 1:50 into fresh SD or SDΔUra medium containing 2% (v/v) ethanol and incubated at 30 °C with shaking. When cultures reached mid-log phase (OD_600_ ≈ 0.5), samples were collected, diluted in PBS, and single-cell fluorescence was measured by flow cytometry to quantify promoter activity under ethanol induction conditions.

### Screening of DBD modules in an *E. coli* repression assay

RpaR_179_ was used as a fixed dimerization domain for DBD screening in the *E. coli* repressor system. The repressor cassette was under the control of an IPTG-inducible PLlac promoter. A final concentration of 20 μM IPTG was used to induce TF expression, and the repression fold change was measured in the absence or presence of 100 μM pC-HSL. Three independent colonies per construct were inoculated into 200 µL LB medium in a 96-well plate and grown overnight at 37 °C with shaking at 800 rpm. A 1-μL aliquot was then transferred to 200 μL M9 glycerol medium containing the appropriate antibiotics and inducers at the indicated concentrations. Cells were incubated at 37 °C with shaking at 800 rpm for approximately 5 h, until the OD_600_ reached approximately 0.3. YFP fluorescence (excitation, 500 nm; emission, 530 nm) and OD_600_ were measured simultaneously using a microplate reader.

### Qualitative and quantitative characterization of CIC-TF modules in yeast

LexA_bs94_ and RpaR_179_ were used as the fixed DBD and LBD for qualitative and quantitative characterization of LBDs and DBDs, respectively. In the qualitative assay, TF expression was induced with 100 ng/mL aTc. Fold change was calculated between the uninduced and maximally induced ligand conditions. Cultures were grown at 30 °C and 800 rpm for 24 h, diluted 1:200 into fresh medium containing the indicated expression inducer and ligand, and incubated for 16 h. Flow-cytometry samples consisted of 20 µL culture and 180 µL PBS (PBS, Proteintech, PR20014). For quantitative parameter estimation, inducer dose-response curves were measured at multiple TF input levels. Seed cultures grown for 24 h were first transferred into fresh SD medium containing the TF-expression inducer and incubated for a further 24 h. Cells were then transferred into fresh medium containing both the TF-expression inducer and the ligand and incubated for 16 h before analysis.

### Assessment of chimeric TF-associated growth burden

Growth impact was evaluated following the procedure described by Guo et al.^91^, with minor modifications. Two single colonies of each test strain were picked and grown overnight in 500 μL SD medium under auxotrophic selection. Overnight cultures were then diluted 1:200 into non-selective SD medium supplemented with the indicated inducer concentrations and grown for 24 h to saturation. For OD_600_ quantification, 20 μL from each well was transferred into 480 μL of the corresponding inducer-containing medium in a second plate, and OD_600_ was measured after 8 h using a Tecan Infinite 200 Pro plate reader (Tecan Group Ltd.). In parallel, cultures from uninduced wells were inoculated into SD medium containing a gradient of inducer concentrations at a ratio of 1:200 and grown for an additional 24 h; these preconditioned cultures were used as the inoculum for OD_600_ measurements on the following day. Measurements were collected over three consecutive days, and all values were normalized to the parental BY4741_pgs093 strain cultured in SD medium and measured on the same day.

### Flow-cytometry acquisition and analysis

Reporter fluorescence was quantified by flow cytometry following the general procedure described previously^10, 91^. Samples were analyzed on a BD FACSCelesta flow cytometer equipped with a high-throughput sampler. yEmCitrine was excited with a 488-nm laser, and fluorescence emission was collected using a 530/30-nm band-pass filter. Data acquisition was performed with FACSDiva software. More than 10,000 events were collected for each sample. Yeast events were selected on the basis of forward-scatter area and side-scatter area, and events outside the cell gate were excluded from subsequent analysis. The same acquisition settings and cell-selection strategy were applied to the samples within each experiment. Flow-cytometry files were analyzed in FlowJo v10.8.1, and the median FITC-A signal of the gated population was used as the fluorescence output for each sample.

### Yeast CIC-TF model training and parameter estimation

We modeled the measured response *T(x)* as a function of total TF concentration *C_TF,all_* and inducer concentration *c_I_* using Eq. (S83), with 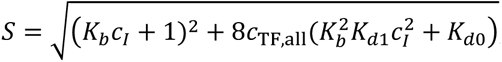; calibration parameters were fixed to *T_0_* = 0.035 and *T_max_* = 35.85. To ensure consistent parameterization across ligand-binding domains (LBDs), we performed hierarchical parameter estimation in three stages: (i) using the complete RpaR_179_ dataset, we fit globally shared kinetic parameters *K_d0_*, *K_b_*, *K_d1_* by minimizing mean-squared error in PyTorch with parameters optimized in log-space (positivity enforced) using Adam for 100,000 iterations with gradient clipping; (ii) fixing *K_d0_*, *K_b_*, *K_d1_* to the Stage 1 values, we independently estimated the DBD-specific association constant *K_A_* for each DBD dataset (150,000 iterations); and (iii) fixing the DBD-specific *K_A_* values from Stage 2, we refit *K_d0_*, *K_b_*, *K_d1_* for each additional LBD to obtain LBD-specific kinetics under a shared DNA-binding framework. Model performance was quantified by *R^2^* computed both globally and within individual experimental conditions.

### Master-curve construction and normalization

A unified master curve was constructed using the sensor response model (Eq. S83), which describes output as a function of total transcription factor concentration *C_TF,all_* and inducer concentration *c_I_*. The maximal response was fixed to *T_max_* = 35.85 RPU based on prior optimization, and each sensor variant was parameterized by its association constant *K_A_* (13 variants; *K_A_* = 0.10– 40.98). A reference sensor with *K_A_* = 6.22 was selected to define the master curve. For each inducer condition (c_I_=0 or 100 μM), experimental measurements from all variants were mapped onto the reference master curve using: *y_corrected_ = y_experimental_ +* [*T_master_(C_TF,all_)* - *T_variant_(C_TF,all_)*] - *T_0, variant_,* where *T_0, variant_* denotes the basal expression of each variant. The deformation term *T_master_* – *T_variant_* was obtained by interpolating the corresponding theoretical response curves at the experimentally sampled *C_TF,all_* values. Alignment quality was quantified by computing *R^2^* separately for each inducer condition across all corrected data points. All computations and visualizations were performed in Python using standard scientific computing libraries.

### Multi-objective simulation and Pareto analysis of multi-sensor CIC-TF SETs

For the panel-level analysis in Fig. 7B, nine CIC LBDs (LasR_177_, RpaR_179_, ER_282–595_, DHBR_282–595_, acVHH, CinR_179_, PR_645–914_, BjaR ^S107R^ and LuxR ^S116Y^) were assigned to nine non-redundant DBD/operator modules selected from an 11-member DBD pool. For each DBD–LBD pair, the full CIC mass-conservation model was evaluated at 512 logarithmically spaced TF input levels between 0.2 and 10 RPU using LBD-specific *K_d_*_0_, *K_b_* and *K_d_*_1_, DBD-specific *K_A_* and *T*_0_, and *T*_max_ = 35.85 RPU. DHBR_282–595_ was evaluated at 100 μM DHB. Up to six representative input states were retained for each DBD–LBD pair, and 15,000 nine-sensor SETs were reproducibly sampled with unique DBD assignments (random seed, 20260808). For each SET, the objectives were mean log_10_(*Δ*Exp), mean log_10_(*FC*), minimum script-defined OI, total TF input and maximum leakage, with the first three maximized and the latter two minimized. Pareto-nondominated SETs were identified across these five objectives. A balanced representative was selected using *J* = 0.27*z_Δ_*_Exp_ + 0.27*z_FC_* + 0.27*z_OI_* − 0.095*z*_input_ − 0.095*z*_leak_. The experimentally implemented nine-CIC architecture was additionally evaluated at its implemented TF input vector using the same model. Full definitions, pair-state reduction and orthogonality calculations are provided in Supplementary Note 8.4.

### Mammalian endpoint parameter estimation and design-space optimization

Mammalian CIC-TF parameters were inferred from stable-cell OFF/ON endpoint measurements using a constrained modular transfer function model. Each measured construct consisted of one DBD, one LBD and VP16. Reporter output was measured at three calibrated TF input levels under uninduced and induced conditions. The OFF state was defined by *c_I_* = 0, whereas the ON state used the inducer concentration applied in the corresponding stable-cell measurement. For a TF containing DBD (d) and LBD (l), the endpoint model approximated the TF pool competent for DNA binding as a dimeric effective species: 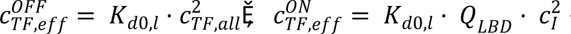 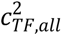 where 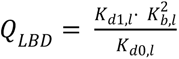The promoter binding weight was then calculated as *Z_d_*_0,*l*_ = *K_A_*_,*d*_ ·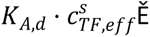 and the predicted output was computed from *T^_d_*_,*l*,*s*_ = *T*_0,*d*_ + *A_d_* · *p_bound_*. Here, *K_A_*_,*d*_ denotes the DBD-associated DNA-binding parameter, *T*_0,*d*_ is the DBD-specific basal output, and *A_d_* is the DBD-specific output amplitude. Thus, the saturated output for DBD (d) is *T*_0,*d*_ + *A_d_*. The main fit constrained TFs sharing the same DBD to share *K_A_*, *T*_0_ and A, and TFs sharing the same LBD to share *K_d_*_0_ and *Q_LBD_*. *K_d_*_0_ was retained as an LBD-specific nuisance parameter because OFF endpoint measurements primarily constrain the product *K_A_* · *K_d_*_0_. The reported LBD-level parameter was 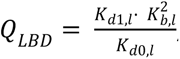, which represents the apparent induced enrichment of the active TF pool relative to the basal state. Because endpoint measurements alone do not uniquely determine the absolute scales of *K_A_* and *K_d_*_0_, the mammalian *K_A_* scale was anchored by fixing lexA_ec87_ to *K_A_* = 9.15. All other DBD-associated *K_A_* values were estimated relative to this anchor. Positive parameters were optimized in log space, and parameters were fitted by nonlinear least squares on the raw output scale. Condition means were fitted with weights proportional to the square root of the number of replicates, equivalent to minimizing replicate-level raw-scale squared error under equal within-condition variance. Model performance was evaluated using raw-scale (R^2^), log-transformed (R^2^), variant-level (R^2^) and residual diagnostics.

For the mammalian design-space analysis in Fig. 6C, the fitted *K_A_* and *Q_LBD_* values were used to position experimentally tested CIC-TF combinations in a two-dimensional landscape. The background landscape was calculated over a grid of *Q_LBD_* and *K_A_* values. For each grid point, *K_d_*_0_, *K_b_*, and *K_d_*_1_ were assigned according to the specified decomposition mode. In the fixed- *K_d_*_1_/*K_d_*_0_ analysis, *K_d_*_1_/*K_d_*_0_ was fixed at 10^5^, and *K_b_* was varied such that 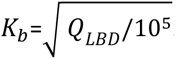. OFF and ON outputs were then calculated using the full dimeric mass-balance model with *c_I_* = 0 and *c_I_* = 100 μM, respectively. For each (*Q_LBD_*, *K_A_*) coordinate, *C_TF,all_* was scanned over the calibrated mammalian expression range of 0.01–3.02 RPU, and the maximum objective value was plotted: 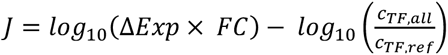 Here, 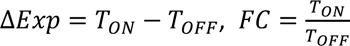 and *c_TF_*_,*ref*_ = 1.0 *RPU*. The resulting landscape therefore represents the best achievable performance at each (*Q_LBD_*, *K_A_*) coordinate after optimizing TF abundance within the experimentally calibrated mammalian input range. Experimentally tested TFs were overlaid according to their OFF/ON-estimated *Q_LBD_* and *K_A_* values, and prioritized designs were selected from measured dose-response-characterized candidates based on the composite score *log*_10_(Δ*Exp* × *FC*).

### Joint mammalian parameter estimation using full dose-response and endpoint data

Mammalian parameters were jointly estimated using full dose-response measurements from 15 CIC-TFs and OFF/ON endpoint measurements from 10 additional CIC-TFs using the full dimeric mass-conservation, promoter-occupancy and transcriptional-output model described in Supplementary Notes 3, 5.3 and 8.3.3 (Eqs. S67, S90 and S94–S99). Both datasets contributed to the final parameter-estimation objective. Variants containing the same LBD shared *K_d_*_0_, *K_b_* and *K_d_*_1_ , whereas variants containing the same DBD shared *K_A_* and *T*_0,variant_ . *K_A_* for LexA_ec87_ was fixed at 9.15, *T*_max_ = 13.61 RPU, and the operator number was fixed at *n* = 7; all remaining *K_A_* and *T*_0,variant_ values were jointly optimized. Positive-valued parameters were estimated in log_10_ coordinates within the bounds specified in Supplementary Note 8.3.3.

Parameter estimation minimized weighted residuals in log_10_(RPU) using bounded scipy.optimize.least_squares (trust-region reflective algorithm; max_nfev = 50,000; ftol = xtol = gtol = 10^−12^; x_scale = ’jac’). Sensor-level weights were initialized to one and adaptively increased for any sensor with log scale *R*^2^ below an internal target of 0.5005. Reweighting continued for at most 24 rounds, and a fit was considered feasible only when every sensor had log scale *R*^2^ ≥ 0.5. Two deterministic 60-parameter initializations were evaluated; among feasible candidates, the parameter set with the smallest pooled unweighted log_10_ mean-squared error was selected. 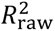 was calculated as a descriptive diagnostic but was not included in the final objective, and no endpoint-shape prior was used.

### Gaussian process-based metabolic optimization

We implemented a Gaussian process (GP)–based iterative optimization workflow to maximize target production in an eight-dimensional regulatory space (LasR_177_, ER_282-595_, PR, XlnR, DHBR_282- 595_, RpaR_179_, LacI, and CinR_179_). All input variables were standardized using a StandardScaler prior to model fitting. The surrogate model was a GP regressor with a radial basis function (RBF) kernel *k*(*x_i_*, *x_j_*) = *C* ⋅ exp(−∥ *x_i_* − *x_j_* ∥^2^/(2*l*^2^)), with hyperparameters optimized using 50 optimizer restarts and a noise term *α* = 10^−3^ . The initial design-of-experiments (DOE) comprised 96 planned induction programs. Owing to errors in inducer dispensing, 32 programs were not implemented as designed and were excluded before model fitting. The remaining 64 correctly executed programs constituted the initial training set (*X*_0_, *y*_0_). In optimization round 1, the fitted GP was used to screen 50,000 uniformly sampled candidates within the predefined bounds; candidates were ranked by predicted mean *μ*(*x*) after filtering by uncertainty *σ*(*x*) < 0.2, and the top 64 were experimentally tested. In optimization round 2, the GP was retrained on all accumulated data (DOE + round 1 measurements) and used for local refinement by perturbing high-performing designs from round 1, *x’_i_* = *x_i_* + *δ* , with element-wise *δ* ∼ *U*(−0.1,0.1), followed by clipping to bounds. Candidates were again evaluated by *μ*(*x*) and *σ*(*x*), filtered with a stricter threshold *σ*(*x*) < 0.15, and the top 64 were selected for experimental validation.

### D-pantothenic acid quantification

The metabolic optimization of vitamin B₅-producing strains was conducted in 2-mL 96-deep-well plates (Axygen, P-2ML-SQ-C-S). Frozen stocks of vitamin B₅-producing strains and reporter strain were streaked onto YPD plates and incubated for 2 days. Single colonies were picked and inoculated into SD medium (with auxotrophic selection) for overnight culture. The cultures were then diluted to an initial OD_600_ of 0.1 and grown for 24 h to reach a saturated cell density state, after which they were transferred to SD medium (with auxotrophic selection) containing 4% glucose and inducers (if required) at an initial OD_600_ of 0.5 and fermented for 96 h. After 16 h of fermentation, 20 µL of reporter culture was mixed with 180 µL PBS for flow-cytometry analysis to quantify gene expression levels. The 96-h fermentation culture from each well was centrifuged at 13,000 × g for 10 min in 1.5-mL tubes. The supernatant was then collected and transferred into sample vials for HPLC analysis. An Agilent HPLC system (Agilent Technologies, USA) equipped with a C18 column (Poroshell 120 EC-C18, 4.6 × 100 mm, 2.7 µm) was used to detect D-pantothenic acid at 200 nm. The elution buffer, composed of water, acetonitrile, and phosphoric acid in a ratio of 950:49:1, was run at a flow rate of 0.6 mL/min.

## Data availability

All data generated or analyzed during this study are included in the supplementary source data files. The nucleotide sequences of synthetic promoters, DBD modules, LBD modules, AD modules and gene sequences involved in vitamin B5 synthesis are provided in Supplementary Tables 9-13. The corresponding transcriptional strength data and vitamin B5 production are available in supplementary source data files.

## Code availability

The custom Python scripts used for data analysis, biophysical modeling, and figure generation during this study are publicly available on https://github.com/CChestnut19/yeast-project.git.

## Acknowledgments

The authors thank Dr. Shuhui Guo and Juhua Du for providing the XlnR sensor, and Dr. Tian Ma for providing the DPA131 strain and vitamin B5 biosynthesis genes.

## Author contributions

Y.C., and S.Z. conceived the study. Y. Z., X. L., G. L., J. X., Z. L., B. S., and Y. C. designed the experiment; Y. Z., X. L., G. L., J. X., Y. Y., and R. W. conducted the experiment; Y. Z., Z. L., X. L., and G. L. analyzed the data; Z. L., B. S., and Y. C. developed the biophysical model; B. W. developed the software GUI; Y. Z., X. L., Z. L., and Y. C. prepared the manuscript; Y.C., S.Z., and F.W. supervised the project. all authors reviewed and approved the final version of the manuscript.

## Declaration of Generative AI and AI-assisted technologies in the manuscript preparation process

During the preparation of this work, the author(s) used ChatGPT (OpenAI) to improve the readability and language of the manuscript and to assist in drafting and debugging code used for data visualization. The author(s) reviewed and edited the output as needed and take full responsibility for the content of the published article.

