## Supplementary information for "Chimeric Induced Cooperativity Opens the Design Space of Eukaryotic Gene Regulation"

### Supplementary Figures

#### Supplementary Figure 1

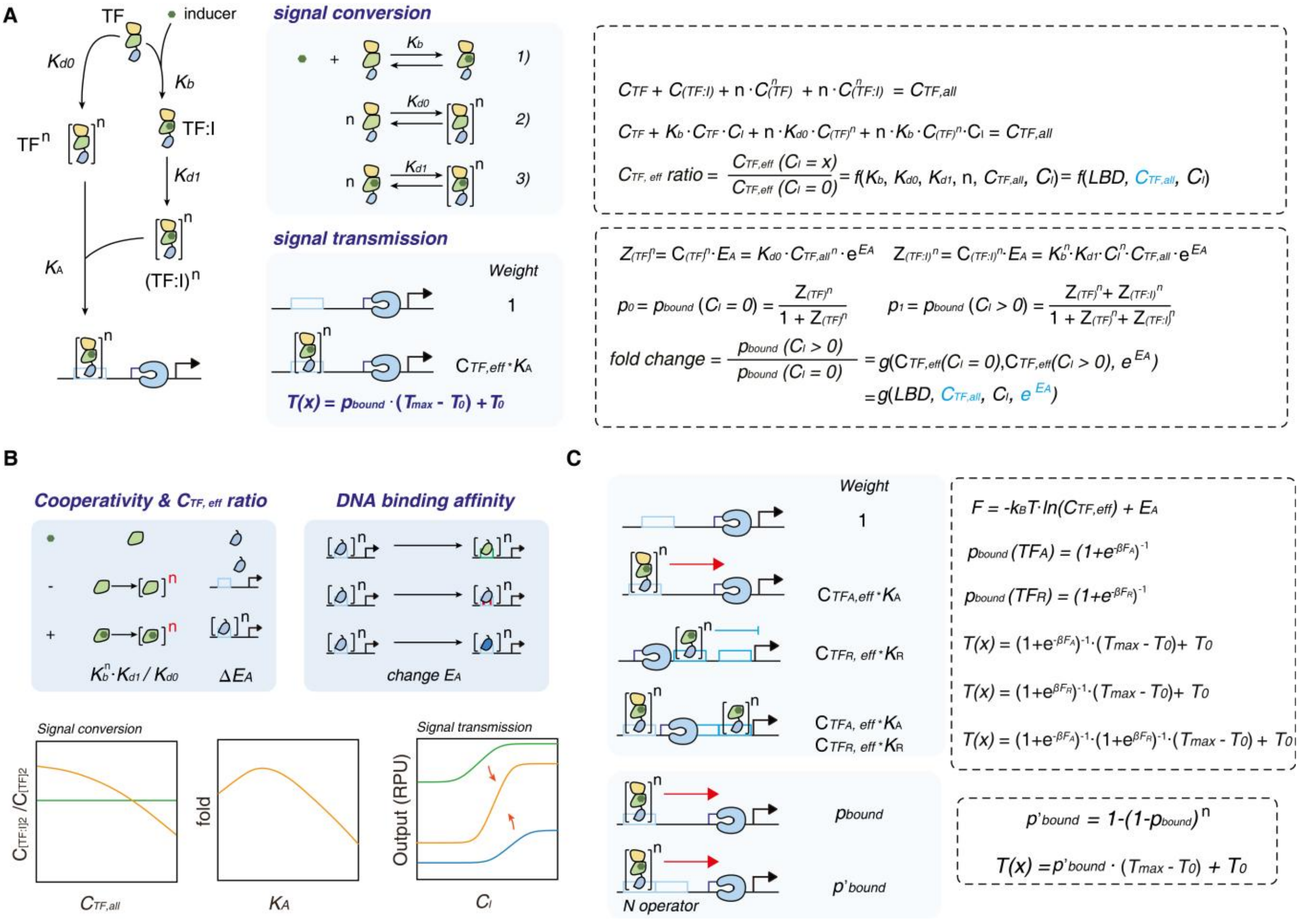

**Supplementary Figure 1. Theoretical framework for CIC-TF-mediated transcriptional regulation. (A)** Biochemical model of CIC-TF signal conversion and signal transmission. Ligand-dependent active-pool formation converts total TF abundance into an apparent DNA-binding-competent effective TF pool, which regulates transcription through DBD–operator binding. Promoter output is modeled as a function of promoter occupancy, and promoter output window defined by $T_0$ and $T_{max}$ ($T(x) = p_{\text{bound}} \cdot (T_{\text{max}} - T_0) + T_0$). **(B)** Key parameters controlling regulatory performance. LBD-associated parameters $K_{d0}$, $K_b$, $K_{d1}$ define the induced-to-uninduced apparent effective TF ratio, while DBD affinity ($K_A$) tunes promoter occupancy and signal transmission. Together, these parameters determine fold-change, output strength, and response shape. **(C)** Partition-function models for activator, repressor, hybrid, and multi-operator promoter architectures. Promoter states are represented by statistical weights, allowing transcriptional output to be predicted from TF occupancy and promoter configuration.

#### Supplementary Figure 2

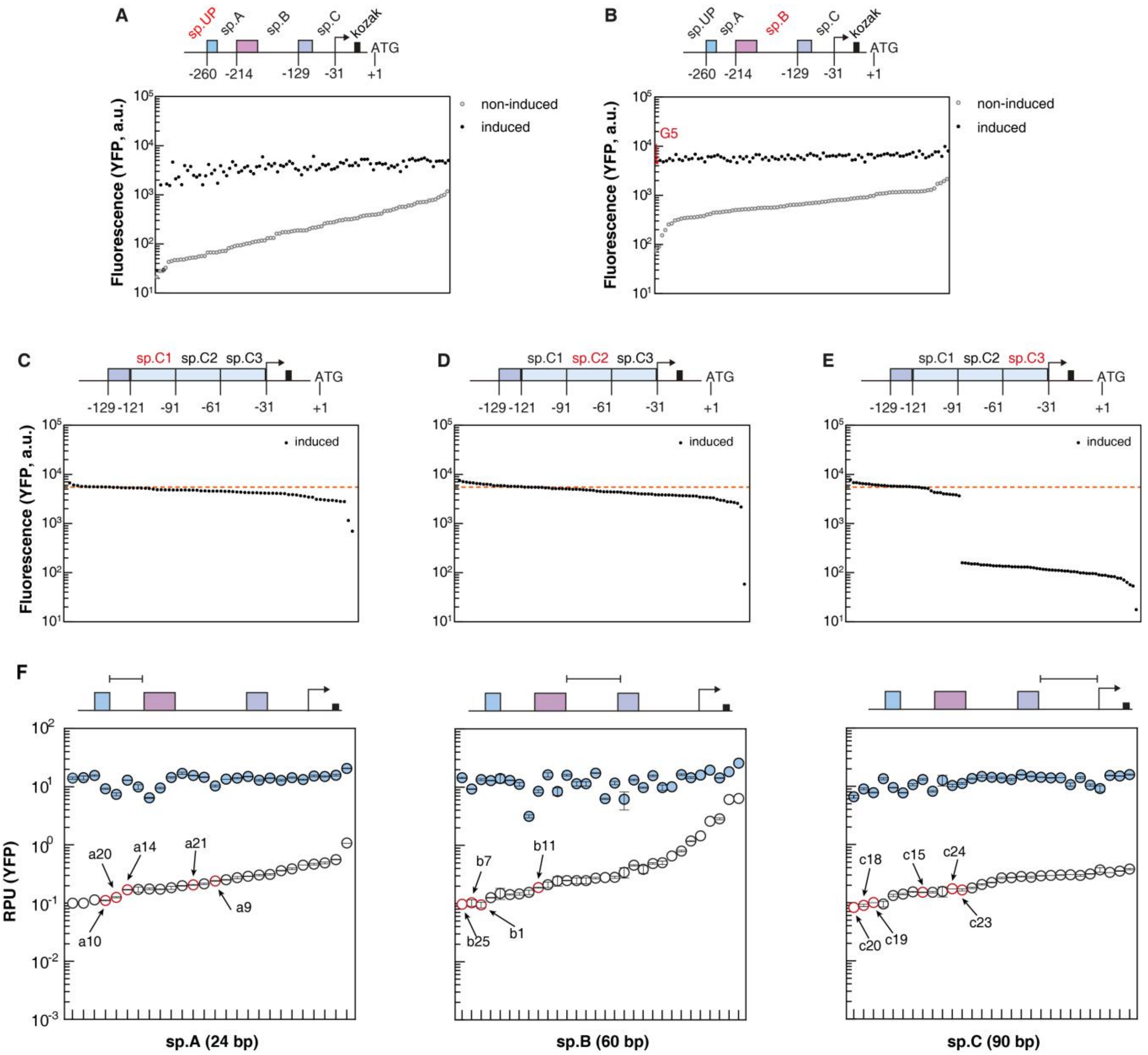

**Supplementary Figure 2. Screening random spacer libraries defines sequence-function relationships in a synthetic $P_{S.a0b1c0\_xlnO}$ promoter. (A-B)** High-throughput screening of random spacer-UP (A) and spacer-B (B) libraries under uninduced and induced conditions to quantify effects on $T_0$ and $T_{max}$. **(C-E)** Segmented random library screening of Spacer C. The full 90-bp Spacer C sequence was subdivided into three equal segments, which are sp.C1 (C), sp.C2 (D), and sp.C3 (E). A random library was constructed for each to assess the effect of sequence variation on $T_{max}$. Data are from a single experiment without biological replicates, as the screen was designed for high-throughput identification of potential hits. **(F)** Libraries of effective sp.A, sp.B and sp.C sequences. Panel F list the sp.A, sp.B and sp.C sequences selected from the screens. The marked spacer was used in the high-fold-change promoters assembled in Fig. 2G. Points represent arithmetic means and error bars indicate sample standard deviations calculated from the available replicate measurements. Biological replicate numbers varied among samples (n = 1–3). No SD was estimated for conditions with n = 1. Missing measurements were not imputed.

#### Supplementary Figure 3

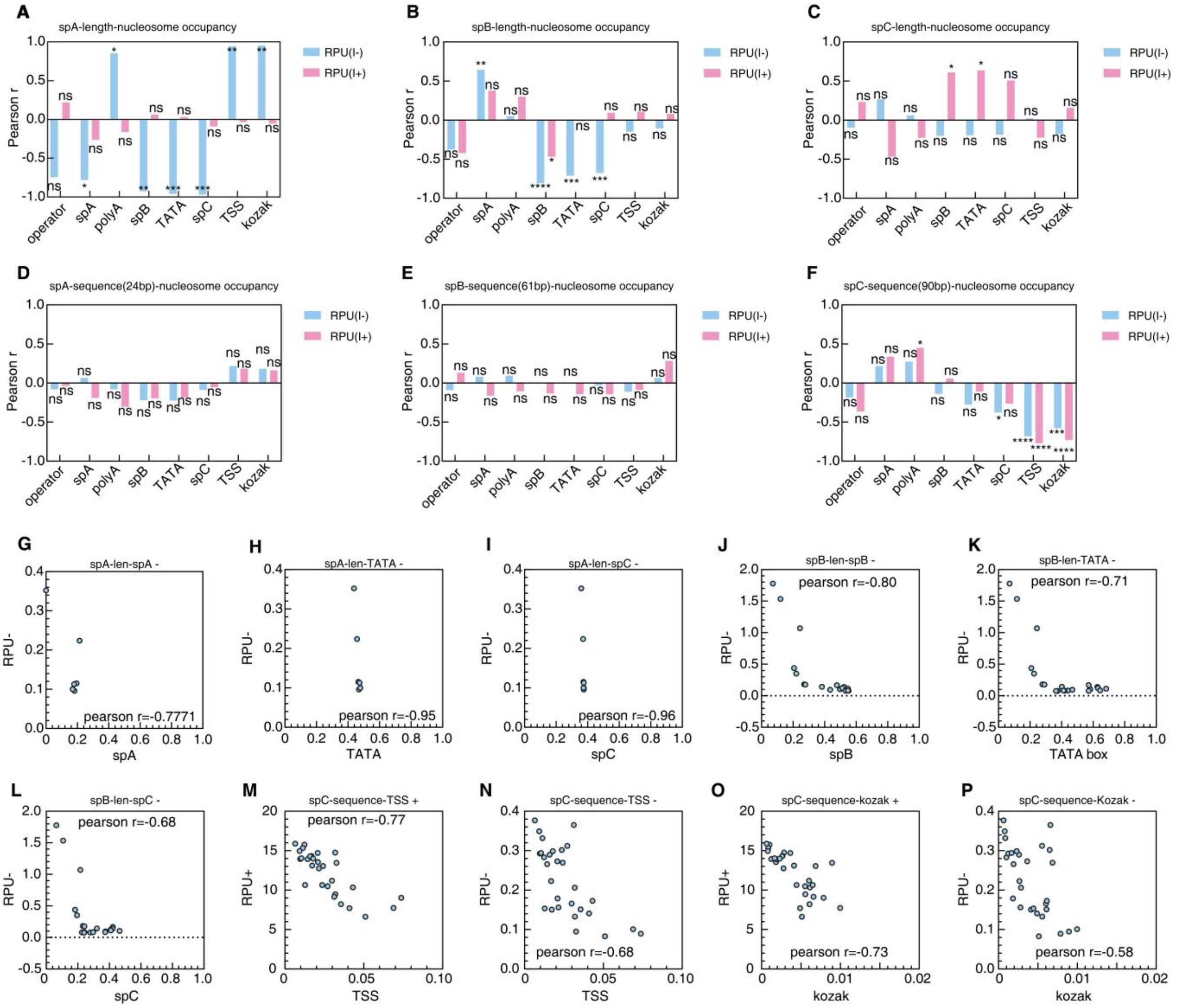

**Supplementary Figure 3. Correlation between nucleosome occupancy and promoter basal strength (*$T_0$*) and maximal output (*$T_{max}$*) in synthetic promoter variants. (A-C)** Overview of the relationship between nucleosome occupancy at the core promoter element and transcriptional output, before (RPU(I-)) and after induction (RPU(I+)), upon incorporation of varying lengths of spA (A), spB (B) and spC (C). **(D-F)** Overview of the relationship between nucleosome occupancy at the core promoter element and transcriptional output, before (RPU(I-)) and after induction (RPU(I+)), upon incorporation of varying sequences of 24-bp spA (D), 61-bp spB (E), and 90-bp spC (F). **(G-I)** Correlation analysis between changes in nucleosome occupancy at the sp.A (G), TATA box (H), and sp.C (I) regions and the induced output RPU(I-), following alterations in sp.A length. **(J–L)** Correlation analysis between changes in nucleosome occupancy at the sp.B (J), TATA box (K), and sp.C (L) regions and the promoter basal output RPU(I-), following alterations in sp.B length. **(M-P)** Spacer C sequence variation influences promoter RPU(I-) and RPU(I+) by modulating nucleosome occupancy at the transcription start site (TSS, M) and Kozak sequence (P). Each point represents one promoter variant. RPU values are means of three independent biological replicates. Pearson correlation coefficients were calculated across promoter variants. Statistical significance was assessed in GraphPad Prism using ordinary simple linear regression with an unconstrained intercept and an F test of the null hypothesis that the slope equals zero (two-sided). P values were not adjusted for multiple correlations. Significance is indicated as follows: ns, $P > 0.05$; *, $P \leq 0.05$; **, $P \leq 0.01$; ***, $P \leq 0.001$; ****, $P \leq 0.0001$.

#### Supplementary Figure 4

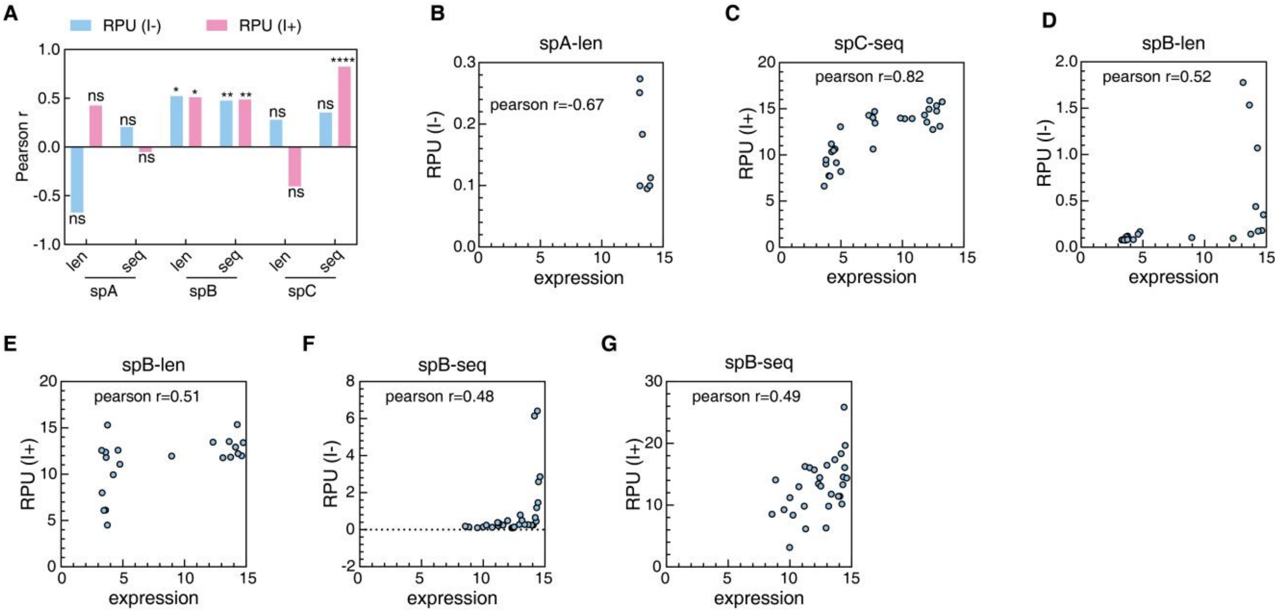

**Supplementary Figure 4. Sequence-function analysis of synthetic promoters based on de Boer and Regev's model[6]. (A)** Overview of the relationship between experimentally determined strength of promoter variants and prediction based on de Boer and Regev's model. **(B)** Correlation between the predicted and measured uninduced strength, RPU(I−), of the promoter variants with different lengths of sp.A. **(C)** Correlation between the predicted and measured induced strength, (RPU(I+)), of the promoter variants with different sequence sp.C (90-bp). **(D-G)** The correlation between predicted and experimental strength of promoter variants with different lengths (D-E) or sequences of sp.B (61-bp, F-G). Predictions were obtained with the de Boer and Regev model. Each point represents one promoter variant. RPU values are means of three independent biological replicates. Pearson correlation coefficients were calculated across promoter variant. Statistical significance was assessed in GraphPad Prism using ordinary simple linear regression with an unconstrained intercept and an F test of the null hypothesis that the slope equals zero (two-sided). P values were not adjusted for multiple correlations. Significance is indicated as follows: ns, $P > 0.05$; *, $P \leq 0.05$; **, $P \leq 0.01$; ***, $P \leq 0.001$; ****, $P \leq 0.0001$.

#### Supplementary Figure 5

**A**

| DBD | operator | Sequence |
|---|---|---|
| lexA*ec*87 | lexO*ec* | TGCTGTATATACTCACAGCA |
| lexA*pa*93 | lexO*pa* | CGCTGTATAAATATACAGTG |
| lexA*vp*91 | lexO*vp* | TAACTGTATAAATAAACAGGAT |
| lexA*pp*91 | lexO*pp* | ATACTGTATATATATACAGTCA |
| lexA*mt*127 | lexO*mt* | TATCGAACACATGTTCGAGA |
| lexA*cg*145 | lexO*cg* | TTCGAACAAATGTTCGA |
| lexA*sa*96 | lexO*sa* | CGAACATATGTTCT |
| lexA*bs*94 | lexO*bs* | AGAACATATGTTCG |
| lexA*pd*114 | lexO*pd* | AGGAACTAATGTTCTG |
| lexA*lm*93 | lexO*lm* | TATAAGAACGTATGTTCGGATA |
| lexA*sm*126 | lexO*sm* | CTGTTCTTGATTTGTTCTA |
| lexA*cc*123 | lexO*cc* | TATGTTCTCGTTATGTTCT |

**B**

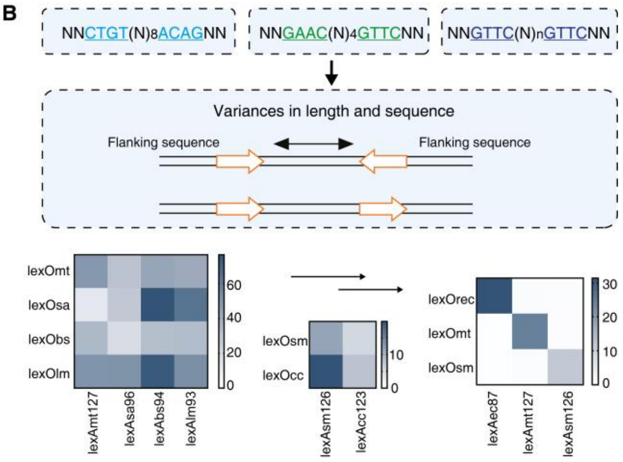

**Supplementary Figure 5. DBD-operator orthogonality screening based on half-site architecture. (A)** Profiles of lexA half-site sequences across closely related bacterial strains. **(B)** Orthogonality test grouped by half-site. Data points represent biological replicates (n = 3).

#### Supplementary Figure 6

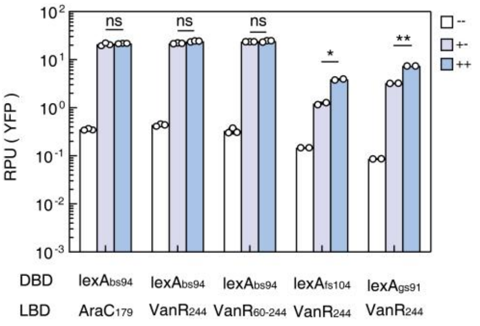

**Supplementary Figure 6. Other functional LBD screening in yeast.** The symbols denote three induction states: "--" indicates no inducer added; "+-" indicates induction of transcription factor expression alone, whereas "++" indicates induction of expression together with the addition of the corresponding ligand that induces transcriptional activation. Data points represent biological replicates (n = 3). Two-way ANOVA with Tukey's post hoc test was performed to analyze the effects of ligand addition. Significance is indicated as ns ($p>0.05$), * ($p < 0.05$), ** ($p < 0.01$), and *** ($p < 0.001$).

#### Supplementary Figure 7

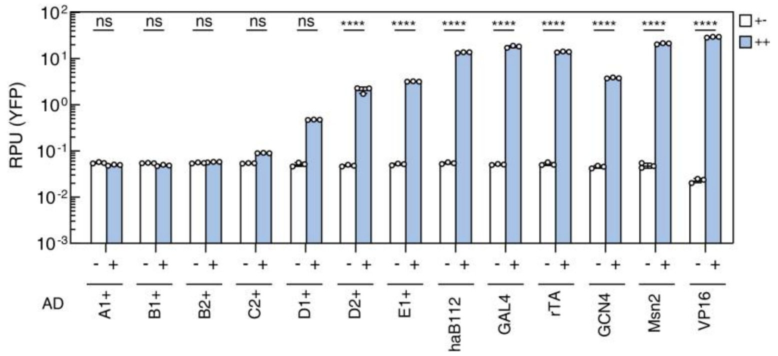

**Supplementary Figure 7. Activating domain screening.** Each activation domain (AD) was fused to the $LexA_{ec87}$ DNA-binding domain and the $RpaR_{179}$ ligand-binding domain to assemble synthetic transcription factors. The transcription factor was placed under the control of the $P_{tet}$ promoter and induced by the addition of anhydrotetracycline (aTc). Upon subsequent addition of pC-HSL ligand, the transcription factor dimerizes and activates expression of YFP. "+–" indicates induction of transcription factor expression alone, whereas "++" indicates induction of expression together with ligand-induced transcription factor dimerization. Data represent the mean ± SD from three biological replicates. Statistical analysis was performed using two-way ANOVA followed by Bonferroni's multiple comparisons test to evaluate the effects of ligand addition. Significance is indicated as ns ($p > 0.05$), * ($p < 0.05$), ** ($p < 0.01$), *** ($p < 0.001$), and **** ($p < 0.0001$).

#### Supplementary Figure 8

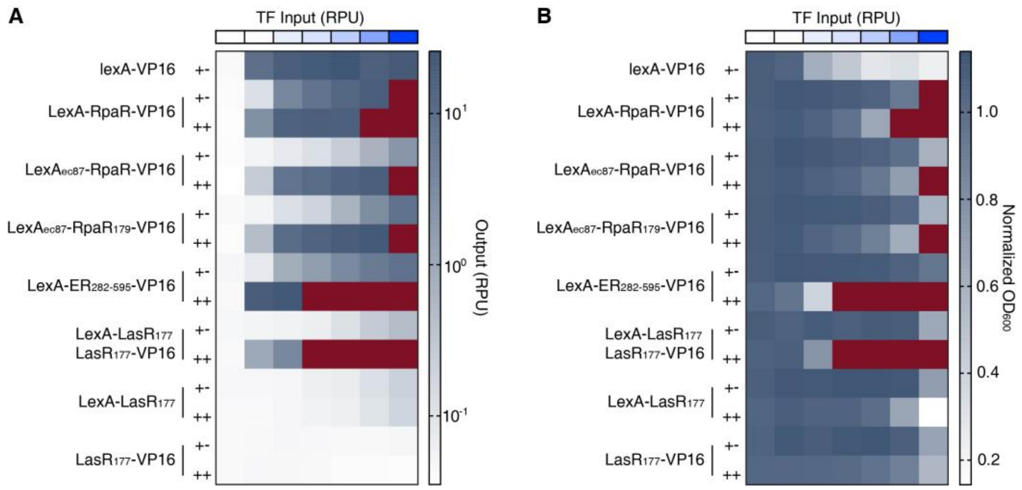

**Supplementary Figure 8. Effects of chimeric TF architectures on transcriptional output and cell growth. (A)** Transcriptional output of the indicated chimeric transcription factors across seven xylose-defined TF-input conditions, shown as output relative promoter units (RPU) on a logarithmic color scale. **(B)** Culture density under the corresponding conditions, shown as $OD_{600}$ normalized to the matched day-specific BY4741 pgs093int strain. Rows marked +− and ++ denote non-induced and induced conditions, respectively. All chimeric TFs were tested at the same xylose concentrations of 0, 0.4, 0.6, 0.8, 1.0, 1.5, and 5.0 mM. Based on the calibration of the xylose-inducible promoter $P_{m115\_xlnO}$, these concentrations correspond to nominal TF-input levels of 0.00, 0.24, 3.32, 5.87, 10.46, 19.50, and 40.74 RPU, respectively. Batch-specific measured TF-input values are provided in the source data. Each heatmap cell represents the arithmetic mean of six measurements obtained from three independent experiments performed on different days, with two technical replicates per experiment.

#### Supplementary Figure 9

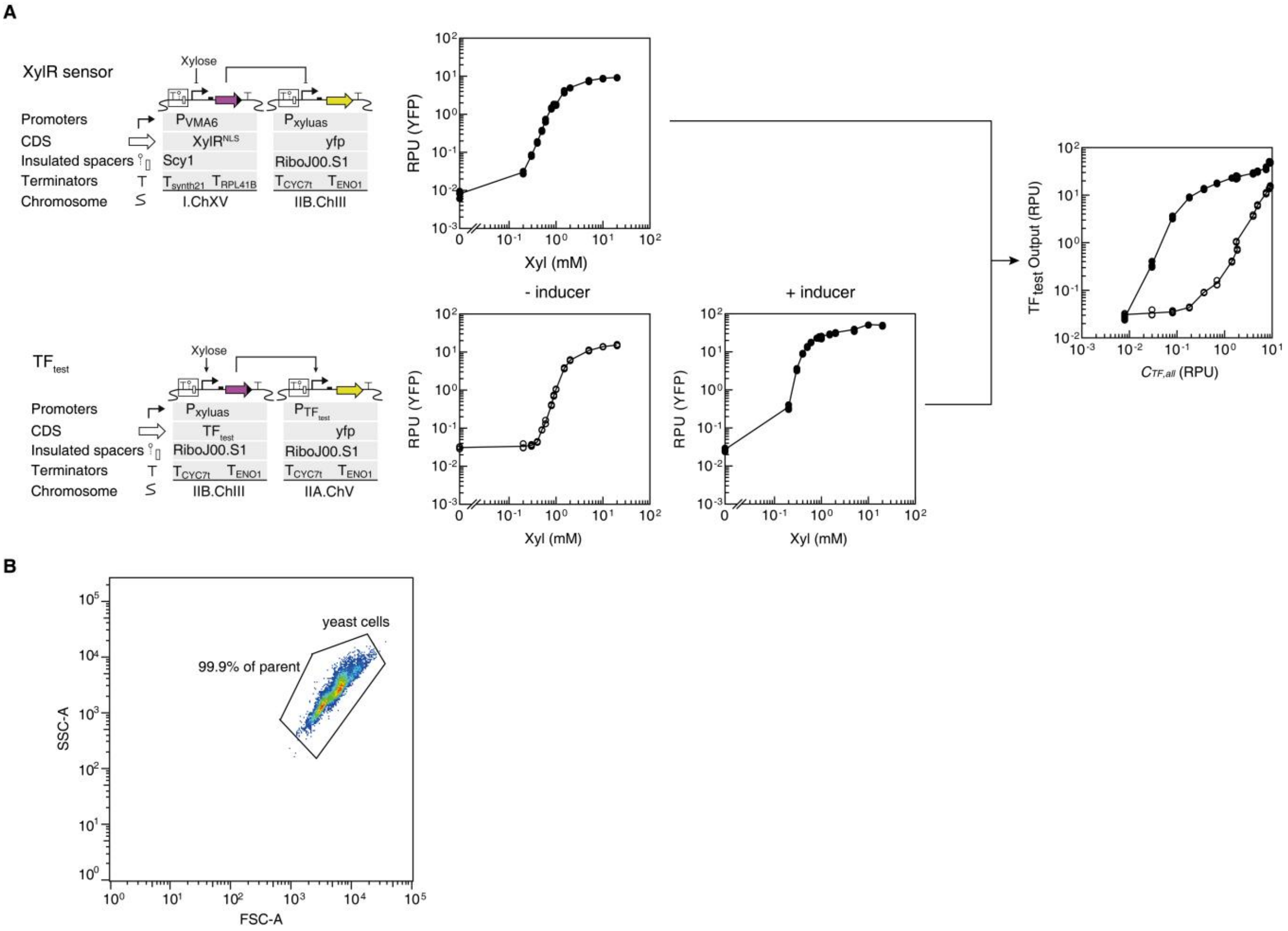

**Supplementary Figure 9. Quantitative testing and flow-cytometry analysis of CIC-TFs in yeast. (A)** The transcription factor under investigation ($TF_{test}$) is expressed from the $P_{xyluas}$ promoter, which is repressed by XylR and inducible by xylose. In a reference strain, $TF_{test}$ is replaced with yEmCitrine, enabling quantitative measurement of TF input levels (in RPUs) at defined xylose concentrations. RPU values are calculated as previously described. **(B)** Yeast gating strategy. Yeast cells were identified using a two-dimensional gate based on forward-scatter area (FSC-A) and side-scatter area (SSC-A). Fluorescence measurements were calculated from events within this yeast-cell gate. The same gating strategy was applied across experimental conditions.

#### Supplementary Figure 10

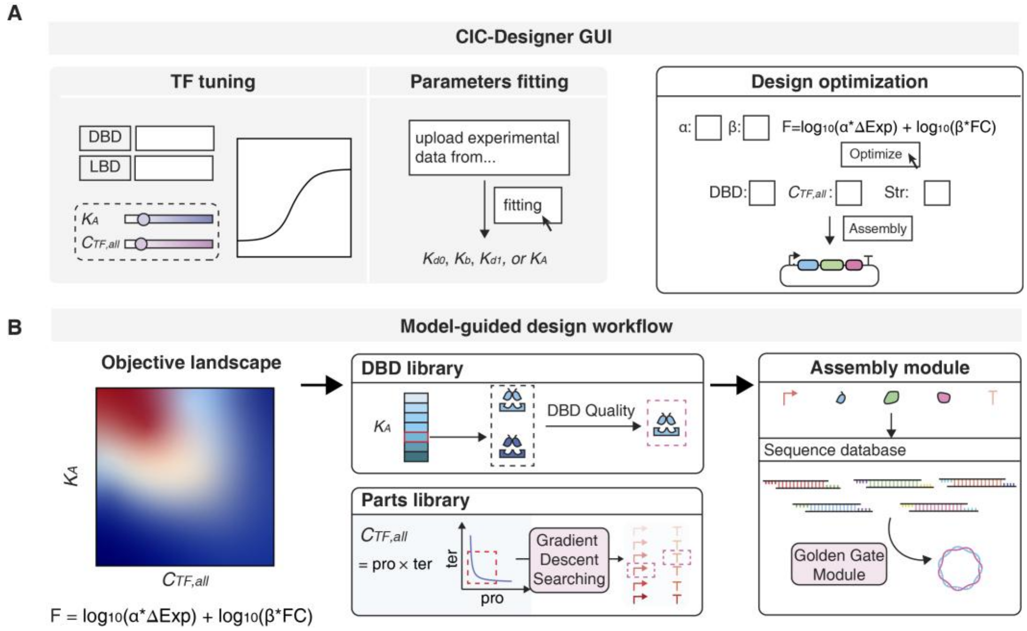

**Supplementary Figure 10. CIC-Designer workflow for model-guided CIC-TF optimization and assembly.** **(A)** Graphical interface for TF tuning, domain-parameter fitting, and user-defined design optimization. Users can adjust $C_{TF,all}$ and $K_A$, upload experimental datasets to fit new domain parameters, and specify objective weights and design constraints $K_A$, $C_{TF,all}$. **(B)** Computational workflow for CIC-TF design. A user-defined objective function is used to search the model-predicted landscape and identify optimal DBD affinity and TF abundance. These targets are mapped to characterized DBDs and promoter–terminator parts, after which the assembly module retrieves DNA parts from the sequence database and returns a candidate construct selected under the specified objective and search constraints.

#### Supplementary Figure 11

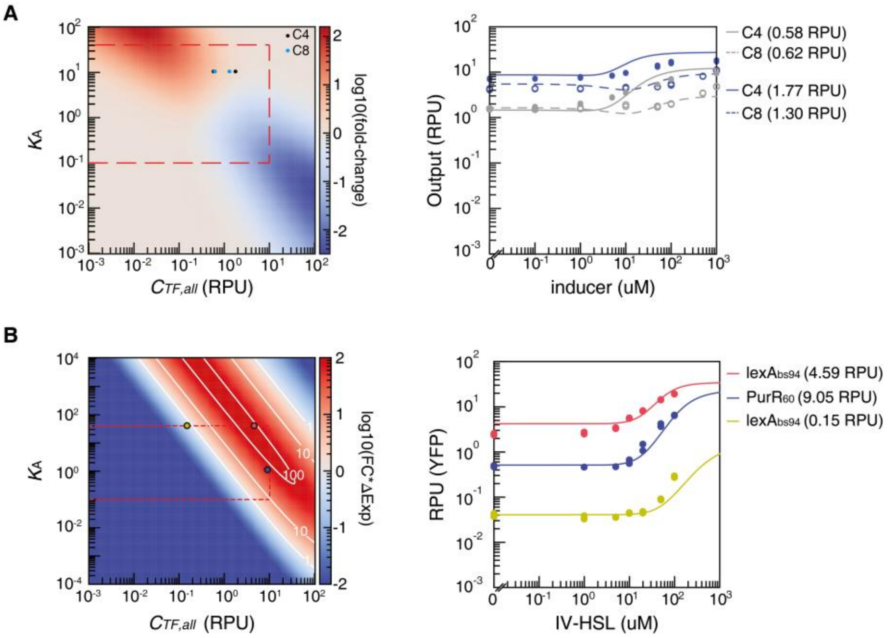

**Supplementary Figure 11.** Model-guided objective optimization for crosstalk reduction and sensor performance enhancement. **(A)** Model-guided design strategy for reducing crosstalk in $SmaR_{179}$-based CIC-TFs. Left, predicted design landscape across $C_{TF,all}$ and DBD affinity ($K_A$), with color indicating $\log_{10}$fold-change. The dashed red box marks the experimentally accessible design region. Points indicate selected designs for experimental validation. Right, measured dose-response curves for selected C4- and C8-HSL-responsive designs. Lines indicate model predictions, and points indicate experimental measurements. **(B)** Model-guided optimization of $BjaR_{180}$ sensor performance by jointly maximizing fold-change (FC) and expression change (ΔExp) Left, predicted objective landscape across $C_{TF,all}$ and $K_A$, with color indicating $\log_{10}(FC \times \Delta Exp)$. White contour lines indicate objective values. Points mark selected DBD–$C_{TF,all}$ designs for validation. Right, measured IV-HSL dose-response curves for the selected designs. Lines indicate model predictions, and points indicate experimental measurements. Data points represent biological replicates (n = 2).

#### Supplementary Figure 12

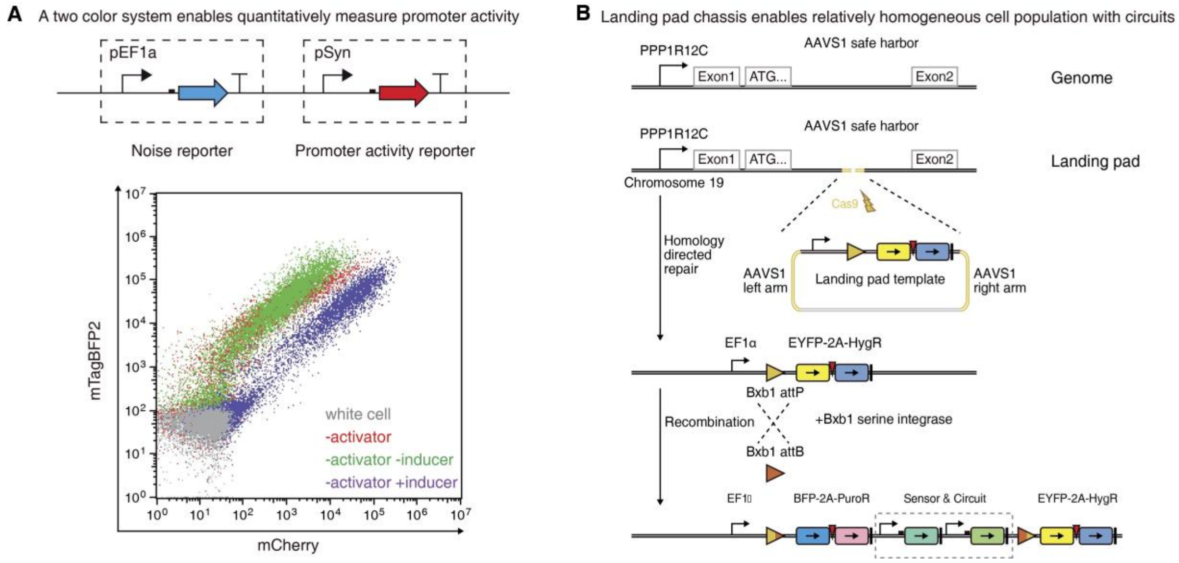

**Supplementary Figure 12. Quantitative characterization of sensor performance in mammalian cells. (A)** Characterization of promoter transcriptional strength using a dual-fluorescence reporter system. The blue fluorescent protein mTagBFP2 is expressed under the constitutive promoter pEF1α, serving as a normalization reference, while mCherry is placed downstream of a synthetic promoter responsive to the target transcription factor (TF). In flow cytometry plots, the negative control is shown as gray dots. Cells with successful circuit integration are highlighted as the green population. Upon inducer addition, the green population shifts to the purple population, indicating inducible promoter activity. During flow cytometry analysis, cells exhibiting blue fluorescence (mTagBFP2) are gated as circuit-containing cells in scatter plots with mCherry fluorescence on the x-axis and mTagBFP2 on the y-axis. **(B)** Schematic of the integration process for sensor testing circuits using a landing pad chassis. Constructs were inserted into a predefined genomic landing pad, allowing for uniform expression of the integrated circuits in the resulting cell population.

#### Supplementary Figure 13

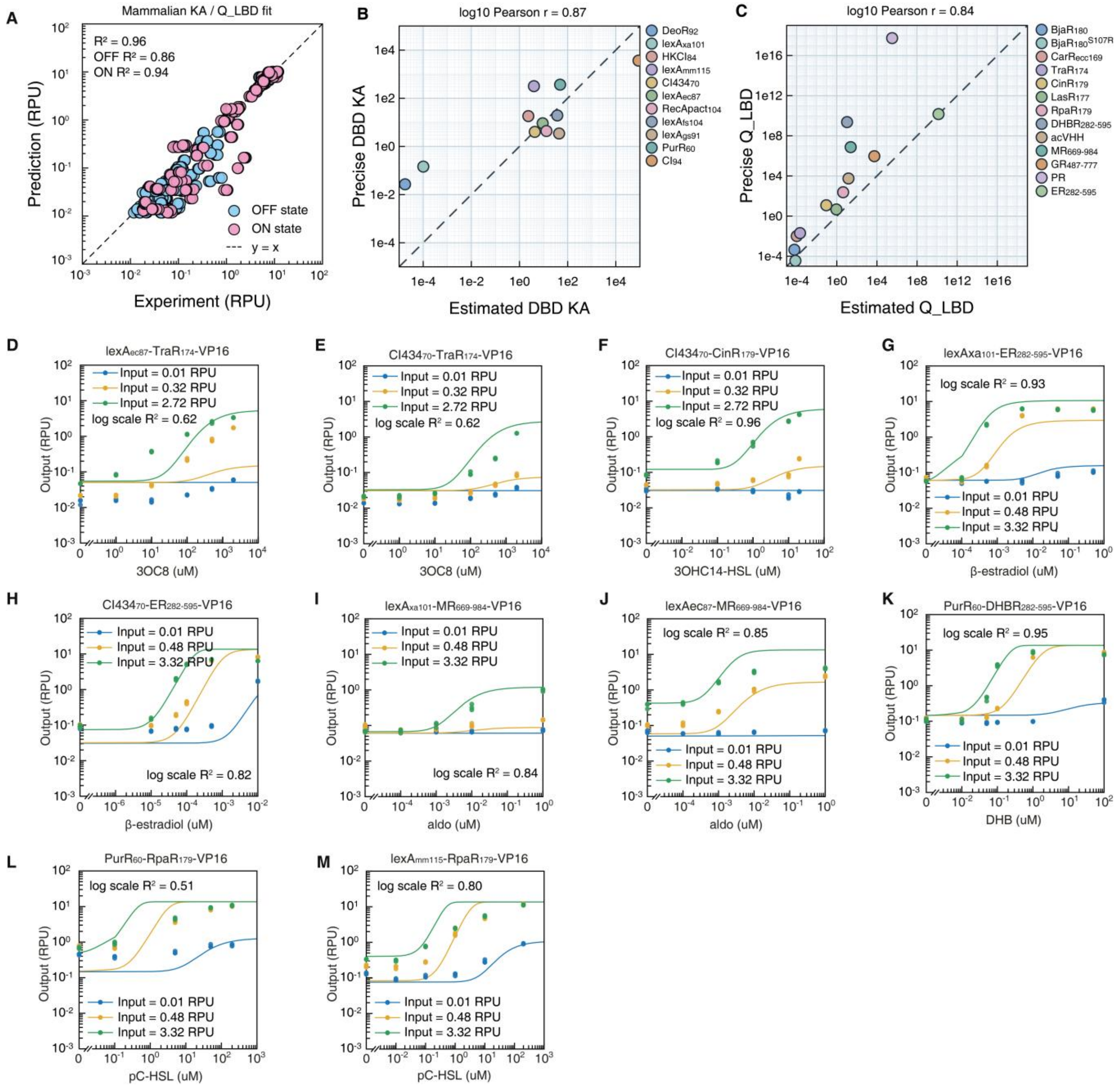

**Supplementary Figure 13. Model calibration and fit assessment of modular transcription factor circuits in mammalian cells.** (A) Experiment versus prediction plot for mammalian CIC-TFs using parameters estimated from sparse OFF/ON measurements. DBD-associated $K_A$ and LBD-associated $Q_{LBD} = K_b^2 \times K_{d1} / K_{d0}$ were inferred from OFF/ON data and then used in the full CIC-TF dimerization transfer model to predict reporter output. Each point represents one measured OFF or ON output value. Blue points indicate OFF-state measurements, and pink points indicate ON-state measurements. The dashed line denotes y = x. Model performance is shown as raw $R^2$ for all data, OFF-state data, and ON-state data. (B) Comparison of DBD binding-strength parameters estimated from sparse OFF/ON data versus parameters obtained from full induction-curve fitting. Each point represents one DBD module. Both axes are plotted on a log scale. The dashed line denotes y = x. Pearson correlation was calculated in log10-transformed parameter space. (C) Comparison of LBD induction-potential parameters estimated from sparse OFF/ON data versus parameters obtained from full induction-curve fitting. $Q_{LBD}$ is defined as $K_{d1} \times K_b^2 / K_{d0}$. Each point represents one LBD module. Both axes are plotted on a log scale. The dashed line denotes y = x. Pearson correlation was calculated in log10-transformed parameter space. Panels D–M represent the transcription factors lexA$_{ec87}$-TraR$_{174}$-VP16, CI434$_{70}$-TraR$_{174}$-VP16, CI434$_{70}$-CinR$_{179}$-VP16, lexA$_{xa101}$-ER$_{282-595}$-VP16, CI434$_{70}$-ER$_{282-595}$-VP16, lexA$_{xa101}$-MR$_{669-984}$-VP16, lexA$_{ec87}$-MR$_{669-984}$-VP16, PurR$_{60}$-DHBR$_{282-595}$-VP16, PurR$_{60}$-RpaR$_{179}$-VP16, lexA$_{mm115}$-RpaR$_{179}$-VP16 respectively. The expression of each transcription factor is driven by the $P_{lac}$ promoter. TF input levels were quantified in relative promoter units (RPUs). Experimental data are shown as dots for three input levels, with corresponding model predictions shown as curves in the same colors. For panels **D–F**, the input levels are: blue, 0.01 RPU; yellow, 0.32 RPU; green, 2.72 RPU. For panels **G–M**, the input levels are: blue, 0.01 RPU; yellow, 0.48 RPU; green, 3.32 RPU. Each condition includes three biological replicates, all of which are individually plotted; in some cases, overlapping values may appear as a single dot.

#### Supplementary Figure 14

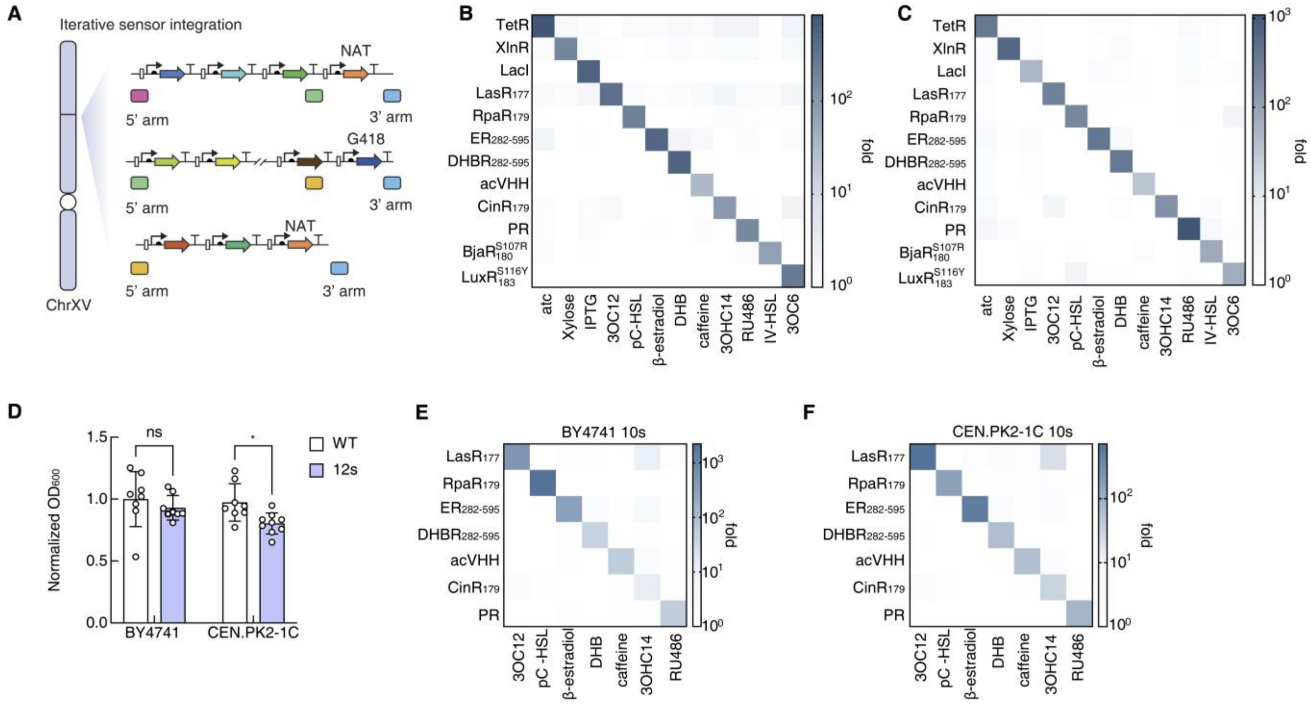

**Supplementary Figure 14. Crosstalk and growth evaluation of CIC-12 in BY4741 and CEN.PK2-1C strain backgrounds. (A)** Iterative chromosomal integration strategy for assembling the multi-sensor chassis. Sensor expression cassettes were sequentially integrated at the ChrXV locus using alternating selectable markers, NAT and G418, flanked by homologous arms for iterative marker replacement and expansion of the sensor array. **(B–C)** Assessment of transcription factor orthogonality in BY4741 12s (B) and CEN.PK2-1C 12s (C) chassis strains. Orthogonality characterization was performed using the lowest tested cognate-inducer concentration that produced a near-saturating response for each sensor. The concentrations used were as follows: 100 ng $mL^{-1}$ for aTc, 10 mM for xylose, 20 mM for IPTG, 1 µM for 3OC12-HSL, 50 µM for pC-HSL, 1 µM for β-estradiol, 2.5 µM for DHB, 1 µM for caffeine, 100 nM for 3OHC14-HSL, 25 µM for RU486, 100 µM for IV-HSL, and 100 µM for 3OC6 (1 mM 3OC6 for CEN.PK2-1C 12s chassis strain). **(D)** Growth evaluation of the BY4741 12s and CEN.PK2-1C 12s strains. Data are presented as individual values with error bars indicating mean ± SD from three independent experiments performed on different days. In each experiment, three distinct single colonies were cultured and measured independently, n = 3 independent clones per condition, yielding nine culture measurements per condition. Normalized $OD_{600}$ values were analyzed using a two-factor model with strain background and 12-sensor integration status as factors. Pre-specified comparisons between WT and 12-sensor strains within each strain background were performed using Šídák's multiple-comparisons test. *, $P < 0.05$; ns, not significant. **(E–F)** Crosstalk evaluation in BY4741 10s (E) and CEN.PK2-1C 10s (F) chassis strains based on non-repetitive reporters. The concentrations used were as follows: 100 ng $mL^{-1}$ for aTc, 10 mM for xylose, 20 mM for IPTG, 1 µM for 3OC12-HSL, 50 µM for pC-HSL, 1 µM for β-estradiol, 2.5 µM for DHB, 1 µM for caffeine, 1 µM for 3OHC14-HSL, 100 µM for RU486.

#### Supplementary Figure 15

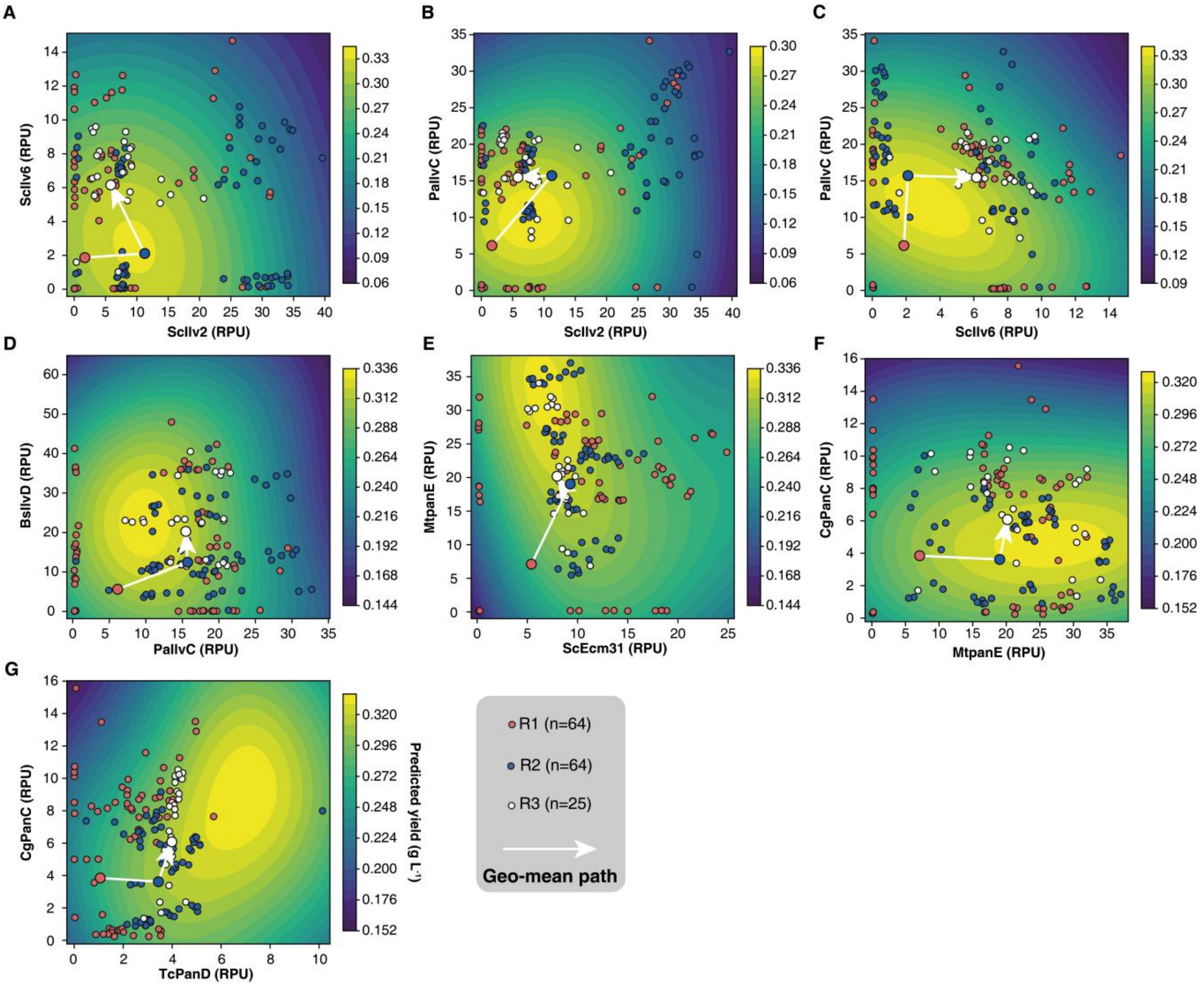

**Supplementary Figure 15. Gaussian-process-predicted yield landscapes across pairwise gene-expression dimensions.** (A–G) Two-dimensional projections of the Gaussian-process (GP)-predicted metabolic-yield landscape for the indicated pairs of gene-expression/RPU dimensions: (A) ScIlv2 versus ScIlv6, (B) ScIlv2 versus PaIlvC, (C) ScIlv6 versus PaIlvC, (D) PaIlvC versus BsIlvD, (E) ScEcm31 versus MtPanE, (F) MtPanE versus CgPanC, (G) TcPanD versus CgPanC. The GP was fitted using 153 complete, non-starred conditions retained across the three optimization rounds, comprising 64 conditions in Round 1, 64 conditions in Round 2, and 25 conditions in Round 3. For each projection, the two indicated gene-expression dimensions were varied over the displayed range, whereas the remaining six dimensions were fixed at their pooled mean values across the 153 training conditions. The colour scale represents the GP-predicted yield (g $l^{-1}$). Each circle represents one tested condition; red, blue and white circles denote Rounds 1, 2 and 3, respectively. The white line connects the coordinate-wise geometric means of Rounds 1, 2 and 3, with each path vertex coloured according to its corresponding round.

### Supplementary Notes

#### Supplementary Note 1: Definition of the Partition Function and Statistical Thermodynamic Model

To quantitatively describe transcriptional regulation mediated by activators and repressors, we adopt a simple thermodynamic modeling framework. In this framework, the promoter is assumed to exist in a set of mutually exclusive microscopic configurations, each corresponding to a distinct combination of transcription factor (TF) binding states. Instead of explicitly modeling RNA polymerase (RNAP) binding, we define the transcriptional output as being proportional to the probability that the promoter occupies a transcriptionally permissive microscopic state.

##### Supplementary Note 1.1: Definition of parameters

We define $p_{\text{bound}}$ as the probability that the promoter adopts a configuration that is competent for transcription initiation. This "transcriptionally active" configuration is implicitly determined by the presence of the activator and the absence of the repressor and does not require RNAP itself to be explicitly included in the statistical weight calculation. Under this simplified representation, TF recruitment or inhibition of RNAP is incorporated into the effective binding constants describing TF–promoter interactions.

Here we define $K_A$, $K_R$ represents the effective Boltzmann weight associated with activator binding or repressor binding, rather than a dissociation constant.

$$\Delta G = G_{bound} - G_{unbound} \tag{S1}$$

$$E_A = -\Delta G_A \tag{S2}$$

$$E_R = -\Delta G_R \tag{S3}$$

$$K_A = e^{-\frac{\Delta G_A}{k_B T}} \tag{S4}$$

$$K_R = e^{-\frac{\Delta G_R}{k_B T}} \tag{S5}$$

Where $\Delta G_A$ and $\Delta G_R$ are the effective free energy changes associated with activator and repressor binding, $E_A$ and $E_R$ is the binding energy of activator and repressor. $k_B$ is the Boltzmann constant, and $T$ is absolute temperature.

##### Supplementary Note 1.2: Statistical Weights of Microscopic States

The statistical weight of each promoter state is defined according to Boltzmann statistics. Assuming that only active TF could bind to promoter, and activator-bound and repressor-bound states contribute non-zero weight, we write:

$$Z_A = K_A \cdot c_{TF_A,eff} \quad (S6)$$

$$Z_R = K_R \cdot c_{TF_R,eff} \quad (S7)$$

where $c_{TF_A,eff}$ and $c_{TF_R,eff}$ represent the concentrations of the active forms of the activator and repressor.

The unbound promoter state is assigned a statistical weight of 1. The total partition function is therefore:

$$Z_{total} = 1 + Z_A + Z_R + Z_A \cdot Z_R \quad (S8)$$

$$Z_{total} = 1 + K_A \cdot c_{TF_A,eff} + K_R \cdot c_{TF_R,eff} + K_A \cdot c_{TF_A,eff} \cdot K_R \cdot c_{TF_R,eff} \quad (S9)$$

**Supplementary Note 1.3: Transcriptional Output Function**

Under this model, the promoter is transcriptionally active only when the activator is bound and the repressor is not bound. Because these states are mutually exclusive, the probability of being in the permissive state equals:

$$p_{bound} = \frac{Z_A}{Z_{total}} = \frac{K_A \cdot c_{TF_A,eff}}{1 + K_A \cdot c_{TF_A,eff} + K_R \cdot c_{TF_R,eff} + K_A \cdot c_{TF_A,eff} \cdot K_R \cdot c_{TF_R,eff}} \quad (S10)$$

And the transcriptional output is modeled as a linear function of $p_{\mathrm{bound}}$. Specifically:

$$T(x) = p_{bound} \cdot (T_{max} - T_0) + T_0 \quad (S11)$$

Where: $T_{max}$ is the maximal transcriptional capacity of the promoter, $T_0$ represents the basal transcription level in the absence of activation. This formulation is consistent with thermodynamic models widely used in bacterial and synthetic promoter modeling[1,8].

**Supplementary Note 1.4: Quantification of transcriptional factor concentration**

Direct and accurate quantification of the absolute intracellular concentration of transcription factors (TFs) in units such as micromolar or per-cell molecular counts remains technically challenging. Existing approaches are frequently labor-intensive, require substantial technical expertise, and introduce considerable quantitative variability. To address these constraints, we employed a widely accepted relative quantification framework that infers TF abundance from the expression of fluorescent reporter genes. TF levels were expressed in relative promoter units (RPU), a dimensionless metric normalized to an internal reference standard. Fluorescence signals were first corrected for variations in cell density through optical-density normalization, and RPU values were subsequently calculated relative to a designated reference strain.

$$RPU = \frac{Fluorescence_{test} - Fluorescence_{Blank}}{Fluorescence_{Ref} - Fluorescence_{Blank}} \quad (S12)$$

In our mathematical framework, the total transcription factor concentration is denoted as $c_{TF,all}$. To enable a direct correspondence between the model and experimental measurements, $c_{TF,all}$ is defined as the relative abundance of monomeric TF proteins quantified in relative promoter units (RPU). Under this convention, $c_{TF,all} = 1$ represents a TF expression level equivalent to that of the reference strain, $c_{TF,all} = 2$ reflects an approximately twofold increase, and higher values scale accordingly. This formulation allows model parameters to be calibrated and validated directly against experimentally reproducible RPU measurements, thereby circumventing the need for absolute protein quantification, which is technically demanding and often inaccessible.

#### Supplementary Note 2: Evaluation of Type I, Type II transcription factors from first principles.

Inducible transcriptional regulation systems can be broadly grouped into three types: Type I allosteric transcription factors, Type II ligand-induced heterodimerization systems, and Type III ligand-induced cooperative DNA-binding systems. To determine which class provides the greatest design space for future synthetic TF engineering, we developed minimal models for each mechanism and evaluated their tunability and modularity.

##### Supplementary Note 2.1: Allosteric TF based transcriptional regulation.

For Type I allosteric transcription factors, the model assumes an equilibrium between an active DNA-binding conformation and an inactive, low-affinity conformation. Ligand binding shifts this equilibrium by selectively stabilizing one of these states. The probability that the TF is in the active state is obtained by summing the statistical weights of all active configurations and normalizing by the total allosteric partition function. This formulation allows quantitative prediction of activation or repression depending on whether ligand binding favors the inactive or the active state. We established the Type I TF model based on Razo and Phillips's work[1].

At the promoter level, transcriptional output is computed using the same thermodynamic model introduced earlier, in which promoter binding is governed by the Boltzmann-weight parameter $K_A$ that quantifies the promoter affinity of the active TF: $T(x) = p_{bound} \cdot T_{max} + T_0$

And the statistical weight of active TF ($Z_A$) and inactive TF ( $Z_I$) is written as:

$$Z_{Active} = 1 + \left(c_I \cdot K_{L,A}\right)^n \tag{S13}$$

$$Z_{Inactive} = e^{\beta \Delta \varepsilon AI} \cdot \left(1 + (c_I \cdot K_{L,I}\right)^n \tag{S14}$$

$$p_{active} = \frac{Z_{Active}}{Z_{Active} + Z_{Inactive}} = \frac{1 + \left(c_I \cdot K_{L,A}\right)^n}{1 + \left(c_I \cdot K_{L,A}\right)^n) + e^{\beta \Delta \varepsilon AI} \cdot \left(1 + (c_I \cdot K_{L,I}\right)^n)} \tag{S15}$$

$$c_{TF,eff} = p_{active} \cdot c_{TF,all} \tag{S16}$$

For activation, combining S6 and S16 equations we would have:

$$p_{bound} = \frac{Z_A}{1 + Z_A} = \frac{\frac{1 + \left(c_I \cdot K_{L,A}\right)^n}{1 + \left(c_I \cdot K_{L,A}\right)^n) + e^{\beta \Delta \varepsilon AI} \cdot \left(1 + (c_I \cdot K_{L,I}\right)^n)} \cdot c_{TF,all} \cdot K_A}{1 + \frac{1 + \left(c_I \cdot K_{L,A}\right)^n}{(1 + \left(c_I \cdot K_{L,A}\right)^n) + e^{\beta \Delta \varepsilon AI} \cdot \left(1 + (c_I \cdot K_{L,I}\right)^n)} \cdot c_{TF,all} \cdot K_A} \tag{S17}$$

While for repression, combining S7 and S16 equations we would have:

$$p_{bound} = \frac{1}{1+Z_R} = \left(1 + \frac{1+\left(c_I \cdot K_{L,A}\right)^n}{\left(1+\left(c_I \cdot K_{L,A}\right)^n\right) + e^{\beta\Delta\varepsilon AI} \cdot \left(1+\left(c_I \cdot K_{L,I}\right)^n\right)} \cdot c_{TF,all} \cdot K_R\right)^{-1} \qquad (S18)$$

Let $S_1 = \left(1+\left(c_I \cdot K_{L,A}\right)^n\right)$, $S_2 = 1+\left(c_I \cdot K_{L,A}\right)^n$, $SS = \left(1+\left(c_I \cdot K_{L,A}\right)^n\right) + e^{\beta\Delta\varepsilon AI} \cdot \left(1+\left(c_I \cdot K_{L,A}\right)^n\right)$,

For activator, fold change= $\frac{p_{bound}(c_I>0)}{p_{bound}(c_I=0)}$ can be written as:

$$foldchange = \frac{\dfrac{K_A \cdot c_{TF,all} \cdot \frac{S_1}{SS}}{1+K_A \cdot c_{TF,all} \cdot \frac{S_1}{SS}}}{\dfrac{K_A \cdot c_{TF,all} \cdot (1+e^{\beta\Delta\varepsilon AI})}{1+K_A \cdot c_{TF,all} \cdot (1+e^{\beta\Delta\varepsilon AI})}} = \frac{S_1 \cdot \left(1+K_A \cdot c_{TF,all} \cdot \left(1+e^{\beta\Delta\varepsilon AI}\right)\right)}{\left(1+e^{\beta\Delta\varepsilon AI}\right) \cdot \left(SS + K_A \cdot c_{TF,all} \cdot S_1\right)} \qquad (S19)$$

In this Equation, the fold-change of a repressor is determined by several key biophysical constraints. First, $K_{L,A}$ and $K_{L,I}$ represent Boltzmann-weighted association strength of the ligand for the active or inactive conformation, respectively, a larger value of $K_{L,A}$ *or* $K_{L,I}$ increases the statistical weight of ligand-bound active or inactive states and favors activation or inactivation. The number of inducer binding sites $n$ is an integer, reflecting the multivalency of the system. The free energy difference $\Delta\varepsilon_{AI}$ between inactive and active states determines the relative probability of each state, with the Boltzmann factor $e^{\beta\Delta\varepsilon_{AI}}$ often denoted as $L$ or $K^*_{RR}$. Ligand concentration $c_I$ is non-negative, and the fold-change equation integrates these parameters to compute the expected gene expression output. The model captures both induction and repression regimes, and the response also depends on experimental variables, such as $c_{TF,all}$, $K_A$, inducer concentration ($c_I$). These constraints ensure the physical plausibility of the allosteric model when applied to transcriptional regulation.

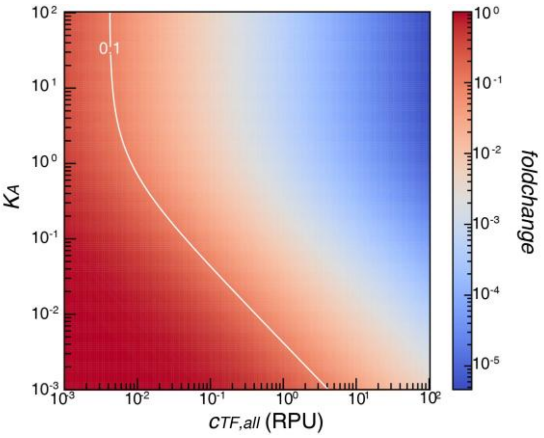

Supplementary Figure 16. Optimization heatmap of a Type I allosteric transcriptional activator plotted as a function of total transcription factor concentration ($c_{\mathrm{TF,all}}$) and the promoter association constant ($K_A$).

Allosteric transcription factors can support both activation and repression, but their signal-sensing and DNA-binding functions are structurally coupled. As a result, changing DNA specificity, retuning ligand response, or inverting regulatory logic often requires system-specific engineering or directed evolution rather than simple modular domain exchange. Their achievable regulatory behavior is therefore constrained by the fixed coupling between sensing, DNA recognition, and transcriptional output.

**Supplementary Note 2.2: Type II TF based transcriptional regulation.**

Type II systems incorporate two tunable parameters: the concentration of a functional component and its binding affinity. Adjusting either parameter changes not only the amplitude of the output but also the response sensitivity, as these two aspects are intrinsically coupled. Consequently, increasing component concentration tends to simultaneously raise maximal output and shift the activation threshold, constraining the ability to optimize different response properties independently.

$$TF_A + TF_B \overset{K_{d0}}{\rightleftharpoons} TF_A{:}TF_B$$

$$TF_A + I \overset{K_b}{\rightleftharpoons} TF_A{:}I$$

$$TF_B + I \overset{K_b}{\rightleftharpoons} TF_B{:}I$$

$$TF_A{:}I + TF_B{:}I \overset{K_{d1}}{\rightleftharpoons} (TF_A{:}I){:}(TF_B{:}I)$$

$$c_{TF_A} + c_{TF_A:I} + c_{TF_A:TF_B} + c_{(TF_A:I):(TF_B:I)} = c_{TF_A,all} \quad (S20)$$

$$c_{TF_B} + c_{TF_B:I} + c_{TF_A:TF_B} + c_{(TF_A:I):(TF_B:I)} = c_{TF_B,all} \quad (S21)$$

$$c_{TF_A} + K_b c_{TF_A} c_I + K_{d0} c_{TF_A} c_{TF_B} + K_b^2 K_{d1} c_{TF_A} c_{TF_B} c_I^2 = c_{TF_A,all} \quad (S22)$$

$$c_{TF_B} + K_b c_{TF_B} c_I + K_{d0} c_{TF_A} c_{TF_B} + K_b^2 K_{d1} c_{TF_A} c_{TF_B} c_I^2 = c_{TF_B,all} \quad (S23)$$

No inducer exists:

$$c_{TF_A} + c_{TF_A:TF_B} = c_{TF_A,all} \quad (S24)$$

$$c_{TF_B} + c_{TF_A:TF_B} = c_{TF_B,all} \quad (S25)$$

$$c_{TF_A} + K_{d0} c_{TF_A:TF_B} = c_{TF_A,all} \quad (S26)$$

$$c_{TF_B} + K_{d0} c_{TF_A:TF_B} = c_{TF_B,all} \quad (S27)$$

Full induction:

$$c_{TF_A} + c_{(TF_A:I):(TF_B:I)} = c_{TF_A,all} \quad (S28)$$

$$c_{TF_B} + c_{(TF_A:I):(TF_B:I)} = c_{TF_B,all} \quad (S29)$$

$$c_{TF_A} + K_b^2 K_{d1} c_{TF_A} c_{TF_B} c_I^2 = c_{TF_A,all} \quad (S30)$$

$$c_{TF_B} + K_b^2 K_{d1} c_{TF_A} c_{TF_B} c_I^2 = c_{TF_B,all} \quad (S31)$$

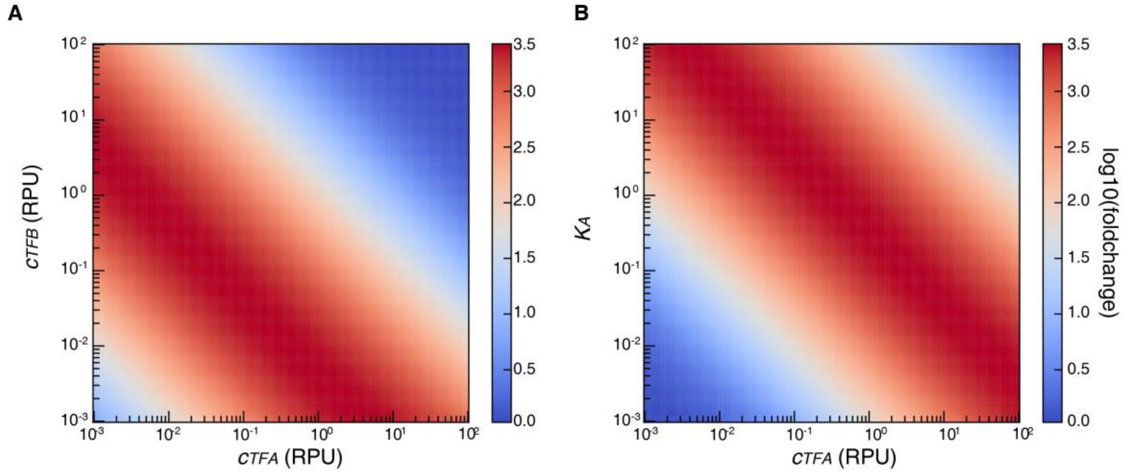

Supplementary Figure 17. Optimization landscapes of a Type II regulatory architecture. **(A)** Log10(fold change) heatmap obtained by varying the concentrations of two regulatory components (A and B). **(B)** Log10(fold change) heatmap obtained by varying the concentration of component A and its binding affinity.

#### Supplementary Note 3: Quantitative model of Type III CIC-TF

##### Supplementary Note 3.1: Equilibrium and mass-conservation model for CIC-TF oligomerization

Signal-induced cooperative transcription factors (TFs) are theoretically capable of forming oligomeric assemblies of arbitrary order ($n \in \mathbb{N}^{+}$). The oligomerization number $n$ dictates the effective cooperativity and amplification potential of the regulatory response. For example, a monomeric TF binds DNA directly, whereas a dimeric TF requires prior dimerization to achieve a DNA-binding–competent state and initiate transcriptional regulation.

In transcriptional regulation, monomeric transcription factors (TFs) that are capable of dimerization can form functional dimers through two mechanistically distinct pathways. (1) Spontaneous dimerization: thermodynamically driven association of monomers in the absence of external signals. (2) Inducer-mediated dimerization: ligand-dependent stabilization of the dimeric state, such as small-molecule binding or hormone-induced conformational changes that promote dimer assembly. Functionally active dimerized TFs subsequently bind to specific promoter sequences to regulate gene expression. This mechanism is mathematically represented as:

$$n\,TF \overset{K_{d0}}{\rightleftharpoons} TF_n$$

$$TF + I \overset{K_b}{\rightleftharpoons} TF{:}I$$

$$n\,TF{:}I \overset{K_{d1}}{\rightleftharpoons} (TF{:}I)n$$

Where $TF$ represents free TF monomer, $TF{:}I$ represents ligand-bound TF monomer, $TF_n$ and $(TF{:}I)n$ represent the active TF oligomer and ligand-bound TF oligomer of size n, respectively, capable of regulating gene expression. $I$ denotes inducer, $n \in \mathbb{N}^{+}$ represents the oligomerization number. The parameter $K_{d0}$ corresponds to the equilibrium constant for spontaneous dimerization of the transcription factor, the theoretical equilibrium constant would have units of 1/RPU, $K_{d0}$ is defined as dimensionless by normalization to a reference concentration. $K_b$ represents the affinity of the inducer for TF monomers, with units of 1/µM, and $K_{d1}$ describes the enhancement of dimerization upon inducer binding and is dimensionless, as the term $K_b^n \cdot c_I^n \cdot K_{d1}$ is dimensionless in the model.

In accordance with the law of mass conservation, the total concentration of the transcription factor across its four distinct states remains constant and is determined by the initial total monomeric TF protein concentration $c_{TF,all}$. This relationship can be mathematically expressed as:

$$c_{TF_n} = K_{d0} \cdot c_{TF}^n \tag{S32}$$

$$c_{TF:I} = K_b \cdot c_I \cdot c_{TF} \tag{S33}$$

$$c_{(TF:I)_n} = K_{d1} \cdot K_b^n \cdot c_I^n \cdot c_{TF}^n \quad (S34)$$

where $c_{TF,all}$ represents the total transcription factor protein concentration in relative promoter units (RPU) and is therefore dimensionless, while $c_I$ denotes the inducer concentration in μM.

Mass conservation gives:

$$c_{TF} + c_{TF:I} + n \cdot c_{TF_n} + n \cdot c_{(TF:I)_n} = c_{TF,all} \quad (S35)$$

$$c_{TF} + K_b \cdot c_I \cdot c_{TF} + n \cdot K_{d0} \cdot c_{TF}^n + n \cdot K_{d1} \cdot K_b^n \cdot c_I^n \cdot c_{TF}^n = c_{TF,all} \quad (S36)$$

For the case of n = 2, the solution for above equations could be written as:

$$c_{TF} = \frac{-(K_b c_I + 1) + \sqrt{(K_b c_I + 1)^2 + 8c_{TF,all}(K_b^2 \cdot c_I^2 \cdot K_{d1} + K_{d0})}}{4(K_b^2 \cdot c_I^2 \cdot K_{d1} + K_{d0})} \quad (S37)$$

$$c_{TF_2} = K_{d0} \cdot \frac{2(K_b c_I + 1)^2 - 2(K_b c_I + 1)\sqrt{(K_b c_I + 1)^2 + 8c_{TF,all}(K_b^2 \cdot c_I^2 \cdot K_{d1} + K_{d0})} + 8c_{TF,all}\left(K_b^2 \cdot c_I^2 \cdot K_{d1} + K_{d0}\right)}{16(K_b^2 \cdot c_I^2 \cdot K_{d1} + K_{d0})^2} \quad (S38)$$

$$c_{TF:I} = K_b \cdot c_I \cdot \frac{-(K_b c_I + 1) + \sqrt{(K_b c_I + 1)^2 + 8c_{TF,all}(K_b^2 \cdot c_I^2 \cdot K_{d1} + K_{d0})}}{4(K_b^2 \cdot c_I^2 \cdot K_{d1} + K_{d0})} \quad (S39)$$

$$c_{(TF:I)_2} = K_b^2 \cdot c_I^2 \cdot K_{d1} \cdot \frac{2(K_b c_I + 1)^2 - 2(K_b c_I + 1)\sqrt{(K_b c_I + 1)^2 + 8c_{TF,all}(K_b^2 \cdot c_I^2 \cdot K_{d1} + K_{d0})} + 8c_{TF,all}\left(K_b^2 \cdot c_I^2 \cdot K_{d1} + K_{d0}\right)}{16(K_b^2 \cdot c_I^2 \cdot K_{d1} + K_{d0})^2} \quad (S40)$$

**Supplementary Note 3.2: Tuning knob for active TF ratio through signal transmission efficiency**

To investigate how the fraction of active transcription factors (TFs) depends on total TF abundance, we examined monomeric and dimeric regulators and plotted the proportion of active TFs before and after induction. The resulting figure reveals a clear distinction between the two classes. For monomeric TFs, the active fraction remains constant upon induction and is unaffected by changes in total TF concentration. In contrast, for dimeric TFs, the active fraction decreases as TF abundance increases. These results show that the signal-transmission efficiency of multimeric TFs can be tuned by adjusting TF concentration, whereas monomeric TFs lack this additional regulatory degree of freedom.

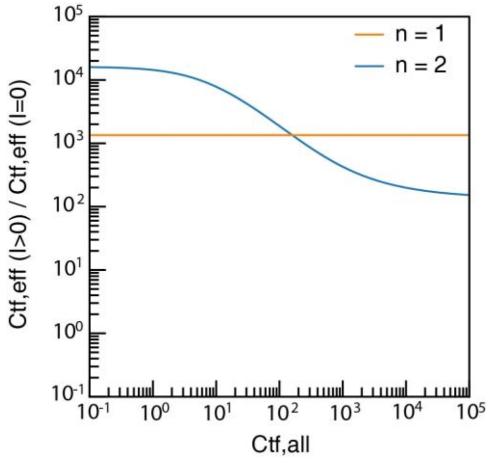

Supplementary Figure 18. Dependence of the active transcription factor (TF) fraction on total TF concentration for monomeric and dimeric regulators. The blue and yellow lines show the active TF ratio before and after induction in the case of n = 1 and n = 2, respectively.

**Supplementary Note 3.3: Modulation of Fold change in Type III CIC-TFs Through the Promoter-Binding Boltzmann Weight $K_A$**

Within our modeling framework, we assume that the binding energies of $TF_n$ and $(TF{:}I)_n$ to DNA are identical, both denoted as $E_A$. Furthermore, $p_{bound}$ is defined as the partition function of promoter output, expressed in terms of the corresponding partition function components under both inducer-free and inducer-present conditions ($Z_{TF_2}$ and $Z_{(TF:I)_2}$).

$$p_0 = p_{bound}(c_I = 0) = \frac{Z_{TF_n}}{1 + Z_{TF_n}} \tag{S41}$$

$$p_1 = p_{bound}(c_I > 0) = \frac{Z_{TF_n} + Z_{(TF:I)_n}}{1 + Z_{TF_n} + Z_{(TF:I)_n}} \tag{S42}$$

Where $p_0$ and $p_1$ correspond to experimental samples that satisfy $c_I = 0$ and $c_I > 0$, respectively, and $c_{TF,eff}$ represents the active form of TF that could bind to DNA. Building on the definitions of $c_{TF_n}$ and $c_{(TF:I)_n}$ provided above, $Z_{TF_n}$ and $Z_{(TF:I)_n}$ can be expressed as functions of $c_{TF_n}$ and $c_{(TF:I)_n}$ in conjunction with $E_A$.

$$e^{E_A} = K_A \tag{S43}$$

$$Z_{TF_n} = c_{TF_n} \cdot e^{E_A} = K_{d0} \cdot c_{TF}^{n} \cdot e^{E_A} \tag{S44}$$

$$Z_{(TF:I)_n} = c_{(TF:I)_n} \cdot e^{E_A} = K_{d1} \cdot K_b^{n} \cdot c_I^{n} \cdot c_{TF}^{n} \cdot e^{E_A} \tag{S45}$$

$$c_{TF,eff} = c_{TF_n} + c_{(TF:I)_n} \tag{S46}$$

$$p_{bound} = \frac{c_{TF,eff} \cdot e^{E_A}}{1 + c_{TF,eff} \cdot e^{E_A}} = \frac{(c_{TF_n} + c_{(TF:I)_n}) \cdot K_A}{1 + (c_{TF_n} + c_{(TF:I)_n}) \cdot K_A} \qquad (S47)$$

When the transcription factor (TF) expression level is held constant, tuning the TF–promoter binding affinity $K_A$(for example, by replacing the DNA-binding domain) provides an effective means to modulate the fold change of $p_{\text{bound}}$ between induced and uninduced states. As shown in Supplementary Fig. 19, the fold-change exhibits a non-monotonic dependence on $K_A$: it first increases and then decreases as the binding affinity strengthens. This trend indicates that $K_A$critically shapes the regulatory efficiency of the system—weak binding fails to recruit sufficient TF dimers, whereas excessively strong binding reduces the relative difference in promoter occupancy before and after induction, thereby diminishing fold-change.

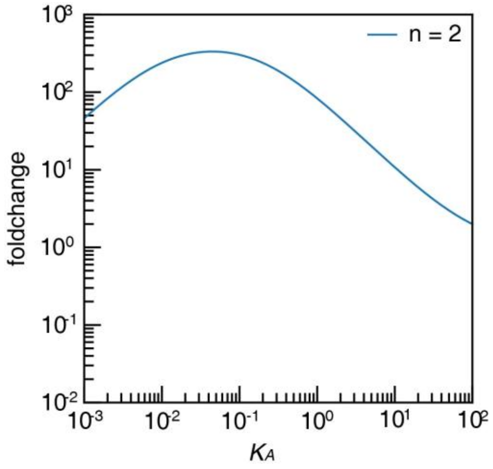

Supplementary Figure 19. Fold change of $p_{\text{bound}}$ as a function of the binding affinity $K_A$.

Therefore, the final promoter output is jointly governed by the oligomerization state of the transcription factor and its DNA-binding energy as shown in Supplementary Fig. 20. Moreover, as demonstrated in **S11** and **S47**, the achievable upper and lower bounds of promoter activity can also be modulated by altering the promoter architecture itself, providing an additional lever for fine-tuning regulatory performance.

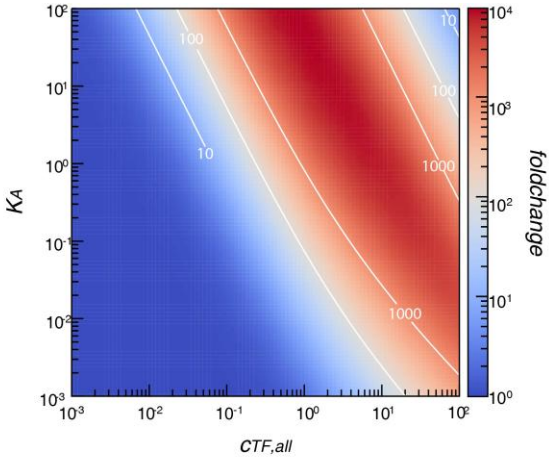

Supplementary Figure 20. Optimization landscapes of a Type III regulatory architecture. Fold-change heatmap obtained by varying the concentrations of total TF and $K_A$.

#### Supplementary Note 4: Apparent cooperative transfer model for chemically diverse CIC modules

The microscopic model described in Supplementary Note 3 provides a mechanistic representation of ligand-induced cooperative transcription factors in which an inactive monomeric TF is converted into a DNA-binding competent oligomeric state. This description is appropriate for simple inducible dimerization modules, such as quorum-sensing-derived LBDs, coiled-coil-like domains, leucine zipper-like domains, or other DD-like modules whose dominant regulatory step can be approximated as ligand-dependent dimer formation. However, the LBD collection characterized in this study also includes nuclear receptor-like modules and other chemically diverse regulatory domains. In these cases, ligand-dependent activation may involve multiple biochemical steps, including ligand binding, conformational activation, chaperone release, nuclear localization, cofactor recruitment, protein stabilization, and dimerization. These processes do not necessarily correspond to a single microscopic dimerization equilibrium.

To avoid introducing separate mechanistic models for each biochemical class of LBD, we used an apparent cooperative transfer model. This formulation preserves the same promoter-level thermodynamic output model while replacing the detailed upstream activation mechanism with an operational mapping from total TF abundance to a DNA-binding competent effective TF pool. The model therefore does not require all LBDs to share the same microscopic reaction pathway. Instead, it requires that their net input-output behavior can be represented by an apparent activation coefficient and an apparent cooperativity within the experimentally tested range.

##### Supplementary Note 4.1: Apparent mapping from total TF abundance to effective TF pool

Let

$$C = C_{TF,all}$$

denote the total abundance of a chimeric TF, quantified in relative promoter units or normalized TF abundance. We define the DNA-binding competent effective TF pool as

$$C_{TF,eff}(I,C) = \Lambda(I)C^{n_{app}} \qquad (S48)$$

where $\Lambda(I)$ is the apparent activation coefficient at inducer condition $I$, and $n_{app}$ is the apparent cooperativity. The parameter $\Lambda(I)$ absorbs all upstream biochemical steps that determine how efficiently total TF abundance is converted into an active DNA-binding pool. The parameter $n_{app}$ describes how strongly the effective pool scales with total TF abundance.

The downstream promoter occupancy is then written as

$$p_{bound}(I,C)=\frac{K_A C_{TF,eff}(I,C)}{1+K_A C_{TF,eff}(I,C)} \quad (S49)$$

where $K_A$ represents the DBD-operator association strength. The transcriptional output is

$$Y(I,C)=T_0+(T_{max}-T_0)\frac{K_A \Lambda(I)C^{n_{app}}}{1+K_A \Lambda(I)C^{n_{app}}} \quad (S50)$$

Here, $T_0$ and $T_{max}$ define the promoter output window. Thus, the final response is governed by three separable layers: effective TF-pool formation, DBD-operator coupling, and promoter output capacity.

**Supplementary Note 4.2: Relationship to the microscopic dimerization model**

For a simple ligand-induced dimerization module, the apparent model reduces to the microscopic CIC-TF model under an appropriate concentration regime. In the dimeric case,

$$n_{app}\approx 2,$$

and the effective TF pool can be approximated as

$$C_{TF,eff}^{(-)}=\Lambda_0 C^2$$

in the uninduced state and

$$C_{TF,eff}^{(+)}=\Lambda_1 C^2$$

in the induced state. In this interpretation, $\Lambda_0$ represents the apparent basal dimerization or basal active-state formation capacity, whereas $\Lambda_1$ represents the apparent induced active-state formation capacity. For ordinary inducible dimerization modules, $\Lambda_0$ is related to the basal dimerization parameter, and $\Lambda_1$ is related to ligand binding and ligand-enhanced dimer formation. Under a low-concentration approximation, these terms can be interpreted as

$$\Lambda_0\sim K_{d0}$$

and

$$\Lambda_1\sim K_b^2 K_{d1} I^2.$$

At saturating inducer concentration, the induced state can be represented more generally as

$$\Lambda_1=\Lambda(I=I_{max}).$$

Therefore, the microscopic dimerization model is a special case of the apparent cooperative transfer model, rather than a separate framework.

**Supplementary Note 4.3: Nuclear receptor-like modules**

For nuclear receptor-like modules, the same promoter-level equation is retained, but $\Lambda(I)$ should not be interpreted as a pure microscopic dimerization constant. Instead, we write

$$\Lambda(I) = K_{dim}\eta(I)^{n_{app}},$$

where $K_{dim}$ denotes the apparent contribution of dimeric or multimeric assembly and $\eta(I)$ represents the ligand-dependent active fraction generated by upstream biochemical processes. These processes may include ligand binding, chaperone release, conformational activation, nuclear localization, cofactor recruitment, and changes in protein stability.

Nuclear receptor-like modules therefore use the same promoter-output function introduced in Supplementary Note 4.1.

$$Y(I, C) = T_0 + (T_{max} - T_0)\frac{K_A\Lambda(I)C^{n_{app}}}{1 + K_A\Lambda(I)C^{n_{app}}}.$$

The distinction is interpretational rather than mathematical. For simple DD-like modules, $\Lambda(I)$ primarily reflects ligand-dependent dimer formation. For nuclear receptor-like modules, $\Lambda(I)$ represents the combined apparent efficiency of all upstream biochemical steps that generate a DNA-binding competent TF pool. This apparent formulation allows chemically diverse LBDs to be compared within a unified parameterization framework without claiming that all modules share the same microscopic mechanism.

**Supplementary Note 4.4: Apparent amplification of upstream expression differences**

The same formulation can also describe amplifier-like modules in which the input is not a small-molecule ligand but the expression level generated by an upstream regulator. If the upstream output $u$ determines the expression level of the amplifier TF,

$$C = \alpha u,$$

then the downstream output becomes

$$Y(u) = T_0 + (T_{max} - T_0)\frac{K_A\Lambda(\alpha u)^{n_{app}}}{1 + K_A\Lambda(\alpha u)^{n_{app}}}.$$

If the module behaves as an apparent dimeric amplifier with $n_{app} = 2$, then a ninefold change in upstream input, for example from $u = 0.1$ to $u = 0.9$, is transformed into an approximately

$$9^2 = 81$$

fold change at the effective TF-pool layer before promoter saturation. If non-cooperative steps, transport limitations, or active-fraction bottlenecks reduce the scaling, the fitted value may satisfy $n_{app} < 2$. Conversely, if additional thresholding or higher-order effects contribute to active-state formation, the fitted value may exceed 2. Therefore, $n_{app}$ provides an operational measure of whether a module functions as a linear scaler, a dimer-like amplifier, or a higher-order thresholding element.

**Supplementary Note 4.5: Apparent parameters reported for module comparison**

For each module, we define the uninduced and induced apparent activation coefficients as

$$\Lambda_0 = \Lambda(I = 0)$$

and

$$\Lambda_1 = \Lambda(I = I_{max}).$$

The apparent activation ratio is

$$R_\Lambda = \frac{\Lambda_1}{\Lambda_0},$$

or, equivalently,

$$Q_\Lambda = \log_{10}\left(\frac{\Lambda_1}{\Lambda_0}\right).$$

$Q_\Lambda$ provides an operational ranking score for LBD or DD modules. However, it is not a sufficient statistic for final sensor performance because the observed output also depends on DBD-operator coupling, TF abundance, and promoter output range:

$$K_A, \quad C_{TF,all}, \quad T_0, \quad T_{max}.$$

We therefore use $\Lambda_0$, $\Lambda_1$, $R_\Lambda$, and $n_{app}$ as module-level parameters, while final sensor optimization is performed using the complete output function.

**Supplementary Note 4.6: Fitting strategy and model comparison**

To test whether the apparent cooperative transfer model is sufficient, we fit expression-gradient data across multiple TF abundance levels. The recommended first-pass input levels are

$$C = 0.03,\ 0.1,\ 0.3,\ 0.9,\ 1.5,\ 3,$$

with an optional higher input such as

$$C = 10$$

to identify saturation or burden-dependent deviations. The points $C = 0.1$ and $C = 0.9$ are particularly important because they represent the intended low and high upstream input states for amplifier-like applications.

For each module, two inducer conditions are first considered:

$$I = 0$$

and

$$I = I_{max}.$$

For nuclear receptor-like modules, the first round should also use no ligand and saturating ligand. Ligand-gradient experiments can then be performed in a second round to verify whether the selected $I_{max}$ condition is close to saturation.

We compare two primary models. The first is a fixed apparent dimer model:

$$Y_s(C) = T_0 + (T_{max} - T_0)\frac{K_A \Lambda_s C^2}{1 + K_A \Lambda_s C^2},$$

where

$$s \in \{0,1\}$$

denotes the uninduced and induced states. The second is a free-cooperativity model:

$$Y_s(C) = T_0 + (T_{max} - T_0)\frac{K_A \Lambda_s C^{n_{app}}}{1 + K_A \Lambda_s C^{n_{app}}}.$$

The fitted parameters are

$$\Lambda_0, \quad \Lambda_1, \quad n_{app}.$$

When $K_A$, $T_0$, and $T_{max}$ are independently calibrated, they are fixed during fitting to reduce parameter non-identifiability. In the first fitting round, the uninduced and induced curves share a common $n_{app}$. If systematic residuals indicate that the two states have different scaling behavior, separate $n_0$ and $n_1$ values may be considered as a diagnostic extension, but this additional freedom is not used as the default model.

Fitting is performed on the raw output scale using nonlinear least squares, weighted nonlinear least squares, or a likelihood-based model when an empirical noise model is available. A general objective function is

$$\min_{\theta} \sum_i w_i \left[Y_i^{obs} - Y_i^{model}(\theta)\right]^2,$$

where

$$\theta = (\Lambda_0, \Lambda_1, n_{app}).$$

If replicate measurements are available, $w_i$ can be set to the inverse variance. Otherwise, replicate variance or an empirical mean-variance relationship can be used to estimate measurement uncertainty.

**Supplementary Note 4.7: Diagnostic use of logit-linearization**

For visualization, the measured output can be transformed into promoter occupancy:

$$p = \frac{Y - T_0}{T_{max} - T_0}.$$

The corresponding logit transformation is

$$\mathrm{logit}(p) = \log\left(\frac{p}{1-p}\right).$$

Because

$$p = \frac{K_A \Lambda C^n}{1 + K_A \Lambda C^n},$$

we obtain

$$\mathrm{logit}(p) = \log(K_A) + \log(\Lambda) + n\log(C).$$

This relationship is useful for diagnostic visualization and approximate estimation of the apparent slope. For example, it can reveal whether a module behaves closer to a linear scaler, a dimer-like amplifier, or a higher-order thresholding element.

However, logit-linearization is not used for final parameter estimation or model selection. The transformation changes the error structure, strongly amplifies noise near $p = 0$ and $p = 1$, and propagates uncertainty in $T_0$ and $T_{max}$ nonlinearly into $p$. Moreover, fluorescence and RPU measurements are typically heteroscedastic. Therefore, all reported apparent parameters are estimated using nonlinear fitting in the raw output space or using an explicit measurement-noise model.

**Supplementary Note 4.8: Criteria for evaluating model adequacy**

The apparent cooperative transfer model is considered adequate when the following criteria are satisfied. First, the model should explain the expression-gradient data with high accuracy, for example with $R^2 > 0.85$ or a sufficiently low normalized root-mean-square error. Second, predictions at the target input states should be accurate. In practice, the predicted outputs at

$$C = 0.1$$

and

$$C = 0.9$$

should deviate from experimental measurements by less than approximately 20-30%, or the predicted fold-change should be within approximately 1.5-fold of the measured value, depending on the experimental noise level.

Third, the residuals should not show systematic structure. Consistent underprediction at low input or overprediction at high input may indicate missing saturation, transport limitation, active-fraction gating, or burden-dependent effects. Fourth, the fitted $n_{app}$ should be stable across bootstrap resampling or replicate fitting. A broad or unstable confidence interval indicates that the dataset is insufficient to identify the scaling exponent.

Finally, for a modular LBD or DD, the fitted parameters

$$\Lambda_0, \quad \Lambda_1, \quad n_{app}$$

should be approximately conserved across different DBDs after accounting for DBD-specific $K_A$. Likewise, when the same module is connected to different downstream promoters, $\Lambda$ and $n_{app}$ should remain approximately unchanged, whereas $T_0$ and $T_{max}$ may vary with promoter architecture. Failure of these invariance tests would indicate coupling between the LBD/DD module and the DBD or promoter context, in which case the apparent cooperative transfer model should be treated as an approximation rather than a fully modular description.

**Supplementary Note 4.9: Diagnostic cascade model for nuclear receptor-like modules**

If the apparent cooperative transfer model exhibits systematic failure for a nuclear receptor-like module, a more explicit diagnostic cascade model can be used. This model is not used as the default fitting framework but can identify whether upstream activation steps limit the response.

For example, the ligand-dependent active fraction can be represented as

$$\eta(C) = \eta_0 + \eta_{max} \frac{C}{K_\eta + C}.$$

The effective TF pool is then

$$C_{TF,eff} = K_{dim}[\eta(C)C]^2.$$

The output becomes

$$Y(C) = T_0 + (T_{max} - T_0)\frac{K_A K_{dim}[\eta(C)C]^2}{1 + K_A K_{dim}[\eta(C)C]^2}.$$

This diagnostic model can test whether chaperone release, nuclear localization, active-fraction formation, or high-input saturation compresses the response. However, because it introduces additional parameters and may reduce identifiability, it is used only when the simpler apparent model shows clear structured residuals.

##### Supplementary Note 4.10: Summary

In summary, all LBD, DD, nuclear receptor-like, and amplifier-like modules are represented by the common effective-pool relationship

$$C_{TF,eff} = \Lambda(I)C_{TF,all}^{n_{app}}.$$

Simple inducible dimerization modules correspond to the special case in which $n_{app} \approx 2$ and $\Lambda(I)$ primarily reflects ligand-dependent dimer formation. Nuclear receptor-like modules are interpreted more generally: ligand binding, chaperone release, conformational activation, localization, cofactor recruitment, and related biochemical steps are absorbed into $\Lambda(I)$ and $n_{app}$. This formulation preserves a unified promoter-level output model while allowing chemically diverse LBDs to be compared, ranked, and optimized within the same quantitative framework.

#### Supplementary Note 5: Assessing Transcriptional Regulation by Type III CIC Transcription Factors.

##### Supplementary Note 5.1: Extraction of effective activation free energy *F*

To systematically identify the molecular factors that influence transcriptional output in the Type III CIC regulatory system, we reformulate the promoter occupancy in a thermodynamic form. The transcription factor (TF) binding probability is written as a logistic function of an effective activation free energy $F$:

$$p_{bound} = \left(1 + c_0 \cdot e^{-\beta F}\right)^{-1} = \frac{c_0 \cdot e^{\beta F}}{1 + c_0 \cdot e^{\beta F}} \qquad (S48)$$

$$p_{bound} = \frac{Z_{TF_n} + Z_{(TF:I)_n}}{1 + Z_{TF_n} + Z_{(TF:I)_n}} = \frac{c_{TF,eff} \cdot e^{E_A}}{1 + c_{TF,eff} \cdot e^{E_A}} \qquad (S49)$$

Where the $c_0 = 1$ RPU serves as the standard-state concentration used to render the exponential factor dimensionless, and $\beta = 1/(k_B T)$is the inverse thermal energy.

With the above two equations,

$$c_0 \cdot e^{\beta F} = c_{TF,eff} \cdot e^{E_A} \qquad (S50)$$

$$F = \frac{1}{\beta} \cdot \ln\left(\frac{c_{TF,eff}}{c_0} \cdot e^{E_A}\right) = k_B T \left(\ln\left(\frac{c_{TF,eff}}{c_0}\right) + E_A\right) \tag{S51}$$

$$c_{TF,eff} = K_{d0} \cdot c_{TF}^2 + K_{d1} \cdot K_b^2 \cdot c_I^2 \cdot c_{TF}^2 = \left(K_b^2 K_{d1} c_I^2 + K_{d0}\right) \cdot c_{TF}^2 \tag{S52}$$

Let $\sqrt{(K_b c_I + 1)^2 + 8c_{TF,all}(K_b^2 K_{d1} c_I^2 + K_{d0})} = S$,

$$\begin{aligned}
c_{TF,eff} &= \left(K_b^2 K_{d1} c_I^2 + K_{d0}\right) \cdot \left(\frac{S - (K_b c_I + 1)}{4(K_b^2 K_{d1} c_I^2 + K_{d0})}\right)^2 \\
&= \left(K_b^2 K_{d1} c_I^2 + K_{d0}\right) \cdot \left(\frac{\big(S - (K_b c_I + 1)\big)\big(S + (K_b c_I + 1)\big)}{4(K_b^2 K_{d1} c_I^2 + K_{d0})\big(S + (K_b c_I + 1)\big)}\right)^2 \\
&= \left(K_b^2 K_{d1} c_I^2 + K_{d0}\right) \cdot \left(\frac{(S^2 - (K_b c_I + 1)^2)}{4(K_b^2 K_{d1} c_I^2 + K_{d0})\big(S + (K_b c_I + 1)\big)}\right)^2 \\
&= \left(K_b^2 K_{d1} c_I^2 + K_{d0}\right) \cdot \left(\frac{8c_{TF,all}(K_b^2 K_{d1} c_I^2 + K_{d0})}{4(K_b^2 K_{d1} c_I^2 + K_{d0})\big(S + (K_b c_I + 1)\big)}\right)^2 \\
&= \left(K_b^2 K_{d1} c_I^2 + K_{d0}\right) \cdot \left(\frac{\left(2c_{TF,all}\right)^2}{\left(S + (K_b c_I + 1)\right)^2}\right)
\end{aligned} \tag{S53}$$

Thus, for Type III systems equipped with a specific ligand-binding domain (LBD), transcriptional output is jointly determined by the total TF concentration and the assembly energy $E_A$. These two parameters independently modulate the maximal activity and the signal-transmission efficiency, respectively.

**Supplementary Note 5.2: Total TF Levels ($c_{TF,all}$) Shape Cooperative Transcriptional Control**

For compact notation, let

$$C \equiv C_{\mathrm{TF,all}},$$
$$A \equiv 1 + K_b C_I,$$
$$D \equiv K_{d0} + K_b^2 K_{d1} C_I^2,$$

where $M \equiv C_{\mathrm{TF}}$ denotes the concentration of free TF monomers. The dimeric mass-conservation equation is

$$C = AM + 2DM^2,$$

and the DNA-binding-competent effective TF pool is

$$C_{\mathrm{TF,eff}} = DM^2.$$

1) For low $c_{TF,all}$,

$$T(x) = \frac{c_{TF,eff} \cdot K_A}{1 + c_{TF,eff} \cdot K_A} \cdot (T_{max} - T_0) + T_0 \tag{S55}$$

$$S(c_{TF} \to 0) = \sqrt{(K_b c_I + 1)^2 + 8c_{TF,all}(K_b^2 K_{d1} c_I^2 + K_{d0})} \approx (K_b c_I + 1), \qquad 8c_{TF,all}D^2 \ll A^2 \qquad (S56)$$

$$c_{TF,eff} = \left(\frac{\left(2c_{TF,all}\right)^2}{\left(S + (K_b c_I + 1)\right)^2}\right) \cdot \left(K_b^2 K_{d1} c_I^2 + K_{d0}\right) \approx \left(\frac{c_{TF,all}}{A}\right)^2 \cdot D = \frac{c_{TF,all}^2}{A^2} \cdot D \qquad (S57)$$

$$T(x) = \frac{c_{TF,all}^2}{\frac{A^2}{D \cdot K_A} + c_{TF,all}^2} \cdot (T_{max} - T_0) + T_0 \qquad (S58)$$

Which is a Hill-like equation for $T(x)$

$$T(x) = T_0 + \frac{(T_{max} - T_0) \cdot c_{TF,all}{}^2}{K^2 + c_{TF,all}{}^2} \qquad (S59)$$

Where $K^2 = \frac{A^2}{D \cdot K_A}$, $T(x) \propto c_{TF,all}{}^2$, indicating cooperative binding with a Hill coefficient n=2.

1) When $c_{TF,all}$ is high,

$$S(c_{TF} \to \infty) = \sqrt{8c_{TF,all}(K_b^2 K_{d1} c_I^2 + K_{d0})} \qquad (S60)$$

$$c_{TF,eff} = \left(K_b^2 K_{d1} c_I^2 + K_{d0}\right) \cdot \left(\frac{\left(2c_{TF,all}\right)^2}{\left(S + (K_b c_I + 1)\right)^2}\right)$$
$$\approx \left(K_b^2 K_{d1} c_I^2 + K_{d0}\right) \cdot \left(\frac{4c_{TF,all}{}^2}{8c_{TF,all}(K_b^2 K_{d1} c_I^2 + K_{d0})}\right) = \frac{c_{TF,all}}{2} \qquad (S61)$$

$$c_{TF,eff} \propto c \qquad (S62)$$

In this regime the effective TF concentration grows approximately linearly with total TF: $c_{\mathrm{TF,eff}}(c) \propto c$.

$$T(x) = \frac{c_{TF,eff} \cdot K_A}{1 + c_{TF,eff} \cdot K_A} \cdot (T_{max} - T_0) + T_0 = \frac{c_{TF,all}}{\frac{2}{K_A} + c_{TF,all}} \cdot (T_{max} - T_0) + T_0 \qquad (S63)$$

Before full saturation, this implies that $T(x)$tends towards a first-order dependence on $c$, and the effective Hill slope $n_{\mathrm{eff}}(c)$ approaches 1. For sufficiently large $c$, $c_{\mathrm{TF,eff}} \cdot K_A \gg 1$ and the output saturates at

$$T(x) \to T_{max} \qquad (S64)$$

The system transitions to non-cooperative binding with a Hill coefficient n=1

2) When $c_{TF,all}$ is intermediate, Because the exact expression transitions smoothly from a quadratic to a linear function, the effective scaling exponent necessarily satisfies

$$c_{\mathrm{TF,eff}} \sim {c_{TF,all}}^{\alpha}, 1 < \alpha < 2 \tag{S65}$$

This exponent directly determines the apparent cooperativity perceived by the promoter because the transcriptional output

$$T(x) = T_0 + (T_{max} - T_0)\frac{c_{\mathrm{TF,eff}}K_A}{1 + c_{\mathrm{TF,eff}}K_A}$$

is approximately linear in $c_{\mathrm{TF,eff}}$ in the intermediate region (where the promoter activation term is steepest). Therefore: $1 < hill\ slope < 2$ in the intermediate regime.

**Supplementary Note 5.3: Promoter occupancy with multiple operator sites.**

According to probability theory, if the probability of a single site being bound is $p_{bound}$, the probability of it remaining unbound is $1 - p_{bound}$. For a system with $n$ independent binding sites, the probability that at least one site is bound is calculated as:

$$p'_{bound} = 1 - (1 - p_{bound})^n \tag{S66}$$

In biological systems, the binding probability $p_{bound}$ is correlated with transcription factor expression levels, inducer concentration, and the DNA-binding affinity of the TF. $p_{bound}$ is formally defined as $p'_{bound}$, the final output is defined as:

$$T(x) = p'_{bound} \cdot (T_{max} - T_0) + T_0 = (1 - [1 - \left(1 + e^{-\beta F}\right)^{-1}]^n) \cdot (T_{max} - T_0) + T_0 \tag{S67}$$

#### Supplementary Note 6: Extension of the Regulatory Model to Repressor and Hybrid Promoters.

##### Supplementary Note 6.1: Functional inversion of CIC-TFs

Our engineered inducible dimerization transcription factors are capable of functional inversion, meaning that they can be designed either as activators or as repressors. Activator- and repressor-type transcription factors which share the same LBD share the same active TF partition. and for which share DBDs, their binding energies to the promoter are assumed to be identical ($E_A = E_R, K_A = K_R$). Under this assumption, both systems differ only in how their promoter occupancy contributes to the transcriptional output.

$$p_{bound}(c_I = 0, R) = \frac{1}{1 + Z_{TF_n}} \quad (S68)$$

$$p_{bound}(c_I > 0, R) = \frac{1}{1 + Z_{TF_n} + Z_{(TF:I)_n}} \quad (S69)$$

We found that there is a clear link between $p_{bound}(A)\ and\ p_{bound}(R)$ by comparing S68-69 and S41-42:

$$p_{bound}(R) = 1 - p_{bound}(A) \quad (S70)$$

And also $p_{bound}$ could be written as:

$$p_{bound}(R) = \left(1 + c_0 \cdot e^{\beta F_R}\right)^{-1} \quad (S71)$$

For an activator, TF binding increases the probability of the ON state, and fold-change is naturally defined as $p_{bound}(c_I > 0)/p_{bound}(c_I = 0)$. For a repressor, TF binding reduces the ON-state probability, so the biologically meaningful fold-change is defined in the opposite direction, $p_{bound}(c_I = 0)/p_{bound}(c_I > 0)$, such that values greater than one consistently represents stronger regulatory effects.

Integrating the analyses in S68–70 and S41–42, we find that the $p_{\text{bound}}$ ratios of activation and repression, computed under the same parameter set ($K_{d0}, K_b, K_{d1}, c_{TF,all}$), satisfy the following equation:

$$\frac{p_{\text{bound}}(c_I > 0, A)}{p_{\text{bound}}(c_I = 0, A)} \cdot \frac{p_{\text{bound}}(c_I = 0, R)}{p_{\text{bound}}(c_I > 0, R)} = \frac{c_{TF,eff}(c_I > 0)}{c_{TF,eff}(c_I = 0)} = constant \quad (S72)$$

Consequently, activator and repressor fold-change expressions are strict reciprocals, leading to a mechanistic symmetry in how promoter occupancy is modulated despite producing opposite transcriptional outcomes. This reciprocal relationship applies to the idealized occupancy term under matched parameters. The realized transcriptional output can differ between activation and repression because promoter geometry, basal activity, steric occlusion, and output-window limits enter downstream of occupancy.

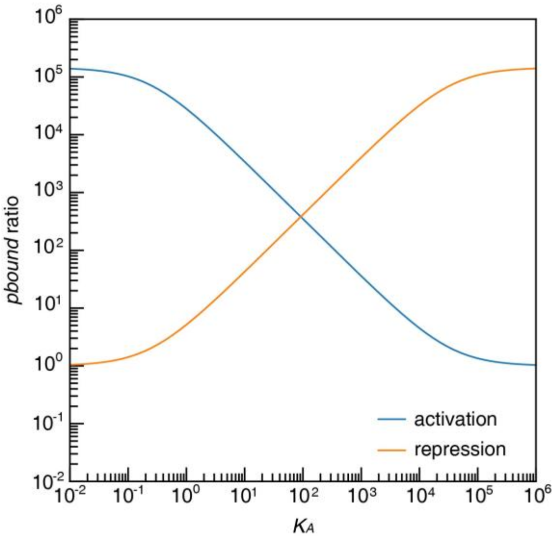

Supplementary Figure 21. Theoretical $p_{bound}$ ratios for a CIC-based activator and a functionally inverted repressor. For the repressor, the ratio reflects the induction-dependent decrease in the transcriptionally permissive, repressor-unbound promoter state; for the activator, it reflects the induction-dependent increase in activator occupancy.

Similarly, by plotting 2D heatmaps for the activator and repressor variants, we found that transcription factors with the same domain architecture can achieve higher dynamic range when configured as activators.

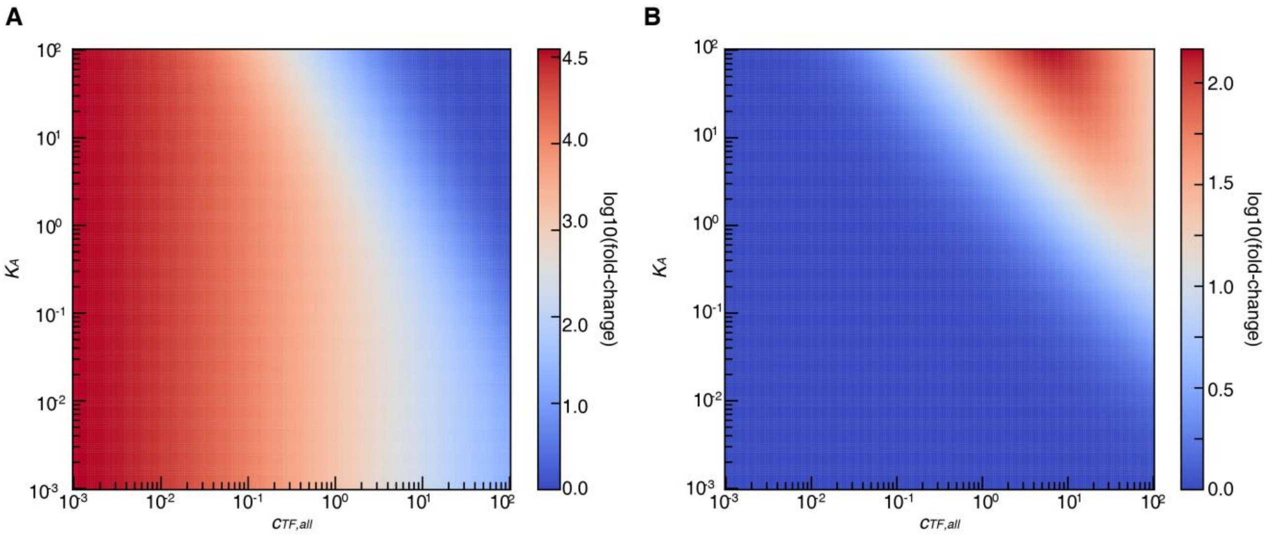

Supplementary Figure 22. Comparison of activator and repressor fold-change within the fixed feasible region.

**Supplementary Note 6.2: Expanding the Thermodynamic Model to Hybrid Promoter Logic**

When activator and repressor coexist, according to S6-S11 equations, we could write the $p_{bound}$ as:

$$p_{bound} = \frac{K_A \cdot c_{TF_A,eff}}{1 + K_A \cdot c_{TF_A,eff} + K_R \cdot c_{TF_R,eff} + K_A \cdot c_{TF_A,eff} \cdot K_R \cdot c_{TF_R,eff}} = p_{bound}(A) \cdot p_{bound}(R) \quad (S73)$$

$$p_{bound} = \left(1 + c_0 \cdot e^{-\beta F_A}\right)^{-1} \cdot \left(1 + c_0 \cdot e^{\beta F_R}\right)^{-1} \quad (S74)$$

Thus, the transcriptional output of hybrid promoter could be written as follows:

$$T(x) = p_{bound} \cdot (T_{max} - T_0) + T_0 = \left(1 + c_0 \cdot e^{-\beta F_A}\right)^{-1} \cdot \left(1 + c_0 \cdot e^{\beta F_R}\right)^{-1} \cdot (T_{max} - T_0) + T_0 \quad (S75)$$

#### Supplementary Note 7: Sensor performance metrics.

For the case of n = 2, the derivation of these expressions yields explicit mathematical expressions for the concentrations of individual transcription factor components. Under the condition of $c_I = 0$, the functional response follows the blue curve; whereas under $c_I \gg c_{TF}$, it follows the orange curve, as depicted in the figure. The figure demonstrates that variations in inducer concentration do not affect the overall shape of the output curve; rather, they result in a translational shift of the curve within its adjustable range.

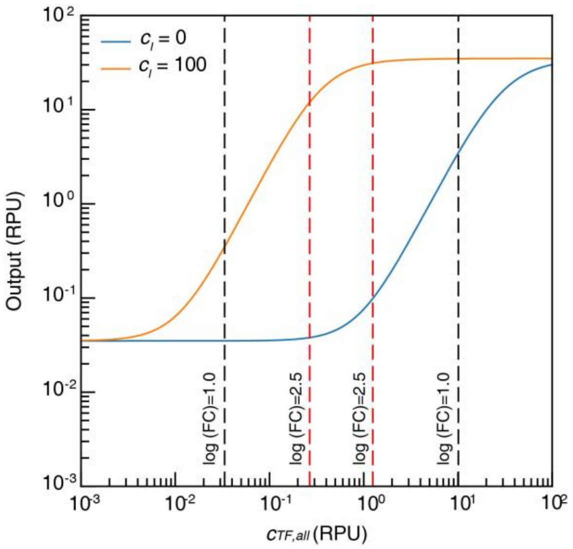

Supplementary Figure 23. The output profile under $c_I = 0\,uM$ and $c_I = 100\,uM$ varying $c_{TF,all}$.

For our transcriptional regulatory systems, the following key metrics are considered:

1) Fold change. In general, when the inducer concentration is nonzero but not saturating, the expression for the regulatory fold change is given as follows:

$$fold\ change = \frac{p_{bound}(c_I = I, n = 2)}{p_{bound}(c_I = 0, n = 2)} = \frac{\dfrac{Z_{TF_2} + Z_{(TF:I)_2}}{1 + Z_{TF_2} + Z_{(TF:I)_2}}}{\dfrac{Z_{TF_2}}{1 + Z_{TF_2}}} \quad (S76)$$

When the inducer concentration is saturating, the expression for the regulatory fold change is given as follows:

$$fold\ change = \frac{p_{bound}(c_I \to \infty, n = 2)}{p_{bound}(c_I = 0, n = 2)} = \frac{\dfrac{Z_{(TF:I)_2}}{1 + Z_{(TF:I)_2}}}{\dfrac{Z_{TF_2}}{1 + Z_{TF_2}}} \quad (S77)$$

While that for a repressor is given by:

$$fold\ change = \frac{p_0}{p_1} = \frac{p_{bound}(c_I = 0, n = 2)}{p_{bound}(c_I > 0, n = 2)} = \frac{\dfrac{1}{1 + Z_{TF_2}}}{\dfrac{1}{1 + Z_{TF_2} + Z_{(TF:I)_2}}} \quad (S78)$$

2) Strength. Similarly, when the inducer concentration is nonzero but not saturating, the expression for the transcriptional strength is given as follows:

$$\Delta\text{Exp} = [p_{bound}(c_I > 0, n = 2) - p_{bound}(c_I = 0, n = 2)] \cdot (T_{max} - T_0) \quad (S79)$$

$$\Delta\text{Exp} = \left( \frac{Z_{TF_2} + Z_{(TF:I)_2}}{1 + Z_{TF_2} + Z_{(TF:I)_2}} (c_I > 0) - \frac{Z_{TF_2}}{1 + Z_{TF_2}} (c_I = 0) \right) \cdot (T_{max} - T_0) \quad (S80)$$

When the inducer concentration is saturating, the expression for the transcriptional strength is given as follows:

$$\Delta\text{Exp} = [p_{bound}(c_I \to \infty, n = 2) - p_{bound}(c_I = 0, n = 2)] \cdot T_{max} = \left( \frac{Z_{(TF:I)_2}}{1 + Z_{(TF:I)_2}} - \frac{Z_{TF_2}}{1 + Z_{TF_2}} \right) \cdot (T_{max} - T_0) (S81)$$

3) Crosstalk. We calculated the signal-to-noise ratios of different inducers responding to the same transcription factor (TF) as a metric of crosstalk:

$$crosstalk = \frac{p_{bound}(c_{I_1} \to \infty, n = 2)}{p_{bound}(c_{I_2} \to \infty, n = 2)} = \frac{\frac{Z_{(TF:I_1)_2}}{1 + Z_{(TF:I_1)_2}}}{\frac{Z_{(TF:I_2)_2}}{1 + Z_{(TF:I_2)_2}}} \quad (S82)$$

#### Supplementary Note 8: Parameters characterization and multi-objective optimization strategy.

##### Supplementary Note 8.1: Parameters characterization in yeast.

We implemented a hierarchical parameter estimation workflow to obtain the kinetic parameters governing the relationship between total transcription factor concentration ($c_{TF,all}$), inducer concentration ($c_I$), and the measured response $T(x)$. The output is formulated as:

$$T(x) = \frac{\left(\frac{\left(2c_{TF,all}\right)^2}{\left(S + (K_b c_I + 1)\right)^2}\right) \cdot K_A}{(K_b^2 K_{d1} c_I^2 + K_{d0}) + \left(\frac{\left(2c_{TF,all}\right)^2}{\left(S + (K_b c_I + 1)\right)^2}\right) \cdot K_A} \cdot (T_{max} - T_0) + T_0 \qquad (S83)$$

where $S = \sqrt{(K_b c_I + 1)^2 + 8c_{TF,all}(K_b^2 K_{d1} c_I^2 + K_{d0})}$.

To ensure consistent parameterization across different ligand-binding domains (LBDs), parameters were estimated in three sequential phases.

**Phase 1: Estimation of globally shared kinetic parameters using the $\mathbf{RpaR_{179}}$ dataset.** The complete quantitative dataset for $RpaR_{179}$ was used to determine the baseline kinetic parameters $K_{d0}$, $K_b$, and $K_{d1}$. Parameter optimization was implemented in PyTorch, with all parameters represented in logarithmic space to enforce positivity. Optimization was performed using the Adam algorithm with a mean-squared error loss over 100,000 iterations. Gradient clipping was applied to enhance numerical stability.

**Phase 2: Determination of DBD-specific association constants**. With the shared kinetic parameters fixed to $RpaR_{179}$ derived values, the association constant $K_A$ for each DBD was independently estimated. For each DBD dataset, $K_A$ was optimized separately for 150,000 iterations, allowing domain-specific DNA-binding affinities to be captured while preserving a consistent kinetic framework.

**Phase 3: Characterization of kinetic parameters for additional ligand-binding domains**. After fixing the DBD-specific $K_A$ values, the kinetic parameters $K_{d0}$, $K_b$, and $K_{d1}$ of other LBDs were characterized based on corresponding datasets. This step allowed each LBD to acquire its own kinetic parameter set while maintaining the shared dissociation constants, enabling meaningful comparisons across LBDs.

**Data preprocessing and model evaluation**. Experimental measurements of total TF concentration, inducer concentration, and fluorescence response were compiled from multiple datasets. The baseline and maximal response parameters, $T_{0,variant}$ in Supplementary table 2 and $T_{\max} = 35.85$, were fixed based on calibration measurements. Model performance was quantified using the log-scale coefficient of determination ($R^2$) computed both globally and within individual experimental conditions, confirming the robustness of the hierarchical estimation strategy.

**Supplementary Note 8.2: Master Curve Construction and Normalization**

The master curve was constructed using a mathematical model describing the sensor response as a function of transcription factor concentration. For a given inducer concentration $c_I$ and total transcription factor concentration ($c_{TF,all}$), the system is governed by the model equations *S(83)* provided above, with $T_{\max} = 35.85$ fixed based on prior parameter optimization. Each sensor variant is characterized by a unique association constant $K_A$, thirteen variants were analyzed, with $K_A$ values spanning 0.10 to 40.98.

To generate a unified master curve, a reference sensor with $K_A = 6.22$ was selected. Experimental measurements from all sensor variants were normalized to this reference using a deformation function-based correction. For each inducer condition ($c_I$ = 0 or 100 μM), corrected responses were computed as:

$$y_{\text{corrected}} = y_{\text{experimental}} + \left[T_{master}(c_{TF,all}) - T_{variant}(c_{TF,all})\right] - T_{0_{variant}} \qquad (S84)$$

The deformation terms $\left[T_{master}(c_{TF,all}) - T_{variant}(c_{TF,all})\right]$ were obtained by interpolating the theoretical response curves at experimentally sampled total transcription factor concentration.

Alignment quality was quantified using the coefficient of determination ($R^2$), computed separately for each inducer condition across all normalized data points. The complete normalized dataset—including corrected values, variant identities, experimental conditions, and visualization color codes—has been archived for reference. All computations and visualizations were performed in Python using standard scientific computing libraries.

**Supplementary Note 8.3: Model transfer for mammalian transcriptional system.**

**Supplementary Note 8.3.1: Stable-cell endpoint parameterization of mammalian DBD and LBD modules.**

To transfer the CIC-TF model to mammalian cells, we used stable-cell OFF/ON endpoint measurements to estimate mammalian host-specific module parameters. Each tested transcription factor consisted of one DNA-binding domain (DBD), one ligand-binding domain (LBD), and VP16. Reporter output was measured at three calibrated TF-input levels under non-induced and induced conditions. The calibrated total TF abundances were treated as fixed experimental inputs:

$$C_{TF,all} = 0.01, 0.40, 3.02 \text{ RPU}.$$

The OFF state was defined as $c_I = 0$. The ON state used the inducer concentration applied in the corresponding stable-cell endpoint measurement. When the same LBD was measured at more than one ON inducer concentration, the inducer concentration was treated as condition-specific, while the intrinsic LBD-associated parameter was shared.

For a TF variant containing DBD d and LBD l, the OFF/ON endpoint response was represented by a dimeric effective-pool approximation. In the uninduced state, the DNA-binding-competent TF pool was written as:

$$c_{TF,eff}^{OFF} = K_{d0,l} \cdot c_{TF,all}^{2} \quad (S85)$$

In the induced state, the effective TF pool was written as:

$$c_{TF,eff}^{ON} = K_{d0,l} \cdot Q_l \cdot c_I^2 \cdot c_{TF,all}^{2} \quad (S86)$$

Here,

$$Q_{LBD} = \frac{K_{d1,l} \cdot K_{b,l}^2}{K_{d0,l}} \quad (S87)$$

which represents the apparent LBD induction potential, or the induced enrichment of the active TF pool relative to basal active-state formation. Therefore, the induced effective pool can also be written as:

$$c_{TF,eff}^{ON} = K_{d1,l} \cdot K_{b,l}^2 \cdot c_I^2 \cdot c_{TF,all}^{2} \quad (S88)$$

The promoter-binding weight was calculated as:

$$Z_{d,l,s} = K_{A,d} \cdot c_{TF,eff}^{s} \quad (S89)$$

where s denotes the OFF or ON state. The probability of promoter occupancy was then:

$$p_{bound} = \frac{Z_{d,l,s}}{1 + Z_{d,l,s}} \quad (S90)$$

The predicted output was calculated as:

$$\hat{T}_{d,l,s} = T_{0,d} + A_d \cdot p_{bound} \quad (S91)$$

Here, $K_{A,d}$ is the DBD-associated DNA-binding parameter, $T_{0,d}$ is the DBD-specific basal output, and $A_d$ is the DBD-specific inducible output amplitude. Thus, the saturated output for DBD d is $T_{0,d} + A_d$. In the primary endpoint-fitting model, TFs sharing the same DBD were constrained to share $K_{A,d}$, $T_{0,d}$, and $A_d$, whereas TFs sharing the same LBD were constrained to share$K_{d0,l}$ and $Q_{lbd}$.

This parameterization decomposes the mammalian response into two modular layers. The DBD layer is represented by $K_A$, which controls the conversion of the effective TF pool into promoter occupancy. The LBD layer is represented by $Q_{LBD} = \frac{K_{d1,l} \cdot K_{b,l}^2}{K_{do,l}}$, which captures ligand-dependent enrichment of the active TF pool. The LBD-specific $K_{d0}$ was retained as a nuisance parameter because the OFF state primarily constrains the product $K_A \, , K_{d0}$. The reported LBD ranking parameter is therefore $Q_{LBD}$ , rather than the individual microscopic parameters $K_b$, $K_{d1}$, and $K_{d0}$.

Because OFF/ON endpoint measurements alone do not uniquely determine the absolute scale of $K_A$ and $K_{d0}$, one DBD was used as an anchor. In the current implementation, lexA$_{ec87}$ was fixed to:

$$K_A = 9.15$$

All other DBD-associated $K_A$ values were estimated relative to this anchor. Changing the anchor rescales the absolute $K_A$ and $K_{d0}$ values but does not change the model-predicted outputs. Thus, the endpoint-derived $K_A$ values should be interpreted as mammalian apparent DBD binding-strength parameters under the chosen anchor, and $Q_{lbd}$ should be interpreted as an apparent LBD induction-potential parameter over the tested concentration range.

**Supplementary Note 8.3.2: Endpoint fitting and mammalian design-space optimization.**

The mammalian endpoint parameters were estimated by nonlinear least-squares fitting on the raw reporter-output scale. Positive parameters, including $K_A$, $K_{d0}$, $Q_{lbd}$, and output amplitudes, were optimized in logarithmic space to enforce positivity. Condition means were fitted with weights proportional to the square root of the number of replicates, which is equivalent to minimizing replicate-level raw-scale squared error when replicate variance is treated as equal within each condition.

The fitting objective was:

$$SSE(\theta) = \ sum_i \cdot w_i[(\,y_i - \hat{y}_i(\theta))]^2 \tag{S92}$$

where $y_i$ is the measured reporter output, $\hat{y}_i(\theta)$ is the model-predicted output, $w_i$ is the condition weight, and theta denotes the fitted parameter set. The overall raw-scale goodness of fit was calculated as:

$$R^2_{raw} \ = \ 1 \ - \ \frac{sum_i(\,y_i - \hat{y}_i(\theta))^2}{sum_i(\,y_i - y_{bar})^2} \tag{S93}$$

For diagnostic purposes, model performance was also evaluated on the log-transformed output scale, by variant-level $R^2$, and by inspection of the largest residuals. The raw-scale $R^2$ mainly reflects reconstruction of high-output ON states, whereas log-scale diagnostics more strongly penalize relative errors in low-output OFF and leaky states. Because the same endpoint dataset was used for fitting and evaluation, the endpoint $R^2$ values quantify in-sample reconstruction accuracy rather than held-out predictive performance.

A stricter control model was also considered in which all DBDs shared a single global output amplitude A. This constrained model provided a lower raw-scale $R^2$ than the primary DBD-specific output-window model, indicating that DBD/promoter contexts contribute measurably to the mammalian output window. Therefore, the primary model used DBD-specific $T_0$ and A while maintaining shared $K_A$ for the same DBD and shared $Q_{lbd}$ for the same LBD.

The fitted mammalian $K_A$ and $Q_{lbd}$ parameters were then used to project experimentally tested CIC-TF combinations into a two-dimensional mammalian design space. In this projection, the x-axis represents:

$$Q_{LBD} = \frac{K_{d1,l} \cdot K_{b,l}^2}{K_{d0,l}}$$

and the y-axis represents:

$$K_A$$

The design-space background was calculated independently from the endpoint-fitting procedure. For each grid point ( $Q_{lbd}$, $K_A$), $K_{d0}$, $K_b$, and $K_{d1}$ were assigned according to a specified decomposition mode. For the fixed-$K_{d1}/K_{d0}$ landscape, $K_{d1}/K_{d0}$ was fixed at:

$$K_{d1} \ / \ K_{d0} = 10^5,$$

and $K_b$ was varied according to:

$$K_b = \sqrt{Q_{LBD}/10\text{^}5}$$

For each ($Q_{LBD}$, $K_A$) coordinate, OFF and ON outputs were calculated using the full dimeric mass-balance model. Let M denote the free TF monomer concentration and let:

$$L = K_b \cdot c_I \tag{S94}$$

The mass-balance equation can be written in quadratic form as:

$$c_{TF.all} = B \cdot M + A \cdot M^2 \tag{S95}$$

where:

$$A = 2K_{d0} + 2K_{d1} \cdot L^2 \tag{S96}$$

and:

$$B = 1 + L \tag{S97}$$

Thus, the free monomer concentration is:

$$M = \frac{-B + \sqrt{B^2 + 4Ac_{TF,all}}}{2A} \tag{S98}$$

The effective TF pool is then:

$$c_{TF,eff} = K_{d0} \cdot M^2 + K_{d1} \cdot (LM)^2 \tag{S99}$$

OFF and ON states were calculated with:

$$c_I = 0$$

and:

$$c_I = 100$$

respectively. Promoter output was calculated from the effective TF pool using the occupancy relation in Eq. S90 and the transcriptional-output function defined in Eq. S11.

For the design-space landscape, a common output window was used to compare all grid points under the same promoter context. The scan used:

$$T_0 = 0.035$$

and:

$$T_{max} = 13.61$$

At each $(Q_{LBD},\ K_A)$ coordinate, $c_{TF,all}$ was scanned over the calibrated mammalian expression range:

$c_{TF,all}$ in [0.01, 3.02] RPU.

For each scanned $c_{TF,all}$, the model computed:

$$\Delta Exp = T_{on} - T_{off}$$

and:

$$Foldchange = \frac{T_{on}}{T_{off}}$$

The design objective was defined as:

$$J = log_{10}(\Delta Exp \cdot Foldchange) - \ log_{10}\left(\frac{c_{TF,all}}{c_{TF,ref}}\right) \qquad (S100)$$

where:

$$c_{TF,ref} = 1.0\ RPU$$

The plotted value for each grid point was the maximum value of J over the scanned $c_{TF,all}$ range:

$$J_{max}(Q_{LBD}, K_A) = \max_{c_{TF,all} \in [0.01,3.02]} J\left(\ Q_{lbd}, K_A, c_{TF,all}\right) \qquad (S101)$$

The corresponding optimal TF abundance was recorded as:

$$c^*_{TF,all} = arg \max_{c_{TF,all} \in [0.01,3.02]} J(\, Q_{LBD}, K_A, c_{TF,all}) \qquad (S102)$$

Therefore, the mammalian landscape represents the best achievable objective value at each $(Q_{LBD}, K_A)$ coordinate after optimizing TF abundance within the experimentally calibrated mammalian input range. Experimentally tested mammalian CIC-TFs were overlaid according to their OFF/ON-estimated $Q_{lbd}$ and $K_A$ values. Prioritized designs were selected from dose-response-characterized candidates based on the composite experimental score:

$$log_{10}(\Delta Exp \cdot Foldchange)$$

This workflow converts sparse stable-cell endpoint measurements into a reusable mammalian parameter map and uses that map to guide quantitative selection of CIC-TFs for full dose-response characterization.

**Supplementary Note 8.3.3: Accurate full-curve fitting of mammalian DBD and LBD parameters.**

The mammalian full dose-response data were fitted with the dimeric mass-conservation model defined in Eqs. S94–S99. The resulting transcription-factor state was propagated through the single-operator occupancy relation (Eq. S90), the seven-operator promoter relation (Eq. S67) and the transcriptional-output relation (Eq. S11). Thus, the fitting procedure used the established mechanistic equations without introducing an additional empirical response function.

Parameter sharing followed the modular architecture of the chimeric transcription factors. All constructs containing the same ligand-binding domain (LBD) shared $K_{d0}$, $K_b$ and $K_{d1}$, whereas all constructs containing the same DNA-binding domain (DBD) shared $K_A$ and $T_{0,\text{variant}}$. The $K_A$ value for $\text{LexA}_{\text{ec87}}$ was fixed at 9.15. The maximum output was fixed at $T_{\max} = 13.61$ RPU and the operator number was fixed at $n = 7$. All other $K_A$ and $T_{0,\text{variant}}$ values were estimated jointly with the LBD parameters. With 13 LBD groups and 11 DBD groups, including one fixed DBD $K_A$ value, the model contained 60 free parameters.

Positive-valued parameters were transformed to $\log_{10}$ coordinates before optimization. Bounds were $-16 \leq \log_{10}(K_{d0}), \log_{10}(K_b), \log_{10}(K_{d1}) \leq 12$ and $-8 \leq \log_{10}(K_A) \leq 8$; each $T_{0,\text{variant}}$ was constrained to $10^{-8} \leq T_{0,\text{variant}} \leq T_{\max} - 10^{-8}$. The objective contained only $\log_{10}$(RPU) residuals. No raw-scale residual term or endpoint-shape prior was included. For observation *i* from sensor *j*, the weighted residual was

$$r_{ji} = \sqrt{w_j}[\log_{10}(\hat{T}_{ji}) - \log_{10}(T_{ji})] \qquad (S103)$$

where $T_{ji}$ and $\hat{T}_{ji}$ denote the measured and predicted outputs, respectively, and $w_j$ is a sensor-level weight. All observations from a given sensor received the same weight, so replicate-rich sensors did not receive a separate point-level weighting rule.

The primary acceptance criterion was $R^2_{\log_{10},j} \geq 0.5$ for every sensor. To provide a small numerical margin, the internal target was 0.5005. After each fit, the shortfall $\delta_j$ for sensor $j$ was calculated as

$$\delta_j = \max(0, 0.5005 - R^2_{\log_{10},j}) \qquad \text{(S104)}$$

and the weight of a sensor below the target was updated according to

$$w_j \leftarrow w_j \times 1.35^{1+4\delta_j} \qquad \text{(S105)}$$

whereas the weight remained unchanged when the target was met. Weights were initialized to one, and the fit–evaluate–reweight cycle was repeated for at most 24 rounds. The final feasibility threshold was $R^2_{\log_{10},j} \geq 0.5$ for every sensor.

Each weighted least-squares problem was solved with scipy.optimize.least_squares using the trust-region reflective algorithm, max_nfev = 50,000, ftol = xtol = gtol = $10^{-12}$ and x_scale = 'jac'. Two deterministic 60-parameter initializations were evaluated: a pure-log initialization and a balanced raw/log initialization inherited from the preceding fit. The initialization affected only the starting point; both candidates were optimized with the same final objective based on $\log_{10}$ residuals and the same constraints.

A candidate was retained only if it satisfied the per-sensor $R^2_{\log_{10},j}$ threshold. Among feasible candidates, the reported parameter set was the one with the smallest pooled, unweighted $\log_{10}$ mean-squared error

$$\mathrm{MSE}_{\log_{10}} = \frac{1}{N}\sum_j \sum_i [\log_{10}(\hat{T}_{ji}) - \log_{10}(T_{ji})]^2 \qquad \text{(S106)}$$

where $N$ is the total number of fitted observations. Raw-scale $R^2_{\mathrm{raw}}$ values were calculated only as descriptive diagnostics and did not contribute to the optimization objective or candidate ranking.

**Supplementary Note 8.4: Multi-objective model evaluation of nine-sensor CIC-TF SETs**

**Supplementary Note 8.4.1: Scope of the panel-level design analysis**

The multi-sensor analysis in Fig. 7B was formulated as a panel-level design-space comparison rather than as independent optimization of individual sensors. Nine CIC ligand-binding domains (LBDs) were fixed: $LasR_{177}$, $RpaR_{179}$, $ER_{282–595}$, $DHBR_{282–595}$, acVHH, $CinR_{179}$, $PR_{645–914}$, $BjaR_{180}^{S107R}$ and $LuxR_{183}^{S116Y}$. Each LBD was assigned to one DNA-binding domain (DBD)/operator module and one total TF input level.

Eleven non-redundant DBD/operator modules were included: $LexA_{bs94}$, $CI434_{70}$, $LexA_{ec87}$, $LexA_{xa101}$, $HKCI_{84}$, $DeoR_{92}$, $PurR_{60}$, $LexA_{gs91}$, $LexA_{fs104}$, $CI_{94}$ OL1 and $LexA_{mm115}$. Each DBD/operator module could be used at most once within a nine-sensor SET. DBD-specific $K_A$ and $T_0$ values were used for the current analysis. Candidate TF inputs were evaluated over 0.2–10 RPU.

The experimentally implemented nine-CIC architecture was additionally evaluated using its fixed DBD–LBD assignments and implemented TF-input values. These values were not forced to lie within the 0.2–10 RPU sampling interval; in particular, the implemented $LuxR_{183}^{S116Y}$ cassette corresponded to 11.57 RPU. The Constructed SET should therefore be interpreted as a model-evaluated implemented reference rather than as an ordinary input-range-constrained sampled candidate.

**Supplementary Note 8.4.2: Single-sensor CIC model**

For each DBD–LBD–input combination, OFF- and ON-state outputs were recalculated using the dimeric mass-conservation and active-pool model defined in Supplementary Note 3 (Eqs. S32–S47), together with the promoter-occupancy and transcriptional-output functions defined in Supplementary Note 1 (Eqs. S10–S11). The DBD-specific $K_A$ and $T_0$ values, LBD-specific $K_{d0}$, $K_b$ and $K_{d1}$ values, total TF input and cognate inducer concentration were supplied to the same deterministic model used for Fig. 7B.

The promoter-output range used $T_{max}$ = 35.85 RPU and the DBD/operator-specific basal output $T_0$. $LuxR_{183}^{S116Y}$ was evaluated with two equivalent operator sites; all other CIC sensors were evaluated with one operator site.

The cognate ON-state reference concentrations were 10 μM 3OC12-HSL for $LasR_{177}$, 100 μM pC-HSL for $RpaR_{179}$, 1 μM β-estradiol for $ER_{282–595}$, 100 μM DHB for $DHBR_{282–595}$, 1 μM caffeine for acVHH, 1 μM 3OHC14-HSL for $CinR_{179}$, 100 μM RU486 for $PR_{645–914}$, 1,000 μM IV-HSL for $BjaR_{180}^{S107R}$ and 1,000 μM 3OC6 for $LuxR_{183}^{S116Y}$. OFF-state output was calculated at zero cognate inducer.

**Supplementary Note 8.4.3: Sensor-level performance metrics**

For each sensor, leakage was the model-predicted OFF output, expression shift was ON minus OFF, and fold-change was the ON/OFF ratio, following the performance-metric definitions in Supplementary Note 7.

A numerical floor of $10^{-9}$ was applied where required to avoid division by zero or logarithms of zero.

In the current Fig. 7B implementation, an explicit non-cognate response model was included for the cross-response of $ER_{282–595}$ to DHB. This off-target response was evaluated at 100 μM DHB using the cross-ligand parameters implemented in the plotting script. The corresponding ER orthogonality index was calculated as

$$OI_{\mathrm{ER}} = \log_{10}\left(\frac{\Delta \mathrm{Exp}_{\beta\text{-estradiol}}}{\Delta \mathrm{Exp}_{\mathrm{DHB}}}\right).$$

For visualization and panel scoring, OI values were bounded to the interval 2.5–4.5. Because other off-target relationships were not explicitly parameterized in this plotting implementation, the remaining sensors were assigned the upper display value of 4.5. Accordingly, the SET-level minimum OI in Fig. 7B should be interpreted as the limiting orthogonality under the explicitly modeled ER–DHB cross-response rather than as a complete quantitative prediction of every possible pairwise ligand cross-response.

**Supplementary Note 8.4.4: Construction of the sampled SET design space**

Each of the 99 possible DBD–LBD pairs was first evaluated at 512 logarithmically spaced TF-input values between 0.2 and 10 RPU. To reduce the pair-level input space before panel assembly, up to six representative input states were retained for each DBD–LBD pair. These included the states maximizing $\log_{10}(\Delta\mathrm{Exp})$, maximizing $\log_{10}(FC)$, minimizing TF input, minimizing leakage, the state closest to 1 RPU and the state maximizing a pair-level composite score

$$J_{\mathrm{pair}} = 0.365 z_{\Delta\mathrm{Exp}} + 0.365 z_{FC} - 0.135 z_{\mathrm{input}} - 0.135 z_{\mathrm{leak}},$$

where each $z$ term was min–max normalized within the 512-point input landscape of the corresponding DBD–LBD pair. Duplicate anchor states were retained only once.

Nine-sensor SETs were then generated reproducibly using random seed 20260808. For each SET, nine distinct DBD/operator modules were sampled without replacement from the 11-member DBD pool and assigned to the nine fixed LBDs. One retained input state was then sampled for each resulting DBD–LBD pair. A total of 15,000 model-evaluated candidate SETs were generated.

This sampling procedure provides a reproducible representation of the multi-sensor trade-off landscape; it does not constitute exhaustive enumeration of the complete $11P9$ architecture space or of the full nine-dimensional continuous TF-input space.

**Supplementary Note 8.4.5: SET-level objectives and Pareto analysis**

For a nine-sensor SET $S$, the panel-level expression-shift score was defined as

$$D(S) = \frac{1}{9}\sum_{i=1}^{9} \log_{10}(\Delta\mathrm{Exp}_i),$$

and the panel-level fold-change score as

$$F(S) = \frac{1}{9}\sum_{i=1}^{9} \log_{10}(FC_i).$$

The limiting orthogonality score was

$$O(S) = \min_i(OI_i),$$

the total TF input burden was

$$B(S) = \sum_{i=1}^{9} C_{\mathrm{TF,all},i},$$

and the maximum sensor leakage was

$$L(S) = \max_i(T_{\mathrm{OFF},i}).$$

Five objectives were considered simultaneously: maximizing $D$, maximizing $F$, maximizing $O$, minimizing $B$, and minimizing $L$. No hard leakage threshold was applied in this implementation. A SET was considered Pareto-nondominated if no other evaluated SET was at least as favorable in all five objectives and strictly better in at least one.

A separate scalar score was used to identify one balanced representative:

$$J = 0.27z_D + 0.27z_F + 0.27z_O - 0.095z_B - 0.095z_L,$$

where each quantity was min–max normalized across the evaluated SET pool. This scalar score was used only to nominate the representative Optimal SET and was not itself the definition of Pareto dominance.

**Supplementary Note 8.4.6: Representative SETs and the implemented reference**

Four SETs were highlighted in Fig. 7B. The ΔExp-biased SET maximized $D(S)$, the FC-biased SET maximized $F(S)$, and the balanced Optimal SET maximized the scalar score $J$. The experimentally implemented nine-CIC portion of CIC-12 was evaluated separately using its fixed DBD–LBD architecture and implemented TF-input vector and is shown as the Constructed SET.

The implemented architecture comprised $\mathrm{HKCI_{84}}$–$\mathrm{LasR_{177}}$, $\mathrm{LexA_{xa101}}$–$\mathrm{RpaR_{179}}$, $\mathrm{LexA_{ec87}}$–$\mathrm{ER_{282–595}}$, $\mathrm{LexA_{mm115}}$–$\mathrm{DHBR_{282–595}}$, $\mathrm{LexA_{gs91}}$–acVHH, $\mathrm{CI434_{70}}$–$\mathrm{CinR_{179}}$, $\mathrm{CI_{94}}$ (OL1)–$\mathrm{PR_{645–914}}$, $\mathrm{PurR_{60}}$–$\mathrm{BjaR_{180}^{S107R}}$ and $\mathrm{LexA_{bs94}}$–$\mathrm{LuxR_{183}^{S116Y}}$. The corresponding implemented TF-input values were 0.960, 2.086, 1.786, 3.133, 1.148, 0.980, 0.668, 2.062 and 11.573 RPU, respectively.

Under the current model calculation, the four highlighted SETs had the following panel metrics:

- **ΔExp-biased:** mean $\log_{10}$(ΔExp) = 1.260; mean $\log_{10}$(FC) = 1.506; minimum OI = 3.00; total TF input = 65.8 RPU; maximum leak = 6.45 RPU.
- **FC-biased:** mean $\log_{10}$(ΔExp) = 0.770; mean $\log_{10}$(FC) = 2.224; minimum OI = 3.69; total TF input = 24.8 RPU; maximum leak = 0.11 RPU.
- **Optimal:** mean $\log_{10}$(ΔExp) = 0.772; mean $\log_{10}$(FC) = 2.079; minimum OI = 4.47; total TF input = 17.4 RPU; maximum leak = 0.26 RPU.
- **Constructed:** mean $\log_{10}$(ΔExp) = 1.066; mean $\log_{10}$(FC) = 2.098; minimum OI = 3.71; total TF input = 24.4 RPU; maximum leak = 0.82 RPU.

Within the reproducibly sampled design space, no sampled candidate dominated the Constructed SET across all five objectives. Relative to the balanced Optimal SET, the implemented architecture retained a larger predicted expression shift and similar mean fold-change, but at the cost of higher total TF input and maximum leakage and a lower limiting OI. Thus, the highlighted SETs illustrate distinct operating regimes in the multi-objective design space rather than a single universally superior architecture.

**Supplementary Note 8.4.7: Interpretation of the panel analysis**

The purpose of this analysis is to test whether independently characterized CIC component parameters can be recombined to compare complete multi-sensor programs under a common quantitative framework. Fig. 7B therefore provides a model-based benchmark of alternative panel-level trade-offs among expression shift, fold-change, orthogonality, TF dosage and basal leakage.

The Constructed SET represents the experimentally implemented CIC-12 architecture evaluated under the same final model and should not be interpreted as experimental ΔExp, fold-change, leakage or OI measurements. Experimental characterization of the implemented chassis is presented separately in Fig. 7C and Supplementary Fig. 14.

**Supplementary Note 8.5: Gaussian process − guided iterative optimization of target production.**

The present study employs a Gaussian process-based iterative optimization framework[7] to enhance target production through an initial DOE (Latin sampling from a 3-level factorial grid) followed by two GP-guided optimization rounds. The methodological approach integrates Bayesian optimization principles with sequential experimental design to efficiently navigate the high-dimensional parameter space.

Initial data preprocessing involved standardization of all input variables using StandardScaler to ensure uniform feature scaling. The Gaussian process regression model was constructed using a composite kernel consisting of a constant kernel multiplied by a radial basis function kernel, formally expressed as:

$$k(x_i, x_j) = C \cdot \exp\left(-\frac{\| x_i - x_j \|^2}{2l^2}\right) \qquad (S107)$$

where C represents the signal variance and $l$ denotes the length-scale parameter. The hyperparameters of this kernel were optimized through 50 restarts of the optimizer to avoid local minima, with a regularization parameter α set to $10^{-3}$ to account for observational noise.

To initialize the surrogate model with experimentally grounded data, we first performed a structured global design over the eight-dimensional regulatory space ($LasR_{177}$, $ER_{282\text{-}595}$, PR, XlnR, $DHBR_{282\text{-}595}$, $RpaR_{179}$, LacI, and $CinR_{179}$). Each dimension was discretized into three levels (low/medium/high), yielding a $3^8$ = 6,561-candidate grid. From this grid, 96 conditions were selected as the planned design-of-experiments set using a Latinized, stratified sampling scheme to provide balanced coverage across dimensions. During experimental implementation, 32 of these conditions were excluded before model fitting because dispensing errors caused the implemented inducer programs to differ from their planned designs. The remaining 64 conditions were correctly dispensed, experimentally quantified, and retained as the initial training set ($X_0$, $y_0$). This dataset was then used to fit the Gaussian process regression model, which subsequently served as the surrogate predictor for large-scale in silico evaluation of candidate points in later rounds.

**GP-guided optimization round 1.**

Using the fitted GP model, we generated 50,000 candidate points uniformly within the same parameter bounds and computed both the predictive mean $\mu(x)$and uncertainty $\sigma(x)$for each candidate. To prioritize reliable predictions, we applied an uncertainty filter ($\sigma(x) < \sigma_{\text{threshold}} = 0.2$) and selected the 64 candidates with the highest predicted yields from the retained set for experimental validation.

**GP-guided optimization round 2.**

We next updated the surrogate model by retraining the GP on all accumulated measurements (the initial DOE dataset plus the 64 designs validated in optimization round 1). To refine solutions locally, we centered sampling around the validated high-performing designs from round 1 and generated perturbed candidates according to

$$x_i' = x_i + \delta \qquad (S108)$$

where $\delta \sim U(-0.1, 0.1)$is an element-wise uniform perturbation vector. All perturbed points were constrained to the predefined parameter bounds via clipping, $x_i' = \text{clip}(x_i', \text{bounds}_{\text{min}}, \text{bounds}_{\text{max}})$. For each candidate, the updated GP returned the predictive mean $\mu(x)$and uncertainty $\sigma(x)$. We applied a stricter uncertainty filter ($\sigma(x) < \sigma_{\text{threshold}} = 0.15$) and the 64 remaining candidates with the highest predicted yields were retained as a provisional computational shortlist. To reduce experimental redundancy and improve coverage of the regulatory space, this shortlist was subsequently pruned by removing candidates with highly similar eight-dimensional gene-expression profiles. After similarity-based redundancy filtering, 25 non-redundant candidates remained and were selected for experimental validation. Thus, 64 denotes the size of the GP-ranked computational shortlist, whereas 25 denotes the number of conditions that were ultimately tested experimentally in this round.

**Supplementary Note 9: Quantitative test and model fitting for activator phenotype.**

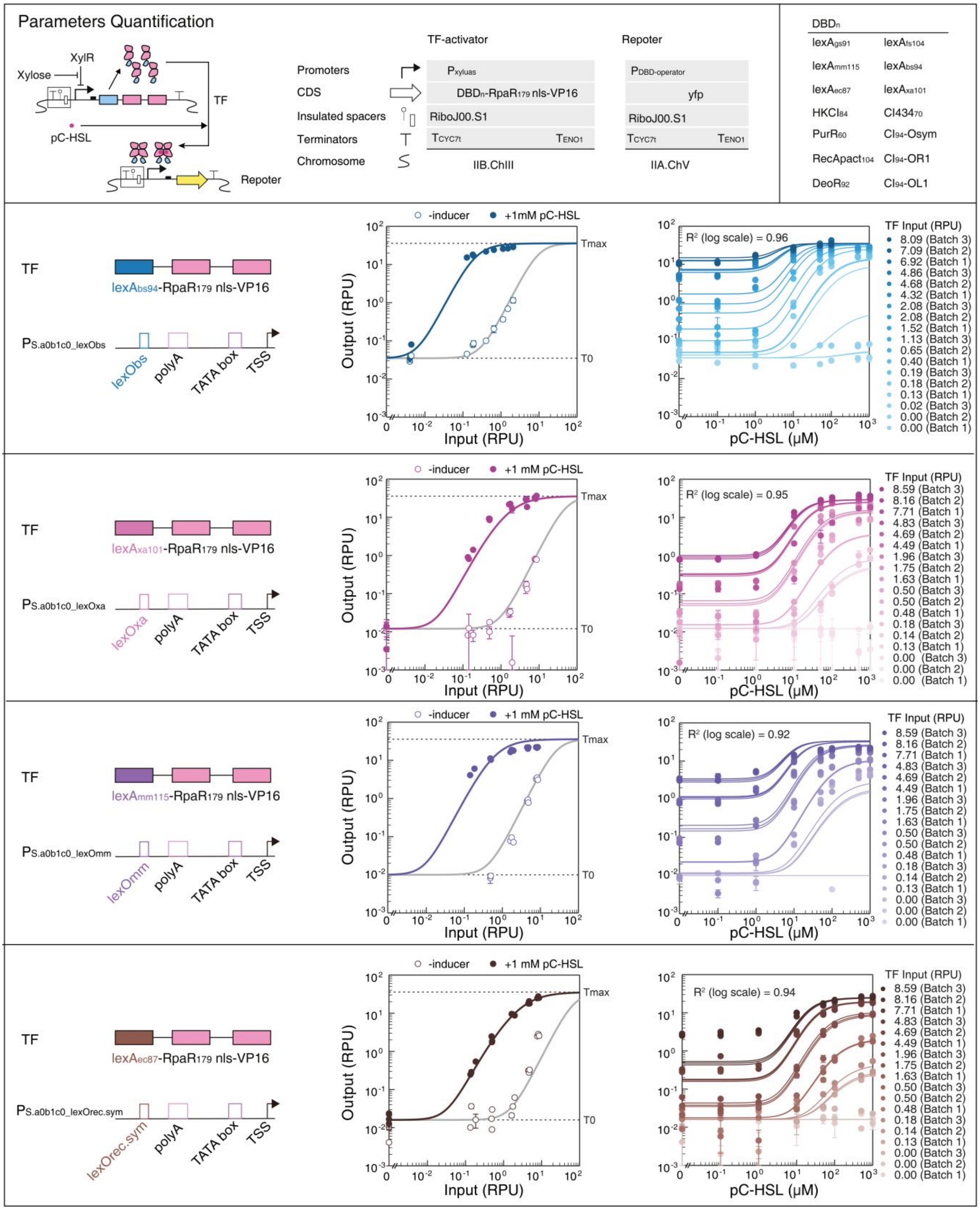

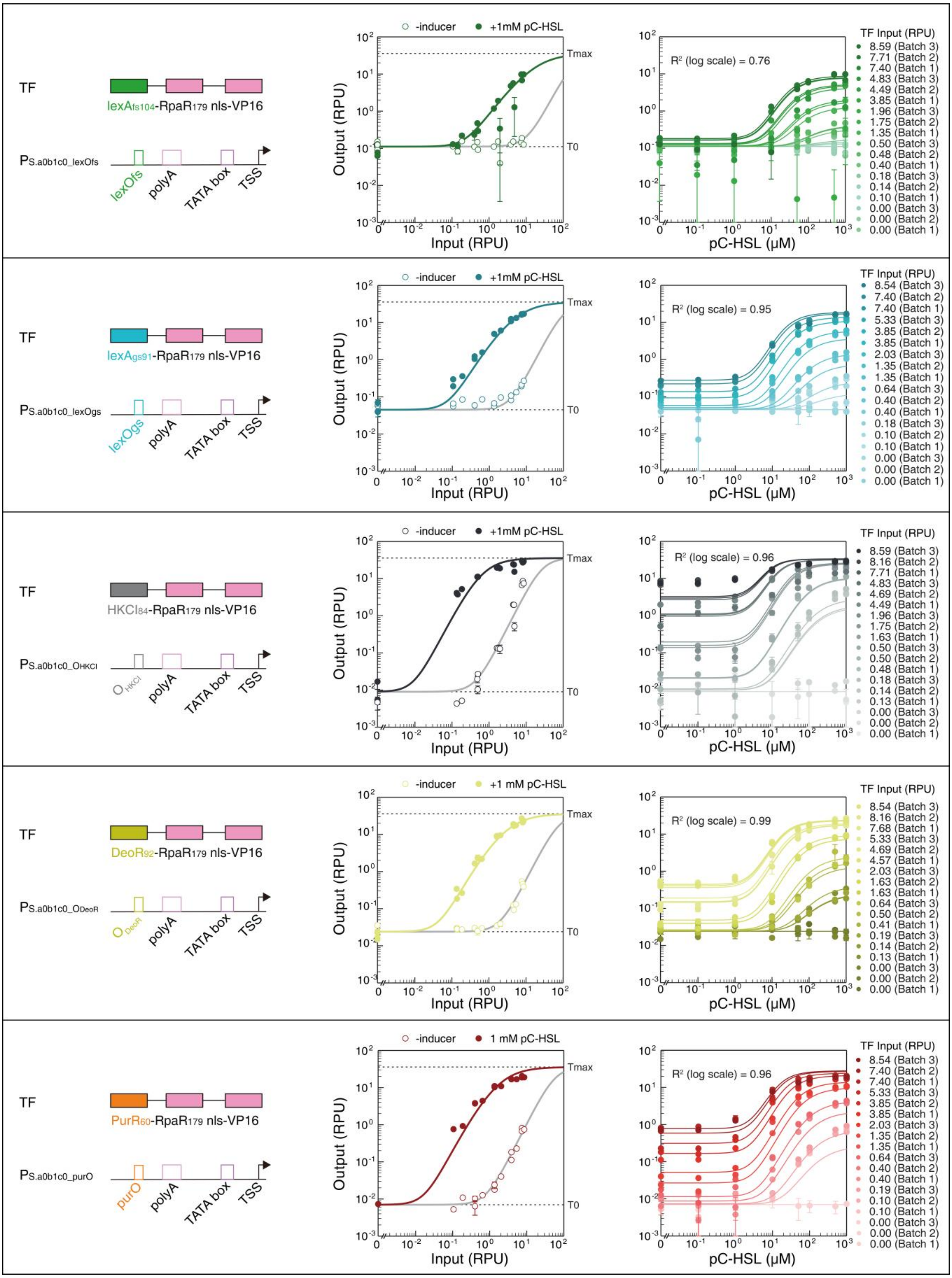

TF
lexAfs104-RpaR179 nls-VP16
PS.a0b1c0_lexOfs
lexOfs
polyA
TATA box
TSS
-inducer
+1mM pC-HSL
Output (RPU)
Input (RPU)
Tmax
T0
R² (log scale) = 0.76
pC-HSL (μM)
TF Input (RPU)
8.59 (Batch 3)
7.71 (Batch 2)
7.40 (Batch 1)
4.83 (Batch 3)
4.49 (Batch 2)
3.85 (Batch 1)
1.96 (Batch 3)
1.75 (Batch 2)
1.35 (Batch 1)
0.50 (Batch 3)
0.48 (Batch 2)
0.40 (Batch 1)
0.18 (Batch 3)
0.14 (Batch 2)
0.10 (Batch 1)
0.00 (Batch 3)
0.00 (Batch 2)
0.00 (Batch 1)
TF
lexAgs91-RpaR179 nls-VP16
PS.a0b1c0_lexOgs
lexOgs
polyA
TATA box
TSS
-inducer
+1mM pC-HSL
Output (RPU)
Input (RPU)
Tmax
T0
R² (log scale) = 0.95
pC-HSL (μM)
TF Input (RPU)
8.54 (Batch 3)
7.40 (Batch 2)
7.40 (Batch 1)
5.33 (Batch 3)
3.85 (Batch 2)
3.85 (Batch 1)
2.03 (Batch 3)
1.35 (Batch 2)
1.35 (Batch 1)
0.64 (Batch 3)
0.40 (Batch 2)
0.40 (Batch 1)
0.18 (Batch 3)
0.10 (Batch 2)
0.10 (Batch 1)
0.00 (Batch 3)
0.00 (Batch 2)
0.00 (Batch 1)
TF
HKCI84-RpaR179 nls-VP16
PS.a0b1c0_OHKCI
OHKCI
polyA
TATA box
TSS
-inducer
+1mM pC-HSL
Output (RPU)
Input (RPU)
Tmax
T0
R² (log scale) = 0.96
pC-HSL (μM)
TF Input (RPU)
8.59 (Batch 3)
8.16 (Batch 2)
7.71 (Batch 1)
4.83 (Batch 3)
4.69 (Batch 2)
4.49 (Batch 1)
1.96 (Batch 3)
1.75 (Batch 2)
1.63 (Batch 1)
0.50 (Batch 3)
0.50 (Batch 2)
0.48 (Batch 1)
0.18 (Batch 3)
0.14 (Batch 2)
0.13 (Batch 1)
0.00 (Batch 3)
0.00 (Batch 2)
0.00 (Batch 1)
TF
DeoR92-RpaR179 nls-VP16
PS.a0b1c0_ODeoR
ODeoR
polyA
TATA box
TSS
-inducer
+1 mM pC-HSL
Output (RPU)
Input (RPU)
Tmax
T0
R² (log scale) = 0.99
pC-HSL (μM)
TF Input (RPU)
8.54 (Batch 3)
8.16 (Batch 2)
7.68 (Batch 1)
5.33 (Batch 3)
4.69 (Batch 2)
4.57 (Batch 1)
2.03 (Batch 3)
1.63 (Batch 2)
1.63 (Batch 1)
0.64 (Batch 3)
0.50 (Batch 2)
0.41 (Batch 1)
0.19 (Batch 3)
0.14 (Batch 2)
0.13 (Batch 1)
0.00 (Batch 3)
0.00 (Batch 2)
0.00 (Batch 1)
TF
PurR60-RpaR179 nls-VP16
PS.a0b1c0_purO
purO
polyA
TATA box
TSS
-inducer
1 mM pC-HSL
Output (RPU)
Input (RPU)
Tmax
T0
R² (log scale) = 0.96
pC-HSL (μM)
TF Input (RPU)
8.54 (Batch 3)
7.40 (Batch 2)
7.40 (Batch 1)
5.33 (Batch 3)
3.85 (Batch 2)
3.85 (Batch 1)
2.03 (Batch 3)
1.35 (Batch 2)
1.35 (Batch 1)
0.64 (Batch 3)
0.40 (Batch 2)
0.40 (Batch 1)
0.19 (Batch 3)
0.10 (Batch 2)
0.10 (Batch 1)
0.00 (Batch 3)
0.00 (Batch 2)
0.00 (Batch 1)

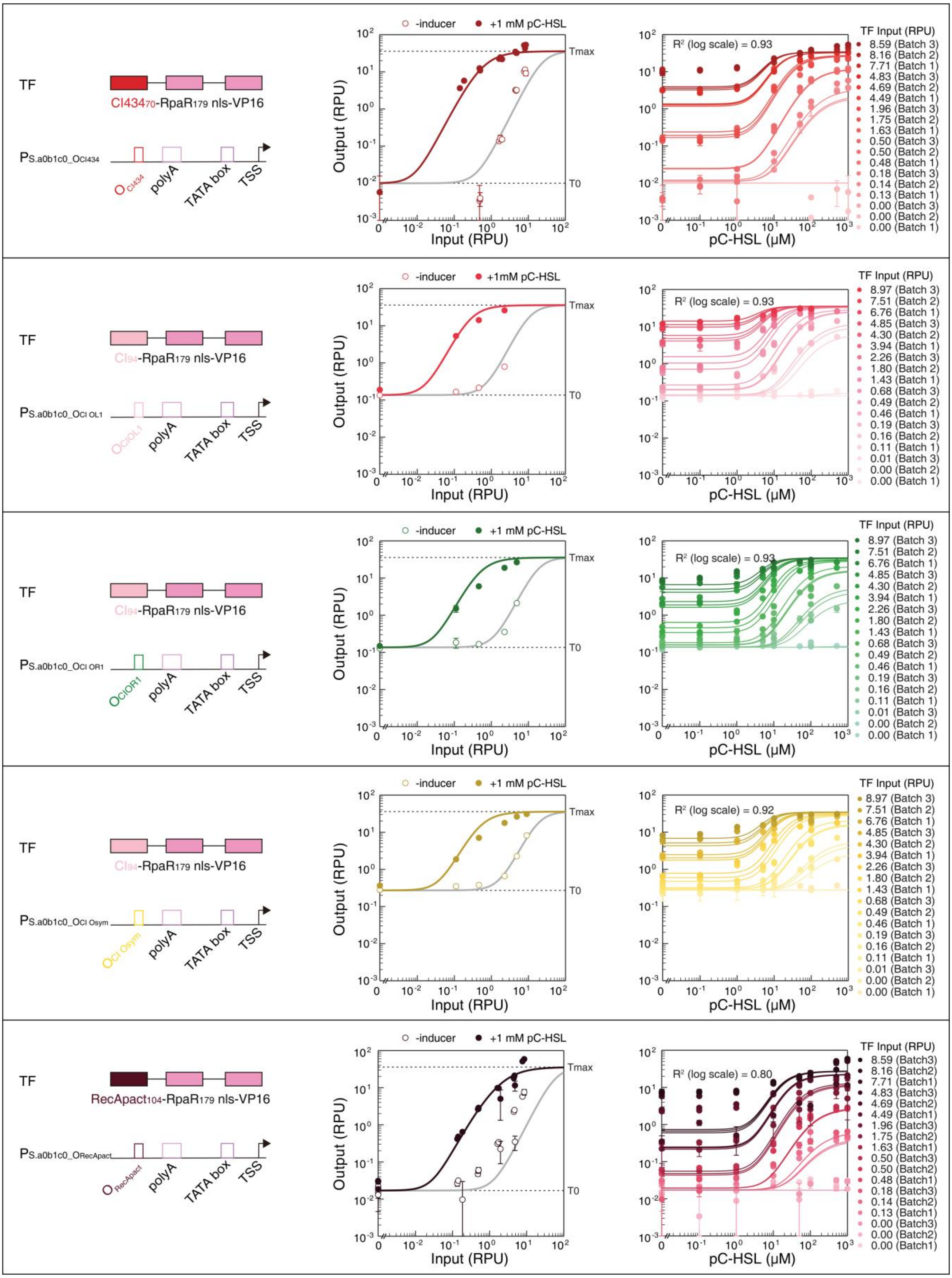

TF
CI434₇₀-RpaR₁₇₉ nls-VP16
PS.a0b1c0_OCI434
OCI434
polyA
TATA box
TSS
-inducer
+1 mM pC-HSL
Output (RPU)
Input (RPU)
Tmax
T0
R² (log scale) = 0.93
pC-HSL (μM)
TF Input (RPU)
8.59 (Batch 3)
8.16 (Batch 2)
7.71 (Batch 1)
4.83 (Batch 3)
4.69 (Batch 2)
4.49 (Batch 1)
1.96 (Batch 3)
1.75 (Batch 2)
1.63 (Batch 1)
0.50 (Batch 3)
0.50 (Batch 2)
0.48 (Batch 1)
0.18 (Batch 3)
0.14 (Batch 2)
0.13 (Batch 1)
0.00 (Batch 3)
0.00 (Batch 2)
0.00 (Batch 1)
TF
CI94-RpaR179 nls-VP16
PS.a0b1c0_OCI OL1
OCIOL1
polyA
TATA box
TSS
-inducer
+1mM pC-HSL
Output (RPU)
Input (RPU)
Tmax
T0
R² (log scale) = 0.93
pC-HSL (μM)
TF Input (RPU)
8.97 (Batch 3)
7.51 (Batch 2)
6.76 (Batch 1)
4.85 (Batch 3)
4.30 (Batch 2)
3.94 (Batch 1)
2.26 (Batch 3)
1.80 (Batch 2)
1.43 (Batch 1)
0.68 (Batch 3)
0.49 (Batch 2)
0.46 (Batch 1)
0.19 (Batch 3)
0.16 (Batch 2)
0.11 (Batch 1)
0.01 (Batch 3)
0.00 (Batch 2)
0.00 (Batch 1)
TF
CI94-RpaR179 nls-VP16
PS.a0b1c0_OCI OR1
OCIOR1
polyA
TATA box
TSS
-inducer
+1 mM pC-HSL
Output (RPU)
Input (RPU)
Tmax
T0
R² (log scale) = 0.93
pC-HSL (μM)
TF Input (RPU)
8.97 (Batch 3)
7.51 (Batch 2)
6.76 (Batch 1)
4.85 (Batch 3)
4.30 (Batch 2)
3.94 (Batch 1)
2.26 (Batch 3)
1.80 (Batch 2)
1.43 (Batch 1)
0.68 (Batch 3)
0.49 (Batch 2)
0.46 (Batch 1)
0.19 (Batch 3)
0.16 (Batch 2)
0.11 (Batch 1)
0.01 (Batch 3)
0.00 (Batch 2)
0.00 (Batch 1)
TF
CI94-RpaR179 nls-VP16
PS.a0b1c0_OCI Osym
OCI Osym
polyA
TATA box
TSS
-inducer
+1 mM pC-HSL
Output (RPU)
Input (RPU)
Tmax
T0
R² (log scale) = 0.92
pC-HSL (μM)
TF Input (RPU)
8.97 (Batch 3)
7.51 (Batch 2)
6.76 (Batch 1)
4.85 (Batch 3)
4.30 (Batch 2)
3.94 (Batch 1)
2.26 (Batch 3)
1.80 (Batch 2)
1.43 (Batch 1)
0.68 (Batch 3)
0.49 (Batch 2)
0.46 (Batch 1)
0.19 (Batch 3)
0.16 (Batch 2)
0.11 (Batch 1)
0.01 (Batch 3)
0.00 (Batch 2)
0.00 (Batch 1)
TF
RecApact104-RpaR179 nls-VP16
PS.a0b1c0_ORecApact
ORecApact
polyA
TATA box
TSS
-inducer
+1 mM pC-HSL
Output (RPU)
Input (RPU)
Tmax
T0
R² (log scale) = 0.80
pC-HSL (μM)
TF Input (RPU)
8.59 (Batch3)
8.16 (Batch2)
7.71 (Batch1)
4.83 (Batch3)
4.69 (Batch2)
4.49 (Batch1)
1.96 (Batch3)
1.75 (Batch2)
1.63 (Batch1)
0.50 (Batch3)
0.50 (Batch2)
0.48 (Batch1)
0.18 (Batch3)
0.14 (Batch2)
0.13 (Batch1)
0.00 (Batch3)
0.00 (Batch2)
0.00 (Batch1)

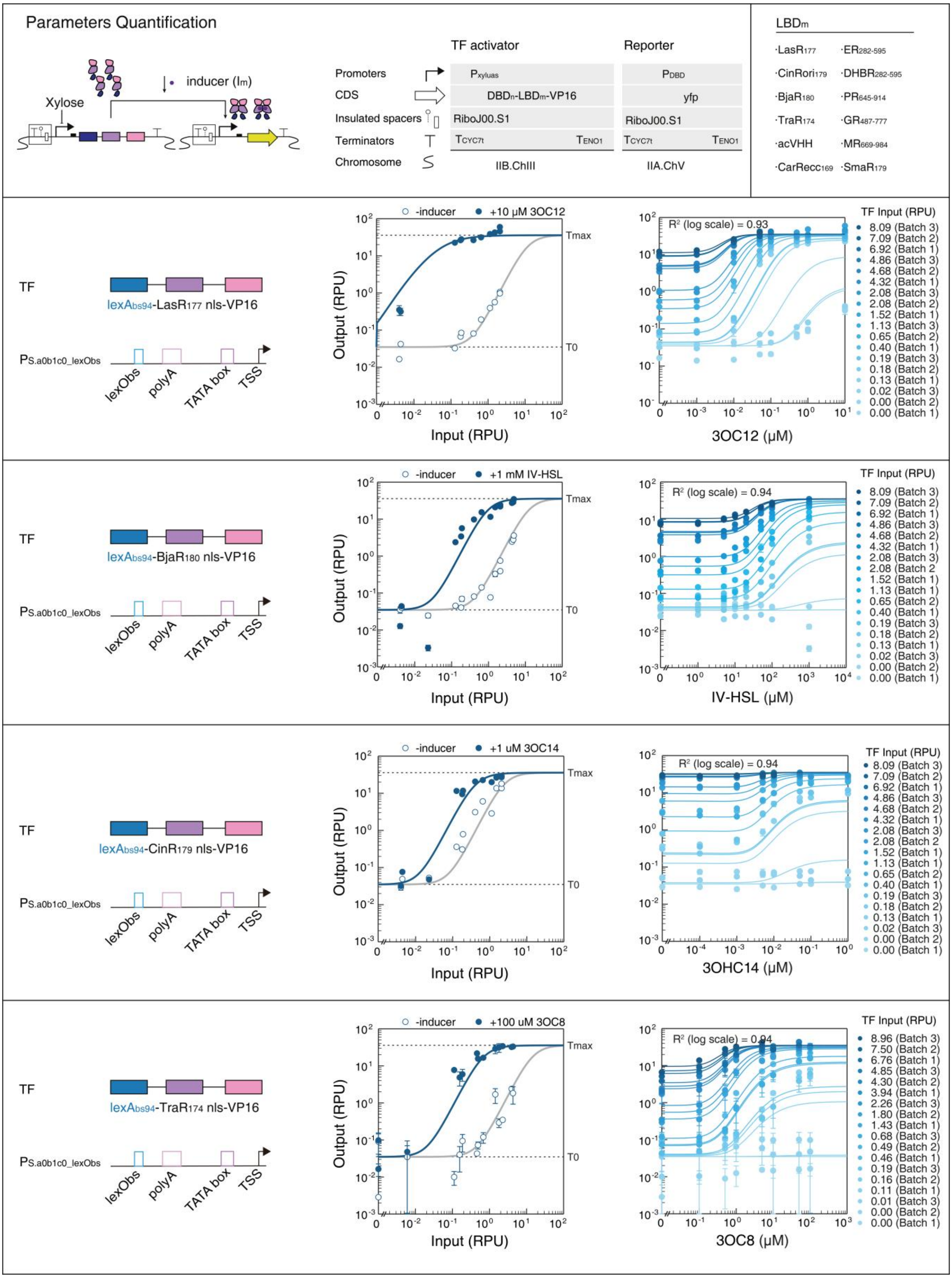

Parameters Quantification
Xylose
inducer (Im)
TF activator
Reporter
Promoters
CDS
Insulated spacers
Terminators
Chromosome
Pxyluas
DBDn-LBDm-VP16
RiboJ00.S1
TCYC7t
TENO1
IIB.ChIII
PDBD
yfp
RiboJ00.S1
TCYC7t
TENO1
IIA.ChV
LBDm
·LasR177
·ER282-595
·CinRori179
·DHBR282-595
·BjaR180
·PR645-914
·TraR174
·GR487-777
·acVHH
·MR669-984
·CarRecc169
·SmaR179
TF
lexAbs94-LasR177 nls-VP16
PS.a0b1c0_lexObs
lexObs
polyA
TATA box
TSS
-inducer
+10 μM 3OC12
Tmax
T0
Output (RPU)
Input (RPU)
R² (log scale) = 0.93
3OC12 (μM)
TF Input (RPU)
8.09 (Batch 3)
7.09 (Batch 2)
6.92 (Batch 1)
4.86 (Batch 3)
4.68 (Batch 2)
4.32 (Batch 1)
2.08 (Batch 3)
2.08 (Batch 2)
1.52 (Batch 1)
1.13 (Batch 3)
0.65 (Batch 2)
0.40 (Batch 1)
0.19 (Batch 3)
0.18 (Batch 2)
0.13 (Batch 1)
0.02 (Batch 3)
0.00 (Batch 2)
0.00 (Batch 1)
TF
lexAbs94-BjaR180 nls-VP16
PS.a0b1c0_lexObs
lexObs
polyA
TATA box
TSS
-inducer
+1 mM IV-HSL
Tmax
T0
Output (RPU)
Input (RPU)
R² (log scale) = 0.94
IV-HSL (μM)
TF Input (RPU)
8.09 (Batch 3)
7.09 (Batch 2)
6.92 (Batch 1)
4.86 (Batch 3)
4.68 (Batch 2)
4.32 (Batch 1)
2.08 (Batch 3)
2.08 (Batch 2
1.52 (Batch 1)
1.13 (Batch 3)
0.65 (Batch 2)
0.40 (Batch 1)
0.19 (Batch 3)
0.18 (Batch 2)
0.13 (Batch 1)
0.02 (Batch 3)
0.00 (Batch 2)
0.00 (Batch 1)
TF
lexAbs94-CinR179 nls-VP16
PS.a0b1c0_lexObs
lexObs
polyA
TATA box
TSS
-inducer
+1 uM 3OC14
Tmax
T0
Output (RPU)
Input (RPU)
R² (log scale) = 0.94
3OHC14 (μM)
TF Input (RPU)
8.09 (Batch 3)
7.09 (Batch 2)
6.92 (Batch 1)
4.86 (Batch 3)
4.68 (Batch 2)
4.32 (Batch 1)
2.08 (Batch 3)
2.08 (Batch 2
1.52 (Batch 1)
1.13 (Batch 3)
0.65 (Batch 2)
0.40 (Batch 1)
0.19 (Batch 3)
0.18 (Batch 2)
0.13 (Batch 1)
0.02 (Batch 3)
0.00 (Batch 2)
0.00 (Batch 1)
TF
lexAbs94-TraR174 nls-VP16
PS.a0b1c0_lexObs
lexObs
polyA
TATA box
TSS
-inducer
+100 uM 3OC8
Tmax
T0
Output (RPU)
Input (RPU)
R² (log scale) = 0.94
3OC8 (μM)
TF Input (RPU)
8.96 (Batch 3)
7.50 (Batch 2)
6.76 (Batch 1)
4.85 (Batch 3)
4.30 (Batch 2)
3.94 (Batch 1)
2.26 (Batch 3)
1.80 (Batch 2)
1.43 (Batch 1)
0.68 (Batch 3)
0.49 (Batch 2)
0.46 (Batch 1)
0.19 (Batch 3)
0.16 (Batch 2)
0.11 (Batch 1)
0.01 (Batch 3)
0.00 (Batch 2)
0.00 (Batch 1)

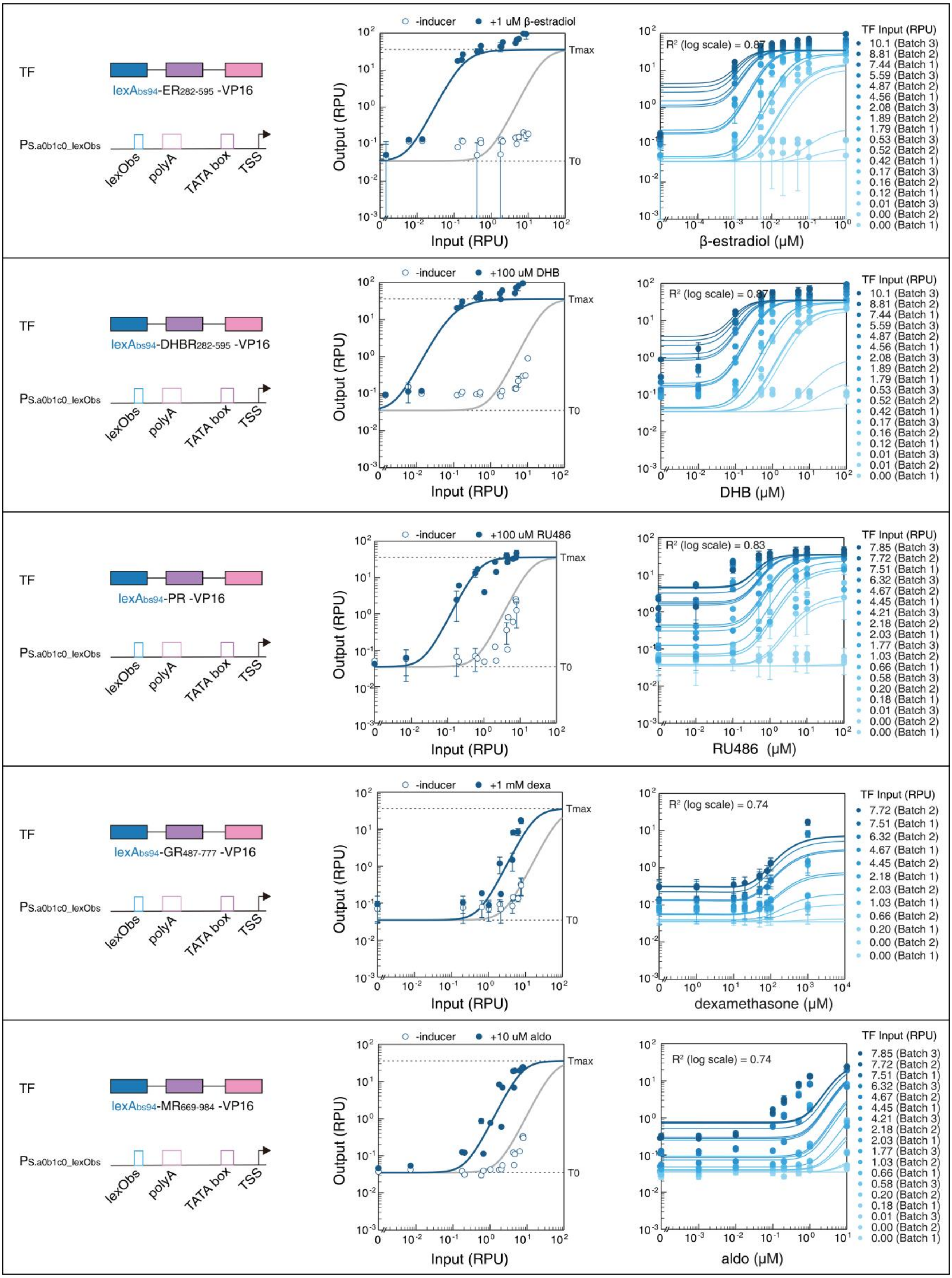

TF
lexAbs94-ER282-595 -VP16
PS.a0b1c0_lexObs
lexObs
polyA
TATA box
TSS
-inducer
+1 uM β-estradiol
Output (RPU)
Input (RPU)
Tmax
T0
R² (log scale) = 0.87
β-estradiol (μM)
TF Input (RPU)
10.1 (Batch 3)
8.81 (Batch 2)
7.44 (Batch 1)
5.59 (Batch 3)
4.87 (Batch 2)
4.56 (Batch 1)
2.08 (Batch 3)
1.89 (Batch 2)
1.79 (Batch 1)
0.53 (Batch 3)
0.52 (Batch 2)
0.42 (Batch 1)
0.17 (Batch 3)
0.16 (Batch 2)
0.12 (Batch 1)
0.01 (Batch 3)
0.00 (Batch 2)
0.00 (Batch 1)
TF
lexAbs94-DHBR282-595 -VP16
PS.a0b1c0_lexObs
lexObs
polyA
TATA box
TSS
-inducer
+100 uM DHB
Output (RPU)
Input (RPU)
Tmax
T0
R² (log scale) = 0.87
DHB (μM)
TF Input (RPU)
10.1 (Batch 3)
8.81 (Batch 2)
7.44 (Batch 1)
5.59 (Batch 3)
4.87 (Batch 2)
4.56 (Batch 1)
2.08 (Batch 3)
1.89 (Batch 2)
1.79 (Batch 1)
0.53 (Batch 3)
0.52 (Batch 2)
0.42 (Batch 1)
0.17 (Batch 3)
0.16 (Batch 2)
0.12 (Batch 1)
0.01 (Batch 3)
0.01 (Batch 2)
0.00 (Batch 1)
TF
lexAbs94-PR -VP16
PS.a0b1c0_lexObs
lexObs
polyA
TATA box
TSS
-inducer
+100 uM RU486
Output (RPU)
Input (RPU)
Tmax
T0
R² (log scale) = 0.83
RU486 (μM)
TF Input (RPU)
7.85 (Batch 3)
7.72 (Batch 2)
7.51 (Batch 1)
6.32 (Batch 3)
4.67 (Batch 2)
4.45 (Batch 1)
4.21 (Batch 3)
2.18 (Batch 2)
2.03 (Batch 1)
1.77 (Batch 3)
1.03 (Batch 2)
0.66 (Batch 1)
0.58 (Batch 3)
0.20 (Batch 2)
0.18 (Batch 1)
0.01 (Batch 3)
0.00 (Batch 2)
0.00 (Batch 1)
TF
lexAbs94-GR487-777 -VP16
PS.a0b1c0_lexObs
lexObs
polyA
TATA box
TSS
-inducer
+1 mM dexa
Output (RPU)
Input (RPU)
Tmax
T0
R² (log scale) = 0.74
dexamethasone (μM)
TF Input (RPU)
7.72 (Batch 2)
7.51 (Batch 1)
6.32 (Batch 2)
4.67 (Batch 1)
4.45 (Batch 2)
2.18 (Batch 1)
2.03 (Batch 2)
1.03 (Batch 1)
0.66 (Batch 2)
0.20 (Batch 1)
0.00 (Batch 2)
0.00 (Batch 1)
TF
lexAbs94-MR669-984 -VP16
PS.a0b1c0_lexObs
lexObs
polyA
TATA box
TSS
-inducer
+10 uM aldo
Output (RPU)
Input (RPU)
Tmax
T0
R² (log scale) = 0.74
aldo (μM)
TF Input (RPU)
7.85 (Batch 3)
7.72 (Batch 2)
7.51 (Batch 1)
6.32 (Batch 3)
4.67 (Batch 2)
4.45 (Batch 1)
4.21 (Batch 3)
2.18 (Batch 2)
2.03 (Batch 1)
1.77 (Batch 3)
1.03 (Batch 2)
0.66 (Batch 1)
0.58 (Batch 3)
0.20 (Batch 2)
0.18 (Batch 1)
0.01 (Batch 3)
0.00 (Batch 2)
0.00 (Batch 1)

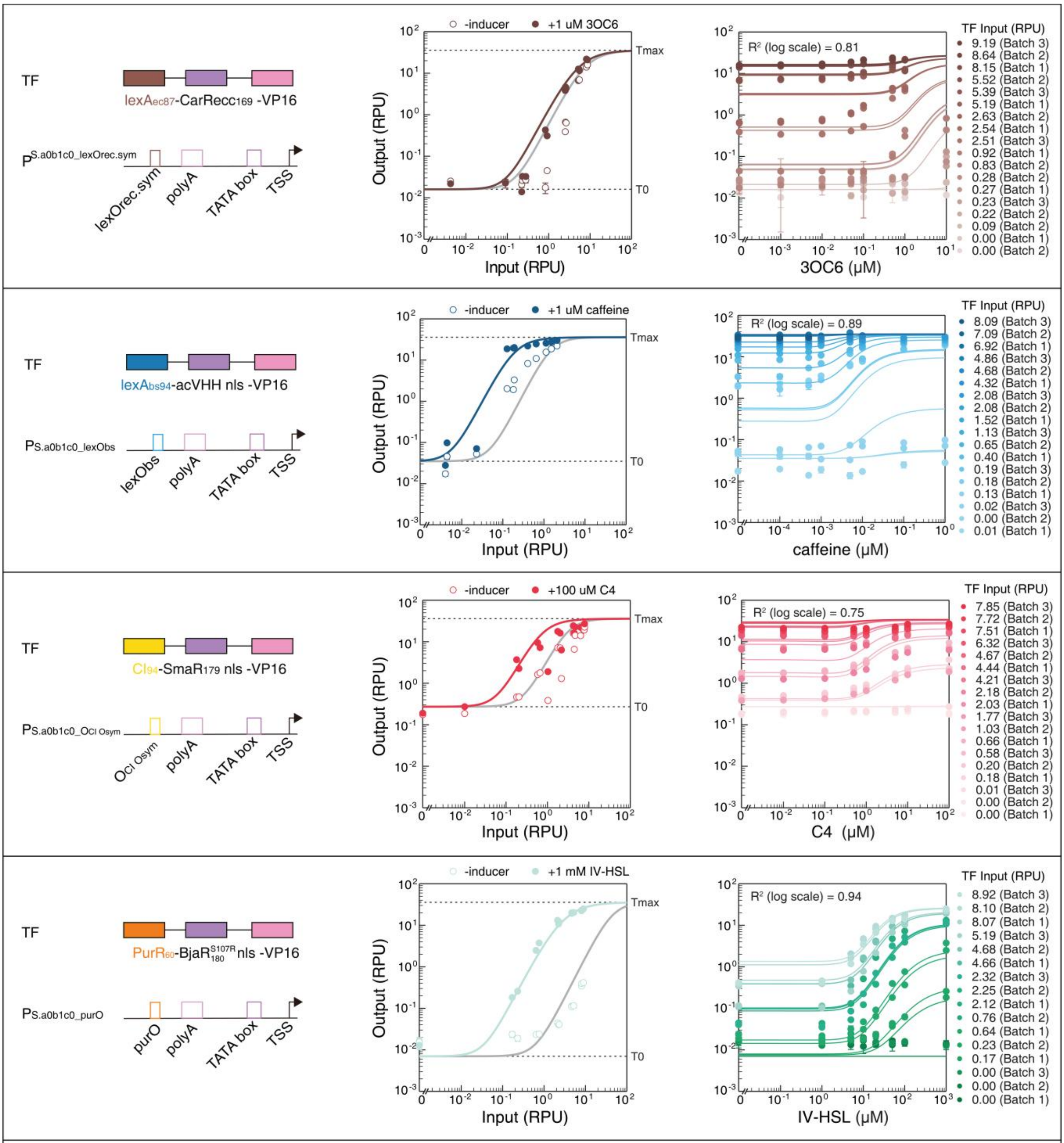

TF
lexAec87-CarRecc169 -VP16
PS.a0b1c0_lexOrec.sym
lexOrec.sym
polyA
TATA box
TSS
-inducer
+1 uM 3OC6
Output (RPU)
Input (RPU)
Tmax
T0
R² (log scale) = 0.81
3OC6 (μM)
TF Input (RPU)
9.19 (Batch 3)
8.64 (Batch 2)
8.15 (Batch 1)
5.52 (Batch 2)
5.39 (Batch 3)
5.19 (Batch 1)
2.63 (Batch 2)
2.54 (Batch 1)
2.51 (Batch 3)
0.92 (Batch 1)
0.83 (Batch 2)
0.28 (Batch 2)
0.27 (Batch 1)
0.23 (Batch 3)
0.22 (Batch 2)
0.09 (Batch 2)
0.00 (Batch 1)
0.00 (Batch 2)
TF
lexAbs94-acVHH nls -VP16
PS.a0b1c0_lexObs
lexObs
polyA
TATA box
TSS
-inducer
+1 uM caffeine
Output (RPU)
Input (RPU)
Tmax
T0
R² (log scale) = 0.89
caffeine (μM)
TF Input (RPU)
8.09 (Batch 3)
7.09 (Batch 2)
6.92 (Batch 1)
4.86 (Batch 3)
4.68 (Batch 2)
4.32 (Batch 1)
2.08 (Batch 3)
2.08 (Batch 2)
1.52 (Batch 1)
1.13 (Batch 3)
0.65 (Batch 2)
0.40 (Batch 1)
0.19 (Batch 3)
0.18 (Batch 2)
0.13 (Batch 1)
0.02 (Batch 3)
0.00 (Batch 2)
0.01 (Batch 1)
TF
CI94-SmaR179 nls -VP16
PS.a0b1c0_OCI Osym
OCI Osym
polyA
TATA box
TSS
-inducer
+100 uM C4
Output (RPU)
Input (RPU)
Tmax
T0
R² (log scale) = 0.75
C4 (μM)
TF Input (RPU)
7.85 (Batch 3)
7.72 (Batch 2)
7.51 (Batch 1)
6.32 (Batch 3)
4.67 (Batch 2)
4.44 (Batch 1)
4.21 (Batch 3)
2.18 (Batch 2)
2.03 (Batch 1)
1.77 (Batch 3)
1.03 (Batch 2)
0.66 (Batch 1)
0.58 (Batch 3)
0.20 (Batch 2)
0.18 (Batch 1)
0.01 (Batch 3)
0.00 (Batch 2)
0.00 (Batch 1)
TF
PurR60-BjaR S107R 180 nls -VP16
PS.a0b1c0_purO
purO
polyA
TATA box
TSS
-inducer
+1 mM IV-HSL
Output (RPU)
Input (RPU)
Tmax
T0
R² (log scale) = 0.94
IV-HSL (μM)
TF Input (RPU)
8.92 (Batch 3)
8.10 (Batch 2)
8.07 (Batch 1)
5.19 (Batch 3)
4.68 (Batch 2)
4.66 (Batch 1)
2.32 (Batch 3)
2.25 (Batch 2)
2.12 (Batch 1)
0.76 (Batch 2)
0.64 (Batch 1)
0.23 (Batch 2)
0.17 (Batch 1)
0.00 (Batch 3)
0.00 (Batch 2)
0.00 (Batch 1)

**Supplementary Note 10: Validation of the Quantitative Model in the prediction of transcriptional repression without Parameter Adjustment.**

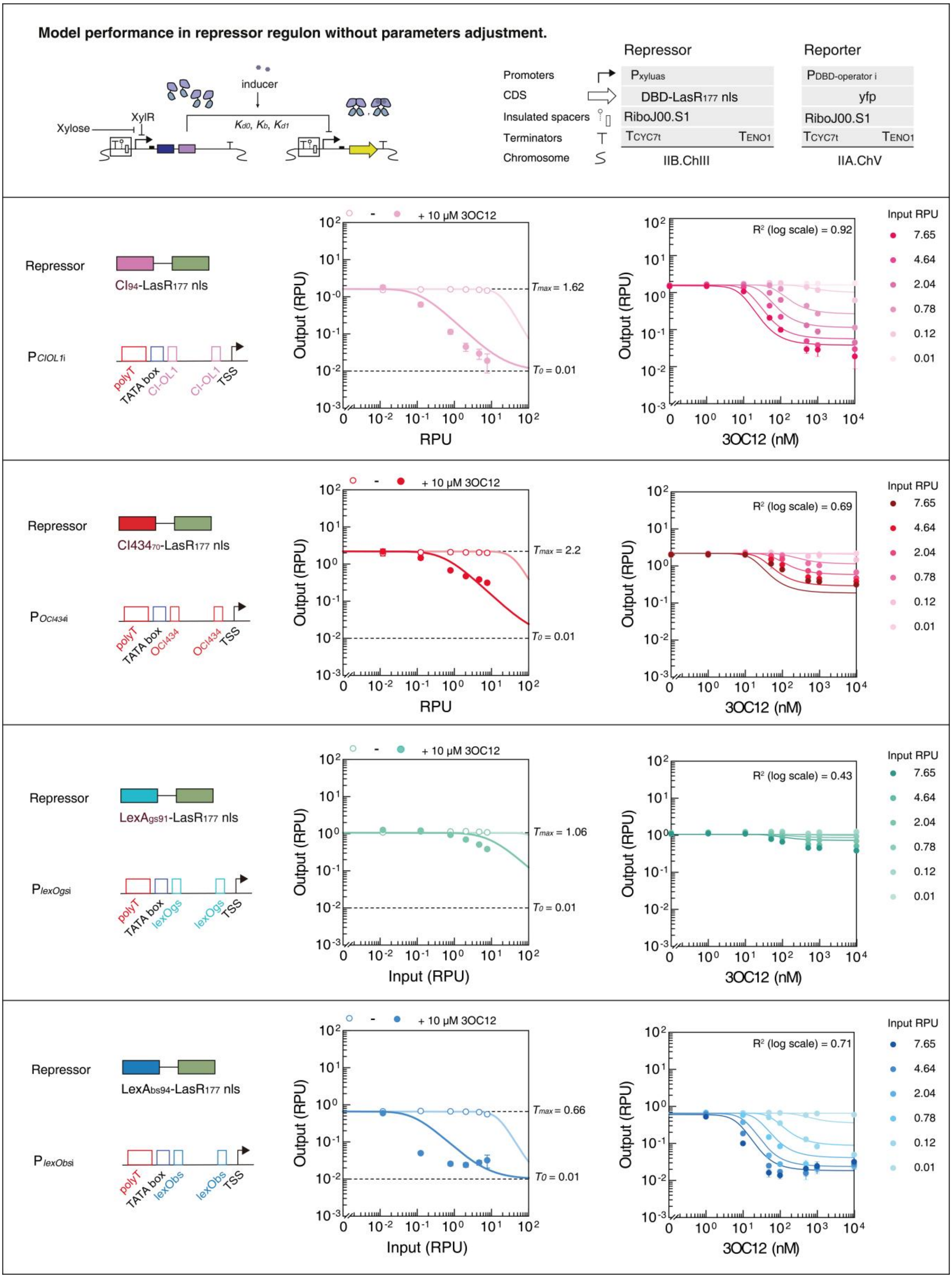

#### Supplementary Note 11: Quantitative test and model fitting for hybrid phenotype.

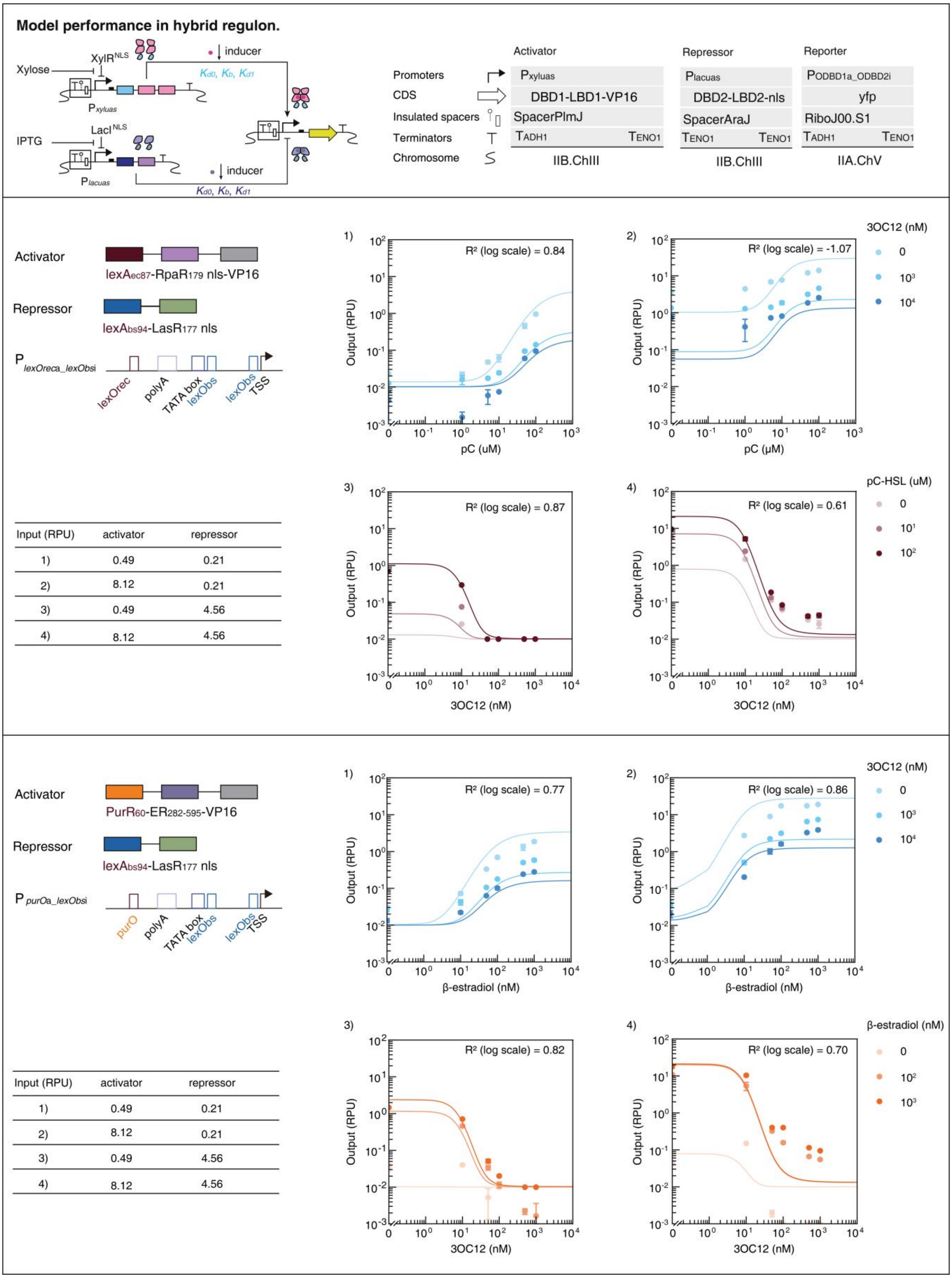

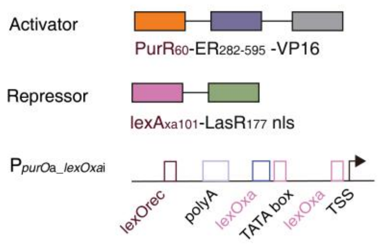

| Input (RPU) | activator | repressor |
|---|---|---|
| 1) | 0.49 | 0.21 |
| 2) | 8.12 | 0.21 |
| 3) | 0.49 | 4.56 |
| 4) | 8.12 | 4.56 |

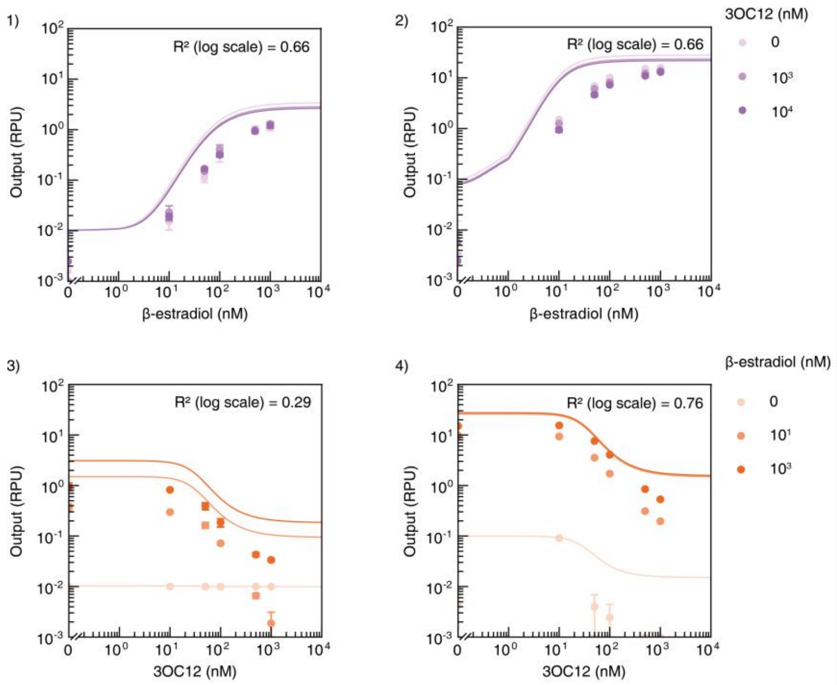

#### Supplementary Note 12: Specification of CIC-12 yeast chassis.

CIC-12 denotes the 12-sensor CIC yeast chassis described in Fig. 7 and Supplementary Fig. 14. The chassis consists of nine model-designed CIC-TFs together with TetR, LacI, and XlnR, assembled by iterative chromosomal integration in BY4741 and CEN.PK2-1C backgrounds. Full plasmid and strain information is provided in Supplementary Tables 7 and 8.

| TF: $TetR^{NLS}$ | Notes |
|---|---|
| $K_A$: n/a | TetR is the repressor of the tetracycline resistance element from *Escherichia coli*, its N-terminal region forms a helix-turn-helix structure and binds DNA. Binding of tetracycline to TetR reduces the repressor affinity for the tetracycline resistance gene (tetA) promoter operator sites. |
| $K_{d0}$: n/a | |
| $K_b$: n/a | |
| $K_{d1}$: n/a | |
| Inducer: Anhydrotetracycline HCl | Structures |
| Source:Sigma-Aldrich | 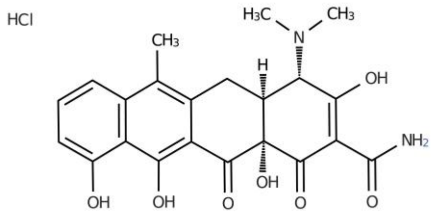 |
| Stock: 100 ng·ul$^{-1}$ | |
| Solvent: DMSO | |
| Max induction: 25 ng·ml$^{-1}$ | |
| Storage: -20°C | |
| $TetR^{NLS}$ in CEN.PK2-1C | |

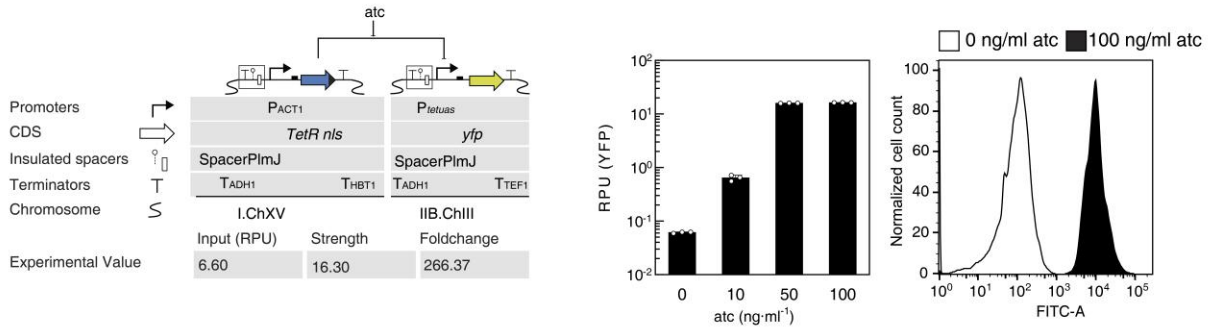

| TF: XlnR$^{NLS}$ | Notes |
|---|---|
| $K_A$: n/a | XlnR is an transcriptional activator of the xylanolytic system from an *Aspergillus nidulans*. |
| $K_{d0}$: n/a | |
| $K_b$: n/a | |
| $K_{d1}$: n/a | |
| Inducer: D-Xylose | Structures |
| Source: Sangon | 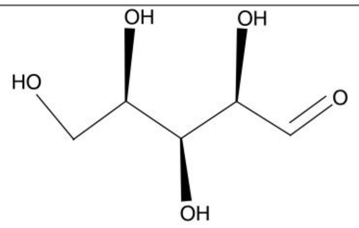 |
| Stock:1 mol·L$^{-1}$ | |
| Solvent: $H_2O$ | |
| Max induction: 10 mmol·L$^{-1}$ | |
| Storage: -20°C | |

XlnR$^{NLS}$ in CEN.PK2-1C

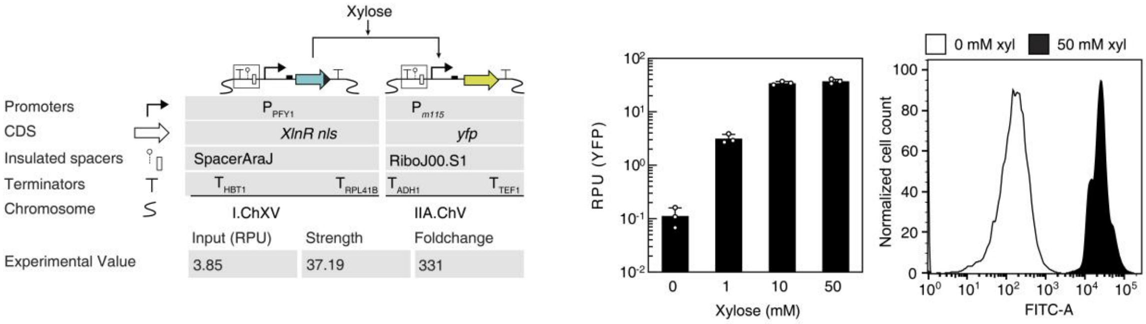

| TF: LacI$^{NLS}$ | Notes |
| --- | --- |
| $K_A$: n/a | LacI is a repressor of the lactose operon from *Escherichia coli*. |
| $K_{d0}$: n/a | |
| $K_b$: n/a | |
| $K_{d1}$: n/a | |
| Inducer: Isopropyl-β-D-thiogalactoside | Structures |
| Source: Sangon | 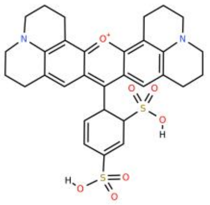 |
| Stock:1 mol·L$^{-1}$ | |
| Solvent: $H_2O$ | |
| Max induction: 10 mmol·L$^{-1}$ | |
| Storage: -20°C | |

LacI$^{NLS}$ in CEN.PK2-1C

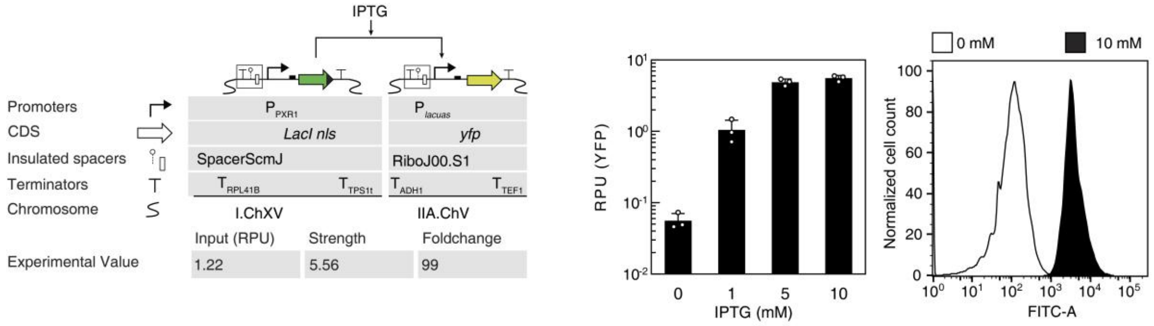

| TF: HKCI$_{84}$-LasR$_{177}$-VP16_opt1 | Notes |
|---|---|
| $K_A$: 5.07 | HKCI$_{84}$ is the DBD of HKCI repressor from Escherichia phage HK022 (Bacteriophage HK022), LasR$_{177}$ is the LBD of LasR, a LuxR family transcriptional regulator from *Pseudomonas aeruginosa*. |
| $K_{d0}$: $1.72 \times 10^{-4}$ | |
| $K_b$: $2.56 \times 10^{-1}$ | |
| $K_{d1}$: $2.25 \times 10^{2}$ | |
| Inducer:N-(3-Oxododecanoyl)-L-homoserine lactone | Structures |
| Source:Sigma-Aldrich | 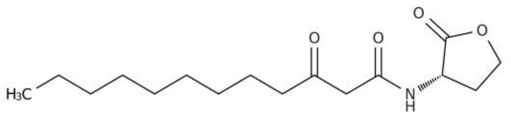 |
| Stock: 25 mM | |
| Solvent: DMSO | |
| Max induction: 1 uM | |
| Storage: -20°C | |

#### Sensor optimization strategy

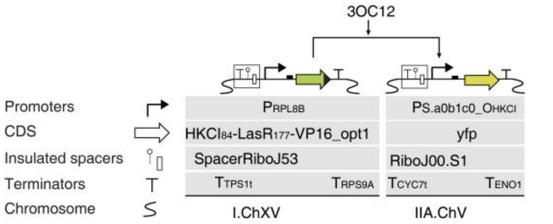

| | Input (RPU) | DBD | $\log_{10}(\Delta Exp)$ | $\log_{10}(FC)$ |
|---|---|---|---|---|
| Global Optimum | 0.46 | CI434$_{70}$ | 1.31 | 3.05 |
| Experimental Value | 0.96 | HKCI$_{84}$ | 1.36 | 2.49 |

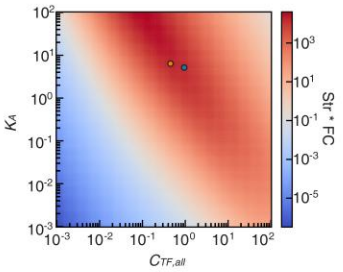

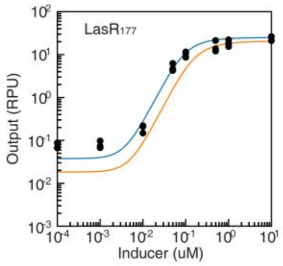

#### Standard promoter and Non-repetitive promoter

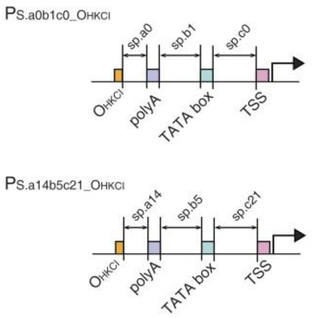

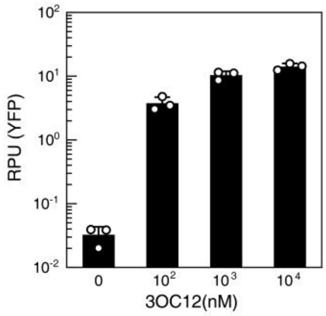

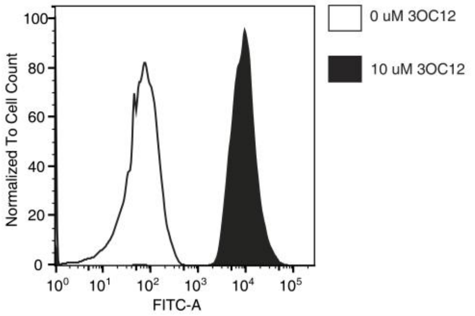

#### HKCI$_{84}$-LasR$_{177}$-VP16_opt1 in CEN.PK2-1C

| | Standard | Non-repetitive |
|---|---|---|
| leakiness | 0.09 | 0.04 |
| Max Output | 34.30 | 23.90 |
| Foldchange | 367 | 563 |

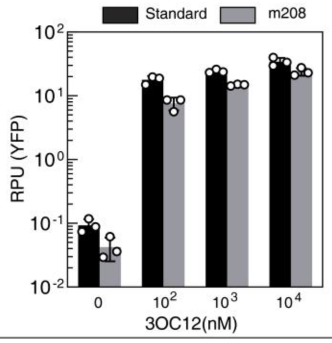

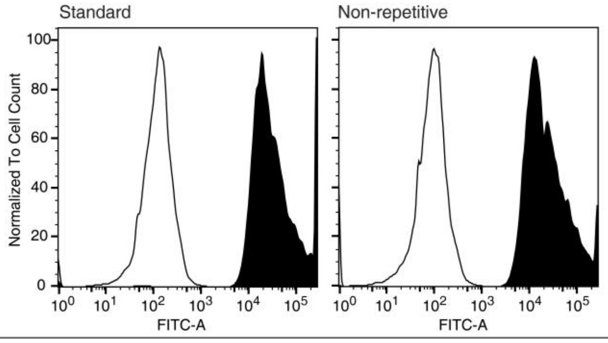

| TF: LexA$_{xa101}$-RpaR$_{179}$-VP16_opt2 | Notes |
|---|---|
| $K_A$: 1.45 | LexA$_{xa101}$ is the DBD of trancriptional repressor lexA from *X. axaonopodis* pv.ciitri str.306, RpaR$_{179}$ is a LBD of HTH-type quorum sensing-dependent transcriptional regulator RpaR from *Rhodopseudomonas palustris*. |
| $K_{d0}$: $2.69 \times 10^{-4}$ | |
| $K_b$: $7.08 \times 10^{-3}$ | |
| $K_{d1}$: $7.61 \times 10^{-1}$ | |
| Inducer: N-(p-Coumaroyl)-L-homoserine lactone | Structures |
| Source: Sigma-Aldrich | 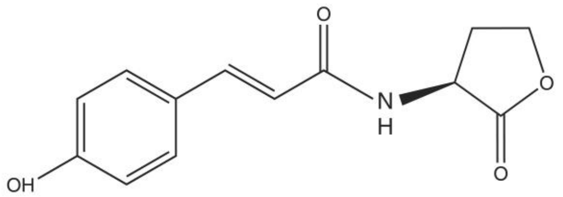 |
| Stock: 100 mM | |
| Solvent: DMSO | |
| Max induction: 50 uM | |
| Storage: -20°C | |

#### Sensor optimization strategy

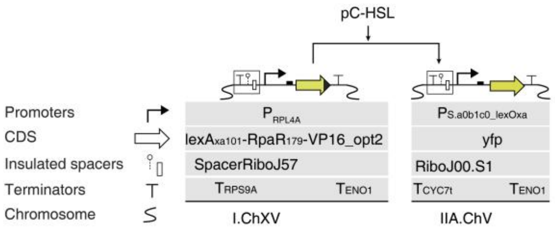

| | Input (RPU) | DBD | log10(ΔExp) | log10(FC) |
|---|---|---|---|---|
| Global Optimum | 0.77 | CI434$_{70}$ | 1.22 | 2.56 |
| Experimental Value | 2.08 | lexA$_{xa101}$ | 1.48 | 2.31 |

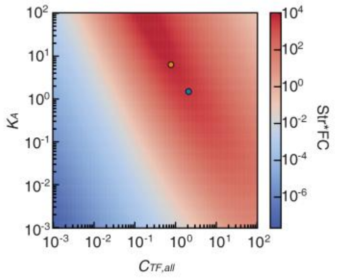

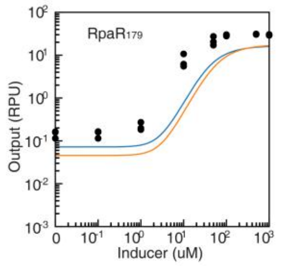

#### Standard promoter and Non-repetitive promoter

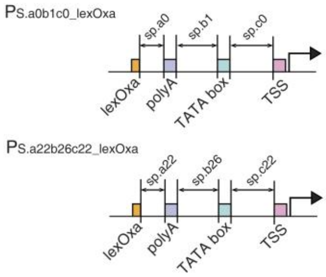

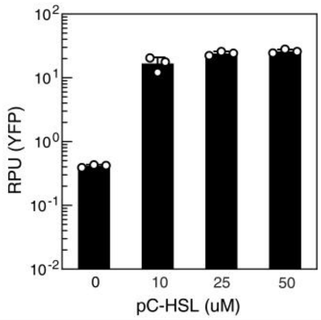

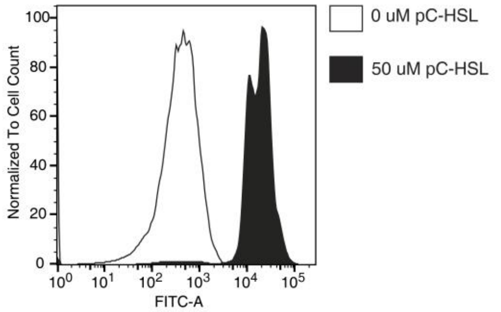

#### LexA$_{xa101}$-RpaR$_{179}$-VP16_opt2 in CEN.PK2-1C

| | Standard | Non-repetitive |
|---|---|---|
| leakiness | 0.06 | 0.15 |
| Max Output | 26.33 | 23.43 |
| Foldchange | 380 | 155 |

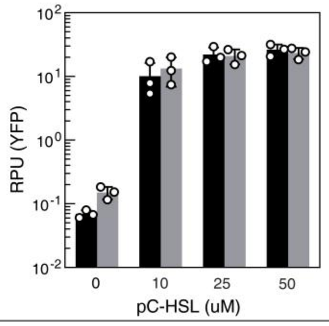

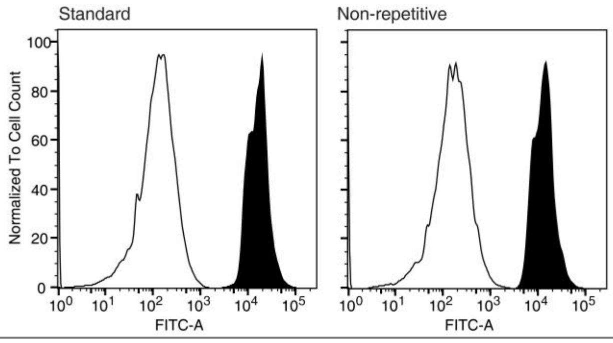

| TF:LexAec87-ER282-595-VP16_opt3 | Notes |
|---|---|
| $K_A$: 1.67 | LexAec87 is the DBD of trancriptional repressor lexA from *E. coli*, ER282-595 is the LBD of estrogen receptor from *Homo sapiens*. |
| $K_{d0}$: 3.40 × $10^{-5}$ | |
| $K_b$: 9.08 | |
| $K_{d1}$: 1.01 | |

| Inducer: beta-estradiol | Structures |
|---|---|
| Source: TCI | 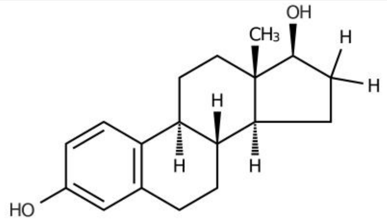 |
| Stock: 10 mM | |
| Solvent: Ethanol | |
| Max induction: 1 uM | |
| Storage: -20°C | |

#### Sensor optimization strategy

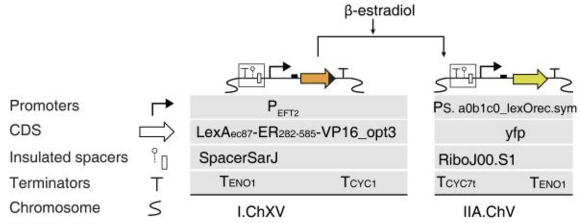

| | Input (RPU) | DBD | $log_{10}$(ΔExp) | $log_{10}$(FC) |
|---|---|---|---|---|
| Global Optimum | 1.24 | HKCl84 | 1.34 | 3.08 |
| Experimental Value | 1.78 | lexAec87 | 1.01 | 2.48 |

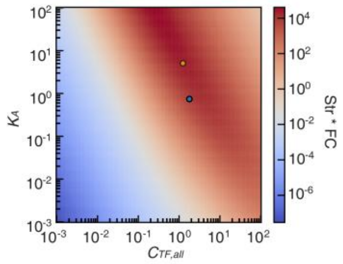

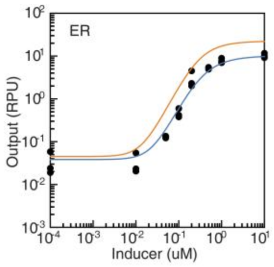

#### Standard promoter and Non-repetitive promoter

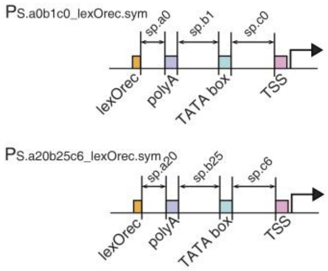

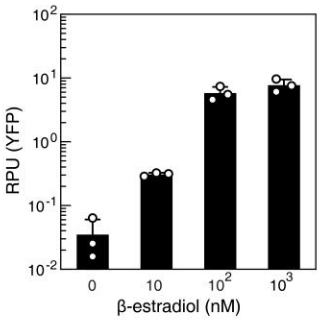

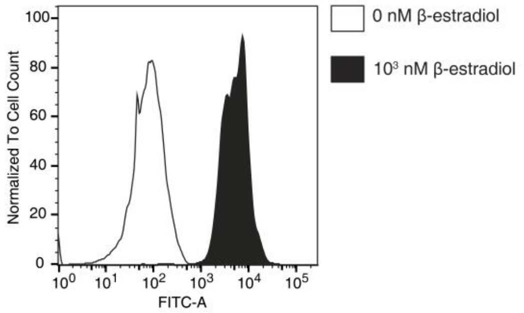

#### LexAec87-ER282-595-VP16_opt3 in CEN.PK2-1C

| | Standard | Non-repetitive |
|---|---|---|
| leakiness | 0.04 | 0.03 |
| Max Output | 15.24 | 12.72 |
| Foldchange | 423 | 473 |

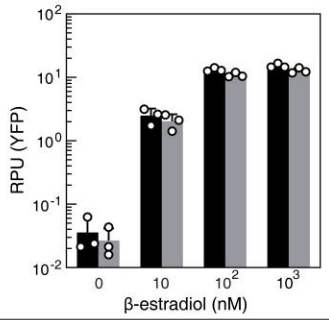

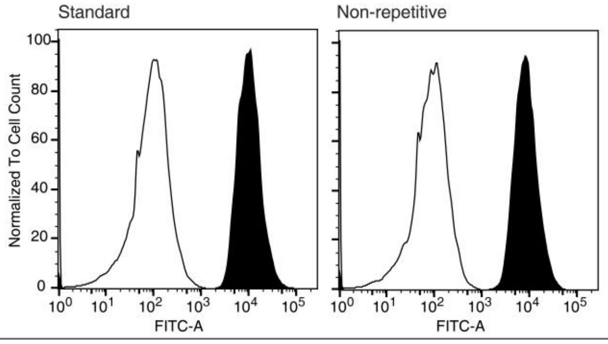

| TF:LexAmm115-DHBR282-595-VP16_opt4 | Notes |
| --- | --- |
| $K_A$: 5.37 | LexAmm115 is the DBD of LexA repressor from *Magnetococcus marinus* MC-1, DHBR282-595 is the LBD of estrogen receptor from *Homo sapiens* with 5 mutations (L346I, A350M, M388Q, G521S, Y526D). |
| $K_{d0}$: $2.74 \times 10^{-5}$ | |
| $K_b$: $5.42 \times 10^{-2}$ | |
| $K_{d1}$: 4.61 | |
| Inducer: 1,2-Bis(4-hydroxyphenyl) ethane-1,2-dione | Structures |
| Source: Mreda | 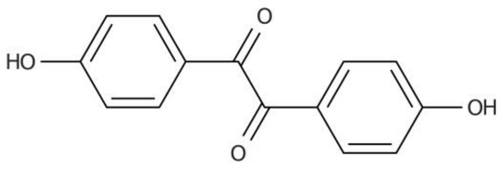 |
| Stock: 250 mM | |
| Solvent: Ethanol | |
| Max induction: 5 uM | |
| Storage: -20°C | |

#### Sensor optimization strategy

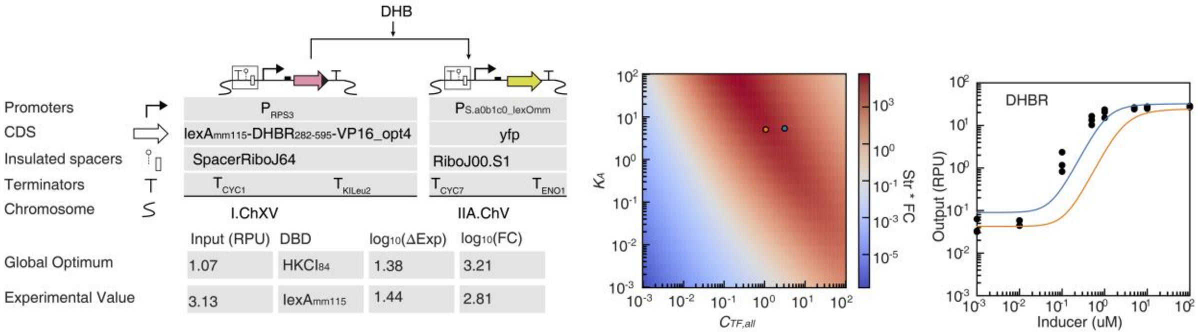

#### Standard promoter and Non-repetitive promoter

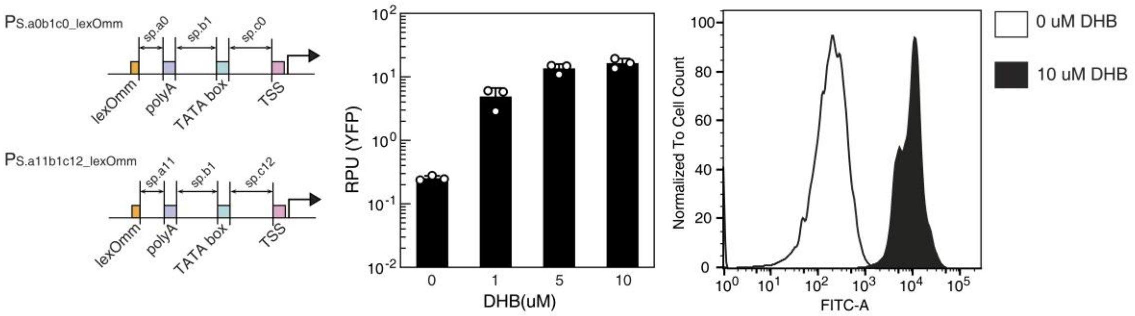

#### LexAmm115-DHBR282-595-VP16_opt4 in CEN.PK2-1C

| | Standard | Non-repetitive |
| --- | --- | --- |
| leakiness | 0.05 | 0.19 |
| Max Output | 30.94 | 22.50 |
| Foldchange | 587 | 116 |

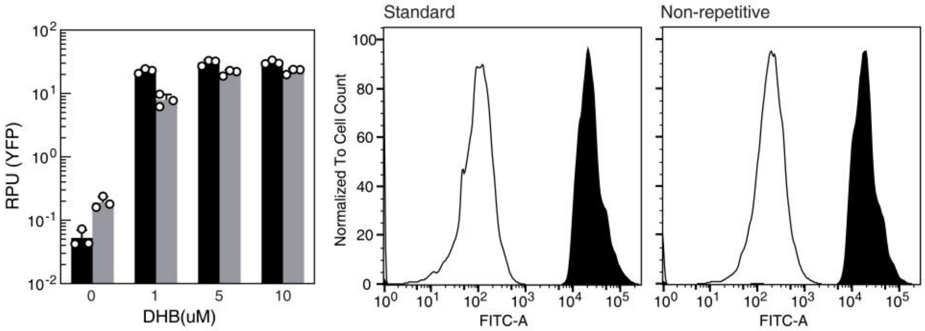

| TF: LexAgs91-acVHH-VP16_opt5 | Notes |
|---|---|
| $K_A$: 0.34 | LexAgs91 is the DBD of lexA repressor from *Geobacter sulfurreducens PCA,* acVHH is an antibody which can homodimerie upon caffeine binding. |
| $K_{d0}$: 1.04 × $10^{-2}$ | |
| $K_b$: 58.84 | |
| $K_{d1}$: 7.47 × $10^{-2}$ | |
| Inducer: Caffeine | Structures |
| Source: GLPBIO | 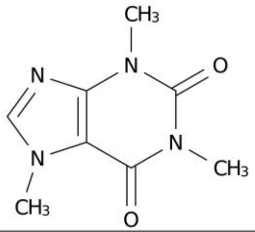 |
| Stock: 10 mM | |
| Solvent: DMSO | |
| Max induction: 100 nM | |
| Storage: -20°C | |

#### Sensor optimization strategy

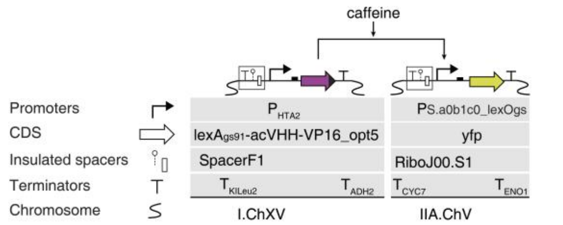

| | Input (RPU) | DBD | log10(ΔExp) | log10(FC) |
|---|---|---|---|---|
| Global Optimum | 0.30 | lexAbs94 | 1.31 | 1.20 |
| Experimental Value | 1.14 | lexAgs91 | 0.62 | 1.68 |

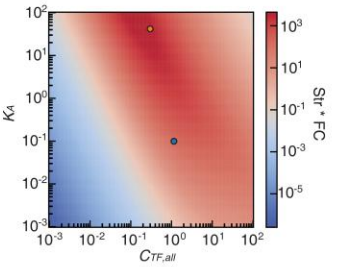

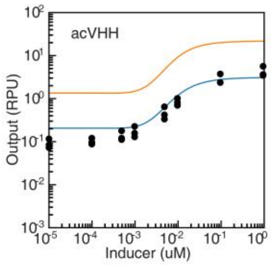

#### Standard promoter and Non-repetitive promoter

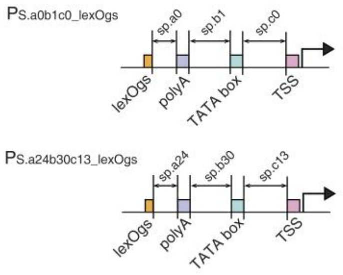

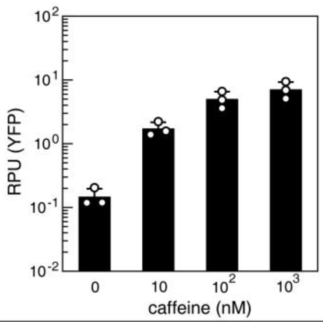

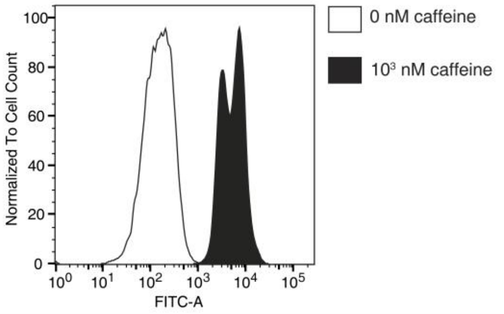

#### LexAgs91-acVHH-VP16_opt5 in CEN.PK2-1C

| | Standard | Non-repetitive |
|---|---|---|
| leakiness | 0.09 | 0.15 |
| Max Output | 4.14 | 6.44 |
| Foldchange | 46 | 43 |

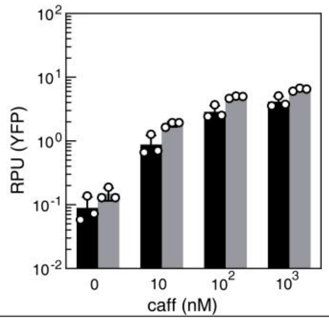

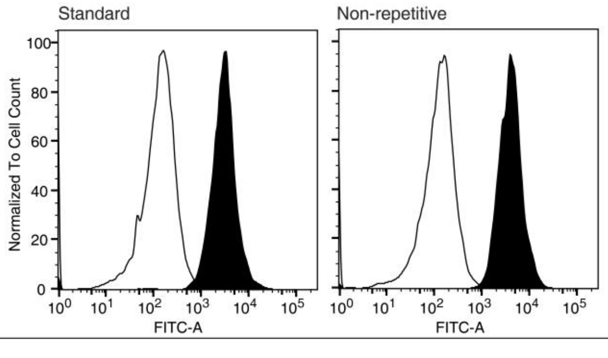

| TF: CI434$_{70}$-CinR$_{179}$-VP16_opt6 | Notes |
|---|---|
| $K_A$: 6.22 | CI434$_{70}$ is the DBD of repressor CI from *E. coli* phage CI434, CinR$_{179}$ is the LBD of a LuxR family transcriptional regulator CinR from *Rhizobium leguminosarum*. |
| $K_{d0}$: $3.91 \times 10^{-3}$ | |
| $K_b$: 56.84 | |
| $K_{d1}$: 0.16 | |
| Inducer: 3-hydroxytetradecanoyl-homoserine lactone | Structures |
| Source: Sigma-Aldrich | 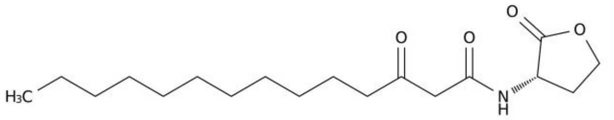 |
| Stock: 25 mM | |
| Solvent: DMSO | |
| Max induction: 100 nM | |
| Storage: -20°C | |

#### Sensor optimization strategy

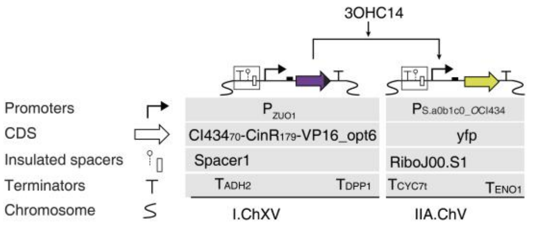

| | Input (RPU) | DBD | $log_{10}(\Delta Exp)$ | $log_{10}(FC)$ |
|---|---|---|---|---|
| Global Optimum | 0.41 | lexA$_{bs94}$ | 1.20 | 1.23 |
| Experimental Value | 0.98 | CI434$_{70}$ | 0.72 | 1.90 |

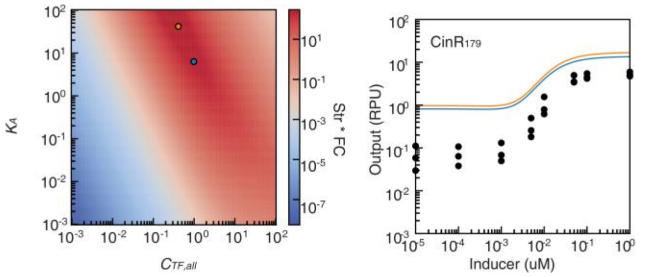

#### Standard promoter and Non-repetitive promoter

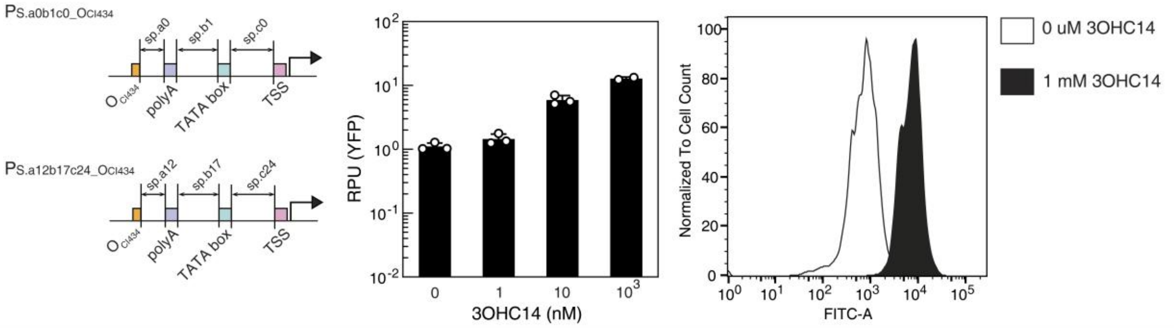

#### CI434$_{70}$-CinR$_{179}$-VP16_opt6 in CEN.PK2-1C

| | Standard | Non-repetitive |
|---|---|---|
| leakiness | 0.03 | 0.35 |
| Max Output | 5.33 | 11.96 |
| Foldchange | 158 | 34 |

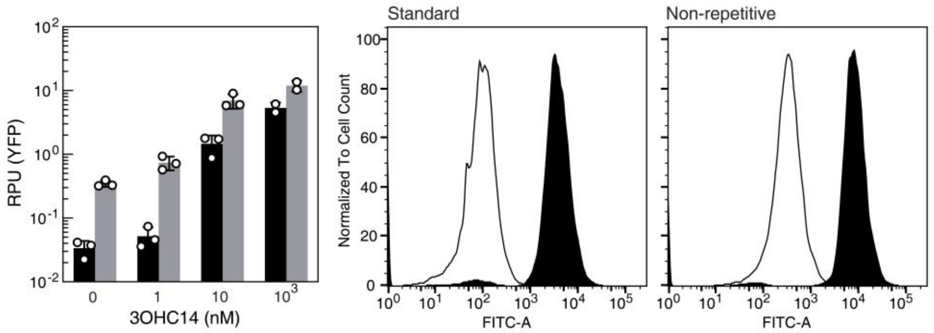

| TF:CI94-PR-VP16_opt7 | Notes |
|---|---|
| $K_A$: 30.09 | CI94 is the DBD of CI repressor from *Enterobacteria* phage lamda, PR is the LBD of progesterone receptor from *Homo sapiens*. |
| $K_{d0}$: $5.93 \times 10^{-5}$ | |
| $K_b$: 0.15 | |
| $K_{d1}$: $5.28 \times 10^{-2}$ | |
| Inducer: RU486 | Structures |
| Source: Selleck | 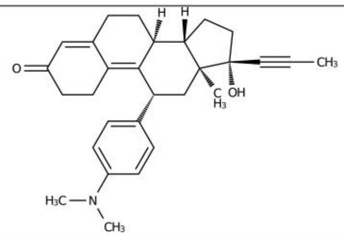 |
| Stock: 100 mM | |
| Solvent: DMSO | |
| Max induction: 10 uM | |
| Storage: -20°C | |

#### Sensor optimization strategy

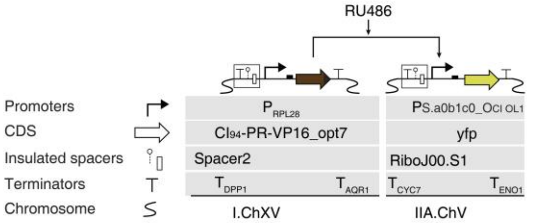

| | Input (RPU) | DBD | log10(ΔExp) | log10(FC) |
|---|---|---|---|---|
| Global Optimum | 2.20 | CI43470 | 1.26 | 2.39 |
| Experimental Value | 0.67 | CI94 (OL1) | 1.21 | 2.31 |

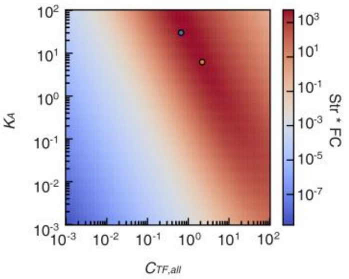

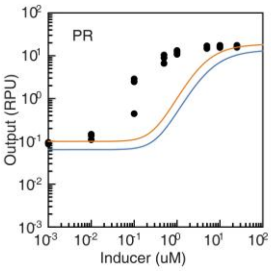

#### Standard promoter and Non-repetitive promoter

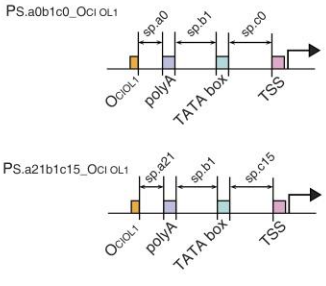

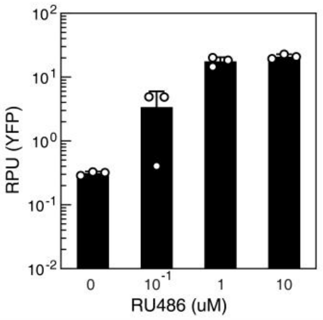

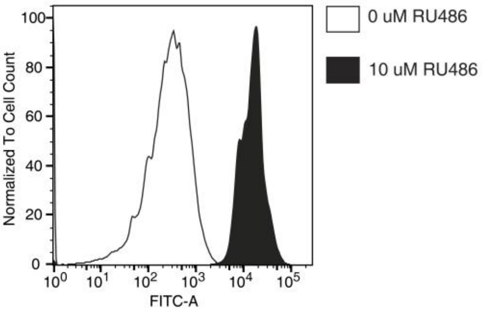

#### CI94-PR-VP16_opt7 in CEN.PK2-1C

| | Standard | Non-repetitive |
|---|---|---|
| leakiness | 0.07 | 0.20 |
| Max Output | 18.67 | 21.69 |
| Foldchange | 257 | 107 |

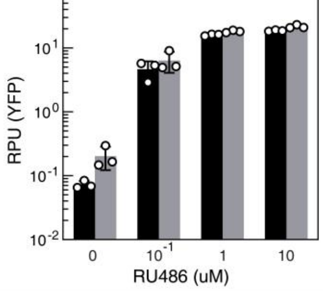

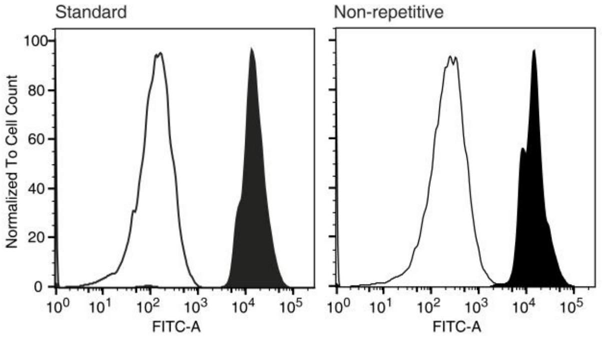

| TF: PurR60-BjaR180$^{S107R}$-VP16_opt8 | Notes |
|---|---|
| $K_A$: 1.13 | PurR60 is the DBD of PurR from *E. coli str.*K-12 substr. MG1655, BjaR180$^{S107R}$ is the LBD of BjaR, a LuxR family transcriptional regulator BjaR from Bradyrhizobium. |
| $K_{d0}$: 4.40 × 10$^{-4}$ | |
| $K_b$: 7.14 × 10$^{-3}$ | |
| $K_{d1}$: 0.17 | |
| Inducer: N-isovaleryl-L-homoserine lactone | Structures |
| Source: Chemically synthesized | 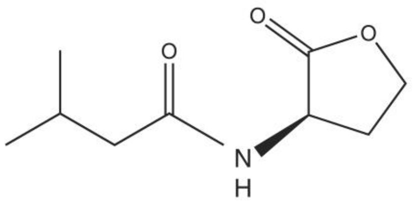 |
| Stock: 100 mM | |
| Solvent: DMSO | |
| Max induction: 400 uM | |
| Storage: -20°C | |

#### Sensor optimization strategy

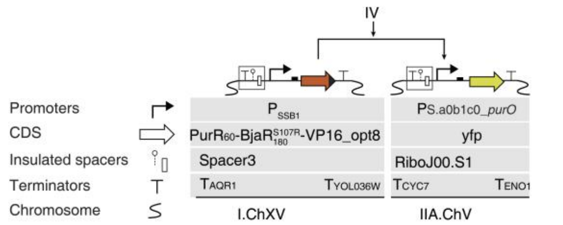

| | Input (RPU) | DBD | ΔExp | fold-change |
|---|---|---|---|---|
| Global Optimum | 0.95 | lexAbs94 | 1.24 | 1.47 |
| Experimental Value | 2.06 | PurR60 | 1.22 | 1.99 |

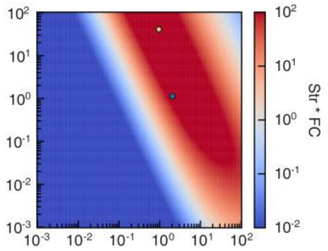

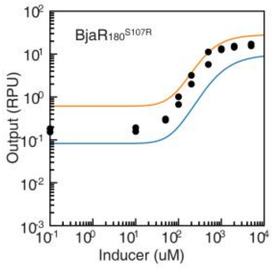

#### Standard promoter and Non-repetitive promoter

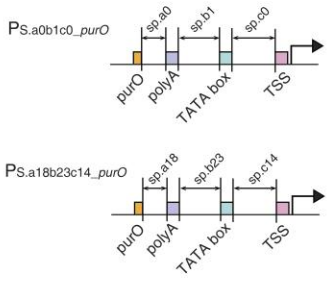

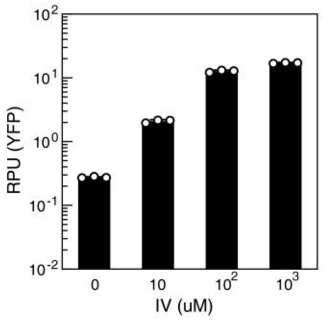

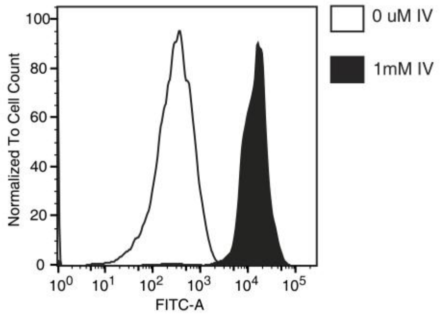

#### PurR60-BjaR180$^{S107R}$-VP16_opt8 in CEN.PK2-1C

| | Standard | Non-repetitive |
|---|---|---|
| leakiness | 0.06 | 0.12 |
| Max Output | 21.95 | 19.21 |
| Foldchange | 340 | 160 |

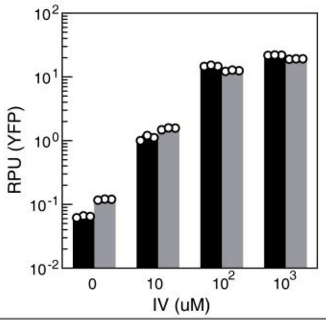

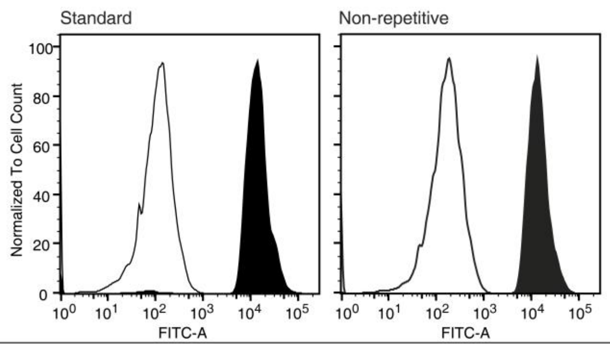

| TF: LexAbs94-LuxR183$^{S116Y}$-VP16_opt9 | Notes |
|---|---|
| $K_A$: 40.98 | lexAbs94 is the DBD of lexA repressor from *Bacillus subtilis*, LuxR183$^{S116Y}$ is the LBD of trancriptional regulator LuxR from *Aliivibrio fischeri*. |
| $K_{d0}$: 9.31 × $10^{-8}$ | |
| $K_b$: 9.05 × $10^{-3}$ | |
| $K_{d1}$: 2.69 × $10^{-5}$ | |
| Inducer: N-(3-Oxohexanoyl)-L-homoserine lactone | Structures |
| Source:Sigma-Aldrich | 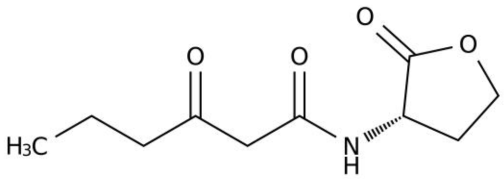 |
| Stock: 100 mM | |
| Solvent: DMSO | |
| Max induction: 1 mM | |
| Storage: -20°C | |

#### Sensor optimization strategy

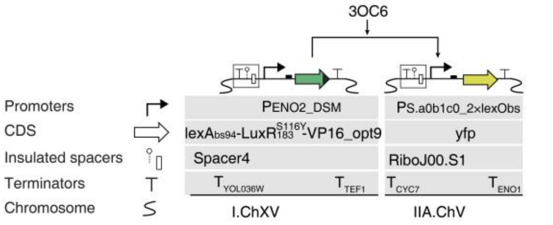

| | Input (RPU) | DBD | ΔExp | fold-change |
|---|---|---|---|---|
| Global Optimum | 10 | lexAbs94 | -0.07 | 1.27 |
| Experimental Value | 11.57 | lexAbs94 | 0.97 | 2.38 |

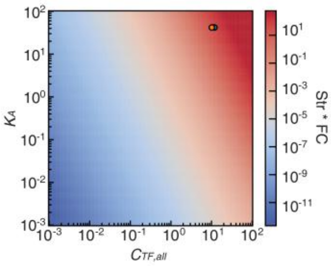

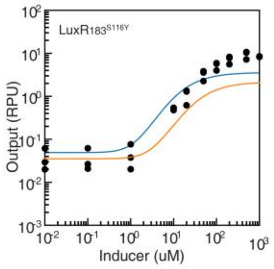

#### Standard promoter and Non-repetitive promoter

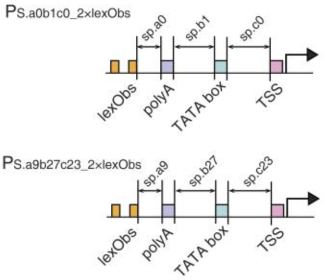

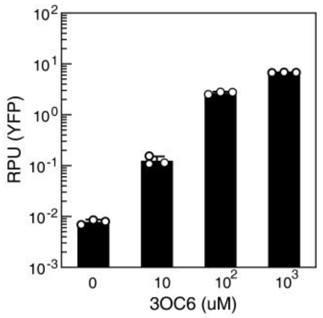

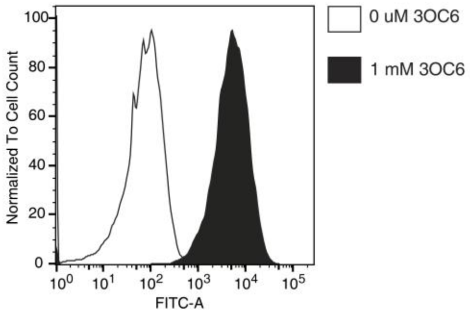

#### LexAbs94-LuxR183$^{S116Y}$-VP16_opt9 in CEN.PK2-1C

| | Standard | Non-repetitive |
|---|---|---|
| leakiness | 0.02 | 0.02 |
| Max Output | 8.96 | 5.52 |
| Foldchange | 403 | 257 |

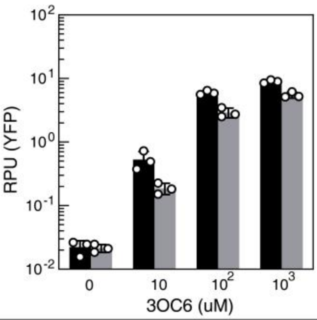

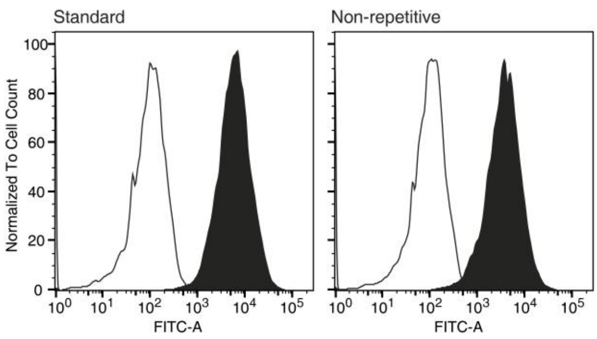

#### Supplementary Tables

##### Supplementary Table 1: Comparison of Type I-III TFs

| Feature | Type I (Allosteric) | Type II (CID) | Type III (CIC) |
|---|---|---|---|
| Prototype | rtTA, LacI | dCas9-FKBP/FRB | CIC-TF / ligand-induced |
| Mechanism | Intramolecular Allostery | Intermolecular | Ligand-induced cooperative |
| DNA binding | Ligand-dependent | Constitutive | Ligand-induced cooperative |
| Cooperativity | >1 (Cooperative) | ≈1 (Non-cooperative) | >1 (Cooperative) |
| Basal State | Bound or Unbound | Bound to DNA | Low DNA-binding-competent |
| Modularity | Low (Structurally coupled) | High | High |
| Toxicity Risk | Low | High (Roadblock) | Lower under tested conditions |

**Supplementary Table 2: Fitting Parameters of DNA binding domain.**

| **DBD** | **operator** | ***Sequence*** | $K_A$ | $T_{0,variant}$ |
|---|---|---|---|---|
| LexA*ec*$_{87}$ | lexOec | TGCTGTATATACTCACAGCA | | |
| | lexO*rec.sym* | TACTGTATGATCATACAGTA | 0.74 | 0.016 |
| | lexO*gc* | TACTGTATGCGCATACAGTA | 19.36 | |
| | lexO*ta* | TACTGTATATATATACAGTA | 9.39 | |
| LexA*bs*$_{94}$ | lexO*bs* | AGAACATATGTTCG | 40.98 | 0.035 |
| LexA*fs*$_{104}$ | lexO*fs* | CTGCACAAAGGTGCAC | 0.10 | 0.111 |
| LexA*gs*$_{91}$ | lexO*gs* | GGTTGACACATGTCAACC | 0.34 | 0.045 |
| LexA*mm*$_{115}$ | lexO*mm* | ACCTAATATTAATAAGGT | 5.37 | 0.010 |
| LexA*xa*$_{101}$ | lexO*xa* | TTAGTAGTAATACTACTAA | 1.45 | 0.012 |
| RecA*pact*$_{104}$ | $O_{RecApact}$ | TACTAATTAATTTAGT | 1.04 | 0.017 |
| DeoR$_{92}$ | $O_{DeoR}$ | TGTTAGAATTCTAACA | 0.61 | 0.024 |
| HKCI$_{84}$ | $O_{HKCI}$ | TGAACCATAAGTTCA | 5.07 | 0.0090 |
| CI$_{94}$ | $O_{CI\ OL1}$ | TACCACTGGCGGTGATA | 30.09 | 0.136 |
| | $O_{CI\ OR1}$ | TACCTCTGGCGGTGATA | 10.37 | 0.136 |
| | $O_{CI\ Osym}$ | TATCACCGGCGGTGATA | 10.53 | 0.272 |
| CI434$_{70}$ | $O_{CI434}$ | TACAAGAAAGTTTGTT | 6.22 | 0.010 |
| PurR$_{60}$ | *purO* | ACGCAAACGTTTGCGT | 1.13 | 0.007 |

**Supplementary Table 3: Fitting Parameters of Ligand binding domain in yeast.**

| **LBD** | **Inducer** | $K_{d0}$ | $K_b$ | $K_{d1}$ |
|---|---|---|---|---|
| acVHH | Caffeine | $1.05\times10^{-2}$ | 58.84 | $7.47\times10^{-1}$ |
| $RpaR_{179}$ | pC-HSL | $2.69\times10^{-4}$ | $7.08\times10^{-3}$ | $7.61\times10^{-1}$ |
| $BjaR_{180}$ | IV-HSL | $1.53\times10^{-4}$ | $2.74\times10^{-3}$ | $5.09\times10^{-2}$ |
| $LasR_{177}$ | 3OC12-HSL | $1.72\times10^{-4}$ | $2.56\times10^{-1}$ | 225.08 |
| $CinR_{179}$ | 3OHC14-HSL | $3.91\times10^{-3}$ | 56.84 | $1.62\times10^{-1}$ |
| $TraR_{174}$ | 3OC8-HSL | $1.10\times10^{-4}$ | $1.82\times10^{-1}$ | $6.04\times10^{-2}$ |
| $CarRecc_{169}$ | 3OC6-HSL | $2.46\times10^{-2}$ | $8.49\times10^{-6}$ | $5.38\times10^{8}$ |
| $SmaR_{179}$ | C4-HSL | $1.00\times10^{-2}$ | $4.59\times10^{-1}$ | $2.08\times10^{-1}$ |
| | C8-HSL | $1.00\times10^{-2}$ | 0.06 | 0.04 |
| | IV-HSL | $1.00\times10^{-2}$ | 3.40 | 0.05 |
| | 3OC12-HSL | $1.00\times10^{-2}$ | 0.91 | 0.10 |
| $ER_{282\text{-}595}$ | β-estradiol | $3.40\times10^{-5}$ | 9.08 | 1.01 |
| $DHBR_{282\text{-}595}$ | DHB | $2.74\times10^{-5}$ | $5.42\times10^{-2}$ | 4.61 |
| PR | RU486 | $5.93\times10^{-5}$ | 0.15 | $5.28\times10^{-2}$ |
| $MR_{669\text{-}984}$ | aldosterone | $8.48\times10^{-6}$ | $5.99\times10^{-2}$ | $3.22\times10^{-3}$ |
| $GR_{487\text{-}777}$ | dexamethasone | $3.30\times10^{-6}$ | $5.13\times10^{-3}$ | $1.07\times10^{-4}$ |
| $PluR_{17\text{-}162}$ | PPYA | $1.05\times10^{-4}$ | $4.55\times10^{-1}$ | $1.76\times10^{-4}$ |
| $PauR_{17\text{-}160}$ | Tapinarof | $1.22\times10^{-4}$ | 1.75 | $4.27\times10^{-4}$ |
| $VqmA_{91\text{-}187}$ | 3,5-Dimethylpyrazin-2-ol | $7.76\times10^{-3}$ | 32.64 | $2.05\times10^{-2}$ |
| $BjaR_{180}^{S107R}$ | IV-HSL | $4.40\times10^{-4}$ | $7.14\times10^{-3}$ | $1.75\times10^{-1}$ |
| $LuxR_{183}^{S116Y}$ | 3OC6-HSL | $9.31\times10^{-8}$ | $9.05\times10^{-3}$ | $2.69\times10^{-5}$ |
| $CepR_{176}$ | C8-HSL | $5.86\times10^{-2}$ | $2.55\times10^{-2}$ | $4.67\times10^{-1}$ |

**Supplementary Table 4: Full-curve fitted apparent LBD parameters in mammalian cells.**

| LBD | Inducer | $K_{d0}$ | $K_b$ | $K_{d1}$ |
|---|---|---|---|---|
| $RpaR_{179}$ | pC-HSL | $9.97\times10^{-7}$ | 0.0371 | 0.171 |
| $BjaR_{180}$ | IV-HSL | $1.04\times10^{-7}$ | $7.87\times10^{-5}$ | 0.00737 |
| $TraR_{174}$ | 3OC8-HSL | $6.52\times10^{-7}$ | 0.00342 | 0.00112 |
| $CinR_{179}$ | 3OHC14-HSL | $3.21\times10^{-5}$ | 0.354 | 0.00305 |
| $CarRecc_{169}$ | 3OC8-HSL | 0.0394 | 0.0136 | 2.16 |
| $BjaR_{180}^{S107R}$ | IV-HSL | $6.79\times10^{-7}$ | $3.56\times10^{-6}$ | 1.87 |
| $LasR_{177}$ | 3OC12-HSL | $3.28\times10^{-7}$ | 0.0089 | 0.019 |
| acVHH | Caffeine | 0.226 | 0.043 | $6.79\times10^{5}$ |
| $ER_{282-595}$ | β-estradiol | $1.04\times10^{-5}$ | 0.0018 | $4.65\times10^{10}$ |
| $DHBR_{282-595}$ | DHB | $3.50\times10^{-13}$ | 0.171 | 0.0278 |
| PR | RU486 | $2.82\times10^{-13}$ | 341 | 1.31 |
| $MR_{669-984}$ | Aldosterone | $3.99\times10^{-5}$ | 183 | 0.00881 |
| $GR_{487-777}$ | Dexamethasone | $1.80\times10^{-12}$ | 0.0136 | 0.00893 |

Units: $K_b$, $\mu M^{-1}$; $K_{d0}$ and $K_{d1}$, $RPU^{-1}$.

**Supplementary Table 5: Full-curve fitted apparent DBD parameters in mammalian cells.**

| DBD | $K_A$ | $T_{0,variant}$ |
|---|---|---|
| $CI434_{70}$ | 4.02 | 0.0309 |
| $lexA_{ec87}$ | 9.15[a] | 0.0504 |
| $lexA_{xa101}$ | 0.145 | 0.0607 |
| $PurR_{60}$ | 367 | 0.146 |
| $lexA_{mm115}$ | 318 | 0.0757 |
| $lexA_{fs104}$ | 19.3 | 0.0207 |
| $CI_{94}$ | $3.65\times10^3$ | 0.0531 |
| $HKCI_{84}$ | 18.2 | 0.11 |
| $lexA_{gs91}$ | 3.44 | 0.0245 |
| $RecApact_{104}$ | 4.35 | 0.203 |
| $DeoR_{92}$ | 0.0267 | 0.0262 |

Units: $K_A$, $RPU^{-1}$; $T_{0,variant}$, RPU. The same canonical DBD shares $K_A$ and $T_{0,variant}$ across sensors. [a] $K_A$ for $lexA_{ec87}$ was fixed at 9.15; all other $K_A$ values and all $T_{0,variant}$ values were jointly optimized. $T_{max}$ = 13.61 RPU and operator number = 7 were fixed globally.

**Supplementary Table 6: Sequences of backbone plasmids for TF and reporter integration.**

| Plasmid | sequence | Restriction enzyme |
|---|---|---|
| XJH01 | ggtctcggctaaattcgagtgaaacacaggaagatcagaaaatcctcatttcatccatattaacaataatttcaaatgtttatttgcat<br>tatttgaaactaggcaagacaagcaacgaaacgtttttgaaaattttgagtattttcaataaatttgtagaggactcagatattgaaaa<br>aaagctacagcaattaatacttgataagaagagtattgagaagggcaacggttcatcatctcatggatctgcacatgaacaaaca<br>ccagagtcaaacgacgttgaaattgaggctactgcgccaattgatgacaatacagacgatgataacaaaccgaagttatctgatg<br>tagaaaaggattaaagatgctaagagatagtgatgatatttcataaataatgtaattctatatatgttaattaccttttttgcgaggcata<br>tttatggtgaaggataagttttgaccatcaaagaaggttaatgtggctgtggtttcagggtccataaagcccacatggataacatta<br>cgcttgctatgtcgtcggaggagatatttattacttttattattctagtttttttacagttatttattaattaattatttttatatgcatgcgaata<br>aaaagtctatatttaagttcttttatttattaatacattttcctctacgagctgtcaccggatgtgctttccggtctgatgagtccgtgag<br>gacgaaacagcctctacaaataattttgtttaagagcaggttgttcatggccgtgcgtatgatgtgggggggctcgggcgttgaaa<br>ccggggttcggagcgccaggggggttttaggtcttcggcttactaaaagccagataacagtatgcatatttgcgcgctgatttttgc<br>ggtataagaatatatactgatatgtatacccgaagtatgtcaaaaagaggtatgctatgaagcagcgtattacagtgacagttgac<br>agcgacagctatcagttgctcaaggcatatatgatgtcaatatctccggtctggtaagcacaaccatgcagaatgaagcccgtcg<br>tctgcgtgccgaacgctggaaagcggaaaatcaggaagggatggctgaggtcgcccggtttattgaaatgaacggctcttttgc<br>tgacgagaacaggggctggtgaaatgcagtttaaggtttacacctataaaagagagagccgttatcgtctgtttgtggatgtacag<br>agtgatattattgacacgcccgggcgacggatggtgatccccctggccagtgcacgtctgctgtcagataaagtctcccgtgaa<br>ctttacccggtggtgcatatcggggatgaaagctggcgcatgatgaccaccgatatggccagtgtgccggtttccgttatcggg<br>gaagaagtggctgatctcagccaccgcgaaaatgacatcaaaaacgccattaacctgatgttctggggaatataagaagaccta<br>atgtctaaaggtgaagaattattcactggtgttgtcccaattttggttgaattagatggtgatgttaatggtcacaaattttctgtctccg<br>gtgaaggtgaaggtgatgctacttacggtaaattgaccttaaaatttatttgtactactggtaaattgccagttccatggccaacctta<br>gtcactactttaggttatggtttgatgtgttttgctagatacccagatcatatgaaacaacatgactttttcaagtctgccatgccagaa<br>ggttatgttcaagaaagaactatttttttcaaagatgacggtaactacaagaccagagctgaagtcaagtttgaaggtgatacctta<br>gttaatagaatcgaattaaaaggtattgattttaaagaagatggtaacattttaggtcacaaattggaatacaactataactctcacaa<br>tgtttacatcatggctgacaaacaaaagaatggtatcaaagttaacttcaaaattagacacaacattgaagatggttctgttcaatta<br>gctgaccattatcaacaaaatactccaattggtgatggtccagtcttgttaccagacaaccattacttatcctatcaatctagattatc<br>caaagatccaaacgaaaagagggatcacatggtcttgttagaatttgttactgctgctggtattacccatggtatggatgaattgta<br>caaataaaagcttttgattaagccttctagtccaaaaaacacgttttttttgtcatttatttcattttcttagaatagtttagtttattcattttat<br>agtcacgaatgttttatgattctatatagggttgcaaacaagcatttttcattttatgttaaaacaatttcaggtttaccttttattctgcttg<br>tggtgacgcgtgtatccgcccgctcttttggtcacccatgtatttaattgcataaataattcttaaaagtggagctagtctatttctattt<br>acatacctctcatttctcatttcctcctgagaaatctgctcgtcagtggtgctcacactgacgaatcatgtacagatcataccgatga<br>ctgcctggcgactcacaactaagcaagacagccggaaccagcgccggcgaacaccactgcatatatggcatatcacaacagt<br>ccacgtctcaagcagttacagagatgttacgaaccactagtgcactgcagtacaagcttgccttgtccccgccgggtcacccgg<br>ccagcgacatggaggcccagaataccctccttgacagtcttgacgtgcgcagctcaggggcatgatgtgactgtcgcccgtac<br>atttagcccatacatccccatgtataatcatttgcatccatacattttgatggccgcacggcgcgaagcaaaaattacggctcctcg<br>ctgcagacctgcgagcagggaaacgctcccctcacagacgcgttgaattgtccccacgccgcgcccctgtagagaaatataaa<br>aggttaggatttgccactgaggttcttctttcatatacttccttttaaaatcttgctaggatacagttctcacatcacatccgaacataaa<br>caacaatgacagtcaacactaagacctatagtgagagagcagaaactcatgcctcaccagtagcacaacgattatttcgattaat<br>ggaactgaagaaaaccaatttatgtgcatcaattgatgttgataccactaaggaattccttgaattaattgataaattgggtccttatg<br>tatgcttaatcaagacacatattgatataatcaatgatttttcctatgaatccactattgaaccattattagaactttcacgtaaacatca<br>atttatgatttttgaagatagaaaatttgctgatattggtaataccgtgaagaaacaatatattggtggagtttataaaattagtagttg | BpiI |

| | | |
|---|---|---|
| | ggcagatattactaatgctcatggtgtcactgggaatggagtagttgaaggattaaaacagggagctaaagaaaccaccaccaa<br>ccaagagccaagagggttattgatgttagctgaattatcatcagtgggatcattagcatatggagaatattctcaaaaaactgttga<br>aattgctaaatccgataaggaatttgttattggatttattgcccaacgtgatatgggtggacaagaagaaggatttgattggcttatta<br>tgacacctggagttggattagatgataaaggtgatggattaggacaacaatatagaactgttgatgaagttgttagcactggaact<br>gatattatcattgttggtagaggattgtttggtaaaggaagagatccagatattgaaggtaaaaggtatagagatgctggttggaat<br>gcttatttgaaaaagactggccaattataaacagtactgacaataaaaagattcttgttttcaagaacttgtcatttgtatagtttttttat<br>attgtagttgttctattttaatcaaatgttagcgtgatttatatttttttcgcctcgacatcatctgcccagatgcgaagttaagtgcgca<br>gaaagtaatatcatgcgtcaatcgtatgtgaatgctggtcgctatactgctgtcgattcgatactaacgccgccatccagtgtcgag<br>agtagagcacttgaatccactgccccgggaatctcggtcgtaatgatttctataatgacgaaaaaaaaaaaattggaaagaaaaa<br>gcttcatggcctttataaaaaggaactatccaatacctcgccagaaccaagtaacagtattttacggggcacaaatcaagaacaat<br>aagacaggactgtaaagatggacgcattgaactccaaagaacaacaagagttccaaaaagtagtggaacaaaagcaaatgaa<br>ggatttcatgcgtttgtactctaatctggtagaaagatgtttcacagactgtgtcaatgacttcacaacatcaaagctaaccaataag<br>gaacaaacatgcatcatgaagtgctcagaaaagttcttgaagcatagcgaacgtgtagggcagcgtttccaagaacaaaacgct<br>gccttgggacaaggcttgggccgataaggtgtactggcgtatatatatctaattatgtatctctggtgtagcccatttttagcatgtaa<br>atataaagaagagacctatcagctcactcaaaggcggtaatacggttatccacagaatcaggggataacgcaggaaagaacat<br>gtgagcaaaaggccagcaaaaggccaggaaccgtaaaaaggccgcgttgctggcgtttttccataggctccgcccccctgac<br>gagcatcacaaaaatcgacgctcaagtcagaggtggcgaaacccgacaggactataaagataccaggcgtttccccctggaa<br>gctccctcgtgcgctctcctgttccgaccctgccgcttaccggatacctgtccgcctttctcccttcgggaagcgtggcgctttctc<br>atagctcacgctgtaggtatctcagttcggtgtaggtcgttcgctccaagctgggctgtgtgcacgaaccccccgttcagcccga<br>ccgctgcgccttatccggtaactatcgtcttgagcccaacccggtaagacacgacttatcgccactggcagcagccactggtaa<br>caggattagcagagcgaggtatgtaggcggtgctacagagttcttgaagtggtggcctaactacggctacactagaagaacagt<br>atttggtatctgcgctctgctgaagccagttaccttcggaaaaagagttggtagctcttgatccggcaaacaaaccaccgctggta<br>gcggtggtttttttgtttgcaagcagcagattacgcgcagaaaaaaaggatctcaagaagatcctttgatcttttctacggggtctg<br>acgctcagtggaacgaaaactcacgttaagggattttggtcatgagattatcaaaaaggatcttcacctagatccttttaaattaaaa<br>atgaagttttaaatcaatctaaagtatatatgagtaaacttggtctgacagagttctgaggtcattactggatctatcaacagcagtcc<br>aagcgagctcgatatcaaattacgccccgccctgccactcatcgcagtactgttgtaattcattaagcattctgccgacatggaag<br>ccatcacaaacggcatgatgaacctgaatcgccagcggcatcagcaccttgtcgccttgcgtataatatttgcccatggtgaaaa<br>cgggggcgaagaagttgtccatattggccacgtttaaatcaaaactggtgaaactcacccagggattggctgagacgaaaaac<br>atattctcaataaaccctttagggaaataggccaggttttcaccgtaacacgccacatcttgcgaatatatgtgtagaaactgccgg<br>aaatcgtcgtggtattcactccagagcgatgaaaacgtttcagtttgctcatggaaaacggtgtaacaagggtgaacactatccca<br>tatcaccagctcaccgtctttcattgccatacgaaattccggatgagcattcatcaggcgggcaagaatgtgaataaaggccgga<br>taaaacttgtgcttatttttctttacggtctttaaaaaggccgtaatatccagctgaacggtctggttataggtacattgagcaactgac<br>tgaaatgcctcaaaatgttctttacgatgccattgggatatatcaacggtggtatatccagtgatttttttctccattttagcttccttagc<br>tcctgaaaatctcgataactcaaaaaatacgcccggtagtgatcttatttcattatggtgaaagttggaacctcttacgtgcccgatc<br>aactcgcgcgtttgccacctgacgtctaagaaaaggaatattcagcaatttgcccgtgccgaagaaaggcccacccgtgaaggt<br>gagcc | |
| LXR370 | agaacagtaaaataaagcaaggtacgtgaaattaatatttttaaatggttctaaccgatgccgaagaactgcgcagtccggttata<br>acgtctgacatgtcctttttttgatttggaatccaaccactcaagtgactctgttcatttactttgcgaaaaatatacccacaaattgccc<br>atcgaaagtgaatcgcaaaccaccttcagactggcaccgacaaagcaaagattatacagacagagtactttatacgtaccgttaa<br>gtctcaagcaaagggttttcttatttactgaacgggtaaagagtatctgggccggcttgccaagatgcaaaccgaataagtatttca<br>aagttgcatttgccttagccgtcctgacaccattggctatttggatattttatattgactttcgtgtacattgatcacatcgactgttctat<br>tggcaaatgaaccacgggcattgactatttttcaggttactactatatattatcatcacgggcaaggattgtaGCTTGCTAT<br>GTCGTCGGAGGAGATATTTATTACTTTTATTATTCTAGTTTTTTACAGTTATTTAT<br>TAATTAATTATTTTTATATGCATGCGAATAAAAAGTCTATATTTAAGTTCTTTTAT | BpiI |

| | |
|---|---|
| | TTATTAATACATTTTCCTCTACGAGCTGTCACCGGATGTGCTTTCCGGTCTGATG<br>AGTCCGTGAGGACGAAACAGCCTCTACAAATAATTTTGTTTAAGAGCAGGTTG<br>TTCATGGCCGTGCGTATGATGTGGGGGGCTCGGGCGTTGAAACCGGGGTTCG<br>GAGCGCCAGGGGGTTTTcaacggcCTAGCATGTGATTAATTAATTATTTTGTTTTTT<br>TTTTGCAGTATAAAAAGTTAGTTTGTTTAAACAACAAACTTTTTTCATTTCTTT<br>TGTTTCCCCTTCTCTTCTTTTAGTTAGTTTGTTTAAACAACAAACTAGAATATC<br>AAGCTACAAAAATAAATAAAAaatgAGGTCTTCTTATATTCCCCAGAACATCAGG<br>TTAATGGCGTTTTTGATGTCATTTTCGCGGTGGCTGAGATCAGCCACTTCTTCC<br>CCGATAACGGAAACCGGCACACTGGCCATATCGGTGGTCATCATGCGCCAGCT<br>TTCATCCCCGATATGCACCACCGGGTAAAGTTCACGGGAGACTTTATCTGACA<br>GCAGACGTGCACTGGCCAGGGGGATCACCATCCGTCGCCCGGGCGTGTCAAT<br>AATATCACTCTGTACATCCACAAACAGACGATAACGGCTCTCTCTTTTATAGGT<br>GTAAACCTTAAACTGCATTTCACCAGCCCCTGTTCTCGTCAGCAAAAGAGCCG<br>TTCATTTCAATAAACCGGGCGACCTCAGCCATCCCTTCCTGATTTTCCGCTTTC<br>CAGCGTTCGGCACGCAGACGACGGGCTTCATTCTGCATGGTTGTGCTTACCAG<br>ACCGGAGATATTGACATCATATATGCCTTGAGCAACTGATAGCTGTCGCTGTCA<br>ACTGTCACTGTAATACGCTGCTTCATAGCATACCTCTTTTTGACATACTTCGGG<br>TATACATATCAGTATATATTCTTATACCGCAAAAATCAGCGCGCAAATATGCATA<br>CTGTTATCTGGCTTTTAGTAAGCCGAAGACCTGGTAGCCCTAAGAAAAAGAGA<br>AAAGTGggtggcagtggcggaagcgggggatcaggtggttctggagggtccGATATCATGCAGGATCT<br>GCCGGGCAACGATAACAGCACCGCGGGCGCGCGTACGAAAAACAATTACGG<br>GTCTACCATCGAGGGCCTGCTCGATCTCCCGGACGACGACGCCCCCGAAGAG<br>GCGGGGCTGGCGGCTCCGCGCCTGTCCTTTCTCCCCGCGGGACACACGCGCA<br>GACTGTCGACGGCCCCCCCGACCGATGTCAGCCTGGGGGACGAatTgCACTTA<br>GACGGCGAGGACGTGGCGATGGCGCATGCCGACGCGCTAGACGATTTCGATC<br>TGGACATGTTGGGGGACGGGGATTCCCCGGGTCCGGGATTTACCCCCCACGA<br>CTCCGCCCCCTACGGCGCTCTGGATATGGCCGACTTCGAGTTTGAGCAGATGT<br>TTACCGATGCCCTTGGAATTGACGAGTACGGTGGGAAGCTTTAAtaaaAGCTTTT<br>GATTAAGCCTTCTAGTCCAAAAAACACGTTTTTTTGTCATTTATTTCATTTTCTT<br>AGAATAGTTTAGTTTATTCATTTTATAGTCACGAATGTTTTATGATTCTATATAGG<br>GTTGCAAACAAGCATTTTTCATTTTATGTTAAAACAATTTCAGGTTTACCTTTT<br>ATTCTGCTTGTGGTGACGCGTGTATCCGCCCGCTCTTTTGGTCACCCATGTATT<br>TAATTGCATAAATAATTCTTAAAAGTGGAGCTAGTCTATTTCTATTTACATACCT<br>CTCATTTCTCATTTCCTCCTGACCAGCTGAGTTTCATGGCTTGAATTACTATGTG<br>CGGGGTAATGTGGCTTGTCGTAGCCTTGGTGAAGTATTTGGCACAAGGTCTAG<br>GGCCTGTTGAGGGAGTGCAGCTTGCCTTGTCCCCGCCGGGTCACCCGGCCAG<br>CGACATGGAGGCCCAGAATACCCTCCTTGACAGTCTTGACGTGCGCAGCTCA<br>GGGGCATGATGTGACTGTCGCCCGTACATTTAGCCCATACATCCCCATGTATAA<br>TCATTTGCATCCATACATTTTGATGGCCGCACGGCGCGAAGCAAAAATTACGG<br>CTCCTCGCTGCAGACCTGCGAGCAGGGAAACGCTCCCCTCACAGACGCGTTG<br>AATTGTCCCCACGCCGCGCCCCTGTAGAGAAATATAAAAGGTTAGGATTTGCC<br>ACTGAGGTTCTTCTTTCATATACTTCCTTTTAAAATCTTGCTAGGATACAGTTCT<br>CACATCACATCCGAACATAAACAACaatGTCTAAGAATATCGTTGTCCTACCGGG<br>TGATCACGTCGGTAAAGAAGTTACTGACGAAGCTATTAAGGTCTTGAATGCCA |

| | |
|---|---|
| | TTGCTGAAGTCCGTCCAGAAATTAAGTTCAATTTCCAACATCACTTGATCGGG<br>GGTGCTGCCATCGATGCCACTGGCACTCCTTTACCAGATGAAGCTCTAGAAGC<br>CTCTAAGAAAGCCGATGCTGTCTTACTAGGTGCTGTTGGTGGTCCAAAATGGG<br>GTACGGGCGCAGTTAGACCAGAACAAGGTCTATTGAAGATCAGAAAGGAATT<br>GGGTCTATACGCCAACTTGAGGCCATGTAACTTTGCTTCTGATTCTTTACTAGA<br>TCTTTCTCCTTTGAAGCCTGAATATGCAAAGGGTACCGATTTCGTCGTCGTTAG<br>AGAATTGGTTGGTGGTATCTACTTTGGTGAAAGAAAAGAAGATGAAGGTGAC<br>GGAGTTGCTTGGGATTCTGAGAAATACAGTGTTCCTGAAGTTCAAAGAATTAC<br>AAGAATGGCTGCTTTCTTGGCATTGCAACAAAACCCACCATTACCAATCTGGT<br>CACTTGACAAGGCTAACGTGCTTGCCTCTTCCAGATTGTGGAGAAAGACTGTT<br>GAAGAAACCATCAAGACTGAGTTCCCACAATTAACTGTTCAGCACCAATTGAT<br>CGATTCTGCTGCTATGATTTTGGTTAAATCACCAACTAAGCTAAACGGTGTTGT<br>TATTACCAACAACATGTTTGGTGATATTATCTCCGATGAAGCCTCTGTTATTCCA<br>GGTTCTTTGGGTTTATTACCTTCTGCATCTCTAGCTTCCCTACCTGACACTAAC<br>AAGGCATTCGGTTTGTACGAACCATGTCATGGTTCTGCCCCAGATTTACCAGC<br>AAACAAGGTTAACCCAATTGCTACCATCTTATCTGCAGCTATGATGTTGAAGTT<br>ATCCTTGGATTTGGTTGAAGAAGGTAGGGCTCTTGAAGAAGCTGTTAGAAATG<br>TCTTGGATGCAGGTGTCAGAACCGGTGACCTTGGTGGTTCTAACTCTACCACT<br>GAGGTTGGCGATGCTATCGCCAAGGCTGTCAAGGAAATCTTGGCTTAAACAGT<br>ACTGACAATAAAAAGATTCTTGTTTTCAAGAACTTGTCATTTGTATAGTTTTTT<br>TATATTGTAGTTGTTCTATTTTAATCAAATGTTAGCGTGATTTATATTTTTTTTCG<br>CCTCGACATCATCTGCCCAGATGCGAAGTTAAGTGCGCAGAAAGTAATATCAT<br>GCGTCAATCGTATGTGAATGCTGGTCGCTATACTGCTGTCGATTCGATACTAAC<br>GCcgccatccagtgtcgacctcCATTactcgtatcgcatgtcggtgcgacacgaaattacaaaatggaatatgttcataggg<br>tagacgaaactatatacgcaatctacatacatttatcaagaaggagaaaaaggaggatgtaaaggaatacaggtaagcaaattg<br>atactaatggctcaacgtgataaggaaaaagaattgcactttaacattaatattgacaaggaggagggcaccacacaaaaagtta<br>ggtgtaacagaaaatcatgaaactatgattcctaatttatatattggaggattttctctaaaaaaaaaaaaaatacaacaaataaaaaa<br>cactcaatgacctgaccatttgatggagtttaagtcaataccttcttgaaccatttcccataatggtgaaagttccctcaagaattta<br>ctctgtcagaaacggccttaacgacgtagtcgacctcctcttcagtactaaatctaccaataccaaatctgatggaagaatgggct<br>aatgcatcatccttacccagcgaGAGACCTATCAGCTCACTCAAAGGCGGTAATACGGTTAT<br>CCACAGAATCAGGGGATAACGCAGGAAAGAACATGTGAGCAAAAGGCCAGC<br>AAAAGGCCAGGAACCGTAAAAAGGCCGCGTTGCTGGCGTTTTTCCATAGGCT<br>CCGCCCCCCTGACGAGCATCACAAAAATCGACGCTCAAGTCAGAGGTGGCGA<br>AACCCGACAGGACTATAAAGATACCAGGCGTTTCCCCCTGGAAGCTCCCTCGT<br>GCGCTCTCCTGTTCCGACCCTGCCGCTTACCGGATACCTGTCCGCCTTTCTCCC<br>TTCGGGAAGCGTGGCGCTTTCTCATAGCTCACGCTGTaggtatctcagttcggtgtaggtcgtt<br>cgctccaagctgggctgtgtgcacgaaccccccgttcagcccgaccgctgcgccttatccggtaactatcgtcttgagcccaac<br>ccggtaagacacgacttatcgccactggcagcagccactggtaacaggattagcagagcgaggtatgtaggcggtgctacag<br>agttcttgaagtggtggcctaactacggctacactagaagaacagtatttggtatctgcgctctgctgaagccagttaccttcgga<br>aaaagagttggtagctcttgatccggcaaacaaaccaccgctggtagcggtggtttttttgtttgcaagcagcagattacgcgca<br>gaaaaaaaggatctcaagaagatcctttgatcttttctacggggtctgacgctcagtggaacgaaaactcacgttaagggattttg<br>gtcatgagattatcaaaaaggatcttcacctagatccttttaaattaaaaatgaagttttaaatcaatctaaagtatatatgagtaaact<br>tggtctgacagagttctgaggtcattactggatctatcaacagcagtccaagcgagctcgatatcaaattacgccccgccctgcc<br>actcatcgcagtactgttgtaattcattaagcattctgccgacatggaagccatcacaaacggcatgatgaacctgaatcgccagc |

| | |
|---|---|
| | ggcatcagcaccttgtcgccttgcgtataatatttgcccatggtgaaaacgggggcgaagaagttgtccatattggccacgtttaaatcaaaactggtgaaactcacccagggattggctgagacgaaaaacatattctcaataaaccctttagggaaataggccaggttttcaccgtaacacgccacatcttgcgaatatatgtgtagaaactgccggaaatcgtcgtggtattcactccagagcgatgaaaacgtttcagtttgctcatggaaaacggtgtaacaagggtgaacactatcccatatcaccagctcaccgtctttcattgccatacgaaattccggatgagcattcatcaggcgggcaagaatgtgaataaaggccggataaaacttgtgcttatttttctttacggtctttaaaaaggccgtaatatccagctgaacggtctggttataggtacattgagcaactgactgaaatgcctcaaaatgttctttacgatgccattgggatatatcaacggtggtatatccagtgatttttttctccattttagcttccTTAGCTCCTGAAAATCTCGATAACTCAAAAAATACGCCCGGTAGTGATCTTATTTCATTATGGTGAAAGTTGGAACCTCTTACGTGCCCGATCAACTCGCGCGTTTGCCACCTGACGTCTAAGAAAAGGAATATTCAGCAATTTGCCCGTGCCGAAGAAAGGCCCACCCGTGAAGGTGAGCCGGTCTCgacccaga |

#### Supplementary Table 7: List of Plasmids

| Plasmid | Features | Origin | Resistance | Figure |
|---|---|---|---|---|
| pXJH17 | $P_{S.\ a0b0c0}$-yEmCitrine-$T_{ENO1}$ | ColE1 | Chl | Fig.2b |
| pXJH20 | $P_{S.\ a0b0c0(adr1)2m}$-yEmCitrine-$T_{ENO1}$ | ColE1 | Chl | Fig.2b |
| pXJH21 | $P_{S.\ a0b0c0\Delta mig1}$-yEmCitrine-$T_{ENO1}$ | ColE1 | Chl | Fig.2b |
| pXJH22 | $P_{S.\ a0b0c0mig1m}$-yEmCitrine-$T_{ENO1}$ | ColE1 | Chl | Fig.2b |
| pXJH26 | $P_{S.\ a0b0c0(adr1)1m}$-yEmCitrine-$T_{ENO1}$ | ColE1 | Chl | Fig.2b |
| pXJh68 | $P_{S.\ a0b0c0\Delta 89}$-yEmCitrine-$T_{ENO1}$ | ColE1 | Chl | Fig.2b |
| pXJH69 | $P_{S.\ a0b0c0\Delta 89k}$-yEmCitrine-$T_{ENO1}$ | ColE1 | Chl | Fig.2b |
| pXJH72 | $P_{S.\ a0b0c0\Delta 89k\_galO}$-yEmCitrine-$T_{ENO1}$ | ColE1 | Chl | Fig.2b |
| pXJH78 | $P_{S.\ a0b0c0\Delta 89k\_4\times kb}$-yEmCitrine-$T_{ENO1}$ | ColE1 | Chl | Fig.2b |
| pXJH79 | $P_{4\times rpaO}$-yEmCitrine-$T_{ENO1}$ | ColE1 | Chl | Fig.2b, Fig.3b |
| pXJH83 | $P_{S.\ a0b0c0\Delta 135k}$-yEmCitrine-$T_{ENO1}$ | ColE1 | Chl | Fig.2b |
| pXJH84 | $P_{tet}$-RpaR-VP16-$T_{ENO1}$ | ColE1 | Chl | Fig.2b, Fig.3b |
| pXJH104 | $P_{S.\ a0b0c0\Delta 89k\_lexOec}$-yEmCitrine-$T_{ENO1}$ | ColE1 | Chl | Fig.2b |
| pXJH105 | $P_{S.\ a0b0c0\Delta 89k\_2\times lexOec}$-yEmCitrine-$T_{ENO1}$ | ColE1 | Chl | Fig.2b |
| pXJH106 | $P_{S.\ a0b0c0\Delta 89k\_3\times lexOec}$-yEmCitrine-$T_{ENO1}$ | ColE1 | Chl | Fig.2b |
| pXJH107 | $P_{S.\ a0b0c0\Delta 89k\_4\times lexOec}$-yEmCitrine-$T_{ENO1}$ | ColE1 | Chl | Fig.2b |
| pXJH119 | $P_{tet}$-$LexA_{ec87}$-$RpaR_{179}$-VP16-$T_{ENO1}$ | ColE1 | Chl | Fig.2b, Fig.3e |
| pXJH120 | $P_{S.\ a0b1c0}$-yEmCitrine-$T_{ENO1}$ | ColE1 | Chl | Fig.2b |
| LXR428 | $P_{xyluas}$-$lexA_{mm115}$-$DHBR_{282-595}$-VP16-$T_{ENO1}$ | ColE1 | Chl | Fig.2c |
| ZY754 | *Spacerup8*- $P_{S.\ a0b1c0\_lexOmm}$-yEmCitrine-$T_{ENO1}$ | ColE1 | Chl | Fig.2c |
| ZY772 | *Spacerup8*- $P_{S.\ a0b1c0\_lexOmm\times 2}$-yEmCitrine-$T_{ENO1}$ | ColE1 | Chl | Fig.2c |
| ZY773 | *Spacerup8*- $P_{S.\ a0b1c0\_lexOmm\times 3}$-yEmCitrine-$T_{ENO1}$ | ColE1 | Chl | Fig.2c |
| ZY774 | *Spacerup8*- $P_{S.\ a0b1c0\_lexOmm\times 4}$-yEmCitrine-$T_{ENO1}$ | ColE1 | Chl | Fig.2c |
| pXJH125 | $P_{S.\ a052c0}$-yEmCitrine-$T_{ENO1}$ | ColE1 | Chl | Fig.2d |
| pXJH128 | $P_{S.\ a071c0}$-yEmCitrine-$T_{ENO1}$ | ColE1 | Chl | Fig.2d |
| pXJH134 | $P_{S.\ a046c0}$-yEmCitrine-$T_{ENO1}$ | ColE1 | Chl | Fig.2d |
| pXJH135 | $P_{S.\ a041c0}$-yEmCitrine-$T_{ENO1}$ | ColE1 | Chl | Fig.2d |
| pXJH137 | $P_{S.\ a036c0}$-yEmCitrine-$T_{ENO1}$ | ColE1 | Chl | Fig.2d |
| pXJH140 | $P_{S.\ a031c0}$-yEmCitrine-$T_{ENO1}$ | ColE1 | Chl | Fig.2d |
| pXJH141 | $P_{S.\ a026c0}$-yEmCitrine-$T_{ENO1}$ | ColE1 | Chl | Fig.2d |
| pXJH142 | $P_{S.\ a016c0}$-yEmCitrine-$T_{ENO1}$ | ColE1 | Chl | Fig.2d |
| pXJH143 | $P_{S.\ a08c0}$-yEmCitrine-$T_{ENO1}$ | ColE1 | Chl | Fig.2d |
| pXJH144 | $P_{S.\ a081c0}$-yEmCitrine-$T_{ENO1}$ | ColE1 | Chl | Fig.2d |
| pXJH145 | $P_{S.\ a091c0}$-yEmCitrine-$T_{ENO1}$ | ColE1 | Chl | Fig.2d |
| pXJH146 | $P_{S.\ a0101c0}$-yEmCitrine-$T_{ENO1}$ | ColE1 | Chl | Fig.2d |
| pXJH147 | $P_{S.\ a0111c0}$-yEmCitrine-$T_{ENO1}$ | ColE1 | Chl | Fig.2d |
| pXJH148 | $P_{S.\ a0121c0}$-yEmCitrine-$T_{ENO1}$ | ColE1 | Chl | Fig.2d |
| pXJH149 | $P_{S.\ a0141c0}$-yEmCitrine-$T_{ENO1}$ | ColE1 | Chl | Fig.2d |
| pXJH150 | $P_{S.\ a0161c0}$-yEmCitrine-$T_{ENO1}$ | ColE1 | Chl | Fig.2d |
| pXJH151 | $P_{S.\ a0181c0}$-yEmCitrine-$T_{ENO1}$ | ColE1 | Chl | Fig.2d |
| pXJH152 | $P_{S.\ a0201c0}$-yEmCitrine-$T_{ENO1}$ | ColE1 | Chl | Fig.2d |
| pXJH153 | $P_{S.\ a0221c0}$-yEmCitrine-$T_{ENO1}$ | ColE1 | Chl | Fig.2d |
| pXJH154 | $P_{S.\ a0241c0}$-yEmCitrine-$T_{ENO1}$ | ColE1 | Chl | Fig.2d |
| pXJH155 | $P_{S.\ a1b1c0}$-yEmCitrine-$T_{ENO1}$ | ColE1 | Chl | Fig.2d |
| pXJH156 | $P_{S.\ a2b1c0}$-yEmCitrine-$T_{ENO1}$ | ColE1 | Chl | Fig.2d |

| | | | | |
|---|---|---|---|---|
| pXJH157 | $P_{S.\ a3b1c0}$-yEmCitrine-$T_{ENO1}$ | ColE1 | Chl | Fig.2d |
| pXJH158 | $P_{S.\ a4b1c0}$-yEmCitrine-$T_{ENO1}$ | ColE1 | Chl | Fig.2d |
| pXJH159 | $P_{S.\ a5b1c0}$-yEmCitrine-$T_{ENO1}$ | ColE1 | Chl | Fig.2d |
| pXJH161 | $P_{S.\ a0b1core1}$-yEmCitrine-$T_{ENO1}$ | ColE1 | Chl | Fig.2d |
| pXJH162 | $P_{S.\ a0b1core2}$-yEmCitrine-$T_{ENO1}$ | ColE1 | Chl | Fig.2d |
| pXJH166 | $P_{S.\ a0b1TDH3}$-yEmCitrine-$T_{ENO1}$ | ColE1 | Chl | Fig.2d |
| pXJH167 | $P_{S.\ a0b1RNR1}$-yEmCitrine-$T_{ENO1}$ | ColE1 | Chl | Fig.2d |
| pXJH168 | $P_{S.\ a0b1TEF2}$-yEmCitrine-$T_{ENO1}$ | ColE1 | Chl | Fig.2d |
| pXJH169 | $P_{S.\ a0b1ALD6}$-yEmCitrine-$T_{ENO1}$ | ColE1 | Chl | Fig.2d |
| pXJH170 | $P_{S.\ a0b1CYC1}$-yEmCitrine-$T_{ENO1}$ | ColE1 | Chl | Fig.2d |
| pXJH171 | $P_{S.\ a0b1LEU2}$-yEmCitrine-$T_{ENO1}$ | ColE1 | Chl | Fig.2d |
| pXJH176 | $P_{S.\ a0b1tdh3}$-yEmCitrine-$T_{ENO1}$ | ColE1 | Chl | Fig.2d |
| pXJH177 | $P_{S.\ a0b1rnr1}$-yEmCitrine-$T_{ENO1}$ | ColE1 | Chl | Fig.2d |
| pXJH178 | $P_{S.\ NAb1c0}$-yEmCitrine-$T_{ENO1}$ | ColE1 | Chl | Fig.2d |
| pXJH227 | $P_{S.\ a0b1c1}$-yEmCitrine-$T_{ENO1}$ | ColE1 | Chl | Fig. S2f |
| pXJH228 | $P_{S.\ a0b1c2}$-yEmCitrine-$T_{ENO1}$ | ColE1 | Chl | Fig. S2f |
| pXJH229 | $P_{S.\ a0b1c3}$-yEmCitrine-$T_{ENO1}$ | ColE1 | Chl | Fig. S2f |
| pXJH230 | $P_{S.\ a0b1c4}$-yEmCitrine-$T_{ENO1}$ | ColE1 | Chl | Fig. S2f |
| pXJH231 | $P_{S.\ a0b1c5}$-yEmCitrine-$T_{ENO1}$ | ColE1 | Chl | Fig. S2f |
| pXJH232 | $P_{S.\ a0b1c6}$-yEmCitrine-$T_{ENO1}$ | ColE1 | Chl | Fig. S2f |
| pXJH233 | $P_{S.\ a0b1c7}$-yEmCitrine-$T_{ENO1}$ | ColE1 | Chl | Fig. S2f |
| pXJH234 | $P_{S.\ a0b1c8}$-yEmCitrine-$T_{ENO1}$ | ColE1 | Chl | Fig. S2f |
| pXJH235 | $P_{S.\ a0b1c9}$-yEmCitrine-$T_{ENO1}$ | ColE1 | Chl | Fig. S2f |
| pXJH236 | $P_{S.\ a0b1c10}$-yEmCitrine-$T_{ENO1}$ | ColE1 | Chl | Fig. S2f |
| pXJH237 | $P_{S.\ a0b1c11}$-yEmCitrine-$T_{ENO1}$ | ColE1 | Chl | Fig. S2f |
| pXJH238 | $P_{S.\ a0b1c12}$-yEmCitrine-$T_{ENO1}$ | ColE1 | Chl | Fig. S2f |
| pXJH239 | $P_{S.\ a0b2c0}$-yEmCitrine-$T_{ENO1}$ | ColE1 | Chl | Fig. S2f |
| pXJH241 | $P_{S.\ a0b3c0}$-yEmCitrine-$T_{ENO1}$ | ColE1 | Chl | Fig. S2f |
| pXJH254 | $P_{S.\ a0b1c13}$-yEmCitrine-$T_{ENO1}$ | ColE1 | Chl | Fig. S2f |
| pXJH255 | $P_{S.\ a0b1c4}$-yEmCitrine-$T_{ENO1}$ | ColE1 | Chl | Fig. S2f |
| pXJH256 | $P_{S.\ a0b1c15}$-yEmCitrine-$T_{ENO1}$ | ColE1 | Chl | Fig. S2f |
| pXJH257 | $P_{S.\ a0b1c16}$-yEmCitrine-$T_{ENO1}$ | ColE1 | Chl | Fig. S2f |
| pXJH258 | $P_{S.\ a0b1c17}$-yEmCitrine-$T_{ENO1}$ | ColE1 | Chl | Fig. S2f |
| pXJH259 | $P_{S.\ a0b1c18}$-yEmCitrine-$T_{ENO1}$ | ColE1 | Chl | Fig. S2f |
| pXJH260 | $P_{S.\ a0b1c19}$-yEmCitrine-$T_{ENO1}$ | ColE1 | Chl | Fig. S2f |
| pXJH261 | $P_{S.\ a0b1c20}$-yEmCitrine-$T_{ENO1}$ | ColE1 | Chl | Fig. S2f |
| pXJH264 | $P_{S.\ a0b4c0}$-yEmCitrine-$T_{ENO1}$ | ColE1 | Chl | Fig. S2f |
| pXJH265 | $P_{S.\ a0b5c0}$-yEmCitrine-$T_{ENO1}$ | ColE1 | Chl | Fig. S2f |
| pXJH266 | $P_{S.\ a0b6c0}$-yEmCitrine-$T_{ENO1}$ | ColE1 | Chl | Fig. S2f |
| pXJH267 | $P_{S.\ a0b7c0}$-yEmCitrine-$T_{ENO1}$ | ColE1 | Chl | Fig. S2f |
| pXJH268 | $P_{S.\ a0b8c0}$-yEmCitrine-$T_{ENO1}$ | ColE1 | Chl | Fig. S2f |
| pXJH269 | $P_{S.\ a0b9c0}$-yEmCitrine-$T_{ENO1}$ | ColE1 | Chl | Fig. S2f |
| pXJH270 | $P_{S.\ a0b10c0}$-yEmCitrine-$T_{ENO1}$ | ColE1 | Chl | Fig. S2f |
| pXJH271 | $P_{S.\ a0b11c0}$-yEmCitrine-$T_{ENO1}$ | ColE1 | Chl | Fig. S2f |
| pXJH272 | $P_{S.\ a0b12c0}$-yEmCitrine-$T_{ENO1}$ | ColE1 | Chl | Fig. S2f |
| pXJH273 | $P_{S.\ a0b13c0}$-yEmCitrine-$T_{ENO1}$ | ColE1 | Chl | Fig. S2f |
| pXJH274 | $P_{S\ a0b14c0}$-yEmCitrine-$T_{ENO1}$ | ColE1 | Chl | Fig. S2f |
| pXJH275 | $P_{S.\ a0b15c0}$-yEmCitrine-$T_{ENO1}$ | ColE1 | Chl | Fig. S2f |
| pXJH276 | $P_{S.\ a0b16c0}$-yEmCitrine-$T_{ENO1}$ | ColE1 | Chl | Fig. S2f |

| | | | | |
|---|---|---|---|---|
| pXJH277 | $P_{S.\ a0b17c0}$-yEmCitrine-$T_{ENO1}$ | ColE1 | Chl | Fig. S2f |
| pXJH278 | $P_{S.\ a0b1c21}$-yEmCitrine-$T_{ENO1}$ | ColE1 | Chl | Fig. S2f |
| pXJH279 | $P_{S.\ a0b1c22}$-yEmCitrine-$T_{ENO1}$ | ColE1 | Chl | Fig. S2f |
| pXJH280 | $P_{S.\ a0b1c23}$-yEmCitrine-$T_{ENO1}$ | ColE1 | Chl | Fig. S2f |
| pXJH281 | $P_{S.\ a0b1c24}$-yEmCitrine-$T_{ENO1}$ | ColE1 | Chl | Fig. S2f |
| pXJH282 | $P_{S.\ a0b1c25}$-yEmCitrine-$T_{ENO1}$ | ColE1 | Chl | Fig. S2f |
| pXJH283 | $P_{S.\ a0b1c26}$-yEmCitrine-$T_{ENO1}$ | ColE1 | Chl | Fig. S2f |
| pXJH284 | $P_{S.\ a0b1c27}$-yEmCitrine-$T_{ENO1}$ | ColE1 | Chl | Fig. S2f |
| pXJH285 | $P_{S.\ a0b1c28}$-yEmCitrine-$T_{ENO1}$ | ColE1 | Chl | Fig. S2f |
| pXJH298 | $P_{S.\ a0b23c0}$-yEmCitrine-$T_{ENO1}$ | ColE1 | Chl | Fig. S2f |
| pXJH299 | $P_{S.\ a0b24c0}$-yEmCitrine-$T_{ENO1}$ | ColE1 | Chl | Fig. S2f |
| pXJH300 | $P_{S.\ a0b25c0}$-yEmCitrine-$T_{ENO1}$ | ColE1 | Chl | Fig. S2f |
| pXJH301 | $P_{S.\ a0b26c0}$-yEmCitrine-$T_{ENO1}$ | ColE1 | Chl | Fig. S2f |
| pXJH302 | $P_{S.\ a0b27c0}$-yEmCitrine-$T_{ENO1}$ | ColE1 | Chl | Fig. S2f |
| pXJH304 | $P_{S.\ a0b29c0}$-yEmCitrine-$T_{ENO1}$ | ColE1 | Chl | Fig. S2f |
| pXJH305 | $P_{S.\ a0b30c0}$-yEmCitrine-$T_{ENO1}$ | ColE1 | Chl | Fig. S2f |
| pXJH318 | $P_{S.\ a6b1c0}$-yEmCitrine-$T_{ENO1}$ | ColE1 | Chl | Fig. S2f |
| pXJH319 | $P_{S.\ a7b1c0}$-yEmCitrine-$T_{ENO1}$ | ColE1 | Chl | Fig. S2f |
| pXJH320 | $P_{S.\ a8b1c0}$-yEmCitrine-$T_{ENO1}$ | ColE1 | Chl | Fig. S2f |
| pXJH321 | $P_{S.\ a9b1c0}$-yEmCitrine-$T_{ENO1}$ | ColE1 | Chl | Fig. S2f |
| pXJH322 | $P_{S.\ a10b1c0}$-yEmCitrine-$T_{ENO1}$ | ColE1 | Chl | Fig. S2f |
| pXJH323 | $P_{S.\ a11b1c0}$-yEmCitrine-$T_{ENO1}$ | ColE1 | Chl | Fig. S2f |
| pXJH324 | $P_{S.\ a12b1c0}$-yEmCitrine-$T_{ENO1}$ | ColE1 | Chl | Fig. S2f |
| pXJH325 | $P_{S.\ a13b1c0}$-yEmCitrine-$T_{ENO1}$ | ColE1 | Chl | Fig. S2f |
| pXJH326 | $P_{S.\ a14b1c0}$-yEmCitrine-$T_{ENO1}$ | ColE1 | Chl | Fig. S2f |
| pXJH327 | $P_{S.\ a15b1c0}$-yEmCitrine-$T_{ENO1}$ | ColE1 | Chl | Fig. S2f |
| pXJH328 | $P_{S.\ a16b1c0}$-yEmCitrine-$T_{ENO1}$ | ColE1 | Chl | Fig. S2f |
| pXJH329 | $P_{S.\ a17b1c0}$-yEmCitrine-$T_{ENO1}$ | ColE1 | Chl | Fig. S2f |
| pXJH330 | $P_{S.\ a18b1c0}$-yEmCitrine-$T_{ENO1}$ | ColE1 | Chl | Fig. S2f |
| pXJH331 | $P_{S.\ a19b1c0}$-yEmCitrine-$T_{ENO1}$ | ColE1 | Chl | Fig. S2f |
| pXJH332 | $P_{S.\ a20b1c0}$-yEmCitrine-$T_{ENO1}$ | ColE1 | Chl | Fig. S2f |
| pXJH333 | $P_{S.\ a21b1c0}$-yEmCitrine-$T_{ENO1}$ | ColE1 | Chl | Fig. S2f |
| pXJH334 | $P_{S.\ a22b1c0}$-yEmCitrine-$T_{ENO1}$ | ColE1 | Chl | Fig. S2f |
| pXJH335 | $P_{S.\ a23b1c0}$-yEmCitrine-$T_{ENO1}$ | ColE1 | Chl | Fig. S2f |
| pXJH336 | $P_{S.\ a24b1c0}$-yEmCitrine-$T_{ENO1}$ | ColE1 | Chl | Fig. S2f |
| pXJH337 | $P_{S.\ a25b1c0}$-yEmCitrine-$T_{ENO1}$ | ColE1 | Chl | Fig. S2f |
| pXJH360 | $P_{S.\ a10b1c20}$-yEmCitrine-$T_{ENO1}$ | ColE1 | Chl | Fig.2f |
| pXJH361 | $P_{S.\ a20b25c18}$-yEmCitrine-$T_{ENO1}$ | ColE1 | Chl | Fig.2f |
| pXJH362 | $P_{S.\ a9b7c19}$-yEmCitrine-$T_{ENO1}$ | ColE1 | Chl | Fig.2f |
| pXJH363 | $P_{S.\ a14b5c21}$-yEmCitrine-$T_{ENO1}$ | ColE1 | Chl | Fig.2f |
| pXJH364 | $P_{S.\ a24b23c22}$-yEmCitrine-$T_{ENO1}$ | ColE1 | Chl | Fig.2f |
| pXJH365 | $P_{S.\ a18b8c15}$ -yEmCitrine-$T_{ENO1}$ | ColE1 | Chl | Fig.2f |
| pXJH366 | $P_{S.\ a22b27c16}$-yEmCitrine-$T_{ENO1}$ | ColE1 | Chl | Fig.2f |
| pXJH367 | $P_{S.\ a21b11c23}$-yEmCitrine-$T_{ENO1}$ | ColE1 | Chl | Fig.2f |
| pXJH368 | $P_{S.\ a12b17c24}$-yEmCitrine-$T_{ENO1}$ | ColE1 | Chl | Fig.2f |
| pXJH369 | $P_{S.\ a19b26c6}$-yEmCitrine-$T_{ENO1}$ | ColE1 | Chl | Fig.2f |
| pXJH370 | $P_{S.\ a6b14c7}$-yEmCitrine-$T_{ENO1}$ | ColE1 | Chl | Fig.2f |
| pXJH371 | $P_{S.\ a7b15c8}$-yEmCitrine-$T_{ENO1}$ | ColE1 | Chl | Fig.2f |
| pXJH372 | $P_{S.\ a21b1c15}$-yEmCitrine-$T_{ENO1}$ | ColE1 | Chl | Fig.2f |

| | | | | |
|---|---|---|---|---|
| pXJH373 | $P_{S.\ a14b1c16}$-yEmCitrine-$T_{ENO1}$ | ColE1 | Chl | Fig.2f |
| pXJH374 | $P_{S.\ a9b1c24}$-yEmCitrine-$T_{ENO1}$ | ColE1 | Chl | Fig.2f |
| pXJH375 | $P_{S.\ a14b25c24}$-yEmCitrine-$T_{ENO1}$ | ColE1 | Chl | Fig.2f |
| pXJH376 | $P_{S.\ a21b25c15}$-yEmCitrine-$T_{ENO1}$ | ColE1 | Chl | Fig.2f |
| pXJH377 | $P_{S.\ a22b26c22}$-yEmCitrine-$T_{ENO1}$ | ColE1 | Chl | Fig.2f |
| pXJH378 | $P_{S.\ a20b25c6}$-yEmCitrine-$T_{ENO1}$ | ColE1 | Chl | Fig.2f |
| pXJH379 | $P_{S.\ a12b7c16}$-yEmCitrine-$T_{ENO1}$ | ColE1 | Chl | Fig.2f |
| pXJH380 | $P_{S.\ a14b8c19}$-yEmCitrine-$T_{ENO1}$ | ColE1 | Chl | Fig.2f |
| pXJH381 | $P_{S.\ a9b27c23}$-yEmCitrine-$T_{ENO1}$ | ColE1 | Chl | Fig.2f |
| pXJH382 | $P_{S.\ a24b23c21}$-yEmCitrine-$T_{ENO1}$ | ColE1 | Chl | Fig.2f |
| LXR453 | $P_{tet}$-lexA-VP16-$T_{ENO1}$ | ColE1 | Chl | Fig.3b |
| pXJH84 | $P_{tet}$-RpaR nls-VP16-$T_{ENO1}$ | ColE1 | Chl | Fig.3b |
| LXR455 | $P_{tet}$-zif268-RpaR nls-VP16-$T_{ENO1}$ | ColE1 | Chl | Fig.3b |
| LXR456 | $P_{tet}$-zif268-$RpaR_{179}$ nls-VP16-$T_{ENO1}$ | ColE1 | Chl | Fig.3b |
| HC83 | $P_{tet}$-lexA-$RpaR_{179}$ nls-VP16-$T_{ENO1}$ | ColE1 | Chl | Fig.3b |
| HC106 | $P_{tet}$-$lexA_{ec87}$-$RpaR_{179}$ nls-VP16-$T_{ENO1}$ | ColE1 | Chl | Fig.3b |
| CYx52 | $P_{4\times lexOec}$-yEmCitrine-$T_{ENO1}$ | ColE1 | Chl | Fig.3b |
| LXR300 | $P_{4\times zifO}$-yEmCitrine-$T_{ENO1}$ | ColE1 | Chl | Fig.3b |
| LXR31 | $P_{llac}$-BCD14-$lexA_{ec87}$-$(GGGGS)_3$-$RpaR_{179}$-$T_7$ | pSC101 | Chl | Fig.3d, Fig.3g |
| LXR33 | $P_{llac}$-BCD14-$lexA_{mt127}$-$(GGGGS)_3$-$RpaR_{179}$-$T_7$ | pSC101 | Chl | Fig.3d |
| LXR34 | $P_{llac}$-BCD14-$lexA_{cg145}$-$(GGGGS)_3$-$RpaR_{179}$-$T_7$ | pSC101 | Chl | Fig.3d |
| LXR35 | $P_{llac}$-BCD14-$lexA_{sa96}$-$(GGGGS)_3$-$RpaR_{179}$-$T_7$ | pSC101 | Chl | Fig.3d |
| LXR36 | $P_{llac}$-BCD14-$lexA_{bs94}$-$(GGGGS)_3$-$RpaR_{179}$-$T_7$ | pSC101 | Chl | Fig.3d, Fig.3g |
| LXR37 | $P_{llac}$-BCD14-$lexA_{lm93}$-$(GGGGS)_3$-$RpaR_{179}$-$T_7$ | pSC101 | Chl | Fig.3d |
| LXR38 | $P_{llac}$-BCD14-$lexA_{pd114}$-$(GGGGS)_3$-$RpaR_{179}$-$T_7$ | pSC101 | Chl | Fig.3d |
| LXR39 | $P_{llac}$-BCD14-$lexA_{sm126}$-$(GGGGS)_3$-$RpaR_{179}$-$T_7$ | pSC101 | Chl | Fig.3d |
| LXR40 | $P_{llac}$-BCD14-$lexA_{cc123}$-$(GGGGS)_3$-$RpaR_{179}$-$T_7$ | pSC101 | Chl | Fig.3d |
| LXR41 | $P_{llac}$-BCD14-$lexA_{pa93}$-$(GGGGS)_3$-$RpaR_{179}$-$T_7$ | pSC101 | Chl | Fig.3d |
| LXR42 | $P_{llac}$-BCD14-$lexA_{vp91}$-$(GGGGS)_3$-$RpaR_{179}$-$T_7$ | pSC101 | Chl | Fig.3d |
| LXR43 | $P_{llac}$-BCD14-$lexA_{pp91}$-$(GGGGS)_3$-$RpaR_{179}$-$T_7$ | pSC101 | Chl | Fig.3d |
| LXR98 | $P_{llac}$-BCD14-$CI_{94}$-$(GGGGS)_3$-$RpaR_{179}$-$T_7$ | pSC101 | Chl | Fig.3d, Fig.3g |
| LXR99 | $P_{llac}$-BCD14-$CI434_{70}$-$(GGGGS)_3$-$RpaR_{179}$-$T_7$ | pSC101 | Chl | Fig.3d, Fig.3g |
| LXR100 | $P_{llac}$-BCD14-$HKCI_{84}$-$(GGGGS)_3$-$RpaR_{179}$-$T_7$ | pSC101 | Chl | Fig.3d, Fig.3g |
| LXR101 | $P_{llac}$-BCD14-$PurR_{60}$-$(GGGGS)_3$-$RpaR_{179}$-$T_7$ | pSC101 | Chl | Fig.3d, Fig.3g |
| LXR139 | $P_{llac}$-BCD14-$ArgR_{93}$-$(GGGGS)_3$-$RpaR_{179}$-$T_7$ | pSC101 | Chl | Fig.3d |
| LXR140 | $P_{llac}$-BCD14-$DeoR_{92}$-$(GGGGS)_3$-$RpaR_{179}$-$T_7$ | pSC101 | Chl | Fig.3d, Fig.3g |
| LXR160 | $P_{llac}$-BCD14-$lexA_{ac89}$-$(GGGGS)_3$-$RpaR_{179}$-$T_7$ | pSC101 | Chl | Fig.3d |
| LXR161 | $P_{llac}$-BCD14-$lexA_{fs104}$-$(GGGGS)_3$-$RpaR_{179}$-$T_7$ | pSC101 | Chl | Fig.3d, Fig.3g |
| LXR162 | $P_{llac}$-BCD14-$lexA_{gs91}$-$(GGGGS)_3$-$RpaR_{179}$-$T_7$ | pSC101 | Chl | Fig.3d, Fig.3g |
| LXR163 | $P_{llac}$-BCD14-$lexA_{mm115}$-$(GGGGS)_3$-$RpaR_{179}$-$T_7$ | pSC101 | Chl | Fig.3d, Fig.3g |
| LXR164 | $P_{llac}$-BCD14-$lexA_{mx100}$-$(GGGGS)_3$-$RpaR_{179}$-$T_7$ | pSC101 | Chl | Fig.3d |
| LXR165 | $P_{llac}$-BCD14-$lexA_{pm92}$-$(GGGGS)_3$-$RpaR_{179}$-$T_7$ | pSC101 | Chl | Fig.3d |
| LXR166 | $P_{llac}$-BCD14-$lexA_{sp87}$-$(GGGGS)_3$-$RpaR_{179}$-$T_7$ | pSC101 | Chl | Fig.3d |
| LXR167 | $P_{llac}$-BCD14-$lexA_{xa101}$-$(GGGGS)_3$-$RpaR_{179}$-$T_7$ | pSC101 | Chl | Fig.3d, Fig.3g |
| LXR168 | $P_{llac}$-BCD14-$lexA_{xa90}$-$(GGGGS)_3$-$RpaR_{179}$-$T_7$ | pSC101 | Chl | Fig.3d |
| LXR169 | $P_{llac}$-BCD14-$lexA_{xf99}$-$(GGGGS)_3$-$RpaR_{179}$-$T_7$ | pSC101 | Chl | Fig.3d |
| LXR170 | $P_{llac}$-BCD14-RecA*pact*$_{104}$-$(GGGGS)_3$-$RpaR_{179}$- | pSC101 | Chl | Fig.3d, Fig.3g |
| LXR172 | $P_{llac}$-BCD14-*Mu*$repc_{82}$-$(GGGGS)_3$-$RpaR_{179}$-$T_7$ | pSC101 | Chl | Fig.3d |
| LXR211 | $P_{llac}$-BCD14-$RutR_{86}$-(GGGGS)3-$RpaR_{179}$-$T_7$ | pSC101 | Chl | Fig.3d |

| | | | | |
|---|---|---|---|---|
| LXR212 | $P_{llac}$-BCD14-BetI$_{80}$-(GGGGS)$_{3}$-RpaR$_{179}$-T$_{7}$ | pSC101 | Chl | Fig.3d |
| LXR213 | $P_{llac}$-BCD14-ArgP$_{98}$-(GGGGS)$_{3}$-RpaR$_{179}$-T$_{7}$ | pSC101 | Chl | Fig.3d |
| LXR214 | $P_{llac}$-BCD14-FruR$_{70}$-(GGGGS)$_{3}$-RpaR$_{179}$-T$_{7}$ | pSC101 | Chl | Fig.3d |
| LXR215 | $P_{llac}$-BCD14-RbsR$_{67}$-(GGGGS)$_{3}$-RpaR$_{179}$-T$_{7}$ | pSC101 | Chl | Fig.3d |
| LXR216 | $P_{llac}$-BCD14-TreR$_{70}$-(GGGGS)$_{3}$-RpaR$_{179}$-T$_{7}$ | pSC101 | Chl | Fig.3d |
| LXR217 | $P_{llac}$-BCD14-NanR$_{107}$-(GGGGS)$_{3}$-RpaR$_{179}$-T$_{7}$ | pSC101 | Chl | Fig.3d |
| LXR218 | $P_{llac}$-BCD14-PdhR$_{83}$-(GGGGS)$_{3}$-RpaR$_{179}$-T$_{7}$ | pSC101 | Chl | Fig.3d |
| LXR219 | $P_{llac}$-BCD14-DgoR$_{89}$-(GGGGS)$_{3}$-RpaR$_{179}$-T$_{7}$ | pSC101 | Chl | Fig.3d |
| LXR220 | $P_{llac}$-BCD14-AllR$_{96}$-(GGGGS)$_{3}$-RpaR$_{179}$-T$_{7}$ | pSC101 | Chl | Fig.3d |
| LXR221 | $P_{llac}$-BCD14-MhpR$_{85}$-(GGGGS)$_{3}$-RpaR$_{179}$-T$_{7}$ | pSC101 | Chl | Fig.3d |
| LXR28 | $P_{lexOec}$-YFP-T$_{7}$ | p15A | Kan | Fig.3d, Fig.3g |
| LXR44 | $P_{lexOmt}$-YFP-T$_{7}$ | p15A | Kan | Fig.3d |
| LXR46 | $P_{lexOcg}$-YFP-T$_{7}$ | p15A | Kan | Fig.3d |
| LXR49 | $P_{lexOsa}$-YFP-T$_{7}$ | p15A | Kan | Fig.3d |
| LXR50 | $P_{lexObs}$-YFP-T$_{7}$ | p15A | Kan | Fig.3d, Fig.3g |
| LXR52 | $P_{lexOlm}$-YFP-T$_{7}$ | p15A | Kan | Fig.3d |
| LXR54 | $P_{lexOpd}$-YFP-T$_{7}$ | p15A | Kan | Fig.3d |
| LXR56 | $P_{lexOsm}$-YFP-T$_{7}$ | p15A | Kan | Fig.3d |
| LXR58 | $P_{lexOcc}$-YFP-T$_{7}$ | p15A | Kan | Fig.3d |
| LXR60 | $P_{lexOpa}$-YFP-T$_{7}$ | p15A | Kan | Fig.3d |
| LXR62 | $P_{lexOvp}$-YFP-T$_{7}$ | p15A | Kan | Fig.3d |
| LXR64 | $P_{lexOpp}$-YFP-T$_{7}$ | p15A | Kan | Fig.3d |
| LXR102 | $P_{CI}$-YFP-T$_{7}$ | p15A | Kan | Fig.3d, Fig.3g |
| LXR103 | $P_{CI434}$-YFP-T$_{7}$ | p15A | Kan | Fig.3d, Fig.3g |
| LXR104 | $P_{HKCI}$-YFP-T$_{7}$ | p15A | Kan | Fig.3d, Fig.3g |
| LXR105 | $P_{purO}$-YFP-T$_{7}$ | p15A | Kan | Fig.3d, Fig.3g |
| LXR142 | $P_{Deo}$-YFP-T$_{7}$ | p15A | Kan | Fig.3d, Fig.3g |
| LXR173 | $P_{lexOac}$-YFP-T$_{7}$ | p15A | Kan | Fig.3d |
| LXR174 | $P_{lexOfs}$-YFP-T$_{7}$ | p15A | Kan | Fig.3d, Fig.3g |
| LXR175 | $P_{lexOgs}$-YFP-T$_{7}$ | p15A | Kan | Fig.3d, Fig.3g |
| LXR176 | $P_{lexOmm}$-YFP-T$_{7}$ | p15A | Kan | Fig.3d, Fig.3g |
| LXR177 | $P_{lexOmx}$-YFP-T$_{7}$ | p15A | Kan | Fig.3d |
| LXR178 | $P_{lexOpm}$-YFP-T$_{7}$ | p15A | Kan | Fig.3d |
| LXR179 | $P_{lexOsp}$-YFP-T$_{7}$ | p15A | Kan | Fig.3d |
| LXR180 | $P_{lexOxa101}$-YFP-T$_{7}$ | p15A | Kan | Fig.3d, Fig.3g |
| LXR181 | $P_{lexOxa90}$-YFP-T$_{7}$ | p15A | Kan | Fig.3d |
| LXR182 | $P_{lexOxf}$-YFP-T$_{7}$ | p15A | Kan | Fig.3d |
| LXR183 | $P_{RecApact}$-YFP-T$_{7}$ | p15A | Kan | Fig.3d, Fig.3g |
| LXR185 | $P_{Murepc}$-YFP-T$_{7}$ | p15A | Kan | Fig.3d |
| LXR222 | $P_{RutR}$-YFP-T$_{7}$ | p15A | Kan | Fig.3d |
| LXR223 | $P_{BetI}$-YFP-T$_{7}$ | p15A | Kan | Fig.3d |
| LXR224 | $P_{ArgP}$-YFP-T$_{7}$ | p15A | Kan | Fig.3d |
| LXR225 | $P_{FruR}$-YFP-T$_{7}$ | p15A | Kan | Fig.3d |
| LXR226 | $P_{RbsR}$-YFP-T$_{7}$ | p15A | Kan | Fig.3d |
| LXR227 | $P_{TreR}$-YFP-T$_{7}$ | p15A | Kan | Fig.3d |
| LXR228 | $P_{NanR}$-YFP-T$_{7}$ | p15A | Kan | Fig.3d |
| LXR229 | $P_{PdhR}$-YFP-T$_{7}$ | p15A | Kan | Fig.3d |
| LXR230 | $P_{DgoR}$-YFP-T$_{7}$ | p15A | Kan | Fig.3d |
| LXR231 | $P_{AllR}$-YFP-T$_{7}$ | p15A | Kan | Fig.3d |

| | | | | |
|---|---|---|---|---|
| LXR232 | $P_{MhpR}$-YFP-$T_7$ | p15A | Kan | Fig.3d |
| pXJH119 | $P_{tet}$-LexA$_{ec87}$-RpaR$_{179}$-VP16-T$_{ENO1}$ | ColE1 | Chl | Fig.3e-f |
| LXR69 | $P_{tet}$-lexA$_{bs94}$-RpaR$_{179}$ nls-VP16-T$_{ENO1}$ | ColE1 | Chl | Fig.3e |
| LXR190 | $P_{tet}$-lexA$_{fs104}$ -RpaR$_{179}$ nls-VP16-T$_{ENO1}$ | ColE1 | Chl | Fig.3e |
| LXR193 | $P_{tet}$-CI$_{94}$-RpaR$_{179}$-VP16-T$_{ENO1}$ | ColE1 | Chl | Fig.3e |
| LXR194 | $P_{tet}$-CI434$_{70}$-RpaR$_{179}$-VP16-T$_{ENO1}$ | ColE1 | Chl | Fig.3e |
| LXR195 | $P_{tet}$-HKCI$_{84}$-RpaR$_{179}$-VP16-T$_{ENO1}$ | ColE1 | Chl | Fig.3e |
| LXR196 | $P_{tet}$-PurR$_{60}$-RpaR$_{179}$-VP16-T$_{ENO1}$ | ColE1 | Chl | Fig.3e |
| LXR197 | $P_{tet}$-DeoR$_{92}$-RpaR$_{179}$-VP16-T$_{ENO1}$ | ColE1 | Chl | Fig.3e |
| LXR198 | $P_{tet}$-RecApact$_{104}$-RpaR$_{179}$-VP16-T$_{ENO1}$ | ColE1 | Chl | Fig.3e |
| LXR234 | $P_{tet}$-lexA$_{gs91}$-RpaR$_{179}$-VP16-T$_{ENO1}$ | ColE1 | Chl | Fig.3e |
| LXR235 | $P_{tet}$-lexA$_{mm115}$-RpaR$_{179}$-VP16-T$_{ENO1}$ | ColE1 | Chl | Fig.3e |
| LXR236 | $P_{tet}$-lexA$_{xa101}$-RpaR$_{179}$-VP16-T$_{ENO1}$ | ColE1 | Chl | Fig.3e |
| LXR74 | $P_{S.\ a0b0c0\Delta89k\_2\times lexObs}$-yEmCitrine-T$_{ENO1}$ | ColE1 | Chl | Fig.3e |
| LXR152 | $P_{S.\ a0b0c0\Delta89k\_2\times lexOrec.sym}$-yEmCitrine -T$_{ENO1}$ | ColE1 | Chl | Fig.3e |
| LXR201 | $P_{S.\ a0b0c0\Delta89k\_2\times lexOfs}$-yEmCitrine-T$_{ENO1}$ | ColE1 | Chl | Fig.3e |
| LXR204 | $P_{S.\ a0b0c0\Delta89k\_2\times OCI\ OL1}$-yEmCitrine-T$_{ENO1}$ | ColE1 | Chl | Fig.3e |
| LXR205 | $P_{S.\ a0b0c0\Delta89k\_2\times OCI434}$-yEmCitrine -T$_{ENO1}$ | ColE1 | Chl | Fig.3e |
| LXR206 | $P_{S.\ a0b0c0\Delta89k\_2\times OHKCI}$-yEmCitrine-T$_{ENO1}$ | ColE1 | Chl | Fig.3e |
| LXR207 | $P_{S.\ a0b0c0\Delta89k\_2\times purO}$-yEmCitrine-T$_{ENO1}$ | ColE1 | Chl | Fig.3e |
| LXR208 | $P_{S.\ a0b0c0\Delta89k\_2\times ODeo}$-yEmCitrine-T$_{ENO1}$ | ColE1 | Chl | Fig.3e |
| LXR209 | $P_{S.\ a0b0c0\Delta89k\_2\times ORecApact}$-yEmCitrine-T$_{ENO1}$ | ColE1 | Chl | Fig.3e |
| LXR237 | $P_{S.\ a0b0c0\Delta89k\_2\times lexOgs}$-yEmCitrine-T$_{ENO1}$ | ColE1 | Chl | Fig.3e |
| LXR238 | $P_{S.\ a0b0c0\Delta89k\_2\times lexOmm}$-yEmCitrine-T$_{ENO1}$ | ColE1 | Chl | Fig.3e |
| LXR239 | $P_{S.\ a0b0c0\Delta89k\_2\times lexOxa}$-yEmCitrine-T$_{ENO1}$ | ColE1 | Chl | Fig.3e |
| LXR111 | $P_{tet}$-lexA$_{ec87}$-BjaR$_{180}$ nls-VP16-T$_{ENO1}$ | ColE1 | Chl | Fig.3f |
| LXR114 | $P_{tet}$-lexA$_{ec87}$-LasR$_{177}$ nls-VP16-T$_{ENO1}$ | ColE1 | Chl | Fig.3f |
| LXR150 | $P_{tet}$-lexA$_{ec87}$-CinR$_{179}$ nls-DBD-VP16-T$_{ENO1}$ | ColE1 | Chl | Fig.3f |
| LXR190 | $P_{tet}$-lexA$_{fs104}$ -RpaR$_{179}$ nls-VP16-T$_{ENO1}$ | ColE1 | Chl | Fig.3f |
| LXR210 | $P_{tet}$-lexA$_{ec87}$-acVHH-VP16-T$_{ENO1}$ | ColE1 | Chl | Fig.3f |
| LXR271 | $P_{tet}$-lexA$_{ec87}$-ER$_{282\text{-}595}$-VP16-T$_{ENO1}$ | ColE1 | Chl | Fig.3f |
| LXR272 | $P_{tet}$-lexA$_{ec87}$-DHBR$_{282\text{-}595}$-VP16-T$_{ENO1}$ | ColE1 | Chl | Fig.3f |
| LXR333 | $P_{xyluas}$-lexA$_{ec87}$-TraR$_{174}$ nls-VP16-T$_{ENO1}$ | ColE1 | Chl | Fig.3f |
| LXR334 | $P_{xyluas}$-lexA$_{ec87}$-CarRecc$_{169}$ nls-VP16-T$_{ENO1}$ | ColE1 | Chl | Fig.3f |
| LXR384 | $P_{xyluas}$-CI$_{94}$-SmaR$_{179}$ nls-VP16-T$_{ENO1}$ | ColE1 | Chl | Fig.3f |
| LXR402 | $P_{xyluas}$-lexA$_{bs94}$-PR-VP16-T$_{ENO1}$ | ColE1 | Chl | Fig.3f |
| LXR404 | $P_{xyluas}$-lexA$_{bs94}$-GR$_{487\text{-}777}$-VP16-T$_{ENO1}$ | ColE1 | Chl | Fig.3f |
| LXR405 | $P_{xyluas}$-lexA$_{bs94}$-MR$_{669\text{-}984}$-VP16-T$_{ENO1}$ | ColE1 | Chl | Fig.3f |
| LXR74 | $P_{S.\ a0b0c0\Delta89k\_2\times lexObs}$-yEmCitrine-T$_{ENO1}$ | ColE1 | Chl | Fig.3f |
| LXR152 | $P_{S.\ a0b0c0\Delta89k\_2\times lexOrec.sym}$-yEmCitrine -T$_{ENO1}$ | ColE1 | Chl | Fig.3f |
| LXR204 | $P_{S.\ a0b0c0\Delta89k\_2\times OCI\ OL1}$-yEmCitrine-T$_{ENO1}$ | ColE1 | Chl | Fig.3f |
| LXR326 | $P_{xyluas}$-lexA$_{bs94}$-LasR$_{177}$ nls-VP16-T$_{ENO1}$ | ColE1 | Chl | Fig.3h |
| LXR327 | $P_{xyluas}$-lexA$_{bs94}$-RpaR$_{179}$ nls-VP16-T$_{ENO1}$ | ColE1 | Chl | Fig.3h |
| LXR329 | $P_{xyluas}$-LexA$_{bs94}$-BjaR$_{180}$ nls-VP16-T$_{ENO1}$ | ColE1 | Chl | Fig.3h |
| LXR330 | $P_{xyluas}$-LexA$_{bs94}$-ER$_{282\text{-}595}$-VP16-T$_{ENO1}$ | ColE1 | Chl | Fig.3h |
| LXR331 | $P_{xyluas}$-LexA$_{bs94}$-DHBR$_{282\text{-}595}$-VP16-T$_{ENO1}$ | ColE1 | Chl | Fig.3h |
| LXR334 | $P_{xyluas}$-lexA$_{ec87}$-CarRecc$_{169}$ nls-VP16-T$_{ENO1}$ | ColE1 | Chl | Fig.3h |
| LXR371 | $P_{xyluas}$-lexA$_{bs94}$-TraR$_{174}$ nls-VP16-T$_{ENO1}$ | ColE1 | Chl | Fig.3h |
| LXR381 | $P_{xyluas}$-lexA$_{mm115}$-CinR$_{179}$ nls-VP16-T$_{ENO1}$ | ColE1 | Chl | Fig.3h |

| | | | | |
|---|---|---|---|---|
| LXR384 | $P_{xyluas}$-CI$_{94}$-SmaR$_{179}$ nls-VP16-T$_{ENO1}$ | ColE1 | Chl | Fig.3h |
| LXR386 | $P_{xyluas}$-HKCI$_{84}$-acVHH-VP16-T$_{ENO1}$ | ColE1 | Chl | Fig.3h |
| LXR402 | $P_{xyluas}$-lexA$_{bs94}$-PR-VP16-T$_{ENO1}$ | ColE1 | Chl | Fig.3h |
| LXR404 | $P_{xyluas}$-lexA$_{bs94}$-GR$_{487\text{-}777}$-VP16-T$_{ENO1}$ | ColE1 | Chl | Fig.3h |
| LXR405 | $P_{xyluas}$-lexA$_{bs94}$-MR$_{669\text{-}984}$-VP16-T$_{ENO1}$ | ColE1 | Chl | Fig.3h |
| LXR348 | $P_{S.\ a0b1c0\_lexOrec.sym}$-yEmCitrine-T$_{ENO}$ | ColE1 | Chl | Fig.3h |
| LXR349 | $P_{S.\ a0b1c0\_lexObs}$-yEmCitrine-T$_{ENO1}$ | ColE1 | Chl | Fig.3h |
| LXR353 | $P_{S.\ a0b1c0\_lexOmm}$-yEmCitrine-T$_{ENO1}$ | ColE1 | Chl | Fig.3h |
| LXR355 | $P_{S.\ a0b1c0\_OHKCI}$-yEmCitrine-T$_{ENO1}$ | ColE1 | Chl | Fig.3h |
| LXR387 | $P_{S.\ a0b1c0\_CIOL1}$-yEmCitrine-T$_{ENO1}$ | ColE1 | Chl | Fig.3h |
| LXR325 | $P_{xyluas}$-lexA$_{bs94}$-acVHH -VP16-T$_{ENO1}$ | ColE1 | Chl | Fig.4b |
| LXR326 | $P_{xyluas}$-lexA$_{bs94}$-LasR$_{177}$ nls-VP16-T$_{ENO1}$ | ColE1 | Chl | Fig.4b |
| LXR327 | $P_{xyluas}$-lexA$_{bs94}$-RpaR$_{179}$ nls-VP16-T$_{ENO1}$ | ColE1 | Chl | Fig.4b-c |
| LXR328 | $P_{xyluas}$-lexA$_{bs94}$-CinR$_{179}$ nls -VP16 | ColE1 | Chl | Fig.4b |
| LXR329 | $P_{xyluas}$-lexA$_{bs94}$-BjaR$_{180}$ nls-VP16-T$_{ENO1}$ | ColE1 | Chl | Fig.4b |
| LXR334 | $P_{xyluas}$-lexA$_{ec87}$-CarRecc$_{169}$ nls-VP16-T$_{ENO1}$ | ColE1 | Chl | Fig.4b |
| LXR371 | $P_{xyluas}$-lexA$_{bs94}$-TraR$_{174}$ nls-VP16-T$_{ENO1}$ | ColE1 | Chl | Fig.4b |
| LXR384 | $P_{xyluas}$-CI$_{94}$-SmaR$_{179}$-VP16 | ColE1 | Chl | Fig.4b |
| LXR436 | $P_{xyluas}$-lexA$_{bs94}$-ER$_{282\text{-}595}$-VP16 | ColE1 | Chl | Fig.4b |
| LXR437 | $P_{xyluas}$-lexA$_{bs94}$-DHBR$_{282\text{-}595}$-VP16 | ColE1 | Chl | Fig.4b |
| LXR402 | $P_{xyluas}$-lexA$_{bs94}$-PR-VP16 | ColE1 | Chl | Fig.4b |
| LXR404 | $P_{xyluas}$-lexA$_{bs94}$-GR$_{487\text{-}777}$-VP16 | ColE1 | Chl | Fig.4b |
| LXR405 | $P_{xyluas}$-lexA$_{bs94}$-MR$_{669\text{-}984}$-VP16 | ColE1 | Chl | Fig.4b |
| ZY404 | $P_{xyluas}$- PurR$_{60}$- BjaR$_{180}^{S107R}$ nls -VP16 | ColE1 | Chl | Fig.4b |
| LXR335 | $P_{xyluas}$-CI434$_{70}$-RpaR$_{179}$ nls-VP16 | ColE1 | Chl | Fig.4b-c |
| LXR327 | $P_{xyluas}$-lexA$_{bs94}$-RpaR$_{179}$ nls-VP16 | ColE1 | Chl | Fig.4b-c |
| LXR337 | $P_{xyluas}$-lexA$_{mm115}$-RpaR$_{179}$ nls-VP16 | ColE1 | Chl | Fig.4b-c |
| LXR338 | $P_{xyluas}$-lexA$_{ec87}$-RpaR$_{179}$ nls-VP16 | ColE1 | Chl | Fig.4b-c |
| LXR339 | $P_{xyluas}$-lexA$_{xa101}$-RpaR$_{179}$ nls-VP16 | ColE1 | Chl | Fig.4b-c |
| LXR340 | $P_{xyluas}$-HKCI$_{84}$-RpaR$_{179}$ nls-VP16 | ColE1 | Chl | Fig.4b-c |
| LXR341 | $P_{xyluas}$-RecApact$_{104}$-RpaR$_{179}$ nls-VP16 | ColE1 | Chl | Fig.4b-c |
| LXR342 | $P_{xyluas}$-DeoR$_{92}$-RpaR$_{179}$ nls-VP16 | ColE1 | Chl | Fig.4b-c |
| LXR343 | $P_{xyluas}$-PurR$_{60}$-RpaR$_{179}$ nls-VP16 | ColE1 | Chl | Fig.4b-c |
| LXR344 | $P_{xyluas}$-CI$_{94}$-RpaR$_{179}$ nls-VP16 | ColE1 | Chl | Fig.4b-c |
| LXR345 | $P_{xyluas}$-lexA$_{gs91}$- RpaR$_{179}$ nls-VP16 | ColE1 | Chl | Fig.4b-c |
| LXR346 | $P_{xyluas}$-lexA$_{fs104}$- RpaR$_{179}$ nls-VP16 | ColE1 | Chl | Fig.4b-c |
| LXR373 | $P_{xyluas}$-lexA$_{xa101}$-LasR$_{177}$ nls-VP16 | ColE1 | Chl | Fig.4b |
| LXR374 | $P_{xyluas}$-lexA$_{fs104}$-LasR$_{177}$ nls-VP16 | ColE1 | Chl | Fig.4b |
| LXR375 | $P_{xyluas}$-RecApact$_{104}$-DHBR$_{282\text{-}595}$ nls-VP16 | ColE1 | Chl | Fig.4b |
| LXR376 | $P_{xyluas}$-lexA$_{gs91}$-DHBR$_{282\text{-}595}$ nls-VP16 | ColE1 | Chl | Fig.4b |
| LXR377 | $P_{xyluas}$-PurR$_{60}$-ER$_{282\text{-}595}$ nls-VP16 | ColE1 | Chl | Fig.4b |
| LXR378 | $P_{xyluas}$-DeoR$_{92}$-ER$_{282\text{-}595}$ nls-VP16 | ColE1 | Chl | Fig.4b |
| LXR379 | CI434$_{70}$-BjaR$_{180}$ nls-VP16 | ColE1 | Chl | Fig.4b |
| LXR380 | $P_{xyluas}$-lexA$_{xa101}$-BjaR$_{180}$ nls-VP16 | ColE1 | Chl | Fig.4b |
| LXR381 | $P_{xyluas}$-lexA$_{mm115}$-CinR$_{179}$ nls-VP16-T$_{ENO1}$ | ColE1 | Chl | Fig.4b |
| LXR383 | CI434$_{70}$-SmaR$_{179}$ nls-VP16 | ColE1 | Chl | Fig.4b |
| LXR384 | $P_{xyluas}$-CI$_{94}$-SmaR$_{179}$ nls-VP16-T$_{ENO1}$ | ColE1 | Chl | Fig.4b |
| LXR385 | $P_{xyluas}$-lexA$_{mm115}$-acVHH nls-VP16 | ColE1 | Chl | Fig.4b |
| LXR386 | $P_{xyluas}$-HKCI$_{84}$-acVHH-VP16-T$_{ENO1}$ | ColE1 | Chl | Fig.4b |

| | | | | |
|---|---|---|---|---|
| LXR421 | $P_{xyluas}$-HKCI$_{84}$-LasR$_{177}$ nls-VP16 | ColE1 | Chl | Fig.4b |
| LXR447 | $P_{xyluas}$-PurR$_{60}$-BjaR$_{180}$ nls-VP16 | ColE1 | Chl | Fig.4b |
| LXR423 | $P_{xyluas}$-lexA$_{gs91}$-acVHH nls-VP16 | ColE1 | Chl | Fig.4b |
| LXR402 | $P_{xyluas}$-lexA$_{bs94}$-PR-VP16-T$_{ENO1}$ | ColE1 | Chl | Fig.4b |
| LXR404 | $P_{xyluas}$-lexA$_{bs94}$-GR$_{487\text{-}777}$-VP16-T$_{ENO1}$ | ColE1 | Chl | Fig.4b |
| LXR405 | $P_{xyluas}$-lexA$_{bs94}$-MR$_{669\text{-}984}$-VP16-T$_{ENO1}$ | ColE1 | Chl | Fig.4b |
| LXR325 | $P_{xyluas}$-lexA$_{bs94}$-acVHH -VP16-T$_{ENO1}$ | ColE1 | Chl | Fig.4d |
| LXR385 | $P_{xyluas}$-lexA$_{mm115}$-acVHH -VP16-T$_{ENO1}$ | ColE1 | Chl | Fig.4d |
| LXR349 | $P_{S.\ a0b1c0\_lexObs}$-yEmCitrine-T$_{ENO1}$ | ColE1 | Chl | Fig.4d |
| LXR353 | $P_{S.\ a0b1c0\_lexOmm}$-yEmCitrine-T$_{ENO1}$ | ColE1 | Chl | Fig.4d |
| LXR433 | $P_{xyluas}$-lexA$_{bs94}$-ER$_{282\text{-}595}$-VP16-T$_{ENO1}$ | ColE1 | Chl | Fig.4e |
| LXR436 | $P_{xyluas}$-DeoR$_{92}$-ER$_{282\text{-}595}$-VP16-T$_{ENO1}$ | ColE1 | Chl | Fig.4e |
| LXR349 | $P_{S.\ a0b1c0\_lexObs}$-yEmCitrine-T$_{ENO1}$ | ColE1 | Chl | Fig.4e |
| LXR357 | $P_{S.\ a0b1c0\_ODeo}$-yEmCitrine-T$_{ENO1}$ | ColE1 | Chl | Fig.4e |
| LXR326 | $P_{xyluas}$-lexA$_{bs94}$-LasR$_{177}$ nls-VP16-T$_{ENO1}$ | ColE1 | Chl | Fig.5c |
| LXR373 | $P_{xyluas}$-lexA$_{xa101}$-LasR$_{177}$ nls-VP16 | ColE1 | Chl | Fig.5c |
| LXR374 | $P_{xyluas}$-lexA$_{fs104}$-LasR$_{177}$ nls-VP16 | ColE1 | Chl | Fig.5c |
| LXR349 | $P_{S.\ a0b1c0\_lexObs}$-yEmCitrine-T$_{ENO1}$ | ColE1 | Chl | Fig.5c |
| LXR354 | $P_{S.\ a0b1c0\_lexOxa}$-yEmCitrine-T$_{ENO1}$ | ColE1 | Chl | Fig.5c |
| LXR361 | $P_{S.\ a0b1c0\_lexOfs}$-yEmCitrine-T$_{ENO1}$ | ColE1 | Chl | Fig.5c |
| WR101 | $P_{xyluas}$-CI$_{94}$-LasR$_{177}$ nls-T$_{ENO1}$ | ColE1 | Chl | Fig.5d-e |
| WR102 | $P_{xyluas}$-CI434$_{70}$-LasR$_{177}$ nls-T$_{ENO1}$ | ColE1 | Chl | Fig.5d-e |
| WR119 | $P_{xyluas}$-lexA$_{gs91}$-LasR$_{177}$ nls-T$_{ENO1}$ | ColE1 | Chl | Fig.5d-e |
| WR121 | $P_{xyluas}$-lexA$_{bs94}$-LasR$_{177}$ nls-T$_{ENO1}$ | ColE1 | Chl | Fig.5d-e |
| WR126 | $P_{lexObsi}$-yEmCitrine-T$_{ENO1}$ | ColE1 | Chl | Fig.5d-e |
| WR127 | $P_{lexOgsi}$-yEmCitrine-T$_{ENO1}$ | ColE1 | Chl | Fig.5d-e |
| CY662 | $P_{cI2.1}$-yEmCitrine-T$_{ENO1}$ | ColE1 | Chl | Fig.5d-e |
| CY952 | $P_{CI434.1}$-yEmCitrine-T$_{ENO1}$ | ColE1 | Chl | Fig.5d-e |
| LXR414 | $P_{xyluas}$-PurR$_{60}$-ER$_{282\text{-}595}$-VP16-T$_{ENO1}$ | ColE1 | Chl | Fig.5i |
| LXR417 | $P_{xyluas}$-PurR$_{60}$-ER$_{282\text{-}595}$-VP16-T$_{ENO1}$ | ColE1 | Chl | Fig.5i |
| LXR419 | $P_{xyluas}$-LexA$_{ec87}$-RpaR$_{179}$ nls-VP16-T$_{ENO1}$ | ColE1 | Chl | Fig.5i |
| LXR411 | $P_{S.\ a0b1c0\_PurRa\text{-}lexObsi}$-yEmCitrine-T$_{ENO1}$ | ColE1 | Chl | Fig.5i |
| LXR415 | $P_{S.\ a0b1c0\_PurRa\text{-}lexOxai}$-yEmCitrine-T$_{ENO1}$ | ColE1 | Chl | Fig.5i |
| LXR420 | $P_{S.\ a0b1c0\_lexOrec.syma\text{-}lexObsi}$-yEmCitrine-T$_{ENO1}$ | ColE1 | Chl | Fig.5i |
| ZY933 | ytetR$^{NLS}$, XlnR, ylacI$^{NLS}$ | ColE1 | Chl | Fig.7c[9] |
| ZY114 | P$_{RPL8B}$-HKCI$_{84}$-LasR$_{177}$ nls-VP16_opt1-T$_{RPS9A}$ | ColE1 | Kan | Fig.7c |
| ZY213 | P$_{RPL4A}$-lexA$_{xa101}$-RpaR$_{179}$ nls-VP16_opt2-T$_{ENO1}$ | ColE1 | Kan | Fig.7c |
| ZY476 | P$_{EFT2}$-lexA$_{ec87}$-ER$_{282\text{-}595}$-VP16_opt3-T$_{CYC1}$ | ColE1 | Kan | Fig.7c |
| ZY517 | P$_{RPS3}$-lexA$_{mm115}$-DHBR$_{282\text{-}595}$-VP16_opt4- | ColE1 | Kan | Fig.7c |
| ZY148 | P$_{HTA2}$-lexA$_{gs91}$-acVHH nls-VP16_opt5-T$_{ADH2.}$ | ColE1 | Kan | Fig.7c |
| ZY395 | P$_{ZUO1}$-CI434$_{70}$-CinR$_{179}$ nls-VP16_opt6 nls-T$_{DPP1}$ | ColE1 | Kan | Fig.7c |
| ZY477 | P$_{RPL28}$-CI$_{94}$-PR-VP16_opt7-T$_{AQR1}$ | ColE1 | Kan | Fig.7c |
| ZY525 | ZY275 ZY114 ZY213 ZY476 ZY517 ZY148 | ColE1 | Chl | Fig.7c |
| ZY428 | P$_{SSB1}$-PurR$_{60}$-BjaR$_{180}^{S107R}$ nls-VP16_opt8- | ColE1 | Kan | Fig.7c |
| ZY602 | P$_{FBA1}$-lexA$_{bs94}$ -LuxR$_{183}^{S116Y}$ nls-VP16_opt9- | ColE1 | Kan | Fig.7c |
| ZY604 | ZY522 ZY428 ZY602 | ColE1 | Chl | Fig.7c |
| gd281 | $P_{tetuas}$-yEmCitrine-T$_{TEF1}$ | ColE1 | Chl | Fig.7c |
| m115 | $P_{xlnO\ m115}$-yEmCitrine-T$_{ENO1}$ | ColE1 | Chl | Fig.7c |
| gd373 | $P_{lacuas}$-yEmCitrine-T$_{ENO1}$ | ColE1 | Chl | Fig.7c |

| | | | | |
|---|---|---|---|---|
| ZY615 | $P_{S.\ a0b1c0\_OHKCI}$-yEmCitrine-$T_{ENO1}$ | ColE1 | Chl | Fig.7c |
| ZY614 | $P_{S.\ a0b1c0\_lexOxa}$-yEmCitrine-$T_{ENO1}$ | ColE1 | Chl | Fig.7c |
| ZY611 | $P_{S.\ a0b1c0\_lexOrec.sym}$-yEmCitrine-$T_{ENO1}$ | ColE1 | Chl | Fig.7c |
| ZY613 | $P_{S.\ a0b1c0\_lexOmm}$-yEmCitrine-$T_{ENO1}$ | ColE1 | Chl | Fig.7c |
| ZY620 | $P_{S.\ a0b1c0\_lexOgs}$-yEmCitrine-$T_{ENO1}$ | ColE1 | Chl | Fig.7c |
| ZY612 | $P_{S.\ a0b1c0\_OCI434}$-yEmCitrine-$T_{ENO1}$ | ColE1 | Chl | Fig.7c |
| ZY622 | $P_{S.\ a0b1c0\_OCI\ OL1}$-yEmCitrine-$T_{ENO1}$ | ColE1 | Chl | Fig.7c |
| ZY618 | $P_{S.\ a0b1c0\_purO}$-yEmCitrine-$T_{ENO1}$ | ColE1 | Chl | Fig.7c |
| ZY753 | $P_{S.\ a0b1c0\_lexObs}$-yEmCitrine-$T_{ENO1}$ | ColE1 | Chl | Fig.7c |
| ZY134 | $P_{S.\ a14b5c21\_OHKCI}$-yEmCitrine-$T_{ENO1}$ | ColE1 | Chl | Fig.7c |
| ZY226 | $P_{S.\ a22b26c22\_lexOxa}$-yEmCitrine-$T_{ENO1}$ | ColE1 | Chl | Fig.7d, Fig.S15 |
| ZY495 | $P_{S.\ a20b25c6\_lexOrec.sym}$-yEmCitrine-$T_{ENO1}$ | ColE1 | Chl | Fig.7d, Fig.S15 |
| ZY573 | $P_{S.\ a17b1c12\_lexOmm}$-yEmCitrine-$T_{ENO1}$ | ColE1 | Chl | Fig.7d, Fig.S15 |
| ZY138 | $P_{S.\ a24b30c13\_lexOgs}$-yEmCitrine-$T_{ENO1}$ | ColE1 | Chl | Fig.7d, Fig.S15 |
| ZY135 | $P_{S.\ a12b17c24\_OCI434}$-yEmCitrine-$T_{ENO1}$ | ColE1 | Chl | Fig.7d, Fig.S15 |
| ZY494 | $P_{S.\ a22b1c15\_OCI\ ol1}$-yEmCitrine-$T_{ENO1}$ | ColE1 | Chl | Fig.7d, Fig.S15 |
| ZY139 | $P_{S.\ a18b23c14\_purO}$-yEmCitrine-$T_{ENO1}$ | ColE1 | Chl | Fig.7d, Fig.S15 |
| ZY675 | $P_{S.\ a9b27c23\_2\times lexObs}$-yEmCitrine-$T_{ENO1}$ | ColE1 | Chl | Fig.7d, Fig.S15 |
| ZY391 | CY571 $P_{lacuas}$-TcpanD-$T_{RPL41B}$ | ColE1 | Chl | Fig.7d-g |
| ZY437 | CY700 $P_{S.\ a12b17c24\_OCI434}$-CgpanC-$T_{GAT2t}$ | ColE1 | Chl | Fig.7d-g |
| ZY589 | CY701 $P_{S.\ a17b1c12\_lexOmm}$-Scecm31-$T_{PDC6t}$ | ColE1 | Chl | Fig.7d-g |
| ZY511 | CY702 $P_{S.\ a20b25c6\_lexOrec.sym}$-Scilv6-$T_{PFY1}$ | ColE1 | Kan | Fig.7d-g |
| ZY512 | CY703 $P_{S.\ a22b1c15\_OCI\ OL1}$-PailvC-$T_{YJR085Ct}$ | ColE1 | Kan | Fig.7d-g |
| ZY224 | CY704 $P_{S.\ a22b26c22\_lexOxa}$-MtpanE-$T_{PRM5t}$ | ColE1 | Kan | Fig.7d-g |
| ZY441 | CY705 $P_{S.\ a14b5c21\_OHKCI}$-Scilv2-$T_{IDP1t}$ | ColE1 | Kan | Fig.7d-g |
| ZY442 | CYx212 $P_{m115\_xlnO}$-BsilvD-$T_{RPL15A}$ | ColE1 | Kan | Fig.7d-g |
| ZY600 | CY1808 ZY391 ZY437 ZY589 ZY511 ZY512 | ColE1 | Chl | Fig.7d-g |

#### Supplementary Table 8: List of Strains

| Strain | Genotype | Selection |
|---|---|---|
| Cye72 | BY4741 *xylR natMX tetR lacI* sensors strain. | [2] |
| BY4741 12s | BY4741 ZY933 int ZY525 int ZY604int | NAT |
| CEN.PK2-1C 12s | CEN.PK2-1C ZY933 int ZY525 int ZY604 int | NAT |
| CEN.PK2-1C 10s | CEN.PK2-1C ZY933 int ZY525 int | G418 |
| CEN.PK2-1C 10s M | CEN.PK2-1C 10s ZY600 int | G418 & Ura |

**Supplementary Table 9: List of promoter sequences tested in this study.**

| Promoter | Sequence (underlined is -35 and -10 region) |
| --- | --- |
| $P_{lexOec}$ | AAATTTATCAAAAAGAGTGTTGAC*tactgtatatatatacagta*TAATTAGATTCAA*tactgtatatatatacagtaT* |
| $P_{lexOmt}$ | CAATCACCTATGAACTGTCGA*TTGACtatcgaacacatgttcgaga*TACTTAGATTCAAT |
| $P_{lexOcg}$ | CAATCACCTATGAACTGTCGATTGACTTCTTGCCATCGACTACGGATACTGAGCAC*ctcgaacactc gtaccattt* |
| $P_{lexOsa}$ | CAATCACCTATGAACTGTCGATTGACTTCTTGCCATCGACTACGGATACTGAGCAC*acgaagaaat gtttgttttg* |
| $P_{lexObs}$ | CAATCACCTATGAACTGTCGATTGACTTTT*agaacatatgttcg*GATACTTAGATTCAAT |
| $P_{lexOlm}$ | CAATCACCTATGAACTGTCGATTGAC*tataagaacgtatgttcggata*CTTAGATTCAAT |
| $P_{lexOpd}$ | CAATCACCTATGAACTGTCGATTGACT*caggaactaatgttctg*GATACTTAGATTCAAT |
| $P_{lexOsm}$ | CAATCACCTATGAACTGTCGATTGA*ctgttcttgattgaacata*GATACTTAGATTCAAT |
| $P_{lexOcc}$ | CAATCACCTATGAACTGTCGATTGACTTCTTGCCATCGACTACGGATACTGAGCAC*tatgttctcgtta gaacatc* |
| $P_{lexOpa}$ | CAATCACCTATGAACTGTCGATTGA*cgctgtataaatatacagtg*ATACTTAGATTCAAT |
| $P_{lexOvp}$ | CAATCACCTATGAACTGTCGATTGA*taactgtataaataaacaggat*ACTTAGATTCAAT |
| $P_{lexOpp}$ | CAATCACCTATGAACTGTCGAT*tgactgtatatatatacagtat*GATACTTAGATTCAAT |
| $P_{CI}$ | CAATCACCTATGAACTGTCGATTGACTTCTTGCCATCGACTACGGATACTGAGCACG*taccactgg cggtgata*TC |
| $P_{CI434}$ | CAATCACCTATGAACTGTCGATTGACTTCTTGCCATCGACTACGGATACTGAGCACG*tacaagaa agtttgtt*ATC |
| $P_{HKCI}$ | CAATCACCTATGAACTGTCGATTGACTTCTTGCCATCGACTACGGATACTGAGCACG*tgaaccata agttcaG*ATC |
| $P_{purO}$ | CAATCACCTATGAACTGTCGATTGACAT*acgcaaacgtttgcgt*GATACTTAGATTCAAT |
| $P_{Deo}$ | CAATCACCTATGAACTGTCGATTGACTTCTTGCCATCGACTACGGATACTGAGCACG*tgttagaatt ctaaca*ATC |
| $P_{lexOac}$ | CAATCACCTATGAACTGTCGATTGACTTCTTGCCATCGACTACGGATACTGAGCAC*gcgaagaaa aggcgaacaac* |
| $P_{lexOfs}$ | CAATCACCTATGAACTGTCGATTGACTTCTTGCCATCGACTACGGATACTGAGCAC*acctgcacaa aggtgcacTG* |
| $P_{lexOgs}$ | CAATCACCTATGAACTGTCGATTGACTTCTTGCCATCGACTACGGATACTGAGCAC*aggttgacac atgtcaacct* |
| $P_{lexOmm}$ | CAATCACCTATGAACTGTCGATTGACTTCTTGCCATCGACTACGGATACTGAGCAC*gacctaatatt aataaggtc* |
| $P_{lexOmx}$ | CAATCACCTATGAACTGTCGATTGACTTCTTGCCATCGACTACGGATACTGAGCAC*gctacaagcc cgttcaggac* |
| $P_{lexOpm}$ | CAATCACCTATGAACTGTCGATTGACTTCTTGCCATCGACTACGGATACTGAGCAC*ggaatatataa gaataacac* |
| $P_{lexOsp}$ | CAATCACCTATGAACTGTCGATTGACTTCTTGCCATCGACTACGGATACTGAGCAC*gctagtcctag agtcctaac* |
| $P_{lexOxa101}$ | CAATCACCTATGAACTGTCGATTGACTTCTTGCCATCGACTACGGATACTGAGCAC*gttagtagtaa tactactaa* |
| $P_{lexOxa90}$ | CAATCACCTATGAACTGTCGATTGACTTCTTGCCATCGACTACGGATACTGAGCAC*gcatgtacaa atgtacaacg* |
| $P_{lexOxf}$ | CAATCACCTATGAACTGTCGATTGACTTCTTGCCATCGACTACGGATACTGAGCAC*gcttagtaata atactaacg* |
| $P_{RecApact}$ | CAATCACCTATGAACTGTCGATTGACTTCTTGCCATCGACTACGGATACTGAGCACG*tactaattaa tttagt*AGC |
| $P_{Murepc}$ | CAATCACCTATGAACTGTCGATTGACTTCTTGCCATCGACTACGGATACTGAGCAC*cttttcagtatta ttcttttctataaagt* |
| $P_{RutR}$ | CAATCACCTATGAACTGTCGATTGACTTCTTGCCATCGACTACGGATACTGAGCACGC*ttgaccat ttggtcaa*TC |
| $P_{BetI}$ | CAATCACCTATGAACTGTCGATTGACTTCTTGCCATCGACTACGGATACTGAGCACGTC*attgga cgttcaat*ATC |
| $P_{ArgP}$ | CAATCACCTATGAACTGTCGATTGACTTCTTGCCATCGACTACGGATACTGAGCAC*cttattagttttt ctgattgcc* |
| $P_{FruR}$ | CAATCACCTATGAACTGTCGATTGACTTCTTGCCATCGACTACGGATACTGAGCAC*ttcttgaaacgt ttcagcgc* |
| $P_{RbsR}$ | CAATCACCTATGAACTGTCGATTGACTTCTTGCCATCGACTACGGATACTGAGCAC*tcagcgaaac gtttcgctga* |
| $P_{TreR}$ | CAATCACCTATGAACTGTCGATTGACTTCTTGCCATCGACTACGGATACTGAGCACTTT*cgggaa cgttcccg*TTT |
| $P_{NanR}$ | CAATCACCTATGAACTGTCGATTGACTTCTTGCCATCGACTACGGATACTGAGCAC*ggtataacag gtataaaggtata* |
| $P_{PdhR}$ | CAATCACCTATGAACTGTCGATTGACTTCTTGCCATCGACTACGGATACTGAGCACTC*attggttat* |

| | |
|---|---|
| | *accaat*TGC |
| $P_{DgoR}$ | CAATCACCTATGAACTGTCGATTGACTTCTTGCCATCGACTACGGATACTGAGCACctaaattgtagtacaacaat |
| $P_{AllR}$ | CAATCACCTATGAACTGTCGATTGACTTCTTGCCATCGACTACGGATACTGAGCACAG*ttggaaaaattttccaa*C |
| $P_{MhpR}$ | CAATCACCTATGAACTGTCGATTGACTTCTTGCCATCGACTACGGATACTGAGCAC*accacgtggaccacgtgtaag* |
| *xlnO* | *gaatttaggctaaagaaagatc* |
| *polyA* | **AATGAAAAAAAAAAAAAAAAAAAA** |
| *TATA box* | **TATAAATA** |
| *TSS* | **AGAATATCAAGCTACAAAAA** |
| $P_{xyluas}$ | CAACGGCCTAGCATGTGATTAATTAATTATTTGTTTTTTTTTGCAGTATAAAAAGTTAGTTTGTTTAAACAACAAACTTTTTTCATTTCTTTTGTTTCCCCTTCTCTTCTTTTAGTTAGTTTGTTTAAACAACAAACT**AGAATATCAAGCTACAAAAA**TAAATAAAA |
| $P_{tet}$ | TTTGTTTTTATTTTTTGCGGTATAAAAATCCCTATCAGTGATAGAGATTTTCTTTTTTTTTTATTTTTCCCTATCAGTGATAGAGAATATCAAGCTACAAAAATAAATAAAA |
| $P_{tetuas}$ | CAACGGCCTAGCATGTGATTAATTAATTTTGTTTTTTATTTTTTGCGGTATAAAAATCCCTATCAGTGATAGAGATTTTCTTTTTTTTTTATTTTTCCCTATCAGTGATAGAGAATATCAAGCTACAAAAATAAATAAAA |
| $P_{lacuas}$ | CCTCCTTGAAACTGAAATTTTAGCATGTGATTAATTAATTTAATTGTGAGCGGATAACAATATAAAAAGAGTGCCAGTAGCGACTTGTTTCACACTCGAATTGTGAGCGGATAACAATAGAATATCAAGCTACAAAAATAAATAAAA |
| $P_{S.\ a0b0c0}$ | TCTCTCCGGTTACAGCCTGTGTAACTGATTAATCCTGCCTTTCTAATCACCATTCTAATGTTTTAATTAAGGGATTTTGTCTTCATTAACGGCTTTCGCTCATAAAAATGTTATGACGTTTTGCCCGCAGGCGGGAAACCATCCACTTCACGAGACTGATCTCCTCTGCCGGAACACCGGGCATCTCCAACTTATAAGTTGGAGAAATAAGAGAATTTCAGATTGAGAG**AATGAAAAAAAAAAAAAAAAAAAA**GGCAGAGGAGAGCATAGAAATGGGGTTCACTTTTTGGTAAAGCTATAGCATGCCTATCACA**TATAAATA**GAGTGCCAGTAGCGACTTTTTTCACACTCGAAATACTCTTACTACTGCTCTCTTGTTGTTTTTATCACTTCTTGTTTCTTCTTGGTAAAT**AGAATATCAAGCTACAAAAA**GCATACAATCAACTATCAACTATTAACTATATCGTAATACACA |
| $P_{S.\ a0b0c0(adr1)2m}$ | ATTAACGGCTTTCGCTCATAAAAATGTTATGACGTTTTGCCCGCAGGCGGGAAACCATCCACTTCACGAGACTGATCTCCTCTGCCGGAACACCGGGCATTGATCGCTTATAAGCGATCAAAATAAGAGAATTTCAGATTGAGAG**AATGAAAAAAAAAAAAAAAAAAAA**GGCAGAGGAGAGCATAGAAATGGGGTTCACTTTTTGGTAAAGCTATAGCATGCCTATCACA**TATAAATA**GAGTGCCAGTAGCGACTTTTTTCACACTCGAAATACTCTTACTACTGCTCTCTTGTTGTTTTTATCACTTCTTGTTTCTTCTTGGTAAAT**AGAATATCAAGCTACAAAAA**GCATACAATCAACTATCAACTATTAACTATATCGTAATACACA |
| $P_{S.\ a0b0c0\Delta mig1}$ | ATTAACGGCTTTCGCTCATAAAAATGTTATGACGTTTTGCCCGCAGGCGGGAAACCATCCACTTCACGAGACTGATCTCCTCTGCCGGAACACCGGGCATCTCCAACTTATAAGTTGGAGAAATAAGAGAATTTCAGATTGAGAG**AATGAAAAAAAAAAAAAAAAAAAA**GGCA**TATAAATA**GAGTGCCAGTAGCGACTTTTTTCACACTCGAAATACTCTTACTACTGCTCTCTTGTTGTTTTTATCACTTCTTGTTTCTTCTTGGTAAAT**AGAATATCAAGCTACAAAAA**GCATACAATCAACTATCAACTATTAACTATATCGTAATACACA |
| $P_{S.\ a0b0c0mig1m}$ | ATTAACGGCTTTCGCTCATAAAAATGTTATGACGTTTTGCCCGCAGGCGGGAAACCATCCACTTCACGAGACTGATCTCCTCTGCCGGAACACCGGGCATCTCCAACTTATAAGTTGGAGAAATAAGAGAATTTCAGATTGAGAG**AATGAAAAAAAAAAAAAAAAAAAA**GGCAGAGGAGAGCATAGAATGAGCTTTCACTTTTTGGTAAAGCTATAGCATGCCTATCACA**TATAAATA**GAGTGCCAGTAGCGACTTTTTTCACACTCGAAATACTCTTACTACTGCTCTCTTGTTGTTTTTATCACTTCTTGTTTCTTCTTGGTAAAT**AGAATATCAAGCTACAAAAA**GCATACAATCAACTATCAACTATTAACTATATCGTAATACACA |
| $P_{S.\ a0b0c0(adr1)1m}$ | ATTAACGGCTTTCGCTCATAAAAATGTTATGACGTTTTGCCCGCAGGCGGGAAACCATCCACTTCACGAGACTGATTGCTTCTGCCGGAACACCGGGCATCTCCAACTTATAAGTTGGAGAAATAAGAGAATTTCAGATTGAGAG**AATGAAAAAAAAAAAAAAAAAAAA**GGCAGAGGAGAGCATAGAAATGGGGTTCACTTTTTGGTAAAGCTATAGCATGCCTATCACA**TATAAATA**GAGTGCCAGTAGCGACTTTTTTCACACTCGAAATACTCTTACTACTGCTCTCTTGTTGTTTTTATCACTTCTTGTTTCTTCTTGGTAAAT**AGAATATCAAGCTACAAAAA**GCATACAATCAACTATCAACTATTAACTATATCGTAATACACA |
| $P_{S.\ a0b0c0\Delta 89}$ | ACACCGGGCATCTCCAACTTATAAGTTGGAGAAATAAGAGAATTTCAGATTGAGAG**AATGAAAAAAAAAAAAAAAAAAAA**GGCAGAGGAGAGCATAGAAATGGGGTTCACTTTTTGGTAAAGCTATAGCATGCCTATCACA**TATAAATA**GAGTGCCAGTAGCGACTTTTTTCACACTCGAAATACTCTTACTACTGCTCTCTTGTTGTTTTTATCACTTCTTGTTTCTTCTTGGTAAAT**AGAATATCAAGCTACAAAAA**GCATACAATCAACTATCAACTATTAACTATATCGTAATACACA |
| $P_{S.\ a0b0c0\Delta 89k}$ | ACACCGGGCATCTCCAACTTATAAGTTGGAGAAATAAGAGAATTTCAGATTGAGAG**AATGAAAAAAAAAAAAAAAAAAAA**GGCAGAGGAGAGCATAGAAATGGGGTTCACTTTTTGGTAAAGCTATAGCATGCCTATCACA**TATAAATA**GAGTGCCAGTAGCGACTTTTTTCACACTCGAAATACTCTTACTACTGCTCTCTTGTTGTTTTTATCACTTCTTGTTTCTTCTTGGTAAAT**AGAATATCAAGCTACAAAAA**TAAATAAAAA |

| | |
|---|---|
| $P_{S.\ a0b0c0\Delta 89k\_gal}$ | ACACCGGGCACGCGCCGCACTGCTCCGAACAATAATAAGAGAATTTCAGATTGAGAG**AATGAAAAAAAAAAAAAAAAAAAA**GGCAGAGGAGAGCATAGAAATGGGGTTCACTTTTTGGTAAAGCTATAGCATGCCTATCACA**TATAAATA**GAGTGCCAGTAGCGACTTTTTTCACACTCGAAATACTCTTACTACTGCTCTCTTGTTGTTTTTATCACTTCTTGTTTCTTCTTGGTAAAT**AGAATATCAAGCTACAAAAA**TAAATAAAAA |
| $P_{S.\ a0b0c0\Delta 89k\_4\times kb}$ | ACACCGGGCAGGGACTTTCCGCTGGGACTTTCCAGGGACTTTCCGCTGGGACTTTCCAATAAGAGAATTTCAGATTGAGAG**AATGAAAAAAAAAAAAAAAAAAAA**GGCAGAGGAGAGCATAGAAATGGGGTTCACTTTTTGGTAAAGCTATAGCATGCCTATCACA**TATAAATA**GAGTGCCAGTAGCGACTTTTTTCACACTCGAAATACTCTTACTACTGCTCTCTTGTTGTTTTTATCACTTCTTGTTTCTTCTTGGTAAAT**AGAATATCAAGCTACAAAAA**TAAATAAAAA |
| $P_{4\times rpaO}$ | ACACCGGGCA*acctgtccgatcggacagta*CCGTTAGAGCTTGACGGGGAAAGCCGG*acctgtccgatcggacagta*CGAACGTGGCGAGAAAGGAAGGGAAGA*acctgtccgatcggacagta*AAGCGCTTAGCGATCCTAGGGCGATCA*acctgtccgatcggacagta*AATAAGAGAATTTCAGATTGAGAG**AATGAAAAAAAAAAAAAAAAAAAA**GGCAGAGGAGAGCATAGAAATGGGGTTCACTTTTTGGTAAAGCTATAGCATGCCTATCACA**TATAAATA**GAGTGCCAGTAGCGACTTTTTTCACACTCGAAATACTCTTACTACTGCTCTCTTGTTGTTTTTATCACTTCTTGTTTCTTCTTGGTAAAT**AGAATATCAAGCTACAAAAA**TAAATAAAAA |
| $P_{S.\ a0b0c0\Delta 135k}$ | AGATTGAGAG**AATGAAAAAAAAAAAAAAAAAAAA**GGCAGAGGAGAGCATAGAAATGGGGTTCACTTTTTGGTAAAGCTATAGCATGCCTATCACA**TATAAATA**GAGTGCCAGTAGCGACTTTTTTCACACTCGAAATACTCTTACTACTGCTCTCTTGTTGTTTTTATCACTTCTTGTTTCTTCTTGGTAAAT**AGAATATCAAGCTACAAAAA**TAAATAAAAA |
| $P_{S.\ a0b0c0\Delta 89k\_lexOec}$ | ACACCGGGCATGCTGTATATACTCACAGCAAATAAGAGAATTTCAGATTGAGAG**AATGAAAAAAAAAAAAAAAAAAAA**GGCAGAGGAGAGCATAGAAATGGGGTTCACTTTTTGGTAAAGCTATAGCATGCCTATCACA**TATAAATA**GAGTGCCAGTAGCGACTTTTTTCACACTCGAAATACTCTTACTACTGCTCTCTTGTTGTTTTTATCACTTCTTGTTTCTTCTTGGTAAAT**AGAATATCAAGCTACAAAAA**TAAATAAAAA |
| $P_{S.\ a0b0c0\Delta 89k\_2\times lexOec}$ | ACACCGGGCATGCTGTATATACTCACAGCATAACTGTATATACACCCAGGGTCTAGGTGCTGTATATACTCACAGCAAATAAGAGAATTTCAGATTGAGAG**AATGAAAAAAAAAAAAAAAAAAAA**GGCAGAGGAGAGCATAGAAATGGGGTTCACTTTTTGGTAAAGCTATAGCATGCCTATCACA**TATAAATA**GAGTGCCAGTAGCGACTTTTTTCACACTCGAAATACTCTTACTACTGCTCTCTTGTTGTTTTTATCACTTCTTGTTTCTTCTTGGTAAAT**AGAATATCAAGCTACAAAAA**TAAATAAAAA |
| $P_{S.\ a0b0c0\Delta 89k\_3\times lexOec}$ | ACACCGGGCATGCTGTATATACTCACAGCATAACTGTATATACACCCAGGGTCTAGGTGCTGTATATACTCACAGCATAACTGTATATACACCCAGGGTCTAGGTGCTGTATATACTCACAGCAAATAAGAGAATTTCAGATTGAGAG**AATGAAAAAAAAAAAAAAAAAAAA**GGCAGAGGAGAGCATAGAAATGGGGTTCACTTTTTGGTAAAGCTATAGCATGCCTATCACA**TATAAATA**GAGTGCCAGTAGCGACTTTTTTCACACTCGAAATACTCTTACTACTGCTCTCTTGTTGTTTTTATCACTTCTTGTTTCTTCTTGGTAAAT**AGAATATCAAGCTACAAAAA**TAAATAAAAA |
| $P_{S.\ a0b0c0\Delta 89k\_4\times lexOec}$ | ACACCGGGCATGCTGTATATACTCACAGCATAACTGTATATACACCCAGGGTCTAGGTGCTGTATATACTCACAGCATAACTGTATATACACCCAGGGTCTAGGTGCTGTATATACTCACAGCATAACTGTATATACACCCAGGGTCTAGGTGCTGTATATACTCACAGCAAATAAGAGAATTTCAGATTGAGAG**AATGAAAAAAAAAAAAAAAAAAAA**GGCAGAGGAGAGCATAGAAATGGGGTTCACTTTTTGGTAAAGCTATAGCATGCCTATCACA**TATAAATA**GAGTGCCAGTAGCGACTTTTTTCACACTCGAAATACTCTTACTACTGCTCTCTTGTTGTTTTTATCACTTCTTGTTTCTTCTTGGTAAAT**AGAATATCAAGCTACAAAAA**TAAATAAAAA |
| $P_{S.\ a0b1c0}$ | ACACCGGGCA*gaatttaggctaaagaaagatc*AATAAGAGAATTTCAGATTGAGAG**AATGAAAAAAAAAAAAAAAAAAAA**GGCAGTCTTGGACAATAGTTATATCACGTTGACTCAGGAAGTGGTGGTGGAGACTGCCACA**TATAAATA**GAGTGCCAGTAGCGACTTTTTTCACACTCGAAATACTCTTACTACTGCTCTCTTGTTGTTTTTATCACTTCTTGTTTCTTCTTGGTAAAT**AGAATATCAAGCTACAAAAA**TAAATAAAAA |
| $P_{S.\ a052c0}$ | ACACCGGGCA*gaatttaggctaaagaaagatc*AATAAGAGAATTTCAGATTGAGAG**AATGAAAAAAAAAAAAAAAAAAAA**GGCAGTCTTGGACAATAGTTATATCACGTTGTGGTGGTGGAGACTGCCACA**TATAAATA**GAGTGCCAGTAGCGACTTTTTTCACACTCGAAATACTCTTACTACTGCTCTCTTGTTGTTTTTATCACTTCTTGTTTCTTCTTGGTAAAT**AGAATATCAAGCTACAAAAA**TAAATAAAAA |
| $P_{S.\ a071c0}$ | ACACCGGGCA*gaatttaggctaaagaaagatc*AATAAGAGAATTTCAGATTGAGAG**AATGAAAAAAAAAAAAAAAAAAAA**GGCAGTCTTGGACAATAGTTATATCACGTTGACTCAGAAACGCAGCAGAAGTGGTGGTGGAGACTGCCACA**TATAAATA**GAGTGCCAGTAGCGACTTTTTTCACACTCGAAATACTCTTACTACTGCTCTCTTGTTGTTTTTATCACTTCTTGTTTCTTCTTGGTAAAT**AGAATATCAAGCTACAAAAA**TAAATAAAAA |
| $P_{S.\ a046c0}$ | ACACCGGGCA*gaatttaggctaaagaaagatc*AATAAGAGAATTTCAGATTGAGAG**AATGAAAAAAAAAAAAAAAAAAAA**GGCAGTCTTGGACAATAGTTATATTGTGGTGGTGGAGACTGCCACA**TATAAATA**GAGTGCCAGTAGCGACTTTTTTCACACTCGAAATACTCTTACTACTGCTCTCTTGTTGTTTTTATCACTTCTTGTTTCTTCTTGGTAAAT**AGAATATCAAGCTACAAAAA**TAAATAAAAA |
| $P_{S.\ a041c0}$ | ACACCGGGCA*gaatttaggctaaagaaagatc*AATAAGAGAATTTCAGATTGAGAG**AATGAAAAAAAAAAAAAAAAAAAA**GGCAGTCTTGGACAATAGTTATGGTGGTGGAGACTGCCACA**TATAAATA** |

| | |
|---|---|
| | GAGTGCCAGTAGCGACTTTTTTCACACTCGAAATACTCTTACTACTGCTCTCTTGTTGTTTTT<br>ATCACTTCTTGTTTCTTCTTGGTAAAT**AGAATATCAAGCTACAAAAA**TAAATAAAAA |
| $P_{S.\ a036c0}$ | ACACCGGGCA*gaatttaggctaaagaaagatc*AATAAGAGAATTTCAGATTGAGAG**AATGAAAAAAAA**<br>**AAAAAAAAAAAA**GGCAGTCTTGGACAATAGTTGGTGGAGACTGCCACA**TATAAATA**GAGTG<br>CCAGTAGCGACTTTTTTCACACTCGAAATACTCTTACTACTGCTCTCTTGTTGTTTTTATCACT<br>TCTTGTTTCTTCTTGGTAAAT**AGAATATCAAGCTACAAAAA**TAAATAAAAA |
| $P_{S.\ a031c0}$ | ACACCGGGCA*gaatttaggctaaagaaagatc*AATAAGAGAATTTCAGATTGAGAG**AATGAAAAAAAA**<br>**AAAAAAAAAAAA**GGCAGTCTTGGACAATGTGGAGACTGCCACA**TATAAATA**GAGTGCCAG<br>TAGCGACTTTTTTCACACTCGAAATACTCTTACTACTGCTCTCTTGTTGTTTTTATCACTTCTT<br>GTTTCTTCTTGGTAAAT**AGAATATCAAGCTACAAAAA**TAAATAAAAA |
| $P_{S.\ a026c0}$ | ACACCGGGCA*gaatttaggctaaagaaagatc*AATAAGAGAATTTCAGATTGAGAG**AATGAAAAAAAA**<br>**AAAAAAAAAAAA**GGCAGTCTTGGACGGAGACTGCCACA**TATAAATA**GAGTGCCAGTAGCG<br>ACTTTTTTCACACTCGAAATACTCTTACTACTGCTCTCTTGTTGTTTTTATCACTTCTTGTTTCT<br>TCTTGGTAAAT**AGAATATCAAGCTACAAAAA**TAAATAAAAA |
| $P_{S.\ a016c0}$ | ACACCGGGCA*gaatttaggctaaagaaagatc*AATAAGAGAATTTCAGATTGAGAG**AATGAAAAAAAA**<br>**AAAAAAAAAAAA**GGCAGTCTTTGCCACA**TATAAATA**GAGTGCCAGTAGCGACTTTTTTCAC<br>ACTCGAAATACTCTTACTACTGCTCTCTTGTTGTTTTTATCACTTCTTGTTTCTTCTTGGTAAAT<br>**AGAATATCAAGCTACAAAAA**TAAATAAAAA |
| $P_{S.\ a08c0}$ | ACACCGGGCA*gaatttaggctaaagaaagatc*AATAAGAGAATTTCAGATTGAGAG**AATGAAAAAAAA**<br>**AAAAAAAAAAAA**GGCACACA**TATAAATA**GAGTGCCAGTAGCGACTTTTTTCACACTCGAAA<br>TACTCTTACTACTGCTCTCTTGTTGTTTTTATCACTTCTTGTTTCTTCTTGGTAAAT**AGAATATC**<br>**AAGCTACAAAAA**TAAATAAAAA |
| $P_{S.\ a081c0}$ | ACACCGGGCA*gaatttaggctaaagaaagatc*AATAAGAGAATTTCAGATTGAGAG**AATGAAAAAAAA**<br>**AAAAAAAAAAAA**GGCAGTCTTGGACAAAAACGCAGCAGGCTTGAAACTAGTTATATCACGT<br>TGACTCAGGAAGTGGTGGTGGAGACTGCCACA**TATAAATA**GAGTGCCAGTAGCGACTTTTT<br>TCACACTCGAAATACTCTTACTACTGCTCTCTTGTTGTTTTTATCACTTCTTGTTTCTTCTTGG<br>TAAAT**AGAATATCAAGCTACAAAAA**TAAATAAAAA |
| $P_{S.\ a091c0}$ | ACACCGGGCA*gaatttaggctaaagaaagatc*AATAAGAGAATTTCAGATTGAGAG**AATGAAAAAAAA**<br>**AAAAAAAAAAAA**GGCAGTCTTGGACAAAAACGCAGCAGGCTTGAAACAGAACGTTACTAG<br>TTATATCACGTTGACTCAGGAAGTGGTGGTGGAGACTGCCACA**TATAAATA**GAGTGCCAGTA<br>GCGACTTTTTTCACACTCGAAATACTCTTACTACTGCTCTCTTGTTGTTTTTATCACTTCTTGT<br>TTCTTCTTGGTAAAT**AGAATATCAAGCTACAAAAA**TAAATAAAAA |
| $P_{S.\ a0101c0}$ | ACACCGGGCA*gaatttaggctaaagaaagatc*AATAAGAGAATTTCAGATTGAGAG**AATGAAAAAAAA**<br>**AAAAAAAAAAAA**GGCAGTCTTGGACAAAAACGCAGCAGGCTTGAAACAGAACGTTACGTG<br>AAGTACCTAGTTATATCACGTTGACTCAGGAAGTGGTGGTGGAGACTGCCACA**TATAAATA**G<br>AGTGCCAGTAGCGACTTTTTTCACACTCGAAATACTCTTACTACTGCTCTCTTGTTGTTTTTAT<br>CACTTCTTGTTTCTTCTTGGTAAAT**AGAATATCAAGCTACAAAAA**TAAATAAAAA |
| $P_{S.\ a0111c0}$ | ACACCGGGCA*gaatttaggctaaagaaagatc*AATAAGAGAATTTCAGATTGAGAG**AATGAAAAAAAA**<br>**AAAAAAAAAAAA**GGCAGTCTTGGACAAAAACGCAGCAGGCTTGAAACAGAACGTTACGTG<br>AAGTACCTGTTGAACGCTAGTTATATCACGTTGACTCAGGAAGTGGTGGTGGAGACTGCCAC<br>A**TATAAATA**GAGTGCCAGTAGCGACTTTTTTCACACTCGAAATACTCTTACTACTGCTCTCTT<br>GTTGTTTTTATCACTTCTTGTTTCTTCTTGGTAAAT**AGAATATCAAGCTACAAAAA**TAAATAA<br>AAA |
| $P_{S.\ a0121c0}$ | ACACCGGGCA*gaatttaggctaaagaaagatc*AATAAGAGAATTTCAGATTGAGAG**AATGAAAAAAAA**<br>**AAAAAAAAAAAA**GGCAGTCTTGGACAAAAACGCAGCAGGCTTGAAACAGAACGTTACGTG<br>AAGTACCTGTTGAACGCAACCAGAGCTTAGTTATATCACGTTGACTCAGGAAGTGGTGGTGG<br>AGACTGCCACA**TATAAATA**GAGTGCCAGTAGCGACTTTTTTCACACTCGAAATACTCTTACT<br>ACTGCTCTCTTGTTGTTTTTATCACTTCTTGTTTCTTCTTGGTAAAT**AGAATATCAAGCTACA**<br>**AAAA**TAAATAAAAA |
| $P_{S.\ a0141c0}$ | ACACCGGGCA*gaatttaggctaaagaaagatc*AATAAGAGAATTTCAGATTGAGAG**AATGAAAAAAAA**<br>**AAAAAAAAAAAA**GGCAGTCTTGGACAAAAACGCAGCAGGCTTGAAACAGAACGTTACGTG<br>AAGTACCTGTTGAACGCAACCAGAGCTGAAGTGGATCTGTAATGCATTAGTTATATCACGTTG<br>ACTCAGGAAGTGGTGGTGGAGACTGCCACA**TATAAATA**GAGTGCCAGTAGCGACTTTTTTC<br>ACACTCGAAATACTCTTACTACTGCTCTCTTGTTGTTTTTATCACTTCTTGTTTCTTCTTGGTA<br>AAT**AGAATATCAAGCTACAAAAA**TAAATAAAAA |
| $P_{S.\ a0161c0}$ | ACACCGGGCA*gaatttaggctaaagaaagatc*AATAAGAGAATTTCAGATTGAGAG**AATGAAAAAAAA**<br>**AAAAAAAAAAAA**GGCAGTCTTGGACAAAAACGCAGCAGGCTTGAAACAGAACGTTACGTG<br>AAGTACCTGTTGAACGCAACCAGAGCTGAAGTGGATCTGTAATGCATGTTAGGTCGAGCAA<br>GTGTGATAGTTATATCACGTTGACTCAGGAAGTGGTGGTGGAGACTGCCACA**TATAAATA**GA<br>GTGCCAGTAGCGACTTTTTTCACACTCGAAATACTCTTACTACTGCTCTCTTGTTGTTTTTATC<br>ACTTCTTGTTTCTTCTTGGTAAAT**AGAATATCAAGCTACAAAAA**TAAATAAAAA |
| $P_{S.\ a0181c0}$ | ACACCGGGCA*gaatttaggctaaagaaagatc*AATAAGAGAATTTCAGATTGAGAG**AATGAAAAAAAA**<br>**AAAAAAAAAAAA**GGCAGTCTTGGACAAAAACGCAGCAGGCTTGAAACAGAACGTTACGTG<br>AAGTACCTGTTGAACGCAACCAGAGCTGAAGTGGATCTGTAATGCATGTTAGGTCGAGCAA<br>GTGTGATTGGCAGTAACCTGGTCCCATAGTTATATCACGTTGACTCAGGAAGTGGTGGTGGA<br>GACTGCCACA**TATAAATA**GAGTGCCAGTAGCGACTTTTTTCACACTCGAAATACTCTTACTA |

| | |
|---|---|
| | CTGCTCTCTTGTTGTTTTTATCACTTCTTGTTTCTTCTTGGTAAAT**AGAATATCAAGCTACAAAAA**TAAATAAAAA |
| $P_{S.\ a0201c0}$ | ACACCGGGCA*gaatttaggctaaagaaagatc*AATAAGAGAATTTCAGATTGAGAG**AATGAAAAAAAAAAAAAAAAAAAA**GGCAGTCTTGGACAAAAACGCAGCAGGCTTGAAACAGAACGTTACGTGAAGTACCTGTTGAACGCAACCAGAGCTGAAGTGGATCTGTAATGCATGTTAGGTCGAGCAAGTGTGATTGGCAGTAACCTGGTCCCATACGCTTAACTGCTCATTGCTAGTTATATCACGTTGACTCAGGAAGTGGTGGTGGAGACTGCCACA**TATAAATA**GAGTGCCAGTAGCGACTTTTTTCACACTCGAAATACTCTTACTACTGCTCTCTTGTTGTTTTTATCACTTCTTGTTTCTTCTTGGTAAAT**AGAATATCAAGCTACAAAAA**TAAATAAAAA |
| $P_{S.\ a0221c0}$ | ACACCGGGCA*gaatttaggctaaagaaagatc*AATAAGAGAATTTCAGATTGAGAG**AATGAAAAAAAAAAAAAAAAAAAA**GGCAGTCTTGGACAAAAACGCAGCAGGCTTGAAACAGAACGTTACGTGAAGTACCTGTTGAACGCAACCAGAGCTGAAGTGGATCTGTAATGCATGTTAGGTCGAGCAAGTGTGATTGGCAGTAACCTGGTCCCATACGCTTAACTGCTCATTGCGTCAAGGAGAACACACTATATAGTTATATCACGTTGACTCAGGAAGTGGTGGTGGAGACTGCCACA**TATAAATA**GAGTGCCAGTAGCGACTTTTTTCACACTCGAAATACTCTTACTACTGCTCTCTTGTTGTTTTTATCACTTCTTGTTTCTTCTTGGTAAAT**AGAATATCAAGCTACAAAAA**TAAATAAAAA |
| $P_{S.\ a0241c0}$ | ACACCGGGCA*gaatttaggctaaagaaagatc*AATAAGAGAATTTCAGATTGAGAG**AATGAAAAAAAAAAAAAAAAAAAA**GGCAGTCTTGGACAAAAACGCAGCAGGCTTGAAACAGAACGTTACGTGAAGTACCTGTTGAACGCAACCAGAGCTGAAGTGGATCTGTAATGCATGTTAGGTCGAGCAAGTGTGATTGGCAGTAACCTGGTCCCATACGCTTAACTGCTCATTGCGTCAAGGAGAACACACTATACTTGGAGCTGTAGTCATTTCTAGTTATATCACGTTGACTCAGGAAGTGGTGGTGGAGACTGCCACA**TATAAATA**GAGTGCCAGTAGCGACTTTTTTCACACTCGAAATACTCTTACTACTGCTCTCTTGTTGTTTTTATCACTTCTTGTTTCTTCTTGGTAAAT**AGAATATCAAGCTACAAAAA**TAAATAAAAA |
| $P_{S.\ a1b1c0}$ | ACACCGGGCA*gaatttaggctaaagaaagatc*AATATCAGATTGAGAG**AATGAAAAAAAAAAAAAAAAAAA**GGCAGTCTTGGACAATAGTTATATCACGTTGACTCAGGAAGTGGTGGTGGAGACTGCCACA**TATAAATA**GAGTGCCAGTAGCGACTTTTTTCACACTCGAAATACTCTTACTACTGCTCTCTTGTTGTTTTTATCACTTCTTGTTTCTTCTTGGTAAAT**AGAATATCAAGCTACAAAAA**TAAATAAAAA |
| $P_{S.\ a2b1c0}$ | ACACCGGGCA*gaatttaggctaaagaaagatc*AATAAGAG**AATGAAAAAAAAAAAAAAAAAAAAA**GGCAGTCTTGGACAATAGTTATATCACGTTGACTCAGGAAGTGGTGGTGGAGACTGCCACA**TATAAATA**GAGTGCCAGTAGCGACTTTTTTCACACTCGAAATACTCTTACTACTGCTCTCTTGTTGTTTTTATCACTTCTTGTTTCTTCTTGGTAAAT**AGAATATCAAGCTACAAAAA**TAAATAAAAA |
| $P_{S.\ a3b1c0}$ | ACACCGGGCA*gaatttaggctaaagaaagatc*AATAAGAGAATGAACTCTGTGTTCAGATTGAGAG**AATGAAAAAAAAAAAAAAAAAAAAA**GGCAGTCTTGGACAATAGTTATATCACGTTGACTCAGGAAGTGGTGGTGGAGACTGCCACA**TATAAATA**GAGTGCCAGTAGCGACTTTTTTCACACTCGAAATACTCTTACTACTGCTCTCTTGTTGTTTTTATCACTTCTTGTTTCTTCTTGGTAAAT**AGAATATCAAGCTACAAAAA**TAAATAAAAA |
| $P_{S.\ a4b1c0}$ | ACACCGGGCA*gaatttaggctaaagaaagatc*AATAAGAGAATGAACTCTGTGTTCGAGCACATTCAGATTGAGAG**AATGAAAAAAAAAAAAAAAAAAAAA**GGCAGTCTTGGACAATAGTTATATCACGTTGACTCAGGAAGTGGTGGTGGAGACTGCCACA**TATAAATA**GAGTGCCAGTAGCGACTTTTTTCACACTCGAAATACTCTTACTACTGCTCTCTTGTTGTTTTTATCACTTCTTGTTTCTTCTTGGTAAAT**AGAATATCAAGCTACAAAAA**TAAATAAAAA |
| $P_{S.\ a5b1c0}$ | ACACCGGGCA*gaatttaggctaaagaaagatc*AATAAGAGAATGAACTCTGTGTTCGAGCACACACTGTGTTGTTCAGATTGAGAG**AATGAAAAAAAAAAAAAAAAAAAAA**GGCAGTCTTGGACAATAGTTATATCACGTTGACTCAGGAAGTGGTGGTGGAGACTGCCACA**TATAAATA**GAGTGCCAGTAGCGACTTTTTTCACACTCGAAATACTCTTACTACTGCTCTCTTGTTGTTTTTATCACTTCTTGTTTCTTCTTGGTAAAT**AGAATATCAAGCTACAAAAA**TAAATAAAAA |
| $P_{S.\ a0b1core1}$ | ACACCGGGCA*gaatttaggctaaagaaagatc*AATAAGAGAATTTCAGATTGAGAG**AATGAAAAAAAAAAAAAAAAAAAA**GGCAGTCTTGGACAATAGTTATATCACGTTGACTCAGGAAGTGGTGGTGGAGACTGCCACA**TATAAATA**GAGTAGCACTGTTGGGCGTGAGTGGAGGCGCCGGTAAAT**AGAATATCAAGCTACAAAAA**TAAATAAAAA |
| $P_{S.\ a0b1core2}$ | ACACCGGGCA*gaatttaggctaaagaaagatc*AATAAGAGAATTTCAGATTGAGAG**AATGAAAAAAAAAAAAAAAAAAAA**GGCAGTCTTGGACAATAGTTATATCACGTTGACTCAGGAAGTGGTGGTGGAGACTGCCACA**TATAAATA**GAGTCGTAGGAGTACTCGATGGTACAGATGAGCATAAAT**AGAATATCAAGCTACAAAAA**TAAATAAAAA |
| $P_{S.\ a0b1TDH3}$ | ACACCGGGCA*gaatttaggctaaagaaagatc*AATAAGAGAATTTCAGATTGAGAG**AATGAAAAAAAAAAAAAAAAAAAA**GGCAGTCTTGGACAATAGTTATATCACGTTGACTCAGGAAGTGGTGGTGGAGACTGCCACA**TATAAATA**GACGGTAGGTATTGATTGTAATTCTGTAAATCTATTTCTTAAACTTCTTAAATTCTACTTTTATAGTTAGTCTTTTTTTAGTT**AGAATATCAAGCTACAAAAA**TAAATAAAAA |
| $P_{S.\ a0b1RNR1}$ | ACACCGGGCA*gaatttaggctaaagaaagatc*AATAAGAGAATTTCAGATTGAGAG**AATGAAAAAAAAAAAAAAAAAAAA**GGCAGTCTTGGACAATAGTTATATCACGTTGACTCAGGAAGTGGTGGTGGAGACTGCCACA**TATAAATA**GGAGCTAATATTTCATTGTTGGAAAATTACTCTACCATAATTGAAGCATATCTCATCCTTTTCATCCTTTTCAACGCAAGAGAGACACCAA**AGAATATCAAGCTACAAAAA**TAAATAAAAA |
| $P_{S.\ a0b1TEF2}$ | ACACCGGGCA*gaatttaggctaaagaaagatc*AATAAGAGAATTTCAGATTGAGAG**AATGAAAAAAAA** |

| | |
|---|---|
| | **AAAAAAAAAAAA**GGCAGTCTTGGACAATAGTTATATCACGTTGACTCAGGAAGTGGTGGTGGAGACTGCCACA**TATAAATA**TTCTCTTGCATTTTCTATTTTTCTCTCTATCTATTCTACTTGTTTATTCCCTTCAAGGTTTTTTTTTAAGGAGTACTTGTTTTTAGAATATACGGTCAACGA**AGAATATCAAGCTACAAAAA**TAAATAAAAA |
| $P_{S.\ a0b1ALD6}$ | ACACCGGGCA*gaatttaggctaaagaaagatc*AATAAGAGAATTTCAGATTGAGAG**AATGAAAAAAAAAAAAAAAAAAAA**GGCAGTCTTGGACAATAGTTATATCACGTTGACTCAGGAAGTGGTGGTGGAGACTGCCACA**TATAAATA**TGTAATAAGAAGTTTGGTAATATTCAATTCGAAGTGTTCAGTCTTTTACTTCTCTTGTTTTATAGAAGAAAAAACATCAAGA**AGAATATCAAGCTACAAAAA**TAAATAAAAA |
| $P_{S.\ a0b1CYC1}$ | ACACCGGGCA*gaatttaggctaaagaaagatc*AATAAGAGAATTTCAGATTGAGAG**AATGAAAAAAAAAAAAAAAAAAAA**GGCAGTCTTGGACAATAGTTATATCACGTTGACTCAGGAAGTGGTGGTGGAGACTGCCACA**TATAAATA**ACTCTTGTTTTCTTCTTTTCTCTAAATATTCTTTCCTTATACATTAGGACCTTTGCAGCATAAATTACTATAC**AGAATATCAAGCTACAAAAA**TAAATAAAAA |
| $P_{S.\ a0b1LEU2}$ | ACACCGGGCA*gaatttaggctaaagaaagatc*AATAAGAGAATTTCAGATTGAGAG**AATGAAAAAAAAAAAAAAAAAAAA**GGCAGTCTTGGACAATAGTTATATCACGTTGACTCAGGAAGTGGTGGTGGAGACTGCCACATTGACAATATTATTTAAGGACCTATTGTTTTTTCCAATAGGTGGTTAGCAATCGTCTTACTTTCTAACTTTTCTTACCTTTTACATTTCAGCAATATATATATATATATATTTCAAGG**AGAATATCAAGCTACAAAAA**TAAATAAAAA |
| $P_{S.\ a0b1tdh3}$ | ACACCGGGCA*gaatttaggctaaagaaagatc*AATAAGAGAATTTCAGATTGAGAG**AATGAAAAAAAAAAAAAAAAAAAA**GGCAGTCTTGGACAATAGTTATATCACGTTGACTCAGGAAGTGGTGGTGGAGACTGCCACA**TATAAATA**GACGGTAGGTATTGATTGTAATTCTGTAAATCTATTTCTTAAACTTGCTCTCTTGTTGTTTTTATCACTTCTTGTTTCTTCTTGGTAAAT**AGAATATCAAGCTACAAAAA**TAAATAAAAA |
| $P_{S.\ a0b1rnr1}$ | ACACCGGGCA*gaatttaggctaaagaaagatc*AATAAGAGAATTTCAGATTGAGAG**AATGAAAAAAAAAAAAAAAAAAAA**GGCAGTCTTGGACAATAGTTATATCACGTTGACTCAGGAAGTGGTGGTGGAGACTGCCACA**TATAAATA**GGAGCTAATATTTCATTGTTGGAAAATTACTCTACCATAATTGAATGCTCTCTTGTTGTTTTTATCACTTCTTGTTTCTTCTTGGTAAAT**AGAATATCAAGCTACAAAAA**TAAATAAAAA |
| $P_{S.\ NAb1c0}$ | ACACCGGGCA*gaatttaggctaaagaaagatc***AATGAAAAAAAAAAAAAAAAAAAA**GGCAGTCTTGGACAATAGTTATATCACGTTGACTCAGGAAGTGGTGGTGGAGACTGCCACA**TATAAATA**GAGTGCCAGTAGCGACTTTTTTCACACTCGAAATACTCTTACTACTGCTCTCTTGTTGTTTTTATCACTTCTTGTTTCTTCTTGGTAAAT**AGAATATCAAGCTACAAAAA**TAAATAAAAA |
| $P_{S.\ a0b1c0}$ | ACACCGGGCA*gaatttaggctaaagaaagatc*AATAAGAGAATTTCAGATTGAGAG**AATGAAAAAAAAAAAAAAAAAAAA**GGCAGTCTTGGACAATAGTTATATCACGTTGACTCAGGAAGTGGTGGTGGAGACTGCCACA**TATAAATA**GAGTGCCAGTAGCGACTTTTTTCACACTCGAAATACTCTTACTACTGCTCTCTTGTTGTTTTTATCACTTCTTGTTTCTTCTTGGTAAAT**AGAATATCAAGCTACAAAAA**TAAATAAAAA |
| $P_{S.\ a0b1c1}$ | ACACCGGGCA*gaatttaggctaaagaaagatc*AATAAGAGAATTTCAGATTGAGAG**AATGAAAAAAAAAAAAAAAAAAAA**GGCAGTCTTGGACAATAGTTATATCACGTTGACTCAGGAAGTGGTGGTGGAGACTGCCACA**TATAAATA**GGCCCGCTGGACTCCACTTTTACAATTCGAATACCTTTGCTCTTTTCCTATTCGACTTTTTTCTTTTCCGTTTTTAAAACTTAAGTATTT**AGAATATCAAGCTACAAAAA**TAAATAAAAA |
| $P_{S.\ a0b1c2}$ | ACACCGGGCA*gaatttaggctaaagaaagatc*AATAAGAGAATTTCAGATTGAGAG**AATGAAAAAAAAAAAAAAAAAAAA**GGCAGTCTTGGACAATAGTTATATCACGTTGACTCAGGAAGTGGTGGTGGAGACTGCCACA**TATAAATA**GGCCCGCTGGACTCCACTTTTACAATTCGAATACCTTTGCTCTTTTCCTATTCGACTTTTATCCATTTTGAATTTACAGTCATCCAACTT**AGAATATCAAGCTACAAAAA**TAAATAAAAA |
| $P_{S.\ a0b1c3}$ | ACACCGGGCA*gaatttaggctaaagaaagatc*AATAAGAGAATTTCAGATTGAGAG**AATGAAAAAAAAAAAAAAAAAAAA**GGCAGTCTTGGACAATAGTTATATCACGTTGACTCAGGAAGTGGTGGTGGAGACTGCCACA**TATAAATA**GGCCCGCTGGACTCCACTTTTACAATTCGAATACCTTTGCTCTTTTCCTATTCGACTTTTTTCTTATCTTTTATTGGCTTCTTGTAACTT**AGAATATCAAGCTACAAAAA**TAAATAAAAA |
| $P_{S.\ a0b1c4}$ | ACACCGGGCA*gaatttaggctaaagaaagatc*AATAAGAGAATTTCAGATTGAGAG**AATGAAAAAAAAAAAAAAAAAAAA**GGCAGTCTTGGACAATAGTTATATCACGTTGACTCAGGAAGTGGTGGTGGAGACTGCCACA**TATAAATA**GGCCCGCTGGACTCCACTTTTACAATTCGATCGTTTCCTCTTGTAATCCGCTTTTTGTCTCTTTTCTTTTCCGTTTTTAAAACTTAAGTATTT**AGAATATCAAGCTACAAAAA**TAAATAAAAA |
| $P_{S.\ a0b1c5}$ | ACACCGGGCA*gaatttaggctaaagaaagatc*AATAAGAGAATTTCAGATTGAGAG**AATGAAAAAAAAAAAAAAAAAAAA**GGCAGTCTTGGACAATAGTTATATCACGTTGACTCAGGAAGTGGTGGTGGAGACTGCCACA**TATAAATA**GGCCCGCTGGACTCCACTTTTACAATTCGATCGTTTCCTCTTGTAATCCGCTTTTTGTCTCTTATCCATTTTGAATTTACAGTCATCCAACTT**AGAATATCAAGCTACAAAAA**TAAATAAAAA |
| $P_{S.\ a0b1c6}$ | ACACCGGGCA*gaatttaggctaaagaaagatc*AATAAGAGAATTTCAGATTGAGAG**AATGAAAAAAAAAAAAAAAAAAAA**GGCAGTCTTGGACAATAGTTATATCACGTTGACTCAGGAAGTGGTGGTGGAGACTGCCACA**TATAAATA**GGCCCGCTGGACTCCACTTTTACAATTCGATCGTTTCCTCTTGTAATCCGCTTTTTGTCTCTTTTCTTATCTTTTATTGGCTTCTTGTAACTT**AGAATATCAAGCTACAAAAA**TAAATAAAAA |

| | |
|---|---|
| $P_{S.\ a0b1c7}$ | ACACCGGGCA*gaatttaggctaaagaaagatc*AATAAGAGAATTTCAGATTGAGAG**AATGAAAAAAAAAAAAAAAAAAAA**GGCAGTCTTGGACAATAGTTATATCACGTTGACTCAGGAAGTGGTGGTGGAGACTGCCACA**TATAAATA**GGCCCGCTGGACTCCACTTTTACAATTCGATTTCTTTTAGATTCTGGTTCTTTGATATTTTTCTTTTCCGTTTTTAAAACTTAAGTATTT**AGAATATCAAGCTACAAAAA**TAAATAAAAA |
| $P_{S.\ a0b1c8}$ | ACACCGGGCA*gaatttaggctaaagaaagatc*AATAAGAGAATTTCAGATTGAGAG**AATGAAAAAAAAAAAAAAAAAAAA**GGCAGTCTTGGACAATAGTTATATCACGTTGACTCAGGAAGTGGTGGTGGAGACTGCCACA**TATAAATA**GGCCCGCTGGACTCCACTTTTACAATTCGATTTCTTTTAGATTCTGGTTCTTTGATATTTATCCATTTTGAATTTACAGTCATCCAACTT**AGAATATCAAGCTACAAAAA**TAAATAAAAA |
| $P_{S.\ a0b1c9}$ | ACACCGGGCA*gaatttaggctaaagaaagatc*AATAAGAGAATTTCAGATTGAGAG**AATGAAAAAAAAAAAAAAAAAAAA**GGCAGTCTTGGACAATAGTTATATCACGTTGACTCAGGAAGTGGTGGTGGAGACTGCCACA**TATAAATA**GGCCCGCTGGACTCCACTTTTACAATTCGATTTCTTTTAGATTCTGGTTCTTTGATATTTTTCTTATCTTTTATTGGCTTCTTGTAACTT**AGAATATCAAGCTACAAAAA**TAAATAAAAA |
| $P_{S.\ a0b1c10}$ | ACACCGGGCA*gaatttaggctaaagaaagatc*AATAAGAGAATTTCAGATTGAGAG**AATGAAAAAAAAAAAAAAAAAAAA**GGCAGTCTTGGACAATAGTTATATCACGTTGACTCAGGAAGTGGTGGTGGAGACTGCCACA**TATAAATA**GGCCCGCTGGACTCCACTTTTACAATTCGAAATTCACTTCTGTTTACTTGGCCCTCTTTTTTCTTTTCCGTTTTTAAAACTTAAGTATTT**AGAATATCAAGCTACAAAAA**TAAATAAAAA |
| $P_{S.\ a0b1c11}$ | ACACCGGGCA*gaatttaggctaaagaaagatc*AATAAGAGAATTTCAGATTGAGAG**AATGAAAAAAAAAAAAAAAAAAAA**GGCAGTCTTGGACAATAGTTATATCACGTTGACTCAGGAAGTGGTGGTGGAGACTGCCACA**TATAAATA**GGCCCGCTGGACTCCACTTTTACAATTCGAAATTCACTTCTGTTTACTTGGCCCTCTTTTATCCATTTTGAATTTACAGTCATCCAACTT**AGAATATCAAGCTACAAAAA**TAAATAAAAA |
| $P_{S.\ a0b1c12}$ | ACACCGGGCA*gaatttaggctaaagaaagatc*AATAAGAGAATTTCAGATTGAGAG**AATGAAAAAAAAAAAAAAAAAAAA**GGCAGTCTTGGACAATAGTTATATCACGTTGACTCAGGAAGTGGTGGTGGAGACTGCCACA**TATAAATA**GGCCCGCTGGACTCCACTTTTACAATTCGAAATTCACTTCTGTTTACTTGGCCCTCTTTTTTCTTATCTTTTATTGGCTTCTTGTAACTT**AGAATATCAAGCTACAAAAA**TAAATAAAAA |
| $P_{S.\ a0b2c0}$ | ACACCGGGCA*gaatttaggctaaagaaagatc*AATAAGAGAATTTCAGATTGAGAG**AATGAAAAAAAAAAAAAAAAAAAA**GGCAGCGCGTTTCTTTCCCTGCCCTAGCATGAAGACCTAGCCTTTGTATCTGTCGTACACA**TATAAATA**GAGTGCCAGTAGCGACTTTTTTCACACTCGAAATACTCTTACTACTGCTCTCTTGTTGTTTTTATCACTTCTTGTTTCTTCTTGGTAAAT**AGAATATCAAGCTACAAAAA**TAAATAAAAA |
| $P_{S.\ a0b3c0}$ | ACACCGGGCA*gaatttaggctaaagaaagatc*AATAAGAGAATTTCAGATTGAGAG**AATGAAAAAAAAAAAAAAAAAAAA**GGCAGCTTTCGATCCGAGTGCCTCAACTGTTCACCCCTTGCTATAACACCCAGTCTACACA**TATAAATA**GAGTGCCAGTAGCGACTTTTTTCACACTCGAAATACTCTTACTACTGCTCTCTTGTTGTTTTTATCACTTCTTGTTTCTTCTTGGTAAAT**AGAATATCAAGCTACAAAAA**TAAATAAAAA |
| $P_{S.\ a0b1c13}$ | ACACCGGGCA*gaatttaggctaaagaaagatc*AATAAGAGAATTTCAGATTGAGAG**AATGAAAAAAAAAAAAAAAAAAAA**GGCAGTCTTGGACAATAGTTATATCACGTTGACTCAGGAAGTGGTGGTGGAGACTGCCACA**TATAAATA**GAGTGCCAGTACTCCACTTTTACAATTCGAAAATACTCTTTTGTAATCCGGCCCTCTTTTTTTATCACTTTTATTGGCTTCTTGTAACTT**AGAATATCAAGCTACAAAAA**TAAATAAAAA |
| $P_{S.\ a0b1c14}$ | ACACCGGGCA*gaatttaggctaaagaaagatc*AATAAGAGAATTTCAGATTGAGAG**AATGAAAAAAAAAAAAAAAAAAAA**GGCAGTCTTGGACAATAGTTATATCACGTTGACTCAGGAAGTGGTGGTGGAGACTGCCACA**TATAAATA**GGCCCGCTGGACTCCACTTTTTCACACTCGTCGTTTCCTCTGTTTACTTGTCTTGTTGTTTTCTTATCTTTTATTGGCTTCTTGGTAAAT**AGAATATCAAGCTACAAAAA**TAAATAAAAA |
| $P_{S.\ a0b1c5}$ | ACACCGGGCA*gaatttaggctaaagaaagatc*AATAAGAGAATTTCAGATTGAGAG**AATGAAAAAAAAAAAAAAAAAAAA**GGCAGTCTTGGACAATAGTTATATCACGTTGACTCAGGAAGTGGTGGTGGAGACTGCCACA**TATAAATA**GGCCCGCTGGAGCGACTTTTTACAATTCGAAATTCACTTCTTGTAATCCGGCCCTCTTTTTTCTTATCTTTTATTGGCTTCTTGTAACTT**AGAATATCAAGCTACAAAAA**TAAATAAAAA |
| $P_{S.\ a0b1c16}$ | ACACCGGGCA*gaatttaggctaaagaaagatc*AATAAGAGAATTTCAGATTGAGAG**AATGAAAAAAAAAAAAAAAAAAAA**GGCAGTCTTGGACAATAGTTATATCACGTTGACTCAGGAAGTGGTGGTGGAGACTGCCACA**TATAAATA**GGCCCGCTGGACTCCACTTTTTCACACTCGAATTCACTTCTTGTAATCCGTCTTGTTGTTTTCTTATCTTTTATTGGCTTCTTGGTAAAT**AGAATATCAAGCTACAAAAA**TAAATAAAAA |
| $P_{S.\ a0b1c17}$ | ACACCGGGCA*gaatttaggctaaagaaagatc*AATAAGAGAATTTCAGATTGAGAG**AATGAAAAAAAAAAAAAAAAAAAA**GGCAGTCTTGGACAATAGTTATATCACGTTGACTCAGGAAGTGGTGGTGGAGACTGCCACA**TATAAATA**TCTGCTCCTTGAAGGCGGTATTCCCCAATTTAGAGTTCTTCCGAGTAATTGTATTGATAAGTTCTCCTTTAGTCTAATTGATGTTAAATT**AGAATATCAAGCTACAAAAA**TAAATAAAAA |
| $P_{S.\ a0b1c18}$ | ACACCGGGCA*gaatttaggctaaagaaagatc*AATAAGAGAATTTCAGATTGAGAG**AATGAAAAAAAAAAAAAAAAAAAA**GGCAGTCTTGGACAATAGTTATATCACGTTGACTCAGGAAGTGGTGGTG |

| | |
|---|---|
| | GAGACTGCCACA**TATAAATA**CCTGTGACATATGCTAGTGTCTACACGGTGGAGTTCATGTTCACAATATACGTTGATTAGCTTTTACATAGTAGACTTCTAATCTATTG**AGAATATCAAGCTACAAAAA**TAAATAAAAA |
| $P_{S.\ a0b1c19}$ | ACACCGGGCA*gaatttaggctaaagaaagatc*AATAAGAGAATTTCAGATTGAGAG**AATGAAAAAAAAAAAAAAAAAAAA**GGCAGTCTTGGACAATAGTTATATCACGTTGACTCAGGAAGTGGTGGTGGAGACTGCCACA**TATAAATA**CTGTAGGTTTCGATTCACTCTCACCTGTGGTGTTCCGTCTTTATAGTGTATTAGTTATCCGTTATTTACTTTATCTCGATGCTACTTTAA**AGAATATCAAGCTACAAAAA**TAAATAAAAA |
| $P_{S.\ a0b1c20}$ | ACACCGGGCA*gaatttaggctaaagaaagatc*AATAAGAGAATTTCAGATTGAGAG**AATGAAAAAAAAAAAAAAAAAAAA**GGCAGTCTTGGACAATAGTTATATCACGTTGACTCAGGAAGTGGTGGTGGAGACTGCCACA**TATAAATA**CTTTCCCATGGTTCAGTTACCTCAGACTCCGTACATAGTATTCCATTTCATTAGGTCCTATAGTACCATATTTCTTCCTTAGTCAAAATT**AGAATATCAAGCTACAAAAA**TAAATAAAAA |
| $P_{S.\ a0b4c0}$ | ACACCGGGCA*gaatttaggctaaagaaagatc*AATAAGAGAATTTCAGATTGAGAG**AATGAAAAAAAAAAAAAAAAAAAA**TGCGTACCCTTATACCCTTCTTTGCATCTGTCCAATCCTTAGTGAACAATGATGGGTAAGC**TATAAATA**GAGTGCCAGTAGCGACTTTTTTCACACTCGAAATACTCTTACTACTGCTCTCTTGTTGTTTTTATCACTTCTTGTTTCTTCTTGGTAAAT**AGAATATCAAGCTACAAAAA**TAAATAAAAA |
| $P_{S.\ a0b5c0}$ | ACACCGGGCA*gaatttaggctaaagaaagatc*AATAAGAGAATTTCAGATTGAGAG**AATGAAAAAAAAAAAAAAAAAAAA**TCGTTGCCAGAAGCTCTGTGCAATTAATGAAGGACTCATGTCACTACACCATCGGGGATGA**TATAAATA**GAGTGCCAGTAGCGACTTTTTTCACACTCGAAATACTCTTACTACTGCTCTCTTGTTGTTTTTATCACTTCTTGTTTCTTCTTGGTAAAT**AGAATATCAAGCTACAAAAA**TAAATAAAAA |
| $P_{S.\ a0b6c0}$ | ACACCGGGCA*gaatttaggctaaagaaagatc*AATAAGAGAATTTCAGATTGAGAG**AATGAAAAAAAAAAAAAAAAAAAA**GGGACCTTTAGCAAACGTCGGTTCCGCAGAGAATGACGTCAAGACGATAGCGCCGCGGGAC**TATAAATA**GAGTGCCAGTAGCGACTTTTTTCACACTCGAAATACTCTTACTACTGCTCTCTTGTTGTTTTTATCACTTCTTGTTTCTTCTTGGTAAAT**AGAATATCAAGCTACAAAAA**TAAATAAAAA |
| $P_{S.\ a0b7c0}$ | ACACCGGGCA*gaatttaggctaaagaaagatc*AATAAGAGAATTTCAGATTGAGAG**AATGAAAAAAAAAAAAAAAAAAAA**GTGTCACTAGATGAAGAGATGCATACCCTTATACAGAACCCTCATAGTTGTAAGTTGCACA**TATAAATA**GAGTGCCAGTAGCGACTTTTTTCACACTCGAAATACTCTTACTACTGCTCTCTTGTTGTTTTTATCACTTCTTGTTTCTTCTTGGTAAAT**AGAATATCAAGCTACAAAAA**TAAATAAAAA |
| $P_{S.\ a0b8c0}$ | ACACCGGGCA*gaatttaggctaaagaaagatc*AATAAGAGAATTTCAGATTGAGAG**AATGAAAAAAAAAAAAAAAAAAAA**AGCATAGTGTCAATTACACTGATACACTAGGTTTTAACCTACCTTCACCACTTGTAAAGGA**TATAAATA**GAGTGCCAGTAGCGACTTTTTTCACACTCGAAATACTCTTACTACTGCTCTCTTGTTGTTTTTATCACTTCTTGTTTCTTCTTGGTAAAT**AGAATATCAAGCTACAAAAA**TAAATAAAAA |
| $P_{S.\ a0b9c0}$ | ACACCGGGCA*gaatttaggctaaagaaagatc*AATAAGAGAATTTCAGATTGAGAG**AATGAAAAAAAAAAAAAAAAAAAA**CTTCGGCTGG**GATACT**CCATGCTCGATTATCAAATCTGCCTCCACACTTGTTCAGTTGGGC**TATAAATA**GAGTGCCAGTAGCGACTTTTTTCACACTCGAAATACTCTTACTACTGCTCTCTTGTTGTTTTTATCACTTCTTGTTTCTTCTTGGTAAAT**AGAATATCAAGCTACAAAAA**TAAATAAAAA |
| $P_{S.\ a0b10c0}$ | ACACCGGGCA*gaatttaggctaaagaaagatc*AATAAGAGAATTTCAGATTGAGAG**AATGAAAAAAAAAAAAAAAAAAAA**AAATGTTAACATGGCGCCCAGTATCTACCGCACGCGGCTGCTTGGTTTAGCCGCTTGAGAAT**TATAAATA**GAGTGCCAGTAGCGACTTTTTTCACACTCGAAATACTCTTACTACTGCTCTCTTGTTGTTTTTATCACTTCTTGTTTCTTCTTGGTAAAT**AGAATATCAAGCTACAAAAA**TAAATAAAAA |
| $P_{S.\ a0b11c0}$ | ACACCGGGCA*gaatttaggctaaagaaagatc*AATAAGAGAATTTCAGATTGAGAG**AATGAAAAAAAAAAAAAAAAAAAA**AGACTACTGGTAGGCTAGCTGTTAGTACTCCTCCTACTGCCAAACTACTATGACAGTGTTC**TATAAATA**GAGTGCCAGTAGCGACTTTTTTCACACTCGAAATACTCTTACTACTGCTCTCTTGTTGTTTTTATCACTTCTTGTTTCTTCTTGGTAAAT**AGAATATCAAGCTACAAAAA**TAAATAAAAA |
| $P_{S.\ a0b12c0}$ | ACACCGGGCA*gaatttaggctaaagaaagatc*AATAAGAGAATTTCAGATTGAGAG**AATGAAAAAAAAAAAAAAAAAAAA**GGCAATCCCCACAATGGGTCCCTGAAGGCGTTTGACGAAGTTGTCTGCAGAGCTGTTTCACA**TATAAATA**GAGTGCCAGTAGCGACTTTTTTCACACTCGAAATACTCTTACTACTGCTCTCTTGTTGTTTTTATCACTTCTTGTTTCTTCTTGGTAAAT**AGAATATCAAGCTACAAAAA**TAAATAAAAA |
| $P_{S.\ a0b13c0}$ | ACACCGGGCA*gaatttaggctaaagaaagatc*AATAAGAGAATTTCAGATTGAGAG**AATGAAAAAAAAAAAAAAAAAAAA**GGCAAGTAACAAGTAGTTTGTTCGATCAGTCCCTGGTCGCAAGCAAGTGGAGGGCGATCACA**TATAAATA**GAGTGCCAGTAGCGACTTTTTTCACACTCGAAATACTCTTACTACTGCTCTCTTGTTGTTTTTATCACTTCTTGTTTCTTCTTGGTAAAT**AGAATATCAAGCTACAAAAA**TAAATAAAAA |
| $P_{S.\ a0b14c0}$ | ACACCGGGCA*gaatttaggctaaagaaagatc*AATAAGAGAATTTCAGATTGAGAG**AATGAAAAAAAAAAAAAAAAAAAA**GGCAATCTGCTACAGGAACTTATTACGGCAACGGCCTGAAGCTCCGATGAGCACTAATCACA**TATAAATA**GAGTGCCAGTAGCGACTTTTTTCACACTCGAAATACTCTTAC |

| | |
|---|---|
| | TACTGCTCTCTTGTTGTTTTTATCACTTCTTGTTTCTTCTTGGTAAAT**AGAATATCAAGCTACAAAAA**TAAATAAAAA |
| $P_{S.\ a0b15c0}$ | ACACCGGGCA*gaatttaggctaaagaaagatc*AATAAGAGAATTTCAGATTGAGAG**AATGAAAAAAAAAAAAAAAAAAAA**GGCAAAATGTAGTCCGCTTCTTGGCGAAAGGTACTGCACTTCAACCATTGCACAAGATCACA**TATAAATA**GAGTGCCAGTAGCGACTTTTTTCACACTCGAAATACTCTTACTACTGCTCTCTTGTTGTTTTTATCACTTCTTGTTTCTTCTTGGTAAAT**AGAATATCAAGCTACAAAAA**TAAATAAAAA |
| $P_{S.\ a0b16c0}$ | ACACCGGGCA*gaatttaggctaaagaaagatc*AATAAGAGAATTTCAGATTGAGAG**AATGAAAAAAAAAAAAAAAAAAAA**GGCAAATGCTCATCGGGTCCGTCAGGCCATGTGGAGTACGCAGGCCTCTCGGGCAGATTCACA**TATAAATA**GAGTGCCAGTAGCGACTTTTTTCACACTCGAAATACTCTTACTACTGCTCTCTTGTTGTTTTTATCACTTCTTGTTTCTTCTTGGTAAAT**AGAATATCAAGCTACAAAAA**TAAATAAAAA |
| $P_{S.\ a0b17c0}$ | ACACCGGGCA*gaatttaggctaaagaaagatc*AATAAGAGAATTTCAGATTGAGAG**AATGAAAAAAAAAAAAAAAAAAAA**GGCAATCGGCAAGAAAGATTTCTATCACTAATGCTCAAGAGAAGCTTCCACGGCATGTCACA**TATAAATA**GAGTGCCAGTAGCGACTTTTTTCACACTCGAAATACTCTTACTACTGCTCTCTTGTTGTTTTTATCACTTCTTGTTTCTTCTTGGTAAAT**AGAATATCAAGCTACAAAAA**TAAATAAAAA |
| $P_{S.\ a0b1c21}$ | ACACCGGGCA*gaatttaggctaaagaaagatc*AATAAGAGAATTTCAGATTGAGAG**AATGAAAAAAAAAAAAAAAAAAAA**GGCAGTCTTGGACAATAGTTATATCACGTTGACTCAGGAAGTGGTGGTGGAGACTGCCACA**TATAAATAT**GGGTGTTGCTGCTGATCTCTTCACTTTCTAATCCACCTTAATTCAGATATCTCTGTCATTATCATTTAGTAACTGCATCTTAACAATTT**AGAATATCAAGCTACAAAAA**TAAATAAAAA |
| $P_{S.\ a0b1c22}$ | ACACCGGGCA*gaatttaggctaaagaaagatc*AATAAGAGAATTTCAGATTGAGAG**AATGAAAAAAAAAAAAAAAAAAAA**GGCAGTCTTGGACAATAGTTATATCACGTTGACTCAGGAAGTGGTGGTGGAGACTGCCACA**TATAAATAT**GGGTGTTGCTGCTGATCTCTTCACTTTCTAATCCACCTTAATTCAGATATCTCTGTCATTTCTATTCGTTAACTGGCTTCTTGCTTCAA**AGAATATCAAGCTACAAAAA**TAAATAAAAA |
| $P_{S.\ a0b1c23}$ | ACACCGGGCA*gaatttaggctaaagaaagatc*AATAAGAGAATTTCAGATTGAGAG**AATGAAAAAAAAAAAAAAAAAAAA**GGCAGTCTTGGACAATAGTTATATCACGTTGACTCAGGAAGTGGTGGTGGAGACTGCCACA**TATAAATAT**GGGTGTTGCTGCTGATCTCTTCACTTTCTTTACTTCCTATCTTTGCGTTGTTTACTCTATATCATTTAGTAACTGCATCTTAACAATTT**AGAATATCAAGCTACAAAAA**TAAATAAAAA |
| $P_{S.\ a0b1c24}$ | ACACCGGGCA*gaatttaggctaaagaaagatc*AATAAGAGAATTTCAGATTGAGAG**AATGAAAAAAAAAAAAAAAAAAAA**GGCAGTCTTGGACAATAGTTATATCACGTTGACTCAGGAAGTGGTGGTGGAGACTGCCACA**TATAAATAT**GGGTGTTGCTGCTGATCTCTTCACTTTCTTTACTTCCTATCTTTGCGTTGTTTACTCTATTCTATTCGTTAACTGGCTTCTTGCTTCAA**AGAATATCAAGCTACAAAAA**TAAATAAAAA |
| $P_{S.\ a0b1c25}$ | ACACCGGGCA*gaatttaggctaaagaaagatc*AATAAGAGAATTTCAGATTGAGAG**AATGAAAAAAAAAAAAAAAAAAAA**GGCAGTCTTGGACAATAGTTATATCACGTTGACTCAGGAAGTGGTGGTGGAGACTGCCACA**TATAAATA**GGAGCGTTCGAGTCGTCTTTAATAGATCATAATCCACCTTAATTCAGATATCTCTGTCATTATCATTTAGTAACTGCATCTTAACAATTT**AGAATATCAAGCTACAAAAA**TAAATAAAAA |
| $P_{S.\ a0b1c26}$ | ACACCGGGCA*gaatttaggctaaagaaagatc*AATAAGAGAATTTCAGATTGAGAG**AATGAAAAAAAAAAAAAAAAAAAA**GGCAGTCTTGGACAATAGTTATATCACGTTGACTCAGGAAGTGGTGGTGGAGACTGCCACA**TATAAATA**GGAGCGTTCGAGTCGTCTTTAATAGATCATAATCCACCTTAATTCAGATATCTCTGTCATTTCTATTCGTTAACTGGCTTCTTGCTTCAA**AGAATATCAAGCTACAAAAA**TAAATAAAAA |
| $P_{S.\ a0b1c27}$ | ACACCGGGCA*gaatttaggctaaagaaagatc*AATAAGAGAATTTCAGATTGAGAG**AATGAAAAAAAAAAAAAAAAAAAA**GGCAGTCTTGGACAATAGTTATATCACGTTGACTCAGGAAGTGGTGGTGGAGACTGCCACA**TATAAATA**GGAGCGTTCGAGTCGTCTTTAATAGATCATTTACTTCCTATCTTTGCGTTGTTTACTCTATATCATTTAGTAACTGCATCTTAACAATTT**AGAATATCAAGCTACAAAAA**TAAATAAAAA |
| $P_{S.\ a0b1c28}$ | ACACCGGGCA*gaatttaggctaaagaaagatc*AATAAGAGAATTTCAGATTGAGAG**AATGAAAAAAAAAAAAAAAAAAAA**GGCAGTCTTGGACAATAGTTATATCACGTTGACTCAGGAAGTGGTGGTGGAGACTGCCACA**TATAAATA**GGAGCGTTCGAGTCGTCTTTAATAGATCATTTACTTCCTATCTTTGCGTTGTTTACTCTATTCTATTCGTTAACTGGCTTCTTGCTTCAA**AGAATATCAAGCTACAAAAA**TAAATAAAAA |
| $P_{S.\ a0b23c0}$ | ACACCGGGCA*gaatttaggctaaagaaagatc*AATAAGAGAATTTCAGATTGAGAG**AATGAAAAAAAAAAAAAAAAAAAA**GGCAGTCTTGGACAATAGTGCAATTAATGAATACAGAACCCTCATCACCACTTGTAACACA**TATAAATA**GAGTGCCAGTAGCGACTTTTTTCACACTCGAAATACTCTTACTACTGCTCTCTTGTTGTTTTTATCACTTCTTGTTTCTTCTTGGTAAAT**AGAATATCAAGCTACAAAAA**TAAATAAAAA |
| $P_{S.\ a0b24c0}$ | ACACCGGGCA*gaatttaggctaaagaaagatc*AATAAGAGAATTTCAGATTGAGAG**AATGAAAAAAAAAAAAAAAAAAAA**GGCAGGCCAGAAGCTCTGATGCATACCCTTATTTAACCTACCTTGGTGGAGACTGCCACA**TATAAATA**GAGTGCCAGTAGCGACTTTTTTCACACTCGAAATACTCTTACTACTGCTCTCTTGTTGTTTTTATCACTTCTTGTTTCTTCTTGGTAAAT**AGAATATCAAGCTACAAAAA**TAAATAAAAA |

| | |
|---|---|
| $P_{S.\ a0b25c0}$ | ACACCGGGCA*gaatttaggctaaagaaagatc*AATAAGAGAATTTCAGATTGAGAG**AATGAAAAAAAA**<br>**AAAAAAAAAAAA**GGCAGACTAGATGAAGAGCTGATACACTAGGACTCAGGAAGTGGTACC<br>CATCGGGGCACA**TATAAATA**GAGTGCCAGTAGCGACTTTTTTCACACTCGAAATACTCTTAC<br>TACTGCTCTCTTGTTGTTTTTATCACTTCTTGTTTCTTCTTGGTAAAT**AGAATATCAAGCTACA**<br>**AAAA**TAAATAAAAA |
| $P_{S.\ a0b26c0}$ | ACACCGGGCA*gaatttaggctaaagaaagatc*AATAAGAGAATTTCAGATTGAGAG**AATGAAAAAAAA**<br>**AAAAAAAAAAAA**GGCAGAGTGTCAATTACATTATATCACGTTGGGACTCATGTCACTAGTTG<br>TAAGTTGCACA**TATAAATA**GAGTGCCAGTAGCGACTTTTTTCACACTCGAAATACTCTTACTA<br>CTGCTCTCTTGTTGTTTTTATCACTTCTTGTTTCTTCTTGGTAAAT**AGAATATCAAGCTACAA**<br>**AAA**TAAATAAAAA |
| $P_{S.\ a0b27c0}$ | ACACCGGGCA*gaatttaggctaaagaaagatc*AATAAGAGAATTTCAGATTGAGAG**AATGAAAAAAAA**<br>**AAAAAAAAAAAA**GGCAGTCTTGGACAATAGCTGATACACTAGGTACAGAACCCTCATACAC<br>CATCGGGGCACA**TATAAATA**GAGTGCCAGTAGCGACTTTTTTCACACTCGAAATACTCTTAC<br>TACTGCTCTCTTGTTGTTTTTATCACTTCTTGTTTCTTCTTGGTAAAT**AGAATATCAAGCTACA**<br>**AAAA**TAAATAAAAA |
| $P_{S.\ a0b29c0}$ | ACACCGGGCA*gaatttaggctaaagaaagatc*AATAAGAGAATTTCAGATTGAGAG**AATGAAAAAAAA**<br>**AAAAAAAAAAAA**GGCAGACTAGATGAAGAGTTATATCACGTTGTTTTAACCTACCTTACACC<br>ATCGGGGCACA**TATAAATA**GAGTGCCAGTAGCGACTTTTTTCACACTCGAAATACTCTTACT<br>ACTGCTCTCTTGTTGTTTTTATCACTTCTTGTTTCTTCTTGGTAAAT**AGAATATCAAGCTACA**<br>**AAAA**TAAATAAAAA |
| $P_{S.\ a0b30c0}$ | ACACCGGGCA*gaatttaggctaaagaaagatc*AATAAGAGAATTTCAGATTGAGAG**AATGAAAAAAAA**<br>**AAAAAAAAAAAA**GGCAGAGTGTCAATTACAATGCATACCCTTAGGACTCATGTCACTGGTG<br>GAGACTGCCACA**TATAAATA**GAGTGCCAGTAGCGACTTTTTTCACACTCGAAATACTCTTAC<br>TACTGCTCTCTTGTTGTTTTTATCACTTCTTGTTTCTTCTTGGTAAAT**AGAATATCAAGCTACA**<br>**AAAA**TAAATAAAAA |
| $P_{S.\ a6b1c0}$ | ACACCGGGCA*gaatttaggctaaagaaagatc*CACCTTCTATTGGATGGGAATGAT**AATGAAAAAAAA**<br>**AAAAAAAAAAAA**GGCAGTCTTGGACAATAGTTATATCACGTTGACTCAGGAAGTGGTGGTGG<br>AGACTGCCACA**TATAAATA**GAGTGCCAGTAGCGACTTTTTTCACACTCGAAATACTCTTACT<br>ACTGCTCTCTTGTTGTTTTTATCACTTCTTGTTTCTTCTTGGTAAAT**AGAATATCAAGCTACA**<br>**AAAA**TAAATAAAAA |
| $P_{S.\ a7b1c0}$ | ACACCGGGCA*gaatttaggctaaagaaagatc*AACATATGAGGAAAGGCCCCAAAA**AATGAAAAAAAA**<br>**AAAAAAAAAAAA**GGCAGTCTTGGACAATAGTTATATCACGTTGACTCAGGAAGTGGTGGTG<br>GAGACTGCCACA**TATAAATA**GAGTGCCAGTAGCGACTTTTTTCACACTCGAAATACTCTTAC<br>TACTGCTCTCTTGTTGTTTTTATCACTTCTTGTTTCTTCTTGGTAAAT**AGAATATCAAGCTACA**<br>**AAAA**TAAATAAAAA |
| $P_{S.\ a8b1c0}$ | ACACCGGGCA*gaatttaggctaaagaaagatc*TCCAAAGTCACAAGTCCCAAAGAA**AATGAAAAAAA**<br>**AAAAAAAAAAAAA**GGCAGTCTTGGACAATAGTTATATCACGTTGACTCAGGAAGTGGTGGT<br>GGAGACTGCCACA**TATAAATA**GAGTGCCAGTAGCGACTTTTTTCACACTCGAAATACTCTTA<br>CTACTGCTCTCTTGTTGTTTTTATCACTTCTTGTTTCTTCTTGGTAAAT**AGAATATCAAGCTAC**<br>**AAAAA**TAAATAAAAA |
| $P_{S.\ a9b1c0}$ | ACACCGGGCA*gaatttaggctaaagaaagatc*CAAGGTAATTGAACGAGCATGAAC**AATGAAAAAAAA**<br>**AAAAAAAAAAAA**GGCAGTCTTGGACAATAGTTATATCACGTTGACTCAGGAAGTGGTGGTG<br>GAGACTGCCACA**TATAAATA**GAGTGCCAGTAGCGACTTTTTTCACACTCGAAATACTCTTAC<br>TACTGCTCTCTTGTTGTTTTTATCACTTCTTGTTTCTTCTTGGTAAAT**AGAATATCAAGCTACA**<br>**AAAA**TAAATAAAAA |
| $P_{S.\ a10b1c0}$ | ACACCGGGCA*gaatttaggctaaagaaagatc*ATAATAGGCATCGAATTAACTATT**AATGAAAAAAAA**<br>**AAAAAAAAAAAA**GGCAGTCTTGGACAATAGTTATATCACGTTGACTCAGGAAGTGGTGGTGG<br>AGACTGCCACA**TATAAATA**GAGTGCCAGTAGCGACTTTTTTCACACTCGAAATACTCTTACT<br>ACTGCTCTCTTGTTGTTTTTATCACTTCTTGTTTCTTCTTGGTAAAT**AGAATATCAAGCTACA**<br>**AAAA**TAAATAAAAA |
| $P_{S.\ a11b1c0}$ | ACACCGGGCA*gaatttaggctaaagaaagatc*AAAGTTCCTCGAAGGATAGCAGAA**AATGAAAAAAAA**<br>**AAAAAAAAAAAA**GGCAGTCTTGGACAATAGTTATATCACGTTGACTCAGGAAGTGGTGGTG<br>GAGACTGCCACA**TATAAATA**GAGTGCCAGTAGCGACTTTTTTCACACTCGAAATACTCTTAC<br>TACTGCTCTCTTGTTGTTTTTATCACTTCTTGTTTCTTCTTGGTAAAT**AGAATATCAAGCTACA**<br>**AAAA**TAAATAAAAA |
| $P_{S.\ a12b1c0}$ | ACACCGGGCA*gaatttaggctaaagaaagatc*AACTAATGGCTACAATGAGTCTAG**AATGAAAAAAAA**<br>**AAAAAAAAAAAA**GGCAGTCTTGGACAATAGTTATATCACGTTGACTCAGGAAGTGGTGGTG<br>GAGACTGCCACA**TATAAATA**GAGTGCCAGTAGCGACTTTTTTCACACTCGAAATACTCTTAC<br>TACTGCTCTCTTGTTGTTTTTATCACTTCTTGTTTCTTCTTGGTAAAT**AGAATATCAAGCTACA**<br>**AAAA**TAAATAAAAA |
| $P_{S.\ a13b1c0}$ | ACACCGGGCA*gaatttaggctaaagaaagatc*ATGGTTGTGTGAAGGTTGATCCAA**AATGAAAAAAAA**<br>**AAAAAAAAAAAA**GGCAGTCTTGGACAATAGTTATATCACGTTGACTCAGGAAGTGGTGGTG<br>GAGACTGCCACA**TATAAATA**GAGTGCCAGTAGCGACTTTTTTCACACTCGAAATACTCTTAC<br>TACTGCTCTCTTGTTGTTTTTATCACTTCTTGTTTCTTCTTGGTAAAT**AGAATATCAAGCTACA**<br>**AAAA**TAAATAAAAA |
| $P_{S.\ a14b1c0}$ | ACACCGGGCA*gaatttaggctaaagaaagatc*AATCAACCACTACACTACAAGAAT**AATGAAAAAAAA**<br>**AAAAAAAAAAAA**GGCAGTCTTGGACAATAGTTATATCACGTTGACTCAGGAAGTGGTGGTG |

| | |
|---|---|
| | GAGACTGCCACA**TATAAATA**GAGTGCCAGTAGCGACTTTTTTCACACTCGAAATACTCTTAC<br>TACTGCTCTCTTGTTGTTTTTATCACTTCTTGTTTCTTCTTGGTAAAT**AGAATATCAAGCTACA**<br>**AAAA**TAAATAAAAA |
| $P_{S.\ a15b1c0}$ | ACACCGGGCA*gaatttaggctaaagaaagatc*GAGTTATACTGAGAGCATATGATA**AATGAAAAAAAAA**<br>**AAAAAAAAAAA**GGCAGTCTTGGACAATAGTTATATCACGTTGACTCAGGAAGTGGTGG<br>AGACTGCCACA**TATAAATA**GAGTGCCAGTAGCGACTTTTTTCACACTCGAAATACTCTTACT<br>ACTGCTCTCTTGTTGTTTTTATCACTTCTTGTTTCTTCTTGGTAAAT**AGAATATCAAGCTACA**<br>**AAAA**TAAATAAAAA |
| $P_{S.\ a16b1c0}$ | ACACCGGGCA*gaatttaggctaaagaaagatc*GTCATGAGTATATGATGAGTGATA**AATGAAAAAAAAA**<br>**AAAAAAAAAAA**GGCAGTCTTGGACAATAGTTATATCACGTTGACTCAGGAAGTGGTGG<br>AGACTGCCACA**TATAAATA**GAGTGCCAGTAGCGACTTTTTTCACACTCGAAATACTCTTACT<br>ACTGCTCTCTTGTTGTTTTTATCACTTCTTGTTTCTTCTTGGTAAAT**AGAATATCAAGCTACA**<br>**AAAA**TAAATAAAAA |
| $P_{S.\ a17b1c0}$ | ACACCGGGCA*gaatttaggctaaagaaagatc*AGGATTATGAGAAGAACCTAGTGG**AATGAAAAAAAA**<br>**AAAAAAAAAAAA**GGCAGTCTTGGACAATAGTTATATCACGTTGACTCAGGAAGTGGTGGTG<br>GAGACTGCCACA**TATAAATA**GAGTGCCAGTAGCGACTTTTTTCACACTCGAAATACTCTTAC<br>TACTGCTCTCTTGTTGTTTTTATCACTTCTTGTTTCTTCTTGGTAAAT**AGAATATCAAGCTACA**<br>**AAAA**TAAATAAAAA |
| $P_{S.\ a18b1c0}$ | ACACCGGGCA*gaatttaggctaaagaaagatc*ATAATAGGGCTACAATACAAGAAT**AATGAAAAAAAAA**<br>**AAAAAAAAAAA**GGCAGTCTTGGACAATAGTTATATCACGTTGACTCAGGAAGTGGTGG<br>AGACTGCCACA**TATAAATA**GAGTGCCAGTAGCGACTTTTTTCACACTCGAAATACTCTTACT<br>ACTGCTCTCTTGTTGTTTTTATCACTTCTTGTTTCTTCTTGGTAAAT**AGAATATCAAGCTACA**<br>**AAAA**TAAATAAAAA |
| $P_{S.\ a19b1c0}$ | ACACCGGGCA*gaatttaggctaaagaaagatc*ATAATAGGACTACACTGAGTCTAG**AATGAAAAAAAA**<br>**AAAAAAAAAAAA**GGCAGTCTTGGACAATAGTTATATCACGTTGACTCAGGAAGTGGTGGTG<br>GAGACTGCCACA**TATAAATA**GAGTGCCAGTAGCGACTTTTTTCACACTCGAAATACTCTTAC<br>TACTGCTCTCTTGTTGTTTTTATCACTTCTTGTTTCTTCTTGGTAAAT**AGAATATCAAGCTACA**<br>**AAAA**TAAATAAAAA |
| $P_{S.\ a20b1c0}$ | ACACCGGGCA*gaatttaggctaaagaaagatc*AACTAATGCATCGAATACAAGAAT**AATGAAAAAAAA**<br>**AAAAAAAAAAAA**GGCAGTCTTGGACAATAGTTATATCACGTTGACTCAGGAAGTGGTGGTG<br>GAGACTGCCACA**TATAAATA**GAGTGCCAGTAGCGACTTTTTTCACACTCGAAATACTCTTAC<br>TACTGCTCTCTTGTTGTTTTTATCACTTCTTGTTTCTTCTTGGTAAAT**AGAATATCAAGCTACA**<br>**AAAA**TAAATAAAAA |
| $P_{S.\ a21b1c0}$ | ACACCGGGCA*gaatttaggctaaagaaagatc*AACTAATGACTACACTTAACTATT**AATGAAAAAAAAA**<br>**AAAAAAAAAAA**GGCAGTCTTGGACAATAGTTATATCACGTTGACTCAGGAAGTGGTGG<br>AGACTGCCACA**TATAAATA**GAGTGCCAGTAGCGACTTTTTTCACACTCGAAATACTCTTACT<br>ACTGCTCTCTTGTTGTTTTTATCACTTCTTGTTTCTTCTTGGTAAAT**AGAATATCAAGCTACA**<br>**AAAA**TAAATAAAAA |
| $P_{S.\ a22b1c0}$ | ACACCGGGCA*gaatttaggctaaagaaagatc*AATCAACCGCTACAATTAACTATT**AATGAAAAAAAAA**<br>**AAAAAAAAAAA**GGCAGTCTTGGACAATAGTTATATCACGTTGACTCAGGAAGTGGTGG<br>AGACTGCCACA**TATAAATA**GAGTGCCAGTAGCGACTTTTTTCACACTCGAAATACTCTTACT<br>ACTGCTCTCTTGTTGTTTTTATCACTTCTTGTTTCTTCTTGGTAAAT**AGAATATCAAGCTACA**<br>**AAAA**TAAATAAAAA |
| $P_{S.\ a23b1c0}$ | ACACCGGGCA*gaatttaggctaaagaaagatc*AATCAACCCATCGAATGAGTCTAG**AATGAAAAAAAA**<br>**AAAAAAAAAAAA**GGCAGTCTTGGACAATAGTTATATCACGTTGACTCAGGAAGTGGTGGTG<br>GAGACTGCCACA**TATAAATA**GAGTGCCAGTAGCGACTTTTTTCACACTCGAAATACTCTTAC<br>TACTGCTCTCTTGTTGTTTTTATCACTTCTTGTTTCTTCTTGGTAAAT**AGAATATCAAGCTACA**<br>**AAAA**TAAATAAAAA |
| $P_{S.\ a24b1c0}$ | ACACCGGGCA*gaatttaggctaaagaaagatc*ATAATAGGTTGAACGAACAAGAAT**AATGAAAAAAAA**<br>**AAAAAAAAAAAA**GGCAGTCTTGGACAATAGTTATATCACGTTGACTCAGGAAGTGGTGGTG<br>GAGACTGCCACA**TATAAATA**GAGTGCCAGTAGCGACTTTTTTCACACTCGAAATACTCTTAC<br>TACTGCTCTCTTGTTGTTTTTATCACTTCTTGTTTCTTCTTGGTAAAT**AGAATATCAAGCTACA**<br>**AAAA**TAAATAAAAA |
| $P_{S.\ a25b1c0}$ | ACACCGGGCA*gaatttaggctaaagaaagatc*ATAATAGGGCTACAATGCATGAAC**AATGAAAAAAAA**<br>**AAAAAAAAAAAA**GGCAGTCTTGGACAATAGTTATATCACGTTGACTCAGGAAGTGGTGGTG<br>GAGACTGCCACA**TATAAATA**GAGTGCCAGTAGCGACTTTTTTCACACTCGAAATACTCTTAC<br>TACTGCTCTCTTGTTGTTTTTATCACTTCTTGTTTCTTCTTGGTAAAT**AGAATATCAAGCTACA**<br>**AAAA**TAAATAAAAA |
| $P_{S.\ a10b1c20}$ | ACACCGGGCA*gaatttaggctaaagaaagatc*ATAATAGGCATCGAATTAACTATT**AATGAAAAAAAAA**<br>**AAAAAAAAAAA**GGCAGTCTTGGACAATAGTTATATCACGTTGACTCAGGAAGTGGTGG<br>AGACTGCCACA**TATAAATA**CTTTCCCATGGTTCAGTTACCTCAGACTCCGTACATAGTATTCC<br>ATTTCATTAGGTCCTATAGTACCATATTTCTTCCTTAGTCAAAATT**AGAATATCAAGCTACAA**<br>**AAA**TAAATAAAAA |
| $P_{S.\ a20b25c18}$ | ACACCGGGCA*gaatttaggctaaagaaagatc*AACTAATGCATCGAATACAAGAAT**AATGAAAAAAAA**<br>**AAAAAAAAAAAA**GGCAGACTAGATGAAGAGCTGATACACTAGGACTCAGGAAGTGGTACA<br>CCATCGGGGCACA**TATAAATA**CCTGTGACATATGCTAGTGTCTACACGGTGGAGTTCATGTTC |

| | |
|---|---|
| | ACAATATACGTTGTTAGCTTTTTACATAGTAGACTTCTAATCTATTG**AGAATATCAAGCTACA**<br>**AAAA**TAAATAAAAA |
| $P_{S.\ a9b7c19}$ | ACACCGGGCA*gaatttaggctaaagaaagatc*CAAGGTAATTGAACGAGCATGAAC**AATGAAAAAAAAA**<br>**AAAAAAAAAAAA**GTGTCACTAGATGAAGAGATGCATACCCTTATACAGAACCCTCATAGTTG<br>TAAGTTGCACA**TATAAATA**CTGTAGGTTTCGATTCACTCTCACCTGTGGTGTTCCGTCTTTAT<br>AGTGTATTAGTTATCCGTTATTTACTTTATCTCGATGCTACTTTAA**AGAATATCAAGCTACAAA**<br>**AA**TAAATAAAAA |
| $P_{S.\ a14b5c21}$ | ACACCGGGCA*gaatttaggctaaagaaagatc*AATCAACCACTACACTACAAGAAT**AATGAAAAAAAAA**<br>**AAAAAAAAAAAA**TCGTTGCCAGAAGCTCTGTGCAATTAATGAAGGACTCATGTCACTACAC<br>CATCGGGGATGA**TATAAATA**TGGGTGTTGCTGCTGATCTCTTCACTTTCTAATCCACCTTAATT<br>CAGATATCTCTGTCATTATCATTTAGTAACTGCATCTTAACAATTT**AGAATATCAAGCTACAA**<br>**AAA**TAAATAAAAA |
| $P_{S.\ a24b23c22}$ | ACACCGGGCA*gaatttaggctaaagaaagatc*ATAATAGGTTGAACGAACAAGAAT**AATGAAAAAAAAA**<br>**AAAAAAAAAAAA**GGCAGTCTTGGACAATAGTGCAATTAATGAATACAGAACCCTCATCACC<br>ACTTGTAACACA**TATAAATA**TGGGTGTTGCTGCTGATCTCTTCACTTTCTAATCCACCTTAATT<br>CAGATATCTCTGTCATTTCTATTCGTTAACTGGCTTCTTGCTTCAA**AGAATATCAAGCTACAA**<br>**AAA**TAAATAAAAA |
| $P_{S.\ a18b8c15}$ | ACACCGGGCA*gaatttaggctaaagaaagatc*ATAATAGGGCTACAATACAAGAAT**AATGAAAAAAAAAA**<br>**AAAAAAAAAAA**AGCATAGTGTCAATTACACTGATACACTAGGTTTTAACCTACCTTCACCAC<br>TTGTAAAGGA**TATAAATA**GGCCCGCTGGAGCGACTTTTTACAATTCGAAATTCACTTCTTGTA<br>ATCCGGCCCTCTTTTTTCTTATCTTTTATTGGCTTCTTGTAACTT**AGAATATCAAGCTACAAA**<br>**AA**TAAATAAAAA |
| $P_{S.\ a22b27c16}$ | ACACCGGGCA*gaatttaggctaaagaaagatc*AATCAACCGCTACAATTAACTATT**AATGAAAAAAAAAA**<br>**AAAAAAAAAAA**GGCAGTCTTGGACAATAGCTGATACACTAGGTACAGAACCCTCATACACC<br>ATCGGGGCACA**TATAAATA**GGCCCGCTGGACTCCACTTTTTCACACTCGAATTCACTTCTTG<br>TAATCCGTCTTGTTGTTTTCTTATCTTTTATTGGCTTCTTGGTAAAT**AGAATATCAAGCTACAA**<br>**AAA**TAAATAAAAA |
| $P_{S.\ a21b11c23}$ | ACACCGGGCA*gaatttaggctaaagaaagatc*AACTAATGACTACACTTAACTATT**AATGAAAAAAAAAA**<br>**AAAAAAAAAAA**AGACTACTGGTAGGCTAGCTGTTAGTACTCCTCCTACTGCCAAACTACTAT<br>GACAGTGTTC**TATAAATA**TGGGTGTTGCTGCTGATCTCTTCACTTTCTTTACTTCCTATCTTTG<br>CGTTGTTTACTCTATATCATTTAGTAACTGCATCTTAACAATTT**AGAATATCAAGCTACAAAA**<br>**A**TAAATAAAAA |
| $P_{S.\ a12b17c24}$ | ACACCGGGCA*gaatttaggctaaagaaagatc*AACTAATGGCTACAATGAGTCTAG**AATGAAAAAAAAA**<br>**AAAAAAAAAAAA**GGCAATCGGCAAGAAAGATTTCTATCACTAATGCTCAAGAGAAGCTTCC<br>ACGGCATGTCACA**TATAAATA**TGGGTGTTGCTGCTGATCTCTTCACTTTCTTTACTTCCTATCT<br>TTGCGTTGTTTACTCTATTCTATTCGTTAACTGGCTTCTTGCTTCAA**AGAATATCAAGCTACA**<br>**AAAA**TAAATAAAAA |
| $P_{S.\ a19b26c6}$ | ACACCGGGCA*gaatttaggctaaagaaagatc*ATAATAGGACTACACTGAGTCTAG**AATGAAAAAAAAA**<br>**AAAAAAAAAAAA**GGCAGAGTGTCAATTACATTATATCACGTTGGGACTCATGTCACTAGTTG<br>TAAGTTGCACA**TATAAATA**GGCCCGCTGGACTCCACTTTTACAATTCGATCGTTTCCTCTTGT<br>AATCCGCTTTTTGTCTCTTTTCTTATCTTTTATTGGCTTCTTGTAACTT**AGAATATCAAGCTAC**<br>**AAAAA**TAAATAAAAA |
| $P_{S.\ a6b14c7}$ | ACACCGGGCA*gaatttaggctaaagaaagatc*CACCTTCTATTGGATGGGAATGAT**AATGAAAAAAAAAA**<br>**AAAAAAAAAAA**GGCAATCTGCTACAGGAACTTATTACGGCAACGGCCTGAAGCTCCGATGA<br>GCACTAATCACA**TATAAATA**GGCCCGCTGGACTCCACTTTTACAATTCGATTTCTTTTAGATT<br>CTGGTTCTTTGATATTTTTCTTTTCCGTTTTTAAAACTTAAGTATTT**AGAATATCAAGCTACAA**<br>**AAA**TAAATAAAAA |
| $P_{S.\ a7b15c8}$ | ACACCGGGCA*gaatttaggctaaagaaagatc*AACATATGAGGAAAGGCCCCAAAA**AATGAAAAAAAAA**<br>**AAAAAAAAAAAA**GGCAAAATGTAGTCCGCTTCTTGGCGAAAGGTACTGCACTTCAACCATT<br>GCACAAGATCACA**TATAAATA**GGCCCGCTGGACTCCACTTTTACAATTCGATTTCTTTTAGAT<br>TCTGGTTCTTTGATATTTATCCATTTTGAATTTACAGTCATCCAACTT**AGAATATCAAGCTACA**<br>**AAAA**TAAATAAAAA |
| $P_{S.\ a21b1c15}$ | ACACCGGGCA*gaatttaggctaaagaaagatc*AACTAATGACTACACTTAACTATT**AATGAAAAAAAAAA**<br>**AAAAAAAAAAA**GGCAGTCTTGGACAATAGTTATATCACGTTGACTCAGGAAGTGGTGGTGG<br>AGACTGCCACA**TATAAATA**GGCCCGCTGGAGCGACTTTTTACAATTCGAAATTCACTTCTTG<br>TAATCCGGCCCTCTTTTTTCTTATCTTTTATTGGCTTCTTGTAACTT**AGAATATCAAGCTACAA**<br>**AAA**TAAATAAAAA |
| $P_{S.\ a14b1c16}$ | ACACCGGGCA*gaatttaggctaaagaaagatc*AATCAACCACTACACTACAAGAAT**AATGAAAAAAAAA**<br>**AAAAAAAAAAAA**GGCAGTCTTGGACAATAGTTATATCACGTTGACTCAGGAAGTGGTGGTG<br>GAGACTGCCACA**TATAAATA**GGCCCGCTGGACTCCACTTTTTCACACTCGAATTCACTTCTT<br>GTAATCCGTCTTGTTGTTTTCTTATCTTTTATTGGCTTCTTGGTAAAT**AGAATATCAAGCTACA**<br>**AAAA**TAAATAAAAA |
| $P_{S.\ a9b1c24}$ | ACACCGGGCA*gaatttaggctaaagaaagatc*CAAGGTAATTGAACGAGCATGAAC**AATGAAAAAAAAA**<br>**AAAAAAAAAAAA**GGCAGTCTTGGACAATAGTTATATCACGTTGACTCAGGAAGTGGTGGTG<br>GAGACTGCCACA**TATAAATA**TGGGTGTTGCTGCTGATCTCTTCACTTTCTTTACTTCCTATCTT<br>TGCGTTGTTTACTCTATTCTATTCGTTAACTGGCTTCTTGCTTCAA**AGAATATCAAGCTACAA**<br>**AAA**TAAATAAAAA |

| | |
|---|---|
| $P_{S.\ a14b25c24}$ | ACACCGGGCA*gaatttaggctaaagaaagatc*AATCAACCACTACACTACAAGAAT**AATGAAAAAAAAAAAAAAAAAAAA**GGCAGACTAGATGAAGAGCTGATACACTAGGACTCAGGAAGTGGTACACCATCGGGGCACA**TATAAATA**TGGGTGTTGCTGCTGATCTCTTCACTTTCTTTACTTCCTATCTTTGCGTTGTTTACTCTATTCTATTCGTTAACTGGCTTCTTGCTTCAA**AGAATATCAAGCTACAAAAA**TAAATAAAAA |
| $P_{S.\ a21b1c15}$ | ACACCGGGCA*gaatttaggctaaagaaagatc*AACTAATGACTACACTTAACTATT**AATGAAAAAAAAAAAAAAAAAAAA**GGCAGACTAGATGAAGAGCTGATACACTAGGACTCAGGAAGTGGTACACCATCGGGGCACA**TATAAATA**GGCCCGCTGGAGCGACTTTTTACAATTCGAAATTCACTTCTTGTAATCCGGCCCTCTTTTTTCTTATCTTTTATTGGCTTCTTGTAACTT**AGAATATCAAGCTACAAAAA**TAAATAAAAA |
| $P_{S.\ a22b26c22}$ | ACACCGGGCA*gaatttaggctaaagaaagatc*AATCAACCGCTACAATTAACTATT**AATGAAAAAAAAAAAAAAAAAAAA**GGCAGAGTGTCAATTACATTATATCACGTTGGGACTCATGTCACTAGTTGTAAGTTGCACA**TATAAATA**TGGGTGTTGCTGCTGATCTCTTCACTTTCTAATCCACCTTAATTCAGATATCTCTGTCATTTCTATTCGTTAACTGGCTTCTTGCTTCAA**AGAATATCAAGCTACAAAAA**TAAATAAAAA |
| $P_{S.\ a20b25c6}$ | ACACCGGGCA*gaatttaggctaaagaaagatc*AACTAATGCATCGAATACAAGAAT**AATGAAAAAAAAAAAAAAAAAAAA**GGCAGACTAGATGAAGAGCTGATACACTAGGACTCAGGAAGTGGTACACCATCGGGGCACA**TATAAATA**GGCCCGCTGGACTCCACTTTTACAATTCGATCGTTTCCTCTTGTAATCCGCTTTTTGTCTCTTTTCTTATCTTTTATTGGCTTCTTGTAACTT**AGAATATCAAGCTACAAAAA**TAAATAAAAA |
| $P_{S.\ a12b7c16}$ | ACACCGGGCA*gaatttaggctaaagaaagatc*AACTAATGGCTACAATGAGTCTAG**AATGAAAAAAAAAAAAAAAAAAAA**GTGTCACTAGATGAAGAGATGCATACCCTTATACAGAACCCTCATAGTTGTAAGTTGCACA**TATAAATA**GGCCCGCTGGACTCCACTTTTTCACACTCGAATTCACTTCTTGTAATCCGTCTTGTTGTTTTCTTATCTTTTATTGGCTTCTTGGTAAAT**AGAATATCAAGCTACAAAAA**TAAATAAAAA |
| $P_{S.\ a14b8c19}$ | ACACCGGGCA*gaatttaggctaaagaaagatc*AATCAACCACTACACTACAAGAAT**AATGAAAAAAAAAAAAAAAAAAAA**AGCATAGTGTCAATTACACTGATACACTAGGTTTTAACCTACCTTCACCACTTGTAAAGGA**TATAAATA**CTGTAGGTTTCGATTCACTCTCACCTGTGGTGTTCCGTCTTTATAGTGTATTAGTTATCCGTTATTTACTTTATCTCGATGCTACTTTAA**AGAATATCAAGCTACAAAAA**TAAATAAAAA |
| $P_{S.\ a9b27c23}$ | ACACCGGGCA*gaatttaggctaaagaaagatc*CAAGGTAATTGAACGAGCATGAAC**AATGAAAAAAAAAAAAAAAAAAAA**GGCAGTCTTGGACAATAGCTGATACACTAGGTACAGAACCCTCATACACCATCGGGGCACA**TATAAATA**TGGGTGTTGCTGCTGATCTCTTCACTTTCTTTACTTCCTATCTTTGCGTTGTTTACTCTATATCATTTAGTAACTGCATCTTAACAATTT**AGAATATCAAGCTACAAAAA**TAAATAAAAA |
| $P_{S.\ a24b23c21}$ | ACACCGGGCA*gaatttaggctaaagaaagatc*ATAATAGGTTGAACGAACAAGAAT**AATGAAAAAAAAAAAAAAAAAAAA**GGCAGTCTTGGACAATAGTGCAATTAATGAATACAGAACCCTCATCACCACTTGTAACACA**TATAAATA**TGGGTGTTGCTGCTGATCTCTTCACTTTCTAATCCACCTTAATTCAGATATCTCTGTCATTATCATTTAGTAACTGCATCTTAACAATTT**AGAATATCAAGCTACAAAAA**TAAATAAAAA |
| $P_{4\times lexO}$ | CTTTTCGAGCTCCCTAGG*tgctgtatatactcacagca*TAACTGTATATACACCCAGGGTCTAGG*tgctgtatatactcacagca*TAACTGTATATACACCCAGGGTCTAGG*tgctgtatatactcacagca*TAACTGTATATACACCCAGGGTCTAGG*tgctgtatatactcacagca*TAACTGTATATACACCCAGGGTCTAGAGCATGTGCTTAACAGATATATAAATGGAAAAGCTGCATAACCACTTTAACTAATACTTTCAACATTTTCAGTTTGTATTACTTCTTATTCAAATGTCATAAAAGTATCAACAAAAAATAAATAAAAA |
| $P_{a0b0c0\Delta89k\_2\times lexObs}$ | ACACCGGGCA*tttagaacatatgttcg*TAACTGTATATACACCCAGGGTCTAGG*tttagaacatatgttcg*AATAAGAGAATTTCAGATTGAGAG**AATGAAAAAAAAAAAAAAAAAAAA**GGCAGAGGAGAGCATAGAAATGGGGTTCACTTTTTGGTAAAGCTATAGCATGCCTATCACA**TATAAATA**GAGTGCCAGTAGCGACTTTTTTCACACTCGAAATACTCTTACTACTGCTCTCTTGTTGTTTTTATCACTTCTTGTTTCTTCTTGGTAAAT**AGAATATCAAGCTACAAAAA**TAAATAAAAA |
| $P_{a0b0c0\Delta89k\_2\times lexOrec.sym}$ | ACACCGGGCA*tactgtatgatcatacagta*TAACTGTATATACACCCAGGGTCTAGG*tactgtatgatcatacagta*AATAAGAGAATTTCAGATTGAGAG**AATGAAAAAAAAAAAAAAAAAAAA**GGCAGAGGAGAGCATAGAAATGGGGTTCACTTTTTGGTAAAGCTATAGCATGCCTATCACA**TATAAATA**GAGTGCCAGTAGCGACTTTTTTCACACTCGAAATACTCTTACTACTGCTCTCTTGTTGTTTTTATCACTTCTTGTTTCTTCTTGGTAAAT**AGAATATCAAGCTACAAAAA**TAAATAAAAA |
| $P_{a0b0c0\Delta89k\_2\times lexOfs}$ | ACACCGGGCA*tactgtatatatatacagta*TAACTGTATATACACCCAGGGTCTAGG*ctgcacaaaggtgcac*AATAAGAGAATTTCAGATTGAGAG**AATGAAAAAAAAAAAAAAAAAAAA**GGCAGAGGAGAGCATAGAAATGGGGTTCACTTTTTGGTAAAGCTATAGCATGCCTATCACA**TATAAATA**GAGTGCCAGTAGCGACTTTTTTCACACTCGAAATACTCTTACTACTGCTCTCTTGTTGTTTTTATCACTTCTTGTTTCTTCTTGGTAAAT**AGAATATCAAGCTACAAAAA**TAAATAAAAA |
| $P_{a0b0c0\Delta89k\_2\times CIOL1}$ | ACACCGGGCA*taccactggcagtggta*TAACTGTATATACACCCAGGGTCTAGG*taccactggcagtggta*AATAAGAGAATTTCAGATTGAGAG**AATGAAAAAAAAAAAAAAAAAAAA**GGCAGAGGAGAGCATAGAAATGGGGTTCACTTTTTGGTAAAGCTATAGCATGCCTATCACA**TATAAATA**GAGTGCCAGTAGCGACTTTTTTCACACTCGAAATACTCTTACTACTGCTCTCTTGTTGTTTTTATCACTTCTTGTTTCTTCTTGGTAAAT**AGAATATCAAGCTACAAAAA**TAAATAAAAA |
| $P_{a0b0c0\Delta89k\_2\times CI434}$ | ACACCGGGCA*tacaagaaagtttgtt*TAACTGTATATACACCCAGGGTCTAGG*tacaagaaagtttgtt*AATAAGAGAATTTCAGATTGAGAG**AATGAAAAAAAAAAAAAAAAAAAA**GGCAGAGGAGAGCATA |

| | |
|---|---|
| | GAAATGGGGTTCACTTTTTGGTAAAGCTATAGCATGCCTATCACA**TATAAATA**GAGTGCCAGTAGCGACTTTTTTCACACTCGAAATACTCTTACTACTGCTCTCTTGTTGTTTTTATCACTTCTTGTTTCTTCTTGGTAAAT**AGAATATCAAGCTACAAAAA**TAAATAAAAA |
| $P_{a0b0c0\Delta89k\_2\times HKCI}$ | ACACCGGGCA*tgaaccataagttca*GTAACTGTATATACACCCAGGGTCTAGG*tgaaccataagttca*GAATAAGAGAATTTCAGATTGAGAG**AATGAAAAAAAAAAAAAAAAAAAA**GGCAGAGGAGAGCATAGAAATGGGGTTCACTTTTTGGTAAAGCTATAGCATGCCTATCACA**TATAAATA**GAGTGCCAGTAGCGACTTTTTTCACACTCGAAATACTCTTACTACTGCTCTCTTGTTGTTTTTATCACTTCTTGTTTCTTCTTGGTAAAT**AGAATATCAAGCTACAAAAA**TAAATAAAAA |
| $P_{a0b0c0\Delta89k\_2\times purO}$ | ACACCGGGCA*acgcaaacgtttgcgt*TAACTGTATATACACCCAGGGTCTAGG*acgcaaacgtttgcgt*AATAAGAGAATTTCAGATTGAGAG**AATGAAAAAAAAAAAAAAAAAAAA**GGCAGAGGAGAGCATAGAAATGGGGTTCACTTTTTGGTAAAGCTATAGCATGCCTATCACA**TATAAATA**GAGTGCCAGTAGCGACTTTTTTCACACTCGAAATACTCTTACTACTGCTCTCTTGTTGTTTTTATCACTTCTTGTTTCTTCTTGGTAAAT**AGAATATCAAGCTACAAAAA**TAAATAAAAA |
| $P_{a0b0c0\Delta89k\_2\times deoO}$ | ACACCGGGCA*tgttagaattctaaca*TAACTGTATATACACCCAGGGTCTAGG*tgttagaattctaaca*AATAAGAGAATTTCAGATTGAGAG**AATGAAAAAAAAAAAAAAAAAAAA**GGCAGAGGAGAGCATAGAAATGGGGTTCACTTTTTGGTAAAGCTATAGCATGCCTATCACA**TATAAATA**GAGTGCCAGTAGCGACTTTTTTCACACTCGAAATACTCTTACTACTGCTCTCTTGTTGTTTTTATCACTTCTTGTTTCTTCTTGGTAAAT**AGAATATCAAGCTACAAAAA**TAAATAAAAA |
| $P_{a0b0c0\Delta89k\_2\times RecApact}$ | ACACCGGGCA*tactaattaatttagt*TAACTGTATATACACCCAGGGTCTAGG*tactaattaatttagt*AATAAGAGAATTTCAGATTGAGAG**AATGAAAAAAAAAAAAAAAAAAAA**GGCAGAGGAGAGCATAGAAATGGGGTTCACTTTTTGGTAAAGCTATAGCATGCCTATCACA**TATAAATA**GAGTGCCAGTAGCGACTTTTTTCACACTCGAAATACTCTTACTACTGCTCTCTTGTTGTTTTTATCACTTCTTGTTTCTTCTTGGTAAAT**AGAATATCAAGCTACAAAAA**TAAATAAAAA |
| $P_{a0b0c0\Delta89k\_2\times lexOgs}$ | ACACCGGGCA*ggttgacacatgtcaacc*TAACTGTATATACACCCAGGGTCTAGG*ggttgacacatgtcaacc*AATAAGAGAATTTCAGATTGAGAG**AATGAAAAAAAAAAAAAAAAAAAA**GGCAGAGGAGAGCATAGAAATGGGGTTCACTTTTTGGTAAAGCTATAGCATGCCTATCACA**TATAAATA**GAGTGCCAGTAGCGACTTTTTTCACACTCGAAATACTCTTACTACTGCTCTCTTGTTGTTTTTATCACTTCTTGTTTCTTCTTGGTAAAT**AGAATATCAAGCTACAAAAA**TAAATAAAAA |
| $P_{a0b0c0\Delta89k\_2\times lexOmm}$ | ACACCGGGCA*acctaatattaataaggt*TAACTGTATATACACCCAGGGTCTAGG*acctaatattaataaggt*AATAAGAGAATTTCAGATTGAGAG**AATGAAAAAAAAAAAAAAAAAAAA**GGCAGAGGAGAGCATAGAAATGGGGTTCACTTTTTGGTAAAGCTATAGCATGCCTATCACA**TATAAATA**GAGTGCCAGTAGCGACTTTTTTCACACTCGAAATACTCTTACTACTGCTCTCTTGTTGTTTTTATCACTTCTTGTTTCTTCTTGGTAAAT**AGAATATCAAGCTACAAAAA**TAAATAAAAA |
| $P_{a0b0c0\Delta89k\_2\times lexOxa}$ | ACACCGGGCA*ttagtagtaatactactaa*TAACTGTATATACACCCAGGGTCTAGG*ttagtagtaatactactaa*AATAAGAGAATTTCAGATTGAGAG**AATGAAAAAAAAAAAAAAAAAAAA**GGCAGAGGAGAGCATAGAAATGGGGTTCACTTTTTGGTAAAGCTATAGCATGCCTATCACA**TATAAATA**GAGTGCCAGTAGCGACTTTTTTCACACTCGAAATACTCTTACTACTGCTCTCTTGTTGTTTTTATCACTTCTTGTTTCTTCTTGGTAAAT**AGAATATCAAGCTACAAAAA**TAAATAAAAA |
| $P_{4\times zifO}$ | GGGGACCAGGTGCCGTAAGGTAGTGGCGTGGGCGCCGTTAGAGCTTGACGGGGAAAGCCGGGCGTGGGCGCGAACGTGGCGAGAAAGGAAGGGAAGAGCGTGGGCGAAGCGCTTAGCGATCCTAGGGCGATCAGCGTGGGCGTAACTGTATATACACCCAGGGTCTAGAGCATGTGCTTAACAGATATATAAATGGAAAAGCTGCATAACCACTTTAACTAATACTTTCAACATTTTCAGTTTGTATTACTTCTTATTCAAATGTCATAAAAGTATCAACAAAAAATAAATAAAA |
| $P_{a0b1c0\_lexOrec.sym}$ | ACACCGGGCATAACTGTATATACACCCAGGGTCTAGG*tactgtatgatcatacagta*AATAAGAGAATTTCAGATTGAGAG**AATGAAAAAAAAAAAAAAAAAAAA**GGCAGTCTTGGACAATAGTTATATCACGTTGACTCAGGAAGTGGTGGTGGAGACTGCCACA**TATAAATA**GAGTGCCAGTAGCGACTTTTTTCACACTCGAAATACTCTTACTACTGCTCTCTTGTTGTTTTTATCACTTCTTGTTTCTTCTTGGTAAAT**AGAATATCAAGCTACAAAAA**TAAATAAAAA |
| $P_{a0b1c0\_lexObs}$ | ACACCGGGCATAACTGTATATACACCCAGGGTCTAGG*tttagaacatatgttcg*AATAAGAGAATTTCAGATTGAGAG**AATGAAAAAAAAAAAAAAAAAAAA**GGCAGTCTTGGACAATAGTTATATCACGTTGACTCAGGAAGTGGTGGTGGAGACTGCCACA**TATAAATA**GAGTGCCAGTAGCGACTTTTTTCACACTCGAAATACTCTTACTACTGCTCTCTTGTTGTTTTTATCACTTCTTGTTTCTTCTTGGTAAAT**AGAATATCAAGCTACAAAAA**TAAATAAAAA |
| $P_{a0b1c0\_lexOmm}$ | ACACCGGGCATAACTGTATATACACCCAGGGTCTAGG*acctaatattaataaggt*AATAAGAGAATTTCAGATTGAGAG**AATGAAAAAAAAAAAAAAAAAAAA**GGCAGTCTTGGACAATAGTTATATCACGTTGACTCAGGAAGTGGTGGTGGAGACTGCCACA**TATAAATA**GAGTGCCAGTAGCGACTTTTTTCACACTCGAAATACTCTTACTACTGCTCTCTTGTTGTTTTTATCACTTCTTGTTTCTTCTTGGTAAAT**AGAATATCAAGCTACAAAAA**TAAATAAAAA |
| $P_{a0b1c0\_HKCI}$ | ACACCGGGCATAACTGTATATACACCCAGGGTCTAGG*tgaaccataagttcag*AATAAGAGAATTTCAGATTGAGAG**AATGAAAAAAAAAAAAAAAAAAAA**GGCAGTCTTGGACAATAGTTATATCACGTTGACTCAGGAAGTGGTGGTGGAGACTGCCACA**TATAAATA**GAGTGCCAGTAGCGACTTTTTTCACACTCGAAATACTCTTACTACTGCTCTCTTGTTGTTTTTATCACTTCTTGTTTCTTCTTGGTAAAT**AGAATATCAAGCTACAAAAA**TAAATAAAAA |
| $P_{a0b1c0\_CIOL1}$ | ACACCGGGCATAACTGTATATACACCCAGGGTCTAGG*taccactggcggtg*ATAAATAAGAGAATTTCAGATTGAGAG**AATGAAAAAAAAAAAAAAAAAAAA**GGCAGTCTTGGACAATAGTTATATCACGTTGACTCAGGAAGTGGTGGTGGAGACTGCCACA**TATAAATA**GAGTGCCAGTAGCGACTTTTTTCACACTCGAAATACTCTTACTACTGCTCTCTTGTTGTTTTTATCACTTCTTGTTTCTTCTTG |

| | |
|---|---|
| | GTAAAT**AGAATATCAAGCTACAAAAA**TAAATAAAAA |
| *P*$_{a0b1c0\_purOa\text{-}lexObsi}$ | ACACCGGGCATAACTGTATATACACCCAGGGTCTAGG*acgcaaacgtttgcgt*AATAAGAGAATTTCAGATTGAGAG**AATGAAAAAAAAAAAAAAAAA**GGCAGTCTTGGACAATAGTTATATCACGTTGACTCAGGAAGTGGTGGTGGAGACTGCCACA**TATAAATA***agaacatatgttcg*GGGGGCCAGCAGCTTGAGAAAGTGTTTATGTTGTTACTTCTTTTTCATTCC*agaacatatgttcg***AGAATATCAAGCTACAAAAA**TAAATAAAAA |
| *P*$_{a0b1c0\_purOa\text{-}lexOxai}$ | ACACCGGGCATAACTGTATATACACCCAGGGTCTAGG*acgcaaacgtttgcgt*AATAAGAGAATTTCAGATTGAGAG**AATGAAAAAAAAAAAAAAAAA**GGCAGTCTTGGACAATAGTTATATCACGTTGACTCAGGAAGTGGTGGTGGAGACTGCCACA**TATAAATA***ttagtagtaatactactaa*GGGGGCCAGCAGCTTGAGAAAGTGTTTATGTTGTTACTTCTTTTTCATTCC*ttagtagtaatactactaa***AGAATATCAAGCTACAAAAA**TAAATAAAAA |
| *P*$_{a0b1c0\_lexOrec.syma\text{-}lexObsi}$ | ACACCGGGCATAACTGTATATACACCCAGGGTCTAGGtactgtatgatcatacagtaAATAAGAGAATTTCAGATTGAGAG**AATGAAAAAAAAAAAAAAAAA**GGCAGTCTTGGACAATAGTTATATCACGTTGACTCAGGAAGTGGTGGTGGAGACTGCCACA**TATAAATA***agaacatatgttcg*GGGGGCCAGCAGCTTGAGAAAGTGTTTATGTTGTTACTTCTTTTTCATTCC*agaacatatgttcg***AGAATATCAAGCTACAAAAA**TAAATAAAAA |

**Supplementary Table 10: Sequence of DBDs used in this study.**

| DBDs | Sequence | Source |
|---|---|---|
| lexA$_{ec87}$ | ATGAAAGCGTTAACGGCCAGGCAACAAGAGGTGTTTGATCTCATCCGTG<br>ATCACATCAGCCAGACAGGTATGCCGCCGACGCGTGCGGAAATCGCGCA<br>GCGTTTGGGGTTCCGTTCCCCAAACGCGGCTGAAGAACATCTGAAGGC<br>GCTGGCACGCAAAGGCGTTATTGAAATTGTTTCCGGCGCATCACGCGGG<br>ATTCGTCTGTTGCAGGAAGAGGAAGAAGGGTTGCCGCTGGTAGGTCGT<br>GTGGCTGCCGGTGAACCA | *E. coli* |
| lexA$_{mt127}$ | ATGTCTAATGATAGTAACGATACGAGTGTAGCGGGAGGAGCTGCGGGGG<br>CGGATTCCCGCGTATTAAGTGCAGATTCTGCGTTGACAGAACGCCAGCG<br>CACAATCTTAGACGTAATCCGCGCCTCAGTGACTTCTCGCGGCTATCCTC<br>CTTCCATTCGTGAGATCGGCGATGCCGTAGGGCTTACTAGTACGTCCTCC<br>GTTGCACACCAACTGCGCACATTGGAACGCAAAGGCTACCTGCGTCGC<br>GATCCAAACCGCCCCCGTGCGGTAAACGTTCGCGGAGCCGACGACGCA<br>GCACTTCCGCCTGTGACTGAAGTCGCGGGGTCTGATGCTCTGCCAGAGC<br>CGACATTCGTACCTGTACTGGGACGTATTGCAGCTGGCGGC | *M. tuberculosis* |
| lexA$_{cg145}$ | ATGTCTGCAATCGAAAAGAAGCCAGCAGGTGCACGAGGCAGTCGCGGT<br>AGCCGCACAGTTAAGACCTTGCCCAACGGAAAACCAGATCCAGCAAGT<br>TTGTCAGATAGGCAGCGCAGGATTTTAGAGGTTATCCGAGATGCTGTGG<br>TTTTGAGGGGTTATCCACCAAGCATTAGGGAAATTGGTGATGCTGCAGG<br>ACTTCAATCCACTTCTTCCGTTGCTTACCAGCTTAAAGAGCTAGAGAAG<br>AAGGGCTTCCTGCGCAGGGACCCTAATAAGCCTCGCGCGGTGGATGTTC<br>GCCACCTTCCAGAAACTGAAAGCCGTTCCTCCAAGGCTGCTACACAGG<br>CAAAGAGCAAGGCCCCTCAGGCCGGGGTCCATGATCCTGAGTTAGCTG<br>GCCAGACCTCATTTGTCCCAGTGGTGGGCAAAATTGCCGCTGGTAGCCC<br>G | *C. glutamicum* |
| lexA$_{sa96}$ | ATGTCTAGAGAATTAACAAAACGACAAAGCGAAATATATAACTATATTAA<br>ACAAGTTGTTCAAACGAAAGGTTATCCGCCTAGTGTTCGCGAAATTGGT<br>GAAGCAGTTGGCTTAGCATCCAGTTCAACTGTTCATGGTCACCTTTCAC<br>GTCTTGAAGAAAAAGGCTATATAAGAAGAGATCCAACGAAACCACGTG<br>CTATAGAAATTGTAAGTGATCAAACAAATGATAATATTAATATGGAAGAA<br>ACGATTCATGTGCCAGTTATTGGTAAAGTCACAGCAGGTGTTCCT | *S. aureus* |
| lexA$_{bs94}$ | ATGTCTACGAAGCTATCAAAAAGGCAACTTGATATCCTCCGTTTTATTAA<br>AGCAGAGGTTAAATCAAAAGGATATCCGCCTTCCGTGAGAGAGATCGGA<br>GAGGCTGTCGGGCTTGCGTCCAGTTCTACTGTCCACGGCCATTTGGCCC<br>GTTTGGAAACAAAAGGGCTGATCAGACGAGATCCGACAAAACCAAGAG<br>CAATAGAAATTCTTGATGAAGAAGTAGACATTCCGCAAAGCCAGGTCGT<br>TAATGTACCGGTCATCGGGAAAGTCACGGCGGGATCTCCT | *B. subtilis* |
| lexA$_{lm93}$ | ATGTCTAAGATCAGTAAGCGCCAACAGGATATTTATGAGTTTATCAAATC<br>AGAGGTTAAGGAAAAGGGTTATCCTCCGAGTGTTCGTGAAATCGGGGA<br>AGCTGTTGGTCTGGCATCTTCGTCGACGGTGCATGGTCACCTGGCACGT<br>CTTGAAGGAAAGGGGCTTATCCGCCGCGATCCAACCAAGCCCCGCGCA<br>ATCGAAATCTTGAGCCTGGAGGACGAAGCAGAGACTCCGAACGTTGTA<br>AATATCCCCATCATTGGCAAAGTGACTGCGGGGATGCCG | *L. monocytogenes* |
| lexA$_{pd114}$ | ATGTCTTATGGTAACAAGAACAAGATTTGTCTGGAAATGTCCGACGTGAT<br>GTACTTGGATTTAACCGAAAAACAGGTTTTGATCTTAGAATTTATCAAAT<br>CCCAGATTACATTGAAGGGGTACCCCCCTGCGGTACGCGAAATTTGCAC<br>GGCCGTGGGTTTACGCTCAACGTCCACGGTGCACAGTCACTTGAATAAA<br>CTGGAAAAGTTGGGTTACATCCGTAAGGACCCCACGAAACCACGCGCC<br>ATTGAAGTACTGGAACGTAGTAAGGTGAATGATGTCTCGGGGGCCAATC<br>AGGAGATTATCGAGTTGCCCTTAGTGGGGCAAATCACTGCTGGTGAGCC<br>A | *P. difficile* |
| lexA$_{sm126}$ | ATGTCTCTGACCCGCAAGCAACAGGAATTGCTTCTGTTCATTCACGAAC<br>GAATGAAGGAATCCGGGGTCCCGCCGTCCTTCGACGAGATGAAGGACG<br>CCTTGGACCTTGCGTCCAAATCGGGCATCCACCGTTTGATCACCGCACT<br>CGAAGAGCGTGGGTTCATTCGCCGGCTACCTAATCGCGCCCGGGCCCTC<br>GAAGTCATCAAGCTGCCGGAGGCATATGCAGGCGCTTCGCAGGTCCGCC<br>GCGGCTTTTCGCCGAGCGTGATCGAAGGCAGTCTGGGCAAGCCTGCTG<br>CGCCGCCCCCCGCACCGAAACCGGCGCCGCCGGCCGAGGCCGCATCGG<br>TCGCCGTTCCGGTCATGGGCCGCATTGCCGCCGGTGTGCCG | *S. meliloti* |
| lexA$_{cc123}$ | ATGTCTCTGACGCGTAAGCAGCACGAGTTGCTTATGTTTATTCACGAGCG<br>CATTAAGGAAACCGGGGTTTCACCCTCGTTCGACGAAATGAAGGAAGC<br>ATTGGATTTAGCATCGAAGTCTGGCATCCATCGCCTTATCACTGCGCTTG<br>AGGAGCGCGGCTTTATTCGCCGCTTGGCACACCGCGCACGCGCGCTTGA<br>GGTTGTCAAGTTGCCGCAGCAGGCGACTGCAGCCGCGCCACCCAAGGG | *C. crescentus* |

| | | |
|---|---|---|
| | ACGTGGTGCGTTTCGTCCACAAGTTTTTGAGGGAGGCGGAGCACCGCCTCCTGCGGCATCCCCTGCCGCTGCAGCTAATGATAGTCGTGAGCTTCCCATCTTAGGGCGTATCGCAGCAGGAACACCA | |
| lexA$_{pa93}$ | ATGTCTCAAAAACTGACTCCTCGTCAAGCAGAGATCCTGTCGTTCATTAAACGCTGTCTGGAGGACCACGGGTTTCCTCCTACGCGTGCTGAGATTGCTCAAGAGTTAGGCTTCAAATCCCCTAACGCAGCAGAGGAACATTTGAAGGCTCTTGCACGTAAAGGGGCCATTGAGATGACACCGGGCGCTTCCCGTGGTATTCGCATCCCAGGCTTCGAGCCCCACGCTGCTAATGATGACGAGGGGCTTCCCGTGATTGGACGTGTGGCCGCTGGTGCCCCA | *P. aeruginosa* |
| lexA$_{vp91}$ | ATGTCTAAGCCGTTAACGCCACGCCAACAACAAGTATTCGATTTGATTAAAAGCAAAATCGATGACACCGGAATGCCACCAACTCGTGCAGAGATTGCGCGTGAGCTTGGCTTCCGCTCTGCGAATGCCGCAGAGGAACACCTAAAAGCGCTTGCTCGTAAGCAAGCCATTGAAATTATTCCGGGCGCATCGCGTGGGATCCGAATCCTGCTTGAAGATGCGGCCAATGACGAACAGGGCTTGCCTCTGATTGGTCAAGTCGCTGCGGGTGAGCCG | *V. parahaemolyticus* |
| lexA$_{pp91}$ | ATGTCTTTGAAACTGACGCCACGCCAAGCCGAAATTCTCGCGTTCATCAAGCGCTGCCTGGAGGACAACGGCTTCCCGCCTACCCGCGCCGAGATTGCTCAGGAGCTGGGCTTCAAGTCGCCCAATGCCGCCGAGGAGCACCTCAAGGCCCTTGCCCGCAAGGGCGCGATCGAAATGACGCCGGGCGCTTCCCGCGGCATCCGCATCCCTGGCCTGGAAGCCAAGGCTGAAGAAGCCGGCCTGCCCATCATCGGCCGGGTCGCTGCCGGTGCGCCG | *P. putida* |
| CI$_{94}$ | atgTCTACAAAAAAAAAGCCCCTGACCCAAGAACAACTTGAGGACGCTCGTCGTTTGAAGGCCATTTACGAAAAGAAGAAGAACGAACTTGGATTGAGCCAGGAGTCAGTAGCTGACAAGATGGGGATGGGGCAATCTGGTGTCGGTGCCTTGTTTAATGGTATTAACGCCTTAAACGCCTATAACGCCGCGCTGTTGGCTAAAATCCTAAAGGTATCTGTAGAAGAGTTTAGTCCATCAATAGCCCGTGAGATCTACGAGATGTACGAGGCCGTTTCCATG | Enterobacteria phage lambda |
| CI434 | atgAGCATCTCATCCAGGGTTAAGTCTAAACGTATCCAATTAGGCCTAAATCAAGCAGAGCTAGCACAGAAAGTCGGTACTACTCAGCAATCAATAGAACAATTAGAGAACGGCAAAACGAAGCGTCCAAGGTTTCTTCCCGAATTAGCTTCCGCGTTAGGAGTATCAGTTGATTGGTTGTTAAATGGGACGAGTGATAGCAACGTGCGT | Bacteriophage 434 |
| HKCI$_{84}$ | ATGGTCCAGCAGAAGGAACGTGAGACGTTTTCTCAACGTCTGGCGTTAGCGTGTGATAAAGCTGGGCTGCCCTTGCATGGGAGGCAAGCAGACTTAGCCGTAAGACTTAAAGTTACGCCTAAGGCTATTAGCAAGTGGTTCAATGGTGAAAGTATACCACGTAAGGACAAAATGGAGAGCCTAGCCAGTGTATTAGGCACCACTGCCGCTTACCTTCACGGGTATGCGGACGATGACGGGATAACAGTCAAC | Bacteriophage HK022 |
| PurR$_{60}$ | atgGCAACAATAAAAGATGTAGCGAAACGAGCAAACGTTTCCACTACAACTGTGTCACACGTGATCAACAAAACACGTTTCGTCGCTGAAGAAACGCGCAACGCCGTGTGGGCAGCGATTAAAGAATTACACTACTCCCCTAGCGCGGTGGCGCGTAGCCTGAAGGTTAACCACACCAAG | *E. coli* str. K-12 substr. MG1655 |
| ArgR$_{93}$ | ATGCGAAGCTCGGCTAAGCAAGAAGAACTAGTTAAAGCATTTAAAGCATTACTTAAAGAAGAGAAATTTAGCTCCCAGGGCGAAATCGTCGCCGCGTTGCAGGAGCAAGGCTTTGACAATATTAATCAGTCTAAAGTCTCGCGGATGTTGACCAAGTTTGGTGCTGTACGTACACGCAATGCCAAAATGGAAATGGTTTACTGCCTGCCAGCTGAACTGGGTGTACCAACCACCTCCAGTCCATTGAAGAATCTGGTGCTGGATATCGACTACAACGAT | *E. coli* str. K-12 substr. MG1655 |
| DeoR$_{92}$ | ATGGAAACACGTCGCGAAGAGCGTATCGGGCAGCTGCTGCAAGAATTAAAACGCAGCGATAAGTTACATCTTAAAGACGCCGCCGCCCTGCTTGGGGTTTCGGAGATGACGATTCGTCGCGATCTGAACAACCACAGTGCGCCCGTCGTTTTGCTCGGCGGCTATATTGTTCTGGAACCGCGCAGTGCCAGCCATTACCTGTTAAGCGATCAAAAATCCCGCCTGGTGGAAGAAAAACGCCGGGCGGCAAAACTGGCTGCGACGCTGGTAGAACCC | *E. coli* str. K-12 substr. MG1655 |
| lexA$_{ac89}$ | atgtctACCAAACGTCAAAAAGAAGTTCTTGATTTCATCACCGGCTTCGTACAACGTAACGGTTATTCCCCGTCCTTTGAAGAGATAGCTCGTGGGCTAAATCTGAAGAGTCTTGCGACGGTTCACAAACACATTACGAACCTACAAAACAAAGGATTACTTGCCAGGGCTCATAACAGATCTAGGAGTTTGGACGTTCTACCTCCCCGTTCTCGTGGTAGATCCGTAGATAGGCTTCCCCTGATGGGCAGGATAGCCGCAGGCCGTCCG | *A. capsulatum ATCC 51196* |
| lexA$_{fs104}$ | atgtctGAAAATACAAACGAGAAAAGAAAAGAGATGACGGCTCGTCAGGAAGAAATATATGAATATATTAAGAAATATTCCAAAGAAAACCACATGCCGCCTACTGTGAGGGAGATCGGTAATCACTTCGACATATCCTCTACCAATGGTGTTCGTTCTATTCTTGCGGCTTTGATTAAAAAAGGATATATAAATCGTAGTCCCCGTCTGTCCCGTGGCATAGAGATCCTATCTGACGACAAAGAGTCGTCTAAAGAGGTTGCCAGTAATACGATAGAAATCCCCATCGTCGGTCGTGTGGCCGCCGGGACCCCG | *F. succinogenes* subsp. succinogenes S85 |

| | | |
|---|---|---|
| lexA$_{gs91}$ | ATGTCTGAAACCCTAACAAGTAGGCAAAGGACTGTCCTTGAATTCATTA<br>CCGCGCACGTTGATCGTCATGGTTATCCGCCGACTATGCGTGAGATTGCC<br>CGTCACCTTAACGTCAACGGCACACTAGGGGTTGCTAAACATTTAGAGG<br>CGCTGGCCAGGAAGGGCTATCTTCAGAGAGAGCCTGGAAACTCCCGTG<br>GGATAACGCTTACTGGGCAAACTAGACACACCCCGACGGTGTCATTACC<br>CGTTATCGGGGTGGTGAGAGCTGGCGTTCCC | *G. sulfurreducens* PCA |
| lexA$_{mm115}$ | atgtctACCGATAGAAAAGAGCACAAAATGAAGCCTGGTAGACGTCCCACT<br>GAAGGGATGACCCCCAGCCAGGAGAGAATCATGCAAGTCATTAGAAAT<br>GCAATAGATAGGGACGGTTTGCCGCCGACCGTGAAGGAGATAGCGGAG<br>GCTCTTGGCATGAAAACTCCCAGCGCGCACGAACAGGTACAGAAGTTA<br>GTGAAAAAGGGGTTTATTAGGAGAACCCCGCGTAAGGCTCGTTCTATAG<br>AGATAGTTGAGCAATCACCAGAAGAGGAACCCGTGGAGAAGAAACCAG<br>ACGTAGCGAGGCTTGTTCCCGTGCCTATTATTGGAGAAGTAGCGGCGGG<br>CATACCC | *M. marinus* MC-1 |
| lexA$_{mx100}$ | atgtctGAAGAGTTGACGGAGAGACAGAGAGAAATCCTATCTTTCATAGTTA<br>AGGAGACGGAAACAAGAGGATTCCCACCAACCATTCGTGAAATTGGTG<br>AACACATGGATATAAGGTCTACCAATGGGGTTAATGATCACTTGAAGGC<br>ACTTGAAAGAAAAGGATACCTAAACCGTGGTGAGCAACAGTCTCGTTC<br>ACTGGTCGCCACTAAAAGGGCAAGGTTACTGCTAGGTTTAGGAGCTAGA<br>AAGGACAGCGGTATGGTGGAGATTCCTCTACTTGGAAAAGTCGCCGCGG<br>GTGCGCCG | *M. xanthus* DK 1622 |
| lexA$_{pm92}$ | atgtctGAGGAACTTACCAAGAGACAATCCCAGGTACTTGACTTTATCAAAT<br>CTTATATGGAAAAAAACGGCTTTGCACCTAGCATAAGGGATATAATGAAG<br>CACTTCAATTTTAAGAGTCCCAGGGCTGCACATAAGCATCTAATAATTCT<br>TGAGAAGAAGGGATATATCGAAAGAAAAAATGTAAGTCGTGGAATAAA<br>GATGATGCCCAAATCTGGCGAAATCTTCGCTACTGAAACATTAGCTCCCG<br>TTTCTGGTAAGATTGCAGCGGGGGATGCC | *P. mobilis* SJ95 |
| lexA$_{sp87}$ | atgtctGAACCACTGACAAGAGCACAGAAAGAGTTATTCGACTGGCTAGTG<br>TCATACATAGACGAAACCCAACACGCACCGAGTATCAGGCAGATGATGA<br>GAGCCATGAATCTTCGTTCCCCCGCTCCTATCCAGTCCAGGCTTGAACGT<br>TTGAGAAACAAGGGTTACGTCGATTGGACCGACGGCAAGGCAAGAACG<br>CTAAGGATCTTACATCAAAAGCCAAAAGGCGTGAGTGTCATAGGCGAAC<br>TTAAAGGAGGGGAGCTG | *Synechocystis sp*. PCC 6803 |
| lexA$_{xa101}$ | GACTTAACTGACACACAGCAGGCAATATTGGCGCTAATTGCTGAACGTA<br>TTGATGCGGATGGGGTTCCCCCAAGTCAGACCGAAATCGCCCGTGCATT<br>TGGGTTCAAAGGCATAAGAGCCGCGCAATATCATTTAGAGGCGCTTGAG<br>CACGCAGGAGCGATCAGAAGAGTACCAGGTCAAGCTAGGGGCATCAGA<br>CTAGCAGGTCAAGGCGCACAGACTAGGACTGCCCCGGTAAGTGAGGTT<br>GCGAGGGATGATGTACTAAGACTTCCCGTGCTAGGAAGGGTCGCAGCG<br>GGACTGCCG | *X. axonopodis* pv. citri str. 306 |
| lexA$_{xa90}$ | atgtctCCAAGTTTACCCCCACAGAGAGCTGCTGTGTTAGCGTTCTTGCAAG<br>AACAGGCTCAAGCCGGGGTGTCTCCCTCACTGGCTGAAATCGCGCAAG<br>CATTCGGATTTGCCAGCCGTAACGCGGCCCAGAAACACGTTCAGGCTTT<br>GGCTGATGCTGGGCTAATAGAGCTACTTCCCAATCAAAAAAGGGGAATA<br>AGGCTGCCAGGCGGGGCAGGTCGTGATGCTCTGTTGGCCTTACCCGTAC<br>TTGGTAGAGTCGCCGCAGGGCTACCG | *X. axonopodis* pv. citri str. 306 |
| lexA$_{xf99}$ | atgtctAGTCTTTCCGATATTCAGCAGGCGATATTGTCCCTAATAACCAACCA<br>CATAAATGCTGACGGAGTTAGCCCCTCTCAGACAGAAATAGCGAGGGCA<br>TTTGGGTTCAAAGGAGTGAGGGCAGTACAGCACCACTTAGATGTTCTGG<br>AGCAGCAGGGCATGATAAGACGTGTGCCGAGACAGGCTCGTGGAATAA<br>GGTTGAAACACCTTACCGAAGTAGACGAGACGGCCTTGGCTTTACAATC<br>CGAAGATGTATTGCGTCTGCCGGTCTTGGGCAGGGTGGCGGCTGGGCAA<br>CCG | *X. fastidiosa* 9a5c |
| RecApact$_{104}$ | ATGCAGCTTGTTACGTCCAACATTAGGCATTTGCGTAAACATGCTGGATT<br>TACTCAGGCGCAGCTTGCCGAAAAGTTGGAGATAAAAAGATCCTTGGTA<br>GGCGCGTATGAGGAGGGTAGGGCGGAACCTAAATTGTCCACGCTAGTAA<br>ACGTCGCTAAGCTGTTCTCAGTTTCCTTAGACGACTTGGTTACACGTGA<br>CTTACAAAGAGAGGGCGCCGCTGGAGGGGCGGATAGATCATCAGGTGG<br>AAACTTAAGGGTCTTGGCGATTACCGTTGATCAAGATGAGAAGGAGAAT<br>ATCGAATTAGTCCCTTAC | *P. actiniarum* strain DSM 19842 |
| Murepc$_{82}$ | ATGAAGTCTAACTTTATCGAGAAGAACAACACGGAGAAGAGTATTTGGT<br>GCTCACCGCAGGAAATTATGGCGGCTGATGGGATGCCGGGGTCAGTCGC<br>CGGAGTGCACTACCGCGCCAATGTGCAGGGCTGGACCAAACAAAAAAA<br>AGAAGGCGTAAAGGGTGGAAAAGCTGTCGAGTATGATGTAATGTCAATG<br>CCAACTAAAGAACGCGAACAAGTGATTGCCCATCTGGGCCTTTCTACGC<br>CT | *Escherichia* phage Mu |
| RutR$_{86}$ | ATGACGCAAGGCGCAGTGAAAACAACGGGTAAACGTTCGCGCGCAGTA | *E. coli* str. K-12 substr. |

| | | |
|---|---|---|
| | AGCGCGAAGAAAAAAGCGATTCTTAGCGCAGCACTGGACACTTTTTCACAATTCGGTTTTCACGGCACAAGGCTGGAGCAGATCGCAGAGTTGGCGGGTGTTTCAAAAACCAATCTGCTGTATTACTTTCCGTCAAAAGAGGCGCTGTATATTGCCGTGCTGCGGCAGATTCTCGATATCTGGCTGGCACCGTTAAAAGCGTTTCGTGAA | MG1655 |
| $BetI_{80}$ | ATGCCCAAATTGGGGATGCAGTCGATCCGGCGCAGACAACTGATCGACGCCACACTGGAAGCAATAAATGAAGTGGGCATGCACGATGCAACGATCGCGCAGATCGCCCGCCGTGCAGGCGTTTCTACGGGGATCATCAGCCACTATTTCAGGGACAAAAATGGTCTGCTGGAAGCAACCATGCGCGATATCACCAGTCAGCTGCGTGACGCGGTTTTGAATCGATTACATGCACTTCCG | *E. coli* str. K-12 substr. MG1655 |
| $ArgP_{98}$ | ATGAAACGCCCGGACTACAGAACATTACAGGCACTGGATGCGGTGATACGTGAACGAGGATTTGAGCGCGCGGCACAAAAGCTGTGCATTACACAATCAGCCGTCTCACAGCGCATTAAGCAACTGGAAAATATGTTCGGGCAGCCGCTGTTGGTGCGTACCGTACCGCCGCGCCCGACGGAACAAGGGCAAAAACTGCTGGCACTGCTGCGCCAGGTGGAGTTGCTGGAAGAAGAGTGGCTGGGCGATGAACAAACCGGTTCGACTCCGCTGCTGCTTTCACTGGCGGTCAAC | *E. coli* str. K-12 substr. MG1655 |
| $FruR_{70}$ | ATGAAACTGGATGAAATCGCTCGGCTGGCGGGAGTGTCGCGGACCACTGCAAGCTATGTTATTAACGGCAAAGCGAAGCAATACCGTGTGAGCGACAAAACCGTTGAAAAAGTCATGGCTGTGGTGCGTGAGCACAATTACCACCCGAACGCCGTGGCAGCTGGGCTTCGTGCTGGACGCACACGTTCTATTGGTCTTGTGATCCCCGAT | *E. coli* str. K-12 substr. MG1655 |
| $RbsR_{67}$ | ATGGCTACAATGAAAGATGTTGCCCGCCTGGCGGGCGTTTCTACCTCAACAGTTTCTCACGTTATCAATAAAGATCGCTTCGTCAGTGAAGCGATTACCGCCAAAGTTGAAGCGGCGATTAAAGAACTCAATTACGCGCCATCAGCTCTGGCGCGTAGCCTCAAACTCAATCAAACACATACCATTGGCATGTTGATCACT | *E. coli* str. K-12 substr. MG1655 |
| $TreR_{70}$ | ATGCAAAATCGGCTGACCATCAAAGATATCGCGCGCTTAAGCGGCGTGGGGAAATCTACAGTTTCCCGGGTGCTGAATAACGAAAGCGGCGTGAGCCAGCTCACCCGCGAGCGTGTTGAAGCAGTGATGAATCAGCATGGATTTTCCCCTTCCCGCTCTGCGCGCGCTATGCGTGGGCAAAGCGATAAAGTGGTCGCCATCATTGTTACC | *E. coli* str. K-12 substr. MG1655 |
| $NanR_{107}$ | ATGGGCCTTATGAACGCATTTGATTCGCAAACCGAAGATTCTTCACCTGCAATTGGTCGCAACTTGCGTAGCCGCCCGCTGGCGCGTAAAAAACTCTCCGAAATGGTGGAAGAAGAGCTGGAACAGATGATCCGCCGTCGTGAATTTGGCGAAGGTGAACAATTACCGTCTGAACGCGAACTGATGGCGTTCTTTAACGTCGGGCGTCCTTCGGTGCGTGAAGCGCTGGCAGCGTTAAAACGCAAAGGTCTGGTGCAAATAAACAACGGCGAACGCGCTCGCGTCTCGCGTCCTTCTGCGGACACTATCATCGGTGAGCTT | *E. coli* str. K-12 substr. MG1655 |
| $PdhR_{83}$ | ATGGCCTACAGCAAAATCCGCCAACCAAAACTCTCCGATGTGATTGAGCAGCAACTGGAGTTTTTGATCCTCGAAGGCACTCTCCGCCCGGGCGAAAAACTCCCACCGGAACGCGAACTGGCAAAACAGTTTGACGTCTCCCGTCCCTCCTTGCGTGAGGCGATTCAACGTCTCGAAGCGAAGGGCTTGTTGCTTCGTCGCCAGGGTGGCGGCACTTTTGTCCAGAGCAGCCTATGGCAAAGCTTCAGC | *E. coli* str. K-12 substr. MG1655 |
| $DgoR_{89}$ | ATGACTCTCAATAAAACCGATCGCATTGTCATTACGCTGGGTAAACAGATCGTTCACGGCAAATACGTGCCAGGCTCGCCGCTTCCGGCTGAGGCGGAACTCTGTGAGGAGTTTGCAACCTCGCGCAACATCATCCGTGAGGTGTTCCGTTCGCTGATGGCGAAGCGGCTGATTGAAATGAAACGTTATCGCGGGGCGTTTGTGGCACCGCGTAACCAGTGGAATTACCTCGACACTGACGTACTGCAATGGGTGCTGGAAAATGAC | *E. coli* str. K-12 substr. MG1655 |
| $AllR_{96}$ | ATGACGGAAGTTAGACGGCGCGGCAGGCCAGGACAGGCGGAGCCTGTGGCACAGAAGGGCGCACAGGCGTTAGAGCGGGGAATTGCGATTCTGCAATATTTGGAAAAAAGTGGGGGAAGTTCGTCGGTTAGCGATATTTCTCTCAATCTGGATTTGCCGCTCTCCACGACCTTTCGCTTGCTGAAGGTTTTACAGGCAGCGGATTTTGTCTATCAGGACAGTCAATTAGGCTGGTGGCATATAGGATTAGGTGTCTTTAACGTCGGTGCGGCGTACATCCATAACCGC | *E. coli* str. K-12 substr. MG1655 |
| $MhpR_{85}$ | ATGCAGAACAATGAGCAGACGGAATACAAAACCGTGCGCGGCTTAACCCGCGGTCTAATGTTATTAAATATGTTAAATAAACTTGATGGCGGTGCCAGCGTCGGGCTGCTGGCGGAACTCAGCGGCCTGCATCGCACCACTGTGCGGCGACTGCTGGAGACGCTGCAGGAAGAGGGATATGTCCGCCGTAGCCCCTCCGATGATAGTTTTCGACTGACCATCAAAGTGCGGCAATTAAGCGAAGGATTTCGTGAC | *E. coli* str. K-12 substr. MG1655 |
| DBP35opt | atgGGTTTCGGTCGTGCGGTAAAAGAGAAGAGAAAAGAACTGGGACTGACGCAGGTGGAGTTCGCTGAAAAGGCAGGTTTATCCAGAAGAACAATCATTAACATAGAGAGAGGGTATATCGTCCCACAAAAAGCTACTAAGGAGAAAATAGCCAAGGCTTTAGGGACAAGCGTTGAGGAGCTTGAACAGGCA | [3] |

| | |
|---|---|
| DBP57 | atGATGGTCTTGACGCCTATGGAGAGAATAGGTGAGTTTATTAAAAGAGC<br>AAGGCGTGAAGCAGGTCTGACTCAGCGTGAGCTTGCGGAGTTAGCGGG<br>AGTCGGGCAGTCTACAGTATCAAGAATAGAGAAGGGTGAGAAGTGTTC<br>ACCAGAATTGGTAGAGAAGATTCTGGAAGCCCTGAGGAAAGTG |
| DBP60 | atGGACATAGAAAAAATTGCAAAAGCTGTAAAAGAGTTGAGGGAAGAGC<br>TTGGACTAACTCAGGCAGAATTTGCCAAGAAAATTGGAATTGGGCAAG<br>GCACGTTGAGTAGGTTTGAAAAGGGCGGCGTTCTTTCACCTAAAACGAT<br>GGAAAGACTGTTAAAAGCACTAGAAAAAGAGTTCGGTTTTGACGTGAA<br>GAAG |
| yEmCitrine | ATGTCTAAAGGTGAAGAATTATTCACTGGTGTTGTCCCAATTTTGGTTGA<br>ATTAGATGGTGATGTTAATGGTCACAAATTTTCTGTCTCCGGTGAAGGTG<br>AAGGTGATGCTACTTACGGTAAATTGACCTTAAAATTTATTTGTACTACTG<br>GTAAATTGCCAGTTCCATGGCCAACCTTAGTCACTACTTTAGGTTATGGT<br>TTGATGTGTTTTGCTAGATACCCAGATCATATGAAACAACATGACTTTTTC<br>AAGTCTGCCATGCCAGAAGGTTATGTTCAAGAAAGAACTATTTTTTTCA<br>AAGATGACGGTAACTACAAGACCAGAGCTGAAGTCAAGTTTGAAGGTG<br>ATACCTTAGTTAATAGAATCGAATTAAAAGGTATTGATTTTAAAGAAGAT<br>GGTAACATTTTAGGTCACAAATTGGAATACAACTATAACTCTCACAATGT<br>TTACATCATGGCTGACAAACAAAAGAATGGTATCAAAGTTAACTTCAAA<br>ATTAGACACAACATTGAAGATGGTTCTGTTCAATTAGCTGACCATTATCA<br>ACAAAATACTCCAATTGGTGATGGTCCAGTCTTGTTACCAGACAACCATT<br>ACTTATCCTATCAATCTAGATTATCCAAAGATCCAAACGAAAAGAGGGAT<br>CACATGGTCTTGTTAGAATTTGTTACTGCTGCTGGTATTACCCATGGTATG<br>GATGAATTGTACAAATAA |

#### Supplementary Table 11: sequence of LBDs used in this study.

| LBDs | Sequence | Source |
|---|---|---|
| LasR$_{177}$ | GCCTTGGTTGACGGTTTTCTTGAGCTGGAACGCTCAAGTGGAAAATTGG<br>AGTGGAGCGCCATCCTGCAGAAGATGGCGAGCGACCTTGGATTCTCGA<br>AGATCCTGTTCGGCCTGTTGCCTAAGGACAGCCAGGACTACGAGAACGC<br>CTTCATCGTCGGCAACTACCCGGCCGCCTGGCGCGAGCATTACGACCGG<br>GCTGGCTACGCGCGGGTCGACCCGACGGTCAGTCACTGTACCCAGAGC<br>GTACTGCCGATTTTCTGGGAACCGTCCATCTACCAGACGCGAAAGCAGC<br>ACGAGTTCTTCGAGGAAGCCTCGGCCGCCGGCCTGGTGTATGGGCTGAC<br>CATGCCGCTGCATGGTGCTCGCGGCGAACTCGGCGCGCTGAGCCTCAGC<br>GTGGAAGCGGAAAACCGGGCCGAGGCCAACCGTTTCATGGAGTCGGTC<br>CTGCCGACCCTGTGGATGCTCAAGGACTACGCACTGCAGAGCGGTGCC<br>GGACTGGCCTTCGAACATCCGGTCAGCAAACCGGTGGTTCTG | *P. aeruginosa* |
| RpaR$_{179}$ | ATTGTGGGTGAAGATCAGCTGTGGGGTCGTCGTACACTGGAATTTGTTG<br>ATAGCGTTGAACGTCTGGAAGCACCGGCACTGATTAGCCGTTTTGAAAG<br>CCTGATTGCAAGCTGTGGTTTTACCGCCTATATCATGGCAGGTCTGCCGA<br>GCCGTAATGCCGGTCTGCCGGAACTGACCCTGGCAAATGGTTGGCCTCG<br>TGATTGGTTTGATCTGTATGTTAGCGAAAACTTTAGCGCAGTTGATCCGG<br>TTCCGCGTTATGGTGCAACCACTGTTCATCCGTTTGTTTGGAGTGATGCA<br>CCGTATGATCGTGACCGTGATCAGGCAGCACATCGTGTTATGACCCGTGC<br>AGCAGAATTTGGTCTGGTTGAAGGTTATTGTATTCCGCTGCATTACGATG<br>ATGGTAGCGCAGCAATTAGTATGGCAGGTGAAGATCCTGATCTGAGTCC<br>GGCAGCCCGTGGTGTAATGCAGCTGGTTAGCATTTATGCACATAGCCGTC<br>TGCGTGTACTGAGCCGTCCGAAACCGATTCGTCGTAAT | *R. palustris* |
| ER$_{282\text{-}595}$ | TCTGCTGGAGACATGAGAGCTGCCAACCTTTGGCCAAGCCCGCTCATGA<br>TCAAACGCTCTAAGAAGAACAGCCTGGCCTTGTCCCTGACGGCCGACC<br>AGATGGTCAGTGCCTTGTTGGATGCTGAGCCCCCCATACTCTATTCCGAG<br>TATGATCCTACCAGACCCTTCAGTGAAGCTTCGATGATGGGCTTACTGAC<br>CAACCTGGCAGACAGGGAGCTGGTTCACATGATCAACTGGGCGAAGAG<br>GGTGCCAGGCTTTGTGGATTTGACCCTCCATGATCAGGTCCACCTTCTAG<br>AATGTGCCTGGTTAGAGATCCTGATGATTGGTCTAGTCTGGCGCTCCATG<br>GAGCACCCAGTGAAGCTACTGTTTGCTCCTAACTTGCTCTTGGACAGGA<br>ACCAGGGAAAATGTGTAGAGGGCATGGTGGAGATCTTCGACATGCTGCT<br>GGCTACATCATCTCGGTTCCGCATGATGAATCTGCAGGGAGAGGAGTTT<br>GTGTGCCTCAAATCTATTATTTTGCTTAATTCTGGAGTGTACACATTTCTG<br>TCCAGCACCCTGAAGTCTCTGGAAGAGAAGGACCATATCCACCGAGTCC<br>TGGACAAGATCACAGACACTTTGATCCACCTGATGGCCAAGGCAGGCCT<br>GACCCTGCAGCAGCAGCACCAGCGGCTGGCCCAGCTCCTCCTCATCCTC<br>TCCCACATCAGGCACATGAGTAACAAAGGCATGGAGCATCTGTACAGCA<br>TGAAGTGCAAGAACGTGGTGCCCCTCTATGACCTGCTGCTGGAGATGCT<br>GGACGCCCACCGCCTACATGCGCCCACTAGCCGTGGAGGGGCATCCGTG<br>GAGGAGACGGACCAAAGCCACTTGGCCACTGCGGGCTCTACTTCATCG<br>CATTCCTTGCAAAAGTATTACATCACGGGGGAGGCAGAGGGTTTCCCTG<br>CCACAGTC | Homo sapiens |
| DHBR$_{282\text{-}595}$ | TCTGCTGGAGACATGAGAGCTGCCAACCTTTGGCCAAGCCCGCTCATGA<br>TCAAACGCTCTAAGAAGAACAGCCTGGCCTTGTCCCTGACGGCCGACC<br>AGATGGTCAGTGCCTTGTTGGATGCTGAGCCCCCCATACTCTATTCCGAG<br>TATGATCCTACCAGACCCTTCAGTGAAGCTTCGATGATGGGCTTAATCAC<br>CAACCTGATGGACAGGGAGCTGGTTCACATGATCAACTGGGCGAAGAG<br>GGTGCCAGGCTTTGTGGATTTGACCCTCCATGATCAGGTCCACCTTCTAG<br>AATGTGCCTGGTTAGAGATCCTGCAGATTGGTCTAGTCTGGCGCTCCATG<br>GAGCACCCAGTGAAGCTACTGTTTGCTCCTAACTTGCTCTTGGACAGGA<br>ACCAGGGAAAATGTGTAGAGGGCATGGTGGAGATCTTCGACATGCTGCT<br>GGCTACATCATCTCGGTTCCGCATGATGAATCTGCAGGGAGAGGAGTTT<br>GTGTGCCTCAAATCTATTATTTTGCTTAATTCTGGAGTGTACACATTTCTG<br>TCCAGCACCCTGAAGTCTCTGGAAGAGAAGGACCATATCCACCGAGTCC<br>TGGACAAGATCACAGACACTTTGATCCACCTGATGGCCAAGGCAGGCCT<br>GACCCTGCAGCAGCAGCACCAGCGGCTGGCCCAGCTCCTCCTCATCCTC<br>TCCCACATCAGGCACATGAGTAACAAAAGCATGGAGCATCTGGACAGC<br>ATGAAGTGCAAGAACGTGGTGCCCCTCTATGACCTGCTGCTGGAGATGC<br>TGGACGCCCACCGCCTACATGCGCCCACTAGCCGTGGAGGGGCATCCGT<br>GGAGGAGACGGACCAAAGCCACTTGGCCACTGCGGGCTCTACTTCATC<br>GCATTCCTTGCAAAAGTATTACATCACGGGGGAGGCAGAGGGTTTCCCT<br>GCCACAGTC | Homo sapiens |
| acVHH | GGCTCTCAGGTTCAATTGGTGGAATCTGGAGGGGGTCTAGTACAGGCAG | Lama glama |

| | | |
|---|---|---|
| | GCGGTTCTCTCCGACTGAGTTGCACAGCCTCCGGTAGGACTGGGACCAT<br>CTACTCAATGGCCTGGTTTCGCCAGGCCCCAGGCAAAGAAAGAGAGTT<br>TCTTGCCACTGTAGGTTGGAGTTCTGGGATCACATACTACATGGATTCAG<br>TTAAAGGAAGATTCACTATCAGCCGAGATAAAGGGAAAAATACTGTGTA<br>CCTCCAGATGGACTCTCTGAAACCGGAGGACACGGCTGTCTACTACTGT<br>ACAGCCACCCGCGCCTGGTCCGTAGGGTATGACTACTGGGGGCAAGGA<br>ACACAGGTAACCGTCTCTAGC | |
| CinR$_{179}$ | ATTGAGAATACCTATAGCGAAAAGTTCGAGTCCGCGTTCGAACAGATCA<br>AGGCGGCGGCCAACGTGGATGCCGCCATCCGTATTCTCCAGGCGGAATA<br>TAACCTCGATTTCGTCACCTACCATCTCGCCCAGACGATCGCGAGCAAG<br>ATCGATTCGCCCTTCGTGCGCACCACCTATCCGGATGCCTGGGTTTCCCG<br>CTACCTCCTCAACAGCTATGTGAAGGTCGATCCGATCGTCAAGCAGGGC<br>TTCGAACGCCAGCTGCCCTTCGACTGGAGCGAGGTCGAACCGACGCCG<br>GAGGCCTATGCCATGCTGGTCGACGCCCAGAAACACGGCATCGGTGGC<br>AATGGCTACTCCATCCCCGTCGCCGACAAGGCGCAGCGCCGCGCCCTGC<br>TGTCGCTGAATGCCCGTATACCGGCCGACGAATGGACCGAGCTTGTGCG<br>CCGCTGCCGCAACGAGTGGATCGAGATCGCCCATCTGATCCACCGCAAG<br>GCCGTCTATGAGCTGCATGGCGAAAACGATCCGGTGCCGGCATTG | *R. leguminosarum* |
| PR | GCTGGTCAGGGTGACCCGGTTCGTGTTGTTCGTGCTCTGGACGCTGTTG<br>CTCTGCCGCAGCCGGTTGGTGTTCCGAACGAATCTCAGGCTCTGTCTCA<br>GCGTTTCACCTTCTCTCCGGGTCAGGACATCCAGCTGATCCCGCCGCTG<br>ATCAACCTGCTGATGTCTATCGAACCGGACGTTATCTACGCTGGTCACGA<br>CAACACCAAACCGGACACCTCTTCTTCTCTGCTGACCTCTCTGAACCAG<br>CTGGGTGAACGTCAGCTGCTGTCTGTTGTTAAATGGTCTAAATCTCTGCC<br>GGGTTTCCGTAACCTGCACATCGACGACCAGATCACCCTGATCCAGTAC<br>TCTTGGATGTCTCTGATGGTTTTCGGTCTGGGTTGGCGTTCTTACAAACA<br>CGTTTCTGGTCAGATGCTGTACTTCGCTCCGGACCTGATCCTGAACGAA<br>CAGCGTATGAAAGAATCTTCTTTCTACTCTCTGTGCCTGACCATGTGGCA<br>GATCCCGCAGGAATTCGTTAAACTGCAGGTTTCTCAGGAAGAATTCCTG<br>TGCATGAAAGTTCTGCTGCTGCTGAACACCATCCCGCTGGAAGGTCTGC<br>GTTCTCAGACCCAGTTCGAAGAAATGCGTTCTTCTTACATCCGTGAACT<br>GATCAAAGCTATCGGTCTGCGTCAGAAAGGTGTTGTTTCTTCTTCTCAGC<br>GTTTCTACCAGCTGACCAAACTGCTGGACAACCTGCACGACCTGGTTAA<br>ACAGCTGCACCTGTACTGCCTGAACACCTTCATCCAGTCTCGTGCTCTG<br>TCTGTTGAATTCCCGGAAATGATGTCTGAAGTTATCGCTGGTTCTACCGC<br>TTCTGGTCGTGCTGGTGCTCCGCCGACCGACGTTTCTCTGGGTGACGAA<br>CTGCACCTGGACGGTGAGGACGTTGCTATGGCTCAC | Homo sapiens |
| BjaR$_{180}^{S107R}$ | AGTGCGGTGGATTATGGCCGTGAAGCCCTGGACTTTATCGAAGGCCTGG<br>GTGTCTACCGCAAAGTGCCGGATGCTATGAACGCGCTGGAAGCGGCCTT<br>TGGCCGTTTTGGTTTCGAAACCATTATCGTCACGGGCCTGCCGAACCCG<br>GATCAGCGTTTCGCTCAAATGGTGCTGGCAAAACGTTGGCCGGCCGGTT<br>GGTTTAATCTGTATACCCAGAACAATTACGATCGTTTCGACCCGGTGGTT<br>CGTCTGTGCCGCCAAAGTGTTAACCCGTTTGAATGGTCCGAAGCCCCGT<br>ATGATGCAGAACTGGAACCGCGTGCAGCTGAAGTTATGAATCGCGCAGG<br>CGACTTTCGTATGTCTCGCGGTTTCATTGTCCCGATCCATGGCCTGACCG<br>GTTACGAAGCGGCCGTTTCACTGGGCGGTGTCCACCTGGATCTGAATCC<br>GCGTTCGAAACCGGCACTGCATCTGATGGCTATGTATGGCTTCGACCAC<br>ATCCGTCGCCTGCTGGAACCGACGCCGTACCCGTCCACCCGTCTG | *B. diazoefficiens* |
| LuxR$_{183}^{S116Y}$ | AAAAACATAAATGCCGACGACACATACAGAATAATTAATAAAATTAAAG<br>CTTGTAGAAGCAATAATGATATTAATCAATGCTTATCTGATATGACTAAAA<br>TGGTACATTGTGAATATTATTTACTCGCGATCATTTATCCTCATTCTATGGT<br>TAAATCTGATATTTCAATCCTAGATAATTACCCTAAAAAATGGAGGCAATA<br>TTATGATGACGCTAATTTAATAAAATATGATCCTATAGTAGATTATTCTAAC<br>TCCAATCATTCACCAATTAATTGGAATATATTTGAAAACAATGCTGTAAAT<br>AAAAAATCTCCAAATGTAATTAAAGAAGCGAAAACATATGGTCTTATCAC<br>TGGGTTTAGTTTCCCTATTCATACGGCTAACAATGGCTTCGGAATGCTTA<br>GTTTTGCACATTCAGAAAAAGACAACTATATAGATAGTTTATTTTTACATG<br>CGTGTATGAACATACCATTAATTGTTCCTTCTCTAGTTGATAATTATCGAA<br>AAATAAATATAGCAAATAATAAATCAAACAACGATTTA | *A. fischeri* |
| TraR$_{174}$ | ATGCAGCACTGGCTGGACAAGCTGACTGATCTTGCCGCGATCGAAGGCG<br>ATGAGTGCATCCTcAAGACCGGGCTGGCGGACATCGCCGACCATTTCGG<br>CTTCACCGGCTATGCCTACCTTCATATCCAGCACAGGCACATCACCGCCG<br>TTACCAACTATCACCGCCAATGGCAATCAACCTACTTCGACAAGAAGTT<br>CGAAGCGCTCGATCCGGTCGTCAAACGCGCGAGGTCCCGGAAGCACAT<br>CTTCACCTGGTCGGGCGAGCACGAGCGGCCGACGCTGTCGAAGGACGA<br>GCGTGCCTTCTATGACCACGCATCCGATTTCGGCATCCGCTCCGGCATCA<br>CAATACCCATCAAGACCGCCAACGGCTTTATGTCGATGTTCACGATGGCA | *A. tumefaciens* |

| | | |
|---|---|---|
| | TCGGACAAGCCGGTGATCGATCTCGATCGGGAGATCGATGCAGTCGCAG<br>CCGCTGCAACCATCGGGCAGATCCATGCCCGCATCTCATTCCTTCGCACC<br>ACCCCTACCGCGGAAGATGCCGCATGGCTC | |
| $GR_{487\text{-}777}$ | AACCTGGAAGCTCGAAAAACAAAGAAAAAAATAAAAGGAATTCAGCA<br>GGCCACTACAGGAGTCTCACAAGAAACCTCTGAAAATCCTGGTAACAA<br>AACAATAGTTCCTGCAACGTTACCACAACTCACCCCTACCCTGGTGTCA<br>CTGTTGGAGGTTATTGAACCTGAAGTGTTATATGCAGGATATGATAGCTC<br>TGTTCCAGACTCAACTTGGAGGATCATGACTACGCTCAACATGTTAGGA<br>GGGCGGCAAGTGATTGCAGCAGTGAAATGGGCAAAGGCAATACCAGGT<br>TTCAGGAACTTACACCTGGATGACCAAATGACCCTACTGCAGTACTCCT<br>GGATGTTTCTTATGGCATTTGCTCTGGGGTGGAGATCATATAGACAACCA<br>AGTGCAAACCTGCTGTGTTTTGCTCCTGATCTGATTATTAATGAGCAGAG<br>AATGACTCTACCCTGCATGTACGACCAATGTAAACACATGCTGTATGTTT<br>CCTCTGAGTTACACAGGCTTCAGGTATCTTATGAAGAGTATCTCTGTATG<br>AAAACCTTACTGCTTCTCTCTTCAGTTCCTAAGGACGGTCTGAAGAGCC<br>AAGAGCTATTTGATGAAATTAGAATGACCTACATCAAAGAGCTAGGAAA<br>AGCCATTGTCAAGAGGGAAGGAAACTCCAGCCAGAACTGGCAGCGGTT<br>TTATCAACTGACAAAACTCTTGGATTCTATGCATGAAGTGGTTGAAAATC<br>TCCTTAACTATTGCTTCCAAACATTTTTGGATAAGACCATGAGTATTGAAT<br>TCCCCGAGATGTTAGCTGAAATCATCACCAATCAGATACCAAAATATTCA<br>AATGGAAATATCAAAAAACTTCTGTTTCATCAAAAG | Homo sapiens |
| $AR_{669\text{-}984}$ | AATTTAGGAGCACGAAAGTCAAAGAAGTTGGGAAAGTTAAAAGGGATT<br>CACGAGGAGCAGCCACAGCAGCAGCAGCCCCCACCCCCACCCCCACCC<br>CCGCAAAGCCCAGAGGAAGGGACAACGTACATCGCTCCTGCAAAAGAA<br>CCCTCGGTCAACACAGCACTGGTTCCTCAGCTCTCCACAATCTCACGAG<br>CGCTCACACCTTCCCCCGTTATGGTCCTTGAAAACATTGAACCTGAAATT<br>GTATATGCAGGCTATGACAGCTCAAAACCAGATACAGCCGAAAATCTGC<br>TCTCCACGCTCAACCGCTTAGCAGGCAAACAGATGATCCAAGTCGTGAA<br>GTGGGCAAAGGTACTTCCAGGATTTAAAAACTTGCCTCTTGAGGACCAA<br>ATTACCCTAATCCGATATTCTTGGATGTGTCTATCATCATTTGCCTTGAGCT<br>GGAGATCGTACAAACATACGAACAGCCAATTTCTCTATTTTGCACCAGA<br>CCTAGTCTTTAATGAAGAGAAGATGCATCAGTCTGCCATGTATGAACTAT<br>GCCAGGGGATGCACCAAATCAGCCTTCAGTTCGTTCGACTGCAGCTCAC<br>CTTTGAAGAATACACCATCATGAAAGTTTTGCTGCTACTAAGCACAATTC<br>CAAAGGATGGCCTCAAAAGCCAGGCTGCATTTGAAGAAATGAGGACAA<br>ATTACATCAAAGAACTGAGGAAGATGGTAACTAAGTGTCCCAACAATTC<br>TGGGCAGAGCTGGCAGAGGTTCTACCAACTGACCAAGCTGCTGGACTC<br>CATGCATGACCTGGTGAGCGACCTGCTGGAATTCTGCTTCTACACCTTCC<br>GAGAGTCCCATGCGCTGAAGGTAGAGTTCCCCGCAATGCTGGTGGAGAT<br>CATCAGCGACCAGCTGCCCAAGGTGGAGTCGGGGAACGCCAAGCCGCT<br>CTACTTCCACCGGAAG | Homo sapiens |
| $AR_{630\text{-}918}$ | CTGAAGAAACTTGGTAATCTGAAACTACAGGAGGAAGGAGAGGCTTCC<br>AGCACCACCAGCCCCACTGAGGAGACAACCCAGAAGCTGACAGTGTCA<br>CACATTGAAGGCTATGAATGTCAGCCCATCTTTCTGAATGTCCTGGAAGC<br>CATTGAGCCAGGTGTAGTGTGTGCTGGACACGACAACAACCAGCCCGA<br>CTCCTTTGCAGCCTTGCTCTCTAGCCTCAATGAACTGGGAGAGAGACAG<br>CTTGTACACGTGGTCAAGTGGGCCAAGGCCTTGCCTGGCTTCCGCAACT<br>TACACGTGGACGACCAGATGGCTGTCATTCAGTACTCCTGGATGGGGCT<br>CATGGTGTTTGCCATGGGCTGGCGATCCTTCACCAATGTCAACTCCAGG<br>ATGCTCTACTTCGCCCCTGATCTGGTTTTCAATGAGTACCGCATGCACAA<br>GTCCCGGATGTACAGCCAGTGTGTCCGAATGAGGCACCTCCCTCAAGAG<br>TTTGGATGGCTCCAAATCACCCCCCAGGAATTCCTGTGCATGAAAGCAC<br>TGCTACTCTTCAGCATTATTCCAGTGGATGGGCTGAAAAATCAAAAATTC<br>TTTGATGAACTTCGAATGAACTACATCAAGGAACTCGATCGTATCATTGC<br>ATGCAAAAGAAAAAATCCCACATCCTGCTCAAGACGCTTCTACCAGCTC<br>ACCAAGCTCCTGGACTCCGTGCAGCCTATTGCGAGAGAGCTGCATCAGT<br>TCACTTTTGACCTGCTAATCAAGTCACACATGGTGAGCGTGGACTTTCC<br>GGAAATGATGGCAGAGATCATCTCTGTGCAAGTGCCCAAGATCCTTTCT<br>GGGAAAGTCAAGCCCATCTATTTCCACACCCAG | Homo sapiens |
| $CepR_{176}$ | ATGGAACTGCGTTGGCAGGATGCGTATCAGCAATTTAGCGCGGCGGAAG<br>ATGAACAGCAACTGTTCCAACGTATCGCAGCTTACTCCAAACGCCTGGG<br>CTTTGAATATTGCTGTTACGGTATCCGTGTTCCGCTGCCGGTCAGCAAAC<br>CGGCGGTTGCAATTTTCGATACCTATCCGGACGGCTGGATGGCACATTAT<br>CAGGCTCAAAACTACATTGAAATCGATTCTACCGTGCGCGACGGTGCGC<br>TGAACACGAATATGATCGTGTGGCCGGATGTTGACCGTATTGATCCGTGC<br>CCGCTGTGGCAGGATGCACGTGACTTTGGCCTGTCAGTCGGTGTGGCAC<br>AAAGCTCTTGGGCAGCACGTGGTGCATTCGGTCTGCTGTCGATCGCACG | *B. cenocepacia* |

| | | |
|---|---|---|
| | TCATGCTGATCGCCTGACCCCGGCAGAAATTAATATGCTGACCCTGCAAA<br>CGAACTGGCTGGCTAATCTGAGTCACTCCCTGATGAGCCGCTTTATGGTT<br>CCGAAACTGTCTCCGGCAGCTGGCGTCACCCTG | |
| VanR | atGGACATGCCCAGAATTAAACCTGGCCAGAGAGTAATGATGGCACTAAG<br>GAAAATGATTGCGTCCGGTGAAATAAAGTCCGGTGAAAGAATTGCTGAG<br>ATCCCTACCGCTGCTGCGCTAGGCGTTTCCAGGATGCCCGTTCGTATAGC<br>CTTACGTTCTTTGGAGCAAGAAGGCTTGGTCGTAAGACTGGGCGCAAG<br>AGGGTATGCGGCCAGGGGAGTCAGCTCCGACCAGATAAGAGACGCGAT<br>AGAGGTGCGTGGTGTTCTGGAAGGGTTTGCCGCGAGGCGTCTTGCAGA<br>AAGAGGTATGACGGCGGAGACACACGCAAGATTCGTGGTATTGATAGCT<br>GAGGGGGAAGCTCTTTTTGCGGCGGGTCGTCTAAACGGCGAGGATCTT<br>GATCGTTATGCGGCTTACAACCAGGCCTTCCACGACACTTTAGTCTCTGC<br>TGCCGGGAACGGAGCAGTCGAATCTGCGCTGGCGAGAAACGGATTCGA<br>GCCGTTTGCGGCGGCCGGTGCCTTGGCCCTAGACCTAATGGACCTTAGT<br>GCAGAGTACGAGCATCTATTGGCAGCTCACAGACAACACCAGGCTGTCT<br>TGGATGCAGTGTCCTGTGGCGACGCAGAGGGCGCGGAAAGGATTATGA<br>GAGATCACGCGCTTGCGGCGATAAGAAATGCGAAGGTGTTCGAAGCGG<br>CAGCCTCAGCGGGAGCGCCATTAGGTGCTGCATGGAGTATTAGAGCCGA<br>C | [4] |
| $PauR_{17-160}$ | ACGATCCCGGAATTAGAAAAGAAAGTACTTAATGCCATAAACTACCTTG<br>ACTTCGAATTCTATTCCATCTACCAAGAAGAGAAGTATCCCTTCACTACT<br>AAGAAATATGTGCTAATAACGAACTTACATACAGAGACTATGGGGAATTA<br>CGACAAGAACGATAACCATGATTCTGACCAGTTAATGAATGGGGACTTT<br>ATATTCTCTATACCGTCAATATATGATGAAAAAACAAGCAAGCACACGGA<br>CTTCATGATCGAAAAGCCGAACGAAAACGAGATAAAGAACATTATAAGT<br>ATAAAGATCAAGTCCTACCACGGCGAAAGATCCATATTCTCTGTGGCAA<br>GGAAAGAGGGTCCGATCTCCCACAATGAGATTCTTAACAAGGCCGGTAT<br>GCTTGGAATTCTTGCTAATGAGTGTATAAAGACCTTT | *P. asymbiotica* |
| $PluR_{17-162}$ | ACGATGCCTGAACTAGAAAAGCGTCTGCTAAACGCCATTAACTATTTGG<br>GCTTTGAGTTCTATAGTATTTACCAAGAGGAGAAATACCCGTTCACAACG<br>AAGAAATACGTGCTGATAACGAATCTGAACATTGAAACAGCCAGTAACT<br>ATAACCAGGATAATAACCATGACAGTGACCAGCTTATGAACGGGGACTA<br>TACCTTTTCTATTCCAAGTGTCTGCAATGACAAGATCTCCAAGAACACTG<br>ACTTCACTATCGAGAAGAGCAATTCAAATAAAAACGAAGTGAAGAACT<br>CACTTAGTATTAAAATCAAAGGTTATCACGGGGAAAGAAGTATTTTCAGC<br>GTAGCTAGAAAGGAAGGAACTATATCTCACAATGAGATACTAAACAAGG<br>CTGGAGTATTAGGCATACTTGCCAATGAATGTATCAAGACGTTC | *P. laumondii* |

#### Supplementary Table 12: Sequence of ADs used in this study.

| ADs | Sequence | Source |
|---|---|---|
| A1+ | TATCAAAAAACTAGAGGTGCTACTGCTAGATCTCCAAGAAGATCTTCTTCTAATCCAACTT<br>GGTGGTTTGATTATTTTTATTGTATGAGA | |
| B1+ | CATAGAAGAACTCATAATTATGAAAAATCTTGTACTTCTTTGGAACCATGGTGGCAATTTG<br>GTATTAATTTGGTTAATTGGCCACCATTG | |
| B2+ | AGAAGAAAAAGAGCTAGAGGTTCTAGACCAACTCATTGGGATTTGTTGTGGGATGAATAT<br>TCTGAAATTTCTGCTTTTTGTAATTTGATT | |
| C2+ | GTTAAAAATACTTGTGTTAGAGAAGGTACTACTGGTAAAGGTACTGATTATTTGGTTATGA<br>ATTGGTGGGATGGTTTTTATGATGTTTTG | |
| D1+ | TCTACTAAAGGTGTTGCTTTGGATGATGTTGAAGGTGATGATGATTTTTATATGTTTGATATT<br>GATATTATTGCTTCTAGATCTACTGGT | |
| D2+ | AATAATGGTACTTCTATTGAAGGTTGTTCTGAAAATATTGATGTTGTTTTGGAAGCTGATTT<br>GGATTTGCCATTGGATTATTTGATTGTT | |
| E1+ | AGATCTGCTAGAGATGATGGTGTTGGTGAAGAAGAAGAAGATGATTATGGTGTTTATTATG<br>ATTTTGATGAATGGTTTGATGAAAATTCT | |
| haB112 | GATATCATGCAGGATCTGCCGGGCAACGATAACAGCACCGCGGGCGAATTTCCAGGTATT<br>ACTTTGAGAATCCAAGAAACTGATATGTTGTATAAAGGTGATACTTTGTATTTGGATTGGTT<br>GGAAGATGGTATTGCTGAATTGGTTTTCGATGCTCCAGGTTCTGTTAATAAATTGGATACTG<br>CTGTTGCTTCTTTGGGTGAAGCTATTGGTGTTTTGGAACAACAATCTGATTTGATTTGGGA<br>AACTTTGACTGTTAAAGATGCTAAAGTTAATTTCGATTCTGGTTTGGAAAAATTCGAAGAA<br>GCTATTCCATCTGCTGATGATTTCGATCCAGTTGCTGAAAGAAGATCCTCTGGTGAATTTA<br>GAGCTGAAAGACATTCTGGTGGTACTGATTTGTGTTTCAAGCTT | |
| GAL4 | AATTTTAATCAAAGTGGGAATATTGCTGATAGCTCATTGTCCTTCACTTTCACTAACAGTAG<br>CAACGGTCCGAACCTCATAACAACTCAAACAAATTCTCAAGCGCTTTCACAACCAATTGC<br>CTCCTCTAACGTTCATGATAACTTCATGAATAATGAAATCACGGCTAGTAAAATTGATGATG<br>GTAATAATTCAAAACCACTGTCACCTGGTTGGACGGACCAAACTGCGTATAACGCGTTTG<br>GAATCACTACAGGGATGTTTAATACCACTACAATGGATGATGTATATAACTATCTATTCGAT<br>GATGAAGATACCCCACCAAACCCAAAAAAAGAG | |
| rTA | AGGGATAGCCGGGAAGGTATGTTCCTGCCAAAGCCGGAAGCGGGCAGTGCCATATCTGA<br>CGTGTTCGAGGGGCGAGAGGTGTGTCAGCCAAAGAGGATCAGGCCCTTCCATCCACCCG<br>GATCCCCGTGGGCCAACCGGCCCCTGCCTGCCTCTTTGGCTCCCACCCCCACAGGACCTG<br>TCCATGAACCGGTCGGATCCCTAACGCCAGCCCCGGTGCCCCAGCCACTTGACCCGGCCC<br>CCGCAGTAACCCCCGAGGCAAGTCATCTGTTGGAGGACCCTGATGAAGAAACCAGTCAG<br>GCCGTGAAGGCCCTAAGGGAGATGGCTGACACTGTTATTCCCCAGAAGGAGGAAGCAGC<br>CATATGTGGACAGATGGACCTGAGCCACCCGCCCCCTCGTGGCCATTTGGACGAACTGAC<br>CACAACACTAGAGTCCATGACAGAGGATTTGAATCTGGACTCCCCCCTGACCCCCGAACT<br>TAATGAAATCTTGGATACATTTCTAAATGATGAATGTCTGCTGCATGCCATGCATATTTCAA<br>CTGGGCTGTCTATTTTTGACACCAGCTTATTT | |
| GCN4 | ATGTTTGAGTATGAAAACCTAGAgGACAACTCTAAAGAATGGACATCCTTGTTTGACAATG<br>ACATTCCAGTTACCACTGACGATGTTTCATTGGCTGATAAGGCAATTGAATCC | |
| Msn2 | ATGACGGTCGACCATGATTTCAATAGCGAAGATATTTTATTCCCCATAGAAAGCATGAGTA<br>GTATACAATACGTGGAGAATAATAACCCAAATAATATTAACAACGATGTTATCCCGTATTCT<br>CTAGATATCAAAAACACTGTCTTAGATAGTGCGGATCTCAATGACATTCAAAATCAAGAAA<br>CTTCACTGAATTTGGGGCTTCCTCCACTATCTTTCGACTCTCCACTGCCCGTAACGGAAAC<br>GATACCATCCACTACCGATAACAGCTTGCATTTGAAAGCTGATAGCAACAAAAATCGCGAT<br>GCAAGAACTATTGAAAATGATAGTGAAATTAAGAGTACTAATAATGCTAGTGGCTCTGGGG<br>CAAATCAATACACAACTCTTACTTCACCTTATCCTATGAACGACATTTTGTACAACATGAA<br>CAATCCGTTACAATCACCGTCACCTTCATCGGTACCTCAAAATCCGACTATAAATCCTCCC<br>ATAAATACAGCAAGTAACGAAACTAATTTATCGCCTCAAACTTCAAATGGTAATGAAACTC<br>TTATATCTCCTCGAGCCCAACAACATACGTCCATTAAAGATAATCGTCTGTCCTTACCTAAT<br>GGTGCTAATTCGAATCTTTTCATTGACACTAACCCAAACAATTTGAACGAAAAACTAAGA<br>AATCAATTGAACTCAGATACAAATTCATATTCTAACTCCATTTCTAATTCAAACTCCAATTC<br>TACGGGTAATTTAAATTCCAGTTATTTTAATTCACTGAACATAGACTCCATGCTAGATGATT<br>ACGTTTCTAGTGATCTCTTATTGAATGATGATGATGATGACACTAATTTATCACGCCGAAGA<br>TTTAGCGACGTTATAACAAACCAA | |
| VP16 | GATATCATGCAGGATCTGCCGGGCAACGATAACAGCACCGCGGGCGCGCGTACGAAAAA<br>CAATTACGGGTCTACCATCGAGGGCCTGCTCGATCTCCCGGACGACGACGCCCCCGAAGA<br>GGCGGGGCTGGCGGCTCCGCGCCTGTCCTTTCTCCCCGCGGGACACACGCGCAGACTGT<br>CGACGGCCCCCCCGACCGATGTCAGCCTGGGGGACGAatTgCACTTAGACGGCGAGGACG<br>TGGCGATGGCGCATGCCGACGCGCTAGACGATTTCGATCTGGACATGTTGGGGGACGGGG<br>ATTCCCCGGGTCCGGGATTTACCCCCCACGACTCCGCCCCCTACGGCGCTCTGGATATGGC<br>CGACTTCGAGTTTGAGCAGATGTTTACCGATGCCCTTGGAATTGACGAGTACGGTGGGAA<br>GCTT | |
| VP16_opt1 | ATGATCATGCAGGATCTACCCGGAAACGACAACTCTACAGCCGGTGCCAGAACGAAAAAT | Sangon |

| | |
|---|---|
| | AATTATGGTTCTACTATTGAAGGACTGCTAGATTTACCCGATGATGACGCCCCTGAGGAGG |
| | CAGGTCTAGCAGCTCCTAGATTGTCCTTCCTGCCCGCAGGTCATACTCGTAGGCTATCAAC |
| | TGCACCCCCAACAGATGTATCTTTAGGCGACGAGTTGCATCTTGACGGCGAGGACGTAGC |
| | TATGGCGCATGCAGACGCCCTAGATGATTTCGATTTGGATATGCTGGGAGATGGGGACAGC |
| | CCGGGGCCGGGCTTTACACCACATGATTCCGCTCCCTACGGTGCCTTAGATATGGCCGACT |
| | TTGAGTTTGAGCAGATGTTCACAGACGCATTAGGGATAGACGAATACGGTGGCAAACTA |
| VP16_opt2 | ATGATTATGCAAGACCTACCTGGGAACGATAACTCTACTGCGGGGGCGAGGACCAAAAAT |
| | AACTATGGGAGCACAATAGAGGGGCTTCTTGACCTGCCCGACGACGATGCCCCCGAAGA |
| | AGCAGGGCTGGCTGCACCGCGTTTAAGTTTCCTACCTGCGGGACATACCAGGAGGCTTTC |
| | AACGGCACCTCCGACAGATGTCTCATTGGGGGATGAGTTACATCTGGACGGAGAGGACG |
| | TAGCAATGGCTCACGCTGACGCGTTAGACGATTTCGATCTGGACATGCTGGGAGATGGCG |
| | ATTCCCCGGGTCCGGGCTTCACTCCCCATGATTCTGCCCCGTATGGGGCATTAGATATGGC |
| | GGATTTTGAATTCGAACAGATGTTTACTGACGCATTAGGTATAGATGAGTATGGTGGTAAA |
| | CTA |
| VP16_opt3 | ATGATTATGCAAGATCTTCCGGGAAACGACAATTCTACAGCGGGCGCCAGGACTAAAAAT |
| | AATTACGGTAGTACGATTGAGGGTCTGCTGGATCTTCCCGATGATGATGCGCCCGAAGAG |
| | GCTGGTCTAGCCGCCCCCCGTTTGTCATTCCTACCGGCAGGCCACACTAGAAGGCTATCTA |
| | CCGCCCCTCCGACGGATGTCTCTCTTGGTGACGAGCTTCACTTGGATGGTGAAGATGTTG |
| | CAATGGCGCATGCAGATGCACTGGACGATTTTGACTTGGATATGCTAGGGGACGGGGATT |
| | CACCCGGCCCAGGATTCACACCGCACGATAGCGCCCCATATGGGGCTCTGGATATGGCCG |
| | ACTTTGAATTTGAGCAGATGTTTACAGACGCACTAGGCATCGACGAGTATGGCGGCAAGC |
| | TG |
| VP16_opt4 | ATGATAATGCAGGACTTACCAGGGAATGACAACTCTACCGCCGGAGCGAGGACTAAAAAT |
| | AATTATGGGTCTACCATCGAGGGGCTTTTAGACCTGCCAGACGATGACGCACCGGAGGAA |
| | GCGGGCCTAGCTGCGCCGCGTTTAAGCTTTCTACCCGCTGGGCACACTAGGCGTCTGAGC |
| | ACAGCTCCACCAACCGATGTGAGCCTAGGAGACGAGTTGCATTTAGACGGCGAGGATGT |
| | GGCTATGGCTCATGCGGATGCCTTAGATGACTTTGACCTGGACATGCTAGGAGATGGAGA |
| | CAGCCCGGGGCCTGGCTTCACCCCACACGACTCCGCGCCTTATGGGGCGTTGGACATGGC |
| | AGATTTCGAGTTCGAGCAAATGTTTACAGACGCCTTAGGAATCGACGAATACGGCGGGAA |
| | GCTT |
| VP16_opt5 | ATGATAATGCAAGACCTACCTGGGAACGACAATAGCACCGCTGGAGCCCGTACAAAGAA |
| | CAATTATGGGTCCACTATAGAAGGTTTATTGGATCTTCCTGACGATGACGCACCGGAGGAG |
| | GCGGGATTGGCTGCGCCACGTTTATCCTTCTTGCCCGCTGGCCACACAAGGAGATTATCA |
| | ACGGCACCTCCCACAGACGTAAGCTTGGGAGATGAGCTACATCTGGACGGTGAGGACGT |
| | TGCGATGGCACACGCTGACGCCCTTGACGACTTTGATCTAGATATGCTTGGTGATGGCGAT |
| | TCCCCGGGTCCAGGATTTACTCCGCACGACTCAGCCCCTTATGGCGCCCTGGACATGGCG |
| | GATTTCGAGTTTGAACAGATGTTTACCGATGCGCTTGGAATCGACGAGTATGGAGGAAAG |
| | CTA |
| VP16_opt6 | ATGATTATGCAGGATCTTCCCGGGAATGATAACTCTACGGCTGGTGCTAGGACAAAGAAC |
| | AATTACGGCTCCACCATCGAGGGATTACTTGACTTACCAGATGACGACGCTCCTGAAGAG |
| | GCCGGATTGGCGGCGCCGAGGCTGTCCTTTCTTCCCGCCGGACACACTAGGAGATTATCC |
| | ACTGCTCCTCCTACCGACGTTAGTTTGGGAGATGAATTGCACCTGGACGGGGAGGACGTA |
| | GCTATGGCGCATGCGGATGCCCTTGATGACTTTGATTTGGACATGCTAGGTGACGGTGATA |
| | GTCCGGGCCCGGGTTTTACACCCCATGATTCAGCCCCTTACGGAGCATTAGATATGGCTGA |
| | CTTTGAGTTTGAACAAATGTTTACTGATGCCTTAGGCATAGATGAATATGGGGGTAAGCTG |
| VP16_opt7 | ATGATAATGCAAGACTTGCCAGGCAATGATAACAGCACCGCCGGTGCTAGAACAAAGAA |
| | CAATTACGGATCTACAATCGAAGGGCTTTTAGATCTGCCTGACGATGACGCGCCTGAGGA |
| | GGCAGGGCTGGCAGCGCCGAGACTATCCTTCTTACCGGCAGGACACACCAGACGTCTTA |
| | GTACGGCACCACCAACGGATGTCTCTCTAGGAGATGAGCTACACTTAGACGGCGAAGATG |
| | TAGCCATGGCCCACGCTGACGCCTTGGACGATTTCGACCTTGATATGTTGGGGGACGGAG |
| | ACTCCCCAGGTCCAGGCTTCACACCCCATGATTCTGCTCCGTATGGGGCTCTTGATATGGC |
| | AGACTTTGAGTTTGAGCAGATGTTTACCGATGCCCTTGGGATAGATGAGTACGGAGGTAA |
| | ATTA |
| VP16_opt8 | ATGATAATGCAAGACTTACCAGGCAACGATAACTCTACTGCCGGAGCAAGGACTAAAAAC |
| | AATTATGGTTCTACAATTGAAGGACTTCTTGACCTACCAGATGACGACGCACCAGAAGAA |
| | GCAGGTCTTGCCGCCCCGCGTTTATCTTTTCTACCCGCGGGTCACACGAGGAGACTTAGT |
| | ACTGCGCCACCAACTGATGTGTCCCTTGGCGACGAACTTCATCTAGATGGCGAAGATGTA |
| | GCTATGGCACATGCTGATGCCCTGGACGATTTTGATTTAGACATGCTAGGTGATGGAGATA |
| | GCCCTGGACCTGGCTTTACTCCCCACGATTCTGCGCCGTATGGCGCTCTAGACATGGCGG |
| | ACTTTGAGTTCGAACAGATGTTCACGGACGCATTAGGTATTGACGAGTATGGGGGGAAGT |
| | TA |
| VP16_opt9 | ATGATTATGCAAGACTTACCGGGTAATGACAACTCTACAGCGGGTGCCCGTACGAAAAAT |
| | AACTACGGTTCTACCATCGAAGGACTTCTAGACTTACCAGATGATGACGCCCCCGAAGAA |
| | GCTGGGCTAGCAGCTCCCAGATTGTCATTTCTGCCGGCAGGTCACACCAGAAGGCTGAGC |
| | ACTGCCCCCACCGACTGATGTGTCTCTGGGAGATGAACTTCACTTGGACGGTGAGGATGTG |
| | GCGATGGCTCATGCCGACGCGCTGGATGATTTCGACCTGGATATGCTTGGCGATGGAGAC |
| | AGTCCGGGTCCGGGATTCACTCCCCATGACAGTGCCCCCTACGGTGCCTTAGACATGGCA |

| | |
|---|---|
| | GACTTTGAGTTTGAGCAAATGTTTACAGACGCTCTTGGAATAGATGAGTATGGTGGTAAA<br>CTA |

#### Supplementary Table 13: Sequence of metabolic genes involved in the VB5 biosynthesis.

| Metabolism | Sequence | Source |
|---|---|---|
| *Sc*ilv2 | **ATG**ATCAGACAGTCTACCTTGAAGAACTTCGCTATTAAGAGATGCTTCCAACATATCGC<br>TTACAGAAATACTCCAGCCATGAGATCTGTTGCTTTGGCTCAAAGATTTTACTCCTCATC<br>TTCTAGGTACTACTCTGCTTCTCCATTGCCAGCTTCTAAAAGACCAGAACCAGCTCCATC<br>TTTTAACGTTGATCCATTGGAACAACCAGCTGAACCATCTAAATTGGCTAAAAAGTTGA<br>GAGCCGAACCAGATATGGATACTTCTTTTGTTGGTTTGACTGGTGGTCAGATCTTCAAC<br>GAAATGATGTCCAGACAAAACGTTGATACCGTTTTTGGTTATCCAGGTGGTGCTATTTTG<br>CCAGTTTATGATGCTATTCACAACTCCGACAAGTTCAACTTCGTTTTGCCAAAACATGA<br>ACAAGGTGCTGGTCACATGGCTGAAGGTTATGCTAGAGCTTCTGGTAAACCAGGTGTT<br>GTTTTGGTTACTTCTGGTCCAGGTGCTACTAATGTTGTTACTCCAATGGCTGATGCTTTC<br>GCTGATGGTATTCCAATGGTTGTTTTTACTGGTCAAGTTCCAACATCCGCTATTGGTACA<br>GATGCTTTTCAAGAAGCTGACGTTGTTGGTATTTCCAGATCTTGTACTAAGTGGAACGT<br>TATGGTTAAGTCCGTTGAAGAATTGCCATTGAGAATCAATGAAGCCTTCGAAATTGCTA<br>CATCAGGTAGACCAGGTCCAGTTTTGGTTGATTTGCCTAAAGATGTTACTGCCGCCATTT<br>TGAGAAATCCAATTCCAACTAAGACTACGTTGCCATCCAATGCCTTGAATCAATTGACAT<br>CAAGAGCCCAAGATGAGTTCGTTATGCAATCTATTAACAAGGCTGCCGACTTGATTAAC<br>TTGGCAAAAAAACCAGTCTTGTACGTTGGTGCTGGTATTTTGAATCATGCTGATGGTCC<br>AAGGTTGCTGAAAGAATTGTCTGATAGAGCACAAATTCCAGTTACTACCACATTGCAAG<br>GTTTGGGTTCTTTCGATCAAGAAGATCCAAAGTCCTTGGACATGTTGGGTATGCATGGT<br>TGTGCTACTGCTAATTTGGCTGTTCAAAACGCCGATTTGATTATTGCTGTTGGTGCTAGA<br>TTCGATGATAGAGTTACTGGTAACATTTCCAAGTTTGCTCCAGAAGCTAGAAGGGCTGC<br>TGCTGAAGGTAGAGGTGGTATTATTCATTTTGAAGTCTCCCCAAAGAATATCAACAAGG<br>TCGTTCAAACCCAAATTGCCGTTGAAGGTGATGCAACTACTAATTTGGGTAAGATGATG<br>AGCAAGATCTTCCCAGTCAAAGAAAGATCTGAATGGTTCGCTCAAATCAACAAGTGGA<br>AGAAAGAATACCCATACGCCTACATGGAAGAAACTCCAGGTTCTAAAATCAAGCCACA<br>AACCGTTATCAAGAAGTTGTCTAAGGTTGCTAACGATACCGGTAGACATGTTATAGTTAC<br>TACTGGTGTTGGTCAACATCAAATGTGGGCTGCTCAACATTGGACTTGGAGAAATCCTC<br>ATACTTTCATTACATCTGGTGGTTTGGGTACTATGGGTTATGGTTTGCCAGCTGCTATTGG<br>TGCTCAAGTTGCTAAACCAGAATCCTTGGTTATTGATATTGATGGTGATGCCTCTTTCAA<br>CATGACCTTGACTGAATTATCTTCCGCTGTTCAAGCTGGTACTCCAGTTAAGATTTTGAT<br>CTTGAACAACGAGGAACAAGGTATGGTTACACAATGGCAGTCTTTGTTCTACGAACATA<br>GATACTCTCATACCCACCAATTGAACCCAGATTTCATTAAGTTGGCTGAAGCCATGGGTT<br>TAAAAGGTTTGAGAGTTAAGAAACAAGAAGAGTTGGACGCCAAGCTGAAAGAGTTTG<br>TTTCTACTAAGGGTCCTGTTTTGTTGGAAGTTGAAGTTGATAAGAAGGTTCCAGTCTTG<br>CCTATGGTTGCTGGTGGTTCAGGTTTGGATGAGTTTATCAATTTTGACCCAGAAGTCGA<br>AAGACAGCAAACTGAATTGAGACATAAGAGAACCGGTGGTAAACAC**TAA** | [5] |
| *Sc*ilv6 | **ATG**CTGAGATCCTTGTTGCAATCTGGTCATAGAAGAGTTGTTGCTTCATCTTGTGCTACT<br>ATGGTCAGATGTTCTTCCTCTTCTACATCTGCTTTGGCTTACAAGCAAATGCATAGACAT<br>GCTACTAGACCACCATTGCCAACTTTGGATACTCCATCTTGGAATGCTAATTCCGCTGTT<br>TCCTCTATTATCTACGAAACTCCAGCTCCATCAAGACAACCTAGAAAACAACATGTCTT<br>GAACTGCTTGGTCCAAAATGAACCAGGTGTTTTGTCAAGAGTTTCTGGTACTTTGGCTG<br>CTAGAGGTTTCAACATTGATTCTTTGGTTGTCTGTAACACCGAAGTCAAAGATTTGTCC<br>AGAATGACCATCGTATTGCAAGGTCAAGATGGTGTTGTTGAACAAGCTAGAAGGCAAA<br>TCGAAGATTTGGTTCCAGTTTACGCTGTTTTGGATTACACCAACTCCGAAATTATCAAGC<br>GTGAATTGGTTATGGCCAGGATTTCTTTGTTGGGTACTGAATACTTTGAGGACTTGTTGT<br>TGCATCACCATACTTCTACTAATGCTGGTGCTGCTGATTCTCAAGAATTGGTTGCTGAAA<br>TCAGAGAGAAGCAATTTCATCCAGCTAATTTGCCAGCTTCCGAAGTCTTGAGATTGAAA<br>CATGAACACTTGAACGACATCACCAACTTGACTAACAATTTCGGTGGTAGAGTTGTCGA<br>TATCTCTGAAACTTCTTGCATCGTTGAATTGTCTGCTAAGCCAACTAGAATTTCCGCTTT<br>CTTGAAATTGGTTGAACCATTCGGTGTTTTGGAATGTGCTAGATCTGGTATGATGGCTTT<br>GCCAAGAACTCCATTGAAAACTTCTACTGAAGAGGCTGCTGATGAAGATGAGAAGATT<br>TCTGAAATCGTGGACATCTCTCAATTGCCACCAGGT**TAA** | |
| *Tc*panD | **ATG**CCTGCTACTGGTGAAGATCAAGATTTGGTTCAAGATTTGATTGAAGAACCTGCTAC<br>TTTTTCTGATGCTGTTTTATCTTCTGATGAAGAATTGTTCCATCAAAAATGTCCAAAACC<br>TGCTCCAATTTATTCTCCTGTTTCTAAACCTGTTTCTTTTGAATCTTTGCCAAATAGAAGA<br>TTGCATGAAGAATTTTTGAGATCTTCTGTTGATGTTTTGTTGCAAGAAGCTGTTTTTGAA<br>GGTACTAATAGAAAAAATAGAGTTTTGCAATGGAGAGAACCTGAAGAATTGAGAAGAT<br>TGATGGATTTTGGTGTTAGATCTGCTCCATCTACACATGAAGAGTTGTTAGAAGTTTTGA<br>AAAAGGTTGTTACTTATTCTGTTAAAACTGGTCATCCATATTTTGTTAATCAATTGTTTTC<br>TGCTGTTGATCCATATGGTTTGGTTGCTCAATGGGCTACTGATGCTTTGAATCCATCTGTT<br>TATACTTATGAAGTTTCTCCTGTTTTTGTTTTGATGGAAGAAGTTGTTTTGAGAGAAATG<br>AGAGCTATTGTTGGTTTTGAAGGTGGTAAAGGTGATGGTATTTTTTGTCCTGGTGGTTCT<br>ATTGCTAATGGTTATGCTATTTCTTGTGCTAGATATAGATTTATGCCTGATATTAAAAAGA<br>AAGGTTTGCATTCTTTGCCAAGATTAGTCTTGTTTACTTCAGAAGATGCTCATTATTCTAT | |

| | |
|---|---|
| | TAAGAAATTGGCTTCTTTTCAAGGTATTGGTACTGATAATGTTTATTTGATTAGAACTGAT<br>GCTAGAGGTAGAATGGATGTTTCTCATTTAGTTGAAGAAATTGAGAGGTCTTTGAGAGA<br>AGGTGCTGCTCCATTTATGGTTTCTGCTACTGCTGGTACTACTGTTATTGGTGCTTTTGAT<br>CCAATTGAAAAAATTGCTGATGTTTGTCAAAAATATAAATTGTGGTTGCATGTTGATGCT<br>GCTTGGGGTGGAGGTGCTTTAGTTTCTGCTAAACATAGACATTTGTTGAAAGGTATTGA<br>AAGAGCTGATTCTGTTACTTGGAATCCACATAAATTGTTGACTGCTCCACAACAATGTTC<br>TACTTTGTTATTGAGACATGAAGGTGTTTTGGCTGAAGCTCACTCTACAAATGCTGCTTA<br>CTTATTTCAGAAAGATAAATTCTATGACACAAAATATGACACTGGTGATAAACATATTCA<br>ATGTGGTAGAAGAGCTGATGTTTTGAAATTTTGGTTTATGTGGAAAGCTAAAGGTACTT<br>CTGGTTTGGAAAAACATGTCGATAAAGTTTTCGAAAATGCTAGATTCTTCACTGATTGTA<br>TTAAAAACAGAGAGGGTTTCGAAATGGTTATTGCTGAACCTGAATATACTAATATTTGTT<br>TTTGGTATGTTCCAAAGTCATTGAGGGGTAGAAAAGATGAAGCAGATTATAAGGATAAA<br>TTGCATAAGGTTGCTCCAAGAATTAAAGAAAGAATGATGAAAGAAGGTTCTATGATGGT<br>TACTTATCAAGCTCAAAAAGGTCACCCAAATTTTTTTAGAATTGTTTTCCAAAATTCTGG<br>TTTGGATAAAGCTGATATGGTTCATTTGGTCGAAGAAATTGAAAGATTGGGTTCTGATTT<br>G**TAA** |
| *Cg*panC | **ATG**CAAGTTGCTACTACAAAGCAAGCTTTGATCGATGCATTGTTGCATCATAAGTCTGTT<br>GGTTTGGTTCCAACTATGGGTGCTTTACATTCTGGTCATGCATCATTGGTTAAAGCTGCA<br>AGAGCTGAAAACGATACAGTTGTTGCATCAATCTTCGTTAACCCATTGCAATTCGAAGC<br>TTTGGGTGACTGTGATGATTACAGAAACTACCCAAGACAATTGGATGCTGATTTGGCAT<br>TGTTAGAAGAAGCTGGTGTTGATATTGTTTTTGCACCAGATGTTGAAGAAATGTATCCA<br>GGTGGTTTGCCATTAGTTTGGGCTAGAACTGGTTCTATTGGTACAAAATTGGAAGGTGC<br>ATCAAGACCAGGTCATTTTGATGGTGTTGCTACTGTTGTTGCAAAGTTGTTTAATTTGGT<br>TAGACCAGATAGAGCTTACTTTGGTCAAAAAGATGCTCAACAAGTTGCAGTTATTAGAA<br>GATTGGTTGCAGATTTGGATATTCCAGTTGAAATTAGACCAGTTCCAATTATTAGAGGTG<br>CTGATGGTTTAGCAGAATCTTCAAGAAACCAAAGATTGTCTGCTGATCAAAGAGCTCA<br>AGCATTGGTTTTACCACAAGTTTTGTCAGGTTTACAAAGAAGAAAAGCTGCAGGTGAA<br>GCTTTAGATATTCAAGGTGCAAGAGATACATTGGCTTCTGCAGATGGTGTTAGATTGGAT<br>CATTTGGAAATTGTTGATCCAGCTACTTTGGAACCATTAGAAATTGATGGTTTGTTAACA<br>CAACCAGCTTTAGTTGTTGGTGCAATTTTTGTTGGTCCAGTTAGATTGATTGATAATATTG<br>AATTA**TAA** |
| *Sc*ecm31 | ATGAACATCATGAAGAGACAGTTGTGCACCTCTTCTAAGAGATTTTTCTCTACCGCTAA<br>AAACGTCGTCAAGTACAACACCATTCAAGACATCAGAAACAAGTACTTCACTGGCACT<br>CCATTGTCTATGTGTACTGCTTACGATTTCATTACTGCTACCTGGGTTAACAAGGCTAAC<br>TGTGATTTGTTGTTGGTTGGTGATTCTTTGGCTATGACTTCTTTGGGTTACGATTCTACCA<br>TTACCTTGTCTTTGAACGAGTTCAAATACCATGTTGCCTCTGTATGTAGAGCTGAAGGTT<br>CTTCTATGGTTGTTGTTGATATGCCATTCGGTACTTTCGAATCCGGTATTTCTGATGGTTT<br>GAAGAACGCCATTGATATCATGAAGTTGGATTCCAAGGTTACCTCCGTTAAGGTTGAAG<br>TTGGTTCTTACACTAAGGATAAGTACGCCATGAAGTTCATCGAAGAATTGTGCTCAAGA<br>GGTATTCCAGTTATGGCTCATATTGGTTTGACTCCACAAAAGGTCCATTCTTTAGGTGGT<br>TACAAAGTCCAAGGTTCCAAGTCTTTGTTGCAAATGCAAGAGTTGTACGAAACCGCCA<br>TGCAGTTGCAAAAAATTGGTTGTTGGTCCATCTTGATCGAATGCGTTCCACATAAGATG<br>GCTCAATTCATTACCTCCAAATTGTCCGTTCCAACCATTGGTATTGGTGCTGGTAACGGT<br>ACTTCTGGTCAAGTTTTGGTTATTTCCGATTTGTTGGGTATGCAAGGTGATTCTGTTCCA<br>AAGTTTGTTAAGCAAGCTGTTAACATGACCGATATTGCTACCCAAGGTCTGAAAGAGTA<br>TATCGCTTCTGTTGAAGATAGGACTTTTCCAGAAAGAGGTACTCATACCTTCAAGGTCA<br>AAGAAGATCTTTGGAACGAGTTTCTGTCCTCCATTAACGAAAAGTAA |
| *Mt*panE | ATGGCTACTGGTATTGCTTTGGTTGGTCCAGGTGCTGTTGGTACTACTGTTGCTGCTTTG<br>TTGCATAAGGCTGGTTATTCTCCATTATTGTGTGGTCATACTCCAAGAGCTGGTATCGAA<br>TTGAGAAGAGATGGTGCTGATCCAATAGTTGTTCCAGGTCCAGTTCATACTTCTCCAAG<br>AGAAGTTGCTGGTCCAGTTGATGTTTTGATTTTGGCTGTTAAGGCTACCCAAAATGATG<br>CTGCTAGACCATGGTTGACTAGATTGTGTGACGAAAGAACTGTTGTTGCTGTCTTGCAA<br>AATGGTGTTGAACAAGTCGAACAAGTTCAACCACATTGTCCATCTTCAGCTGTTGTACC<br>AGCTATAGTTTGGTGTTCTGCTGAAACTCAACCACAAGGTTGGGTTAGATTGAGAGGTG<br>AAGCTGCATTGGTTGTTCCAACTGGTCCAGCTGCAGAACAATTTGCTGGTTTGTTAAGA<br>GGTGCTGGTGCTACAGTTGATTGTGATCCAGATTTTACTACTGCTGCTTGGAGAAAGTT<br>GTTGGTTAATGCTTTGGCTGGTTTCATGGTTTTGTCTGGTAGAAGATCTGCCATGTTTAG<br>AAGGGATGATGTTGCTGCATTGAGTAGAAGATATGTTGCTGAATGTTTGGCAGTTGCTA<br>GAGCTGAAGGTGCTAGATTGGATGATGATGTAGTTGATGAAGTTGTCAGATTGGTTAGA<br>TCTGCTCCACAAGATATGGGTACTTCTATGTTGGCTGATAGAGCTGCTCATAGACCATTG<br>GAATGGGATTTGAGAAACGGTGTTATCGTTAGAAAAGCTAGAGCACATGGTTTGGCTAC<br>TCCAATTTCTGATGTTTTAGTTCCTTTGTTGGCTGCTGCTTCTGATGGTCCTGGTTAA |
| *Pa*ilvC | ATGAGGGTTTTCTACGATAAGGATTGCGACTTGTCCATTATCCAAGGTAAGAAGGTTGC<br>CATTATTGGTTATGGTTCTCAAGGTCATGCTCATGCTTGTAACTTGAAAGATTCTGGTGT<br>TGATGTTACCGTCGGTTTGAGATCTGGTTCTGCTACTGTTGCTAAAGCTGAAGCTCATG<br>GTTTGAAAGTTGCTGATGTTAAGACTGCTGTTGCTGCTGCAGATGTTGTTATGATTTTGA<br>CCCCAGATGAATTTCAAGGCAGGTTGTACAAAGAAGAAATCGAGCCAAATTTGAAGAA |

| | |
|---|---|
| | GGGTGCTACTTTGGCTTTTGCCCATGGTTTTTCCATCCACTACAATCAAGTTGTTCCAAG<br>AGCTGATTTGGACGTTATTATGATTGCTCCAAAAGCTCCAGGTCATACCGTTAGATCTGA<br>ATTTGTCAAAGGTGGTGGTATCCCAGATTTGATTGCTATCTATCAAGATGCTTCTGGTAA<br>CGCTAAGAACGTTGCTTTGTCTTATGCTTGTGGTGTTGGTGGTGGAAGAACTGGTATTAT<br>TGAAACTACTTTCAAGGACGAAACCGAAACCGATTTGTTTGGTGAACAAGCTGTTTTG<br>TGTGGTGGTTGTGTTGAATTGGTTAAGGCTGGTTTTGAAACCTTGGTTGAAGCTGGTTA<br>TGCTCCAGAAATGGCTTATTTTGAATGCTTGCACGAGTTGAAGTTGATCGTTGATTTGAT<br>GTACGAAGGTGGTATTGCCAACATGAACTACTCCATTTCTAACAACGCTGAATACGGTG<br>AATATGTTACTGGTCCAGAAGTTATTAACGCCGAATCAAGAGCTGCTATGAGAAATGCT<br>TTGAAGAGAATCCAAGATGGTGAATACGCCAAGATGTTCATTACTGAAGGTGCTGCTAA<br>TTACCCATCTATGACTGCTTATAGAAGAAACAATGCTGCCCATCCAATCGAACAAATTGG<br>TGAAAAATTGAGAGCTATGATGCCATGGATTGCTGCTAACAAAATCGTTGACAAGTCCA<br>AGAACTAA |
